# Developing and characterising decellularized extracellular matrix hydrogels to bio-fabricate female reproductive tissues

**DOI:** 10.1101/2024.08.24.609513

**Authors:** E. Ribes Martinez, Y. Franko, R. Franko, G.A. Ferronato, A.E.S. Viana, E Windenbach, J. B. Stoeckl, T. Frohlich, M.A.M.M. Ferraz

## Abstract

This study investigated the development and characterization of decellularized extracellular matrix (dECM) hydrogels tailored for the bio-fabrication of female reproductive tissues, specifically targeting cortex, endometrium, medulla, and oviduct tissues. We aimed to evaluate the cytocompatibility, biomechanical properties, and overall efficacy of these dECMs in promoting cell viability, proliferation, and differentiation. Our findings revealed that these dECMs exhibited high biocompatibility with embryo development and cell viability, supporting micro vascularization and cellular differentiation without the need for external growth factors. These hydrogels displayed biomechanical properties that closely mimicked native tissues, which was vital for maintaining their functional integrity and supporting cellular activities. The printability assessments showed that dECMs, particularly those from cortex tissues, achieved high precision in replicating the intended structures, though challenges such as low porosity remained. The bioprinted constructs demonstrated robust cell growth, with over 97% viability observed by day 7, indicating their suitability for cell culture. This work represented a significant advancement in reproductive tissue bio-fabrication, demonstrating the potential of dECM-based hydrogels in creating structurally and functionally viable tissue constructs. By tailoring each dECM to match the unique biomechanical properties of different tissues, we paved the way for more effective and reliable applications in reproductive medicine and tissue engineering.

**Highlights:**

- Developed decellularized extracellular matrix (dECM) bio-inks for bio-fabrication of female reproductive tissues.
- Demonstrated high biocompatibility with embryo development and cell viability.
- Achieved accurate bioprinting, maintaining structural integrity.
- Promoted micro vascularization and cell differentiation without added growth factors.

## 1. Introduction

The female reproductive tract is a complex system composed of different organs that go through extreme remodelling during physiological and pathological processes [1–6]. The coordinated functionality of the ovaries, oviducts, and uterus provides the optimal microenvironment for fertilisation and embryo development. In Addition to the physiological remodelling during the oestrus cycle and pregnancy stages, female reproductive pathologies also promote extracellular and cellular modifications [7]. Current *in vitro* models used to study female reproductive organs are primarily based on 2D cultures, which do not capture the complexity of the native tissue, such as cell polarisation and functionality, and cell-cell interaction [8–12]. Recently, 3D models, such as spheroids and organoids, have been used to recreate the cell-cell and cell-extracellular matrix (ECM) interactions [13–16]. However, such systems lack the full complexity of different cell type interactions and, due to their conformation, need trained personnel and expensive equipment (micromanipulator) to be used for *in vitro* analysis, such as *in vitro* fertilisation (IVF) and secretions collection [17–19]. Organ explants from biopsied tissues have also been used [20–22]. Although tissue explants present different cell types and extracellular matrices, it has short viability before necrosis is observed, which makes it difficult to mimic the endocrine changes of the estrous cycle.

Bioengineering has allowed the successful *in vitro* mimicking of the female reproductive microenvironment, such as the use of microfluidics to recreate an oviduct-on-a-chip and a female menstrual cycle-on-a-chip [23,24,24,25]. However, such systems were limited in the different cell types of crosstalk and lacked a more biomimetic ECM structure. Therefore, there is still a need to recreate models to study these organs in which their multicellular components, morphology, and ECM structure are better bio-mimicked in a dynamic system. In that regard, 3D printing technology has become a powerful tool in the bio-fabrication field [26]. The possibility to reproduce and mimic the complex architecture of tissues allow us to create more representative models of different multifaceted tissues [27,28]. 3D printing allows for regulating the spatial distribution deposition of embedded cells in a bio-ink, modulating the architecture of construct designs and enhancing the cell-ECM interaction, similarly to the native tissue [29,30]. Bio-printing was already used to develop multiple organ models, such as liver, kidney, bone, cartilage, and vasculature [7,31–36].

The range of materials used to bioprint is vast and varies from synthetic polymers, such as polyethylene glycol (PEG) [37,38], poly(epsilon-caprolactone) (PCL) [39,40], pluronic [41], polyglycolic acid (PGA) [42], poly (lactic-co-glycolic acid) (PLGA) [43] and polylactic acid (PLA) [44] to natural polymers, such as alginate [45–47], gelatine [48,49], hyaluronic acid [50,51], fibrin [52,53], collagen [54,55], and matrigel [56,57]. However, polymers should be carefully selected when working with female reproductive tissues due to the potential toxic effects of the polymer on gametes and embryos [58–60]. For instance, methacrylate has been shown to induce fetal malformation, and a reduced rate of embryo cleavage and blastocyst formation [61–63]. Similarly, polyethylene glycol (PEG) has been shown to impair embryo development [64–66]. In that regard, decellularized ECM (dECM) is gaining a significant protagonist role in biofabrication [67]. Due to its complex composition, non-toxicity, and preservation of most of the components from the native tissue, it is an optimal option as bio-ink [68]. dECM was already used for bioprinting several tissues, such as adipose, heart, cartilage, vessel, and placenta, which showed higher viability and gene expression of tissue-specific cells, compared to traditional bio-inks such as alginate, gelatine, or collagen; proving long-term viability and functionality [67,69,70]. However, the use of bioprinting with more biomimetic inks, such as the dECM, for creating female reproductive tissues have not been explored yet.

In this study, our previously characterised bio-mechanical properties of female reproductive tissues (ovary, oviduct, and endometrium) [71] were used as baseline data for the biofabrication of *in vitro* female reproductive models. To biofabricate such models, we developed a protocol to decellularize ovarian, oviductal, and endometrial ECM, which was then used to produce bioinks with rheological properties similar to the tissues of origin. The dECM bioinks were non-toxic to primary cell lines and to *in vitro* produced embryos. The printability of the dECM inks was tested and long-term culture that promoted growth and differentiation of primary epithelial and stromal cells from endometrial and oviductal tissues, as well as cortex stromal cells and spheroids from ovarian tissues within each dECM were investigated.

## 2. Materials and methods

### 2.1 Extracellular matrix decellularization

Female reproductive tracts were collected from the Munich slaughterhouse and carried to the lab for tissue isolation at RT within 2-3 h after collection. The ages of the animals were noted, and the oestrus cycle was determined according to ovarian morphology based on follicle size and corpus luteum (CL) presence, as previously described [72]. Ovaries were dissected, and the ovarian cortex was separated from the medulla to isolate and decellularize both tissues separately. Oviducts were isolated, and surrounding connective tissue was removed. The uterus was opened, and the endometrium was dissected from the two horns. Tissues were stored at - 80°C before starting the process. A total of 5 g of each tissue were homogenised using a blender with 50 mL of sterile ultra-pure water. Cortex, medulla and oviduct were blended for 2 min, and endometrium for 30 s. Contents from the blender were transferred to a 50 mL tube and centrifuge for 5 min at 3,000 rpm and 4°C. The supernatant was removed using a metal strainer with a sterile metal filter to avoid tissue loss. Tissues were then incubated in 500 mL of ultra-pure water in sterile glass bottles and incubated at 4°C in a rotator for 6 h to remove red blood cells. After 6 h, tissues were transferred into a new glass bottle with 4% sodium deoxycholate (SDC; 1 g of tissue per 10 mL of SDC solution). Tissues were incubated at 4°C in a rotator for 24 h, transferred into a new sterile glass bottle with 500 mL of sterile ultra-pure water, and incubated at 4°C in a rotator overnight. Tissues were then transferred to a 50 mL centrifuge tube, filled to 50 mL with ultra-pure water, vortexed for 15 s, and centrifuged for 5 min at 3,000 rpm and 4°C; this wash was repeated up to 20 times. At each wash, a 1 mL aliquot was collected and frozen at -20°C for later quantification of remaining detergent. Tissues were then treated with a DNAse solution (5 μg mL^-1^ in ultra-pure water) for 3 h in rotation at RT, centrifuged for 5 min at 3,000 rpm and 4°C, and the DNase solution was discarded. Two washes were performed, and tissues were frozen at -80°C for at least 1 h before lyophilization. Frozen tissues were frozen-dried overnight using the freeze dryer ZS-10N (Zinscien Scientific Freeze Dryer ZS-N10).

### 2.2 dECM hydrogel formation

The lyophilized dECMs were cut into small pieces and cryo-milled (FRITSCH pulverisette 23). Briefly, cut lyophilized tissues were placed in a 10 cm ball recipient with three 10 mm beads, frozen in liquid nitrogen for 1.5 min and milled for 4 min at a vibration speed of 30 s^-1^. The cryo-milling cycles were repeated 10-15 times until the dECM was fully pulverized. The powder was then dissolved in 0.01 M HCl and 0.1 mg mL^-1^ of pepsin in ultra-pure water in a final concentration of 20 mg mL^-1^ for 72 h under constant rotation at RT, aliquoted and stored at -80°C until use. If bubbles were observed, samples were centrifuged for 5 min at 1,500 rpm at 4°C before aliquoting and storage.

### 2.3 Determination of remaining detergents after dECM washes

Quantification of any remaining SDC in washes was determined by a methylene blue (MB) assay [73]. Briefly, a 0.0125% MB solution was prepared with sterile ultra-pure water. A standard curve was prepared, with following concentrations: 0, 0.125% and 0.25% of SDC in sterile ultra-pure water. Each standard and sample were mixed with 0.0125% MB at a ratio 1:10, vortexed for 30 s, mixed with chloroform at a ratio 1:2, vortexed for 1 min, and incubated for 30 min in the dark to allow for phase separation. Next, 150 μl from the chloroform phase were pipetted in a 96 well plate, and the absorbance of MB-SDC complexes was measured at 630 nm using a plate reader (BMG Labtech), the detergent concentration was extrapolated from the standard curve.

### 2.4 dECM DNA and RNA isolation and quantification

DNA purification was performed using the Genomic DNA Purification Kit (Thermo Scientific) according to the manufacturer’s instructions, using between 30-35 mg of tissue and 200 μl of the hydrogel. Commercial collagen (3 mg/ml, Cellink), was also used for DNA extraction as controls. DNA was quantified in samples using a Qubit (Invitrogen). RNA purification was performed using a Thermo Scientific GeneJET RNA Purification Kit, according to the manufacturer’s instructions, using between 30-35 mg of tissue and 200 μl of the hydrogel. The same controls used for DNA analysis were also used for RNA analysis. Quantification of RNA in samples was performed using a Qubit.

### 2.5 Proteomics of dECM

Decellularized samples were mixed with 8 M urea/ 50 mM ammonium bicarbonate and sonicated using a cup resonator (Sonopuls BR 30, Bandelin) for 5 min 40 sec (10 sec pulse, 20 sec pause) at 4°C. Afterwards, the samples were homogenised using QIAshredders (Qiagen) for 1 min at 21,000 g. The protein concentration was then determined using the Pierce 660 nm assay (Thermo Fisher Scientific). Prior to digestion, samples were reduced in 4 mM dithiothreitol (DTT) and 2 mM tris(2-carboxyethyl) phosphine for 30 min at 56°C and alkylated in 8 mM iodoacetamide in the dark for 30 min at RT. Alkylation was quenched using a final concentration of 10 mM DTT. For the first digestion step, samples were incubated with Lys C (enzyme/substrate: 1/100; Wako, Neuss, Germany) for 4 h at 37°C. The samples were then diluted to a concentration of 1 M urea and underwent a second digestion step with porcine trypsin (enzyme/substrate: 1/50; Promega, Madison, WI, USA) at 37°C for 18 h. After digestion the samples were dried using a vacuum concentrator.

Samples were then dissolved in 0.1% formic acid and aliquots of 1 µg peptides were injected into Ultimate 3000 nano liquid chromatograph coupled with a QExactive HF-X mass spectrometer (both Thermo Fisher Scientific). More specifically, peptides were first trapped on a trap column (PepMap 100 µm × 2 cm, 5 µm particles, Thermo Fisher Scientific) using a flow rate of 5 µL min^-1^ and 0.1% formic acid. Peptides were then eluted using a two-step gradient, first from 5% to 20% of solvent B in 80 min followed by a 9-min increase to 40% solvent B. Solvent A: 0.1% formic acid in water; B: 0.1% formic acid in acetonitrile. The spectra were acquired using data independent acquisition with 50 × 12 m/z-wide isolation windows in the range of 400–1,000 m/z.

The raw mass spectrometry data was searched against the bovine subset of UniProt (Accession date: 24.05.2024) using DIA-NN 1.8.1 [74] in the library-free mode. False discovery rate was controlled to be at 1%. Data analysis was performed using R and Perseus [75]. For quantitative comparisons proteins were filtered for at least 70% valid values in at least one condition and then imputed from a normal distribution. Matrisome proteins were then selected and categorised into Core matrisome (Collagnes, Glycoproteins and Proteoglycans) and Non-core matrisome (affiliated, regulators and secreted factors) using the *Bos Taurus* matrisome dataset (downloaded from “The matrisome project: https://sites.google.com/uic.edu/matrisome/matrisome-annotations/bos-taurus-cow?authuser=0, on June 14th 2024) [76]. Proportions of matrisome proteins from and within different categories in each tissue were calculated by grouping data by matrisome category and summarising the proportion of each category’s protein count relative to the total protein count in the tissue.

### 2.6 Gelation kinetics of dECM

Hydrogels (10 mg mL^-1^ dECM, non-enriched and enriched with 0.5% and 1% alginate) were prepared and neutralised by mixing 11% 10X PBS with sterile 1 M NaOH (target pH 7.4-7.8), 15 mM HEPES and 1X PBS to reach the final volume at 4°C. A total of 100 μl of neutralised hydrogel was added to each well of a 96-well plate on ice in triplicates. Before measurement, the plate was centrifuged for 5 min at 1,200 rpm at 4°C to remove bubbles and incubated for 10 min at RT, to prevent condensation in dishes that can interfere with absorbance measurements. The plate reader (BMG Labtech) was pre-warmed to 38.5°C before use. The turbidity of each well was measured at 405 nm every 2 min for 1 h [77]. Three individual analyses were performed on three batches of each dECM (*n* = 3 per hydrogel type and condition). The normalised absorbance (NA) was calculated using the following formula:

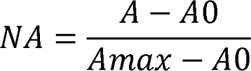

where A is the absorbance to be normalised, A*0* is the absorbance at time 0, and A*max* is the maximum absorbance measured [78].

The kinetic parameters including lag time (TLag; point at which the linear section of the gelation curve intersects with 0% absorbance), time to half gelation (T1/2; moment when absorbance reaches 50%), and time to complete gelation (T1; when absorbance hits 100%), were analysed for the various dECMs groups [78].

### 2.7 Porosity measurements of dECM

Porosity of the crosslinked hydrogels were assessed by liquid displacement assay as previously described [79]. Briefly, volume of the constructs (V) was measured by liquid displacement, gels were then frozen at -80°C for 1 h, and lyophilized overnight. Frozen-dried samples were weighed to determine the dry weight of the hydrogel (M1), immersed in pure ethanol, and incubated overnight at RT. After incubation, constructs were removed from the tubes with ethanol, excess was dried out, and wet weight (M2) was measured. Porosity was determined with the following formula:

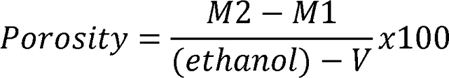

where M1 (g) is the dry weight of the constructs, M2 (g) the wet weight of the constructs, V (cm^3^) is the volume of the dECM constructs, and ⍴ is the density of ethanol (0.798 g cm^-3^).

### 2.8 dECM rheological properties analyses

For initial assessment of hydrogel rheological properties, 20 µL drops of a 10 mg mL^-1^ hydrogel with different alginate supplementation (0, 0.5 and 1%) were prepared, as described above, for all tissues (endometrium, oviduct, cortex and medulla), kept in cell culture media (DMEM/F12 supplemented with 10% FBS and 1% AA) at 38.5°C, 5% CO_2,_ and atmospheric air, and nanoindentation measurements were performed 24h later.

To ensure the stability of hydrogels during nanoindentation, tissue pieces were mounted in a 35mm petri dish by applying a thin layer of a commercial super glue then moving the hydrogel on top of the glue. After a few seconds the hydrogel was covered with culture media (RT) and immediately processed. Hydrogels were indented using a Piuma Chiaro nanoindenter (Optics 11). Indentation depth was set to 5 μm. The probe had a glass spherical tip (tip radius 26 μm) mounted on an individually calibrated cantilever with a spring constant of ∼0.5LJNLJm^−1^. Deformation of the cantilever following contact with the biological sample was measured by tracking the phase-shift in light, reflected from the back of the cantilever. The Young’s modulus (YM) and the effective Young’s modulus (efYM) were calculated using the built-in PIUMA software based on a linear Hertzian contact model build on the first 10% of the force–distance curve.

The elastic and viscoelastic moduli (E’ and E”, respectively) were determined by using the same probe and equipment, using the indentation mode, with frequencies of 1, 5, 10, and 20 Hz, 300 nm amplitude, 1 s slope time, initial relaxation of 10 s, relaxation of 2 s and 5 μm indentation depth. Furthermore, in all experiments, the Piuma attachment mode with a 650 μm distance and 5 s wait was used to prevent probes from starting in contact with hydrogels, a z-above surface of 10 μm, a speed of 80 μm s^-1^ and a threshold of 0.0001 were used in all measurements. All experiments were performed at RT and in culture media to reduce sample attachment.

### 2.9 dECM preparation for embryo culture

Due to the low adherence of dECM hydrogels to the 4-well embryo culture dish, fibronectin (SigmaAldrich) was used to coat the bottom of the plate and promote the adherence of the dECM. Briefly, 25 μL of fibronectin (20 ng uL^-1^) were spreaded in the 4 well plates and incubated for 1 h at 38.5°C. Then, 200 µL of OviECM or EndoECM supplemented with 1% alginate, prepared as previously described, were placed on top, incubated for 45 min at 38.5°C, and 500 µL of warm crosslinking solution supplemented with 0.1 U mL^-1^ Thrombin were carefully added to the top. The dECM plate was incubated for 15 min at 38.5°C, and the crosslinking solution was removed carefully to avoid detachment of the dCEM discs. dECM discs were incubated with ECS media (previously described by Santos et al. [80]) overnight for a washing step and then it was changed to fresh ECS media and incubated overnight again for media equilibration for embryo culture. On the day of *in vitro* maturation (IVM), the coated wells were prepared with oviductal dECM; on the day of *in vitro* fertilisation (IVF), the ECS was changed to fresh ECS media, and the plate was prepared with oil to equilibrate and receive the presumptive zygotes on *in vitro* culture (IVC) day, the next day. On the day of IVC, another plate containing endometrial dECM was prepared. On day 2, the ECS media was changed to fresh ECS media and prepared to transfer the embryos on the next day.

### 2.10 In vitro oocyte maturation (IVM), fertilisation (IVF), and embryo culture (IVC)

Bovine ovaries were obtained from a local slaughterhouse and the follicles between 2-8 mm were aspirated using an 18-gauge needle connected to a vacuum aspiration system. The precipitated pellets containing the cumulus-oocyte complexes (COCs) were transferred to a 9 cm Petri dish and 4 mL of wash medium (TCM 199 - GIBCO, buffered with 2.5% HEPES, supplemented with 10% FBS, 0.2 mM sodium pyruvate, 50 μg mL^-1^ gentamicin sulphate) previously warmed at 38.5°C, was added. Around 200 COCs were selected for each replicate, washed three more times in the same media, and transferred to maturation medium (TCM 199 - GIBCO, buffered with 25 mM sodium bicarbonate, supplemented with 10% FBS, 0.025 mM sodium pyruvate, 2.5 μg mL^-1^ gentamicin sulphate, 0.1 UI mL^-1^ recombinant human FSH) for a last wash before being divided into four wells (30-50 COCs per well) in a pre-equilibrate 4-well plate with 500 µl of maturation media at 38.5°C and 5% CO_2_ in humidified atmospheric air (21% O_2_) for 22 h.

Following, the COCs were washed using fertilisation medium (BO-IVF, IVF Bioscience, UK) and transferred to new pre-equilibrated IVF-wells with 400 µl of fertilisation medium in an incubator at 38.5°C and 5% CO_2_ in humidified atmospheric air (21% O_2_). Frozen bull sperm were thawed at 37°C for 30 s, the content of the straws was added to 2 mL of semen preparation medium (BO-SemenPrep™, IVF Bioscience, UK), centrifuged for 5 min at 330xg, the pellet resuspended with 2 mL of the same media, and again centrifuged (330xg, 5 min). The supernatant was removed, and the remaining pellet was resuspended with 500 μL of fertilisation media. Motility and concentration of thawed sperm were assessed with a mobile computer-assisted sperm analyzer iSperm® mCASA (Aidmics Biotechnology, Taipei, Taiwan). Then, a final concentration of 2.0 x 10^6^ sperm mL^-1^ was added to each of the IVF-wells for incubation with the COCs at 38.5°C, 5% CO_2_, in humidified atmospheric air (21% O_2_) for 18 h.

Before starting the IVC, the 4-well plate was pre-equilibrated overnight in the incubator at 38.5°C, in a humidified atmosphere of 5% O_2_, 5% CO_2_, containing 500 μl of IVC media - ECS and 400 μl of mineral oil (MP BiomedicalsTM) per well with or without oviECM (prepared as described above). The presumptive zygotes were denuded by vortexing in a 15 mL tube with 1 mL of wash medium. Then, the denuded zygotes were washed two more times in the wash medium, one time in the ECS media, transferred to the 4-well plate, and incubated at 38.5°C, in a humidified atmosphere of 5% O_2_, 5% CO_2_. On day 3 post-fertilization, the cleavage rate was checked, and the embryos were transferred to a new pre-equilibrated plate containing the control and endometrium dECM discs. On day 7, the blastocyst rate was checked, and the blastocysts were stained for apoptosis assessment.

To measure the levels of apoptosis, the blastocysts were stained with NucBlue™ Live ReadyProbes™ Reagent (Hoechst 33342; for nuclear staining), and CellEvent™ Caspase 3/7 Detection Reagents in 300 µL of pre-equilibrated ECS media and incubated for 30 min at 38.5°C, 5% O_2_, and 5% CO_2_. The blastocysts were washed once in PBS containing 0.1% polyvinylpyrrolidone and imaged immediately after placing them on a glass slide. Images were taken on an EVOS M7000 (Thermo Fisher Scientific). Each blastocyst was imaged individually with a x20 magnification by taking ZLJstacks with a 2 µm step size. The total number of cells and the apoptotic cells (caspase positive nuclei) were counted, and the apoptosis rate was determined as the percentage of caspase positive cells from the total counted cells.

### 2.11 Stromal and epithelial cell and endometrial gland isolation and culture

#### Endometrial glands and stromal cells

Bovine endometrium from animals older than 45 months at the phases 1 and 2 of the estrus cycle [72] were collected from a local slaughterhouse. Endometrium was dissected and washed with PBS supplemented with 100 Units mL^-1^ Penicillin and 100 μg mL^-1^ Streptomycin (P/S; Gibco, P/S). Pieces of tissue were placed into a sterile 50 mL centrifuge tube with 20 mL of DMEM/F12 enriched with 1% AA (Antibiotic Antimycotic; Corning) and incubated for 20 min in constant shaking at 38.5°C. Pieces were collected onto a petri dish and cut using sterile surgical scissors into 1 x 1 mm. Chopped tissue was placed into a sterile 50 mL centrifuge tube with 20 mL of HBSS supplemented with 0.4 mg mL^-1^ Collagenase V (Sigma, C-9263), 1.25 U mL^-1^ Dispase II (Sigma, D4693), and 1% AA, and incubated for 45-50 minutes in constant shaking at 38.5°C. Every 5 min, the solution was pipetted up and down 60 times to help digestion. Enzymatic reaction was stopped by adding 2 mL of FBS. Subsequently, the content from the tube was filtered using a 100 µm on top of a 70 µm strainer in the middle and a 40 µm strainer at the bottom, in a sterile 50 mL centrifuge tube. For stromal cells collection, the flow through was centrifuged at 1,200 rpm for 10 min at RT. A total of 1x10^6^ cells were seeded in a 100 mm dish and cultured in DMEM/F12 supplemented with 10% FBS and 1% AA at 38.5°C, 5% CO_2,_ and atmospheric air. Media change was performed every other day. The endometrial glands were collected from the 70 µm filter by inverting it towards a sterile 50 mL centrifuge tube and washing the glands out with PBS enriched with 1% AA using a 5 mL syringe with a 21G needle. The sterile 50 mL centrifuge tube was centrifuge at 1,200 rpm for 10 min at RT. The pellet was measured and placed in a 100 mm dish and cultured in DMEM/F12 supplemented with 10% FBS and 1% AA at 38.5°C, 5% CO_2,_ and atmospheric air overnight to posterior experimental procedure.

#### Oviductal epithelial cells

Bovine oviducts from animals older than 45 months at the phases 1 and 2 of the estrus cycle were collected from a local slaughterhouse. Oviduct was isolated, by removing surrounding connective tissue and washed with PBS supplemented with P/S. Tubes were opened and interior epithelial cells scraped with carbon steel surgical blades size 10 (Swann-MortonR), and washed in DMEM/F12 supplemented with 1% AA. Epithelial cells were then placed in a 100 mm dish with DMEM/F12 supplemented with 1% AA and 10% FBS, and cultured at 38.5°C, 5% CO_2,_ and atmospheric air overnight to form oviductal vesicles to be used in experiments.

#### Oviductal stromal cells

Following the collection of oviductal epithelial cells, the remaining tissue was washed three times in PBS with 1% AA. The tissue was then cut into 1x1 mm pieces and 1 g of tissue was transferred to a 20 mL digestion solution, containing 0.4 mg ml^-1^ Collagenase V (Sigma-Aldrich, St. Louis, MO, USA) 1.25 U mL^-1^ Dispase II (Sigma-Aldrich, St. Louis, MO, USA) and 1 % AA in DMEM/F12. This tube was placed in a shaker inside the incubator, and incubated for 40 min, while disrupting the tissue pieces with a cut pipette tip every 10 min. At the end of digestion, 20 mL of culture media (DMEM/F12 with 10% FBS and 1% AA) was added in the solution to stop the digestion. This 40 mL of solution containing the cells was then passed through a 40 µm cell strainer. The flow-through was centrifuged for 5 min at 1,500 rpm, resuspended in culture media and cultured in 100 mm culture dishes. To reduce the epithelial cell contamination, differential plating was performed when the dishes reached 80% confluency. For the differential plating, media was removed, and the culture dish was washed with PBS with 1% AA. 5 mL of 0.25% Trypsin (Corning, Mediatech Inc. Manassas, VA, USA) was added to the plate and incubated for exactly 2 min in the incubator, ensuring that only stromal cells detach. After 2 min, cell suspension was carefully removed, and transferred into a falcon tube with 5 mL of culture media and pelleted, resuspended, replated in 100 mm culture dishes with 10 mL media, and incubated for exactly 5 min, ensuring that only stromal cells attach. After 5 min, the media was carefully removed, and the plate was carefully washed twice with PBS, ensuring that any non-attached cell debris would be washed away. The plate was then supplied with 10 mL of culture media and cultured at 38.5°C, and 5% CO_2_. Differential plating was performed if/when epithelial cell population reached above 15% of cultured cells.

#### Ovarian stromal cells

Cortical and medullar stromal cells were isolated separately from bovine ovaries collected from a local slaughterhouse. The ovaries were transported in Saline solution (0.9% NaCl) supplemented with 100 Units mL^-1^ Penicillin and 100 μg mL^-1^ Streptomycin. After further washes with 70% Ethanol followed with two washes of PBS supplemented with P/S, tissues were isolated. Avoiding corpus luteum and corpus albicans, the cortex was separated from the medulla using a microtome blade and a custom made 3D-printed slicer as shown in [81]. Approximately 2 mm sized cortex and medulla pieces were then collected, placed on a 100 mm cell culture dish, previously scraped for better attachment, and left to dry for 1 h. The tissue pieces were then covered with cell culture media (DMEM/F12 supplemented with 10% FBS, 1% AA and 2.5 μg mL^-1^ gentamicin sulphate) at 38.5°C, 5% CO_2,_ and atmospheric air. The dishes with tissues were cultured until fully confluent, with media change every second day. In case of cross-contamination with epithelial cells, differential plating was performed as described for oviductal stromal cells. For the purpose of the experiments conducted using cortex and medulla stroma cells, only cell lines that had gone through a maximum of 4 passages were used.

### 2.12 dECM hydrogel formation and stroma cell encapsulation

The hydrogel was prepared according to López-Martínez et al. protocol with some modifications [77]. Briefly, 20 mg mL^-1^ dECM ink of EndoECM, CorECM, MedECM, or OviECM was mixed with 11% 10X PBS, 15 mM HEPES, and the pH was neutralised with NaOH 1 M (target pH 7.4-7.8). 1X PBS was added to reach the desired final volume in the case of hydrogels exempt from cells. For preparing hydrogels containing alginate, a 4% alginate stock was used, and alginate was added after pH neutralisation; all procedures were performed on ice. Hydrogels were thoroughly mixed and spun down to remove bubbles. Cells were detached from the culture plate by trypsinization, concentration was determined, and the desired amount of cells was pelleted by centrifugation at 1,200 rpm for 5 min at RT. The pellet was then resuspended in 1X PBS to reach the final volume, and mixed with the neutralised hydrogel reaching a final concentration of 10 mg mL^-1^ dECM and 1x10^6^ cells mL^-1^. 20 or 100 μl drops were prepared. Dishes with drops were incubated at 38.5°C, 5% CO_2_ for 45 min. Then, a pre-warmed and filtered (0.2 μm filter) crosslinking solution (11 mM CaCl_2_, 10 mM HEPES, and 0.1 U mL^-1^ thrombin) was carefully added to the top of the hydrogels and incubated at 38.5°C, 5% CO_2_ for 15 min. After incubation, crosslinking solution was carefully discarded, and crosslinked gels were placed into a 48-well plate with DMEM/F12 media supplemented with 10% FBS and 1% AA and cultured at 38.5°C and 5% CO_2_. Culture media was changed every other day, and hydrogels were analysed for viability and drop area on days 1, 7, and 14. On day 7, gels were also analysed for cell proliferation.

### 2.13 Endometrial glands and stromal cell encapsulation

EndoECM was prepared in a 96-well plate, with a final volume of 100 μl per replicate. 96-well plates were coated as described, and EndoECM was neutralised prior to adding stromal cells with 1X PBS. Stroma cells were prepared and counted, as described before, to obtain a final concentration of 10 mg mL^-1^ dECM and 1x10^6^ cells mL^-1^. Before adding 1X PBS with endometrial stromal cells, glands were collected from a 100 mm dish into a 20 μm cell strainer, 0.2 ul of gland pellet per 1 ul of EndoECM was added in the 1X PBS with stromal cells and gently mixed, then added in the neutralised EndoECM. 1% alginate was added, and 100 μL of EndoECM with cells were pipetted in the coated wells to be crosslinked as previously described. Cells were cultured using ECS media supplemented with 5% FBS and 1% AA. Media were changed every other day. Cells were co-cultured for 7 days, fixed in 4% paraformaldehyde for 1 h at RT and processed for immunofluorescence.

### 2.14 Oviductal epithelial vesicles and stromal cell encapsulation

OviECM was prepared as previously described for embedding oviductal epithelial vesicles and stromal cells together. Oviductal epithelial vesicles were collected after self-formation overnight and passed through a 100 and a 40 μm strainer. Vesicles collection and embedding were performed as described for endometrial glands, but with a 0.05 μl of pellet per 1 μl of ECM ratio. OviECM with oviductal epithelial cells and stromal oviductal cells were kept in culture with DMEM/F12 supplemented with 10% FBS and 1% AA. Media change was performed every other day. Cells were co-cultured for 7 days, fixed in 4% paraformaldehyde for 1 h at RT and processed for immunofluorescence.

### 2.15 Ovarian cortex cell isolation and spheroid culture

Ovarian cortex cells were isolated from cows older than 45 months at the phase 1 and 2 of the estrus cycle. Cortex tissue was dissected from the ovary as previously described and pieces were collected onto a petri dish and cut using sterile surgical scissors into 1 x 1 mm. Similarly to endometrial gland isolation, chopped tissue was placed into a sterile 50 mL centrifuge tube with 20 mL of DMEM/F12 supplemented with 0.4 mg mL^-1^ Collagenase V (Sigma, C-9263), 1.25 U mL^-1^ Dispase II (Sigma, D4693), and 1% AA, and incubated for 40 minutes in constant shaking at 38.5°C. Every 5 min, the solution was pipetted up and down 60 times to help digestion. Enzymatic reaction was stopped by adding 2 mL of FBS. Subsequently, the content from the tube was filtered using a 100 µm on top of a 70 µm strainer in the middle and a 20 µm strainer at the bottom, in a sterile 50 mL centrifuge tube. The flow through was centrifuged at 1,200 rpm for 10 min at RT. A total of 5,000 cells were seeded in a V-shaped bottom 96 well plate previously coated with a non-adherence solution (Stem Cells Technology) according to the manufacturer instruction, to facilitate spheroid formation. Spheroids were cultured in DMEM/F12 supplemented with 10% FBS and 1% AA at 38.5°C, 5% CO_2,_ and atmospheric air for 7 days. At day 7, spheroids were collected and embedded in 10 mg mL^-1^ CorECM enriched with 1% alginate previously neutralised as detailed above. The embedding was performed using a sandwich approach were a layer of 50 µl of neutralised CorECM was added in a fibrinogen coated well from a 96 well plate, the spheroids were deposited, and on top another 50 ul of the same CorECM were added on top. The 96 well plate was incubated at 38.5°C, 5% CO_2_ for 45 min. Then, a pre-warmed and filtered (0.2 μm filter) crosslinking solution (11 mM CaCl_2_, 10 mM HEPES, and 0.1 U mL^-1^ thrombin) was carefully added to the top of the hydrogels and incubated at 38.5°C, 5% CO_2_ for 15 min. After incubation, crosslinking solution was carefully discarded, and crosslinked gels were cultured with DMEM/F12 media supplemented with 10% FBS and 1% AA at 38.5°C and 5% CO_2._ Spheroids were cultured on hydrogel for 7 days, fixed in 4% paraformaldehyde for 1 h at RT and processed for immunofluorescence.

### 2.16 Cell viability and proliferation analysis

For determining cell viability, constructs were washed with phenol red-free DMEMF/12 for 10 min at 38.5°C, incubated for 30 min in phenol red-free DMEMF/12 with 100 nM mL^-1^ Image-iT™ DEAD Green™ viability (Invitrogen) and NucBlue™ Live Cell Stain ReadyProbes™ reagent (Invitrogen; for nuclear staining) according to the manufacturer’s instructions. Constructs were then washed three times in 1x PBS for 10 min each and imaged using an EVOS M7000 fluorescence microscope (Thermo Fisher Scientific). Positive dead cells and total cells were imaged with a x4 magnification by taking ZLJstacks with a 20 µm step size. The total number of cells and the dead cells (DEAD-Green positive nuclei) were counted, and the dead cells rate was determined as the percentage of positive cells from the total counted cells.

Cell proliferation was assessed using a Click-iT® Plus EdU Assay kit according to the manufacturer’s instructions (Invitrogen). Constructs were imaged in phenol red-free media using an EVOS M7000 fluorescence microscope. Positive proliferating cells and total cells were imaged with a x4 magnification by taking ZLJstacks with a 20 µm step size. The total number of cells and the proliferating cells (EdU positive nuclei) were counted, and the proliferation rate was determined as the percentage of positive cells from the total counted cells.

### 2.17 Immunofluorescence

For immunofluorescence, fixed samples were washed with 1X PBS and incubated with a permeabilization solution (1% Triton X-100 in 1X PBS) for 30 min at RT. Following permeabilization, samples were washed with 1X PBS and blocked for 1 hour at RT using a blocking buffer containing 0.1% Triton X-100, 3% BSA, and 5% goat serum in 1X PBS. Primary antibodies were prepared in the blocking buffer at the following concentrations: 1:100 for rabbit anti-VE-cadherin (ab10987564), 1:100 for mouse anti-vimentin (ab8069), 1:300 for rabbit anti-cytokeratin (ab9377), and 1:200 for mouse anti-acetylated alpha-tubulin (ab3201). VE-cadherin was used to stain the microvasculature in ovarian spheroids embedded in CorECM. Mouse anti-vimentin (ab8069) and rabbit anti-cytokeratin (ab9377) were used in combination to stain oviductal epithelial cells and oviductal stromal cells, as well as endometrial stromal cells and epithelial glandular cells. Mouse anti-acetylated alpha-tubulin was used to stain oviductal epithelial cilia. Samples were incubated with primary antibodies overnight at RT with gentle shaking, followed by three washes with 1X PBS, each for 15 min. Secondary antibodies (Alexa Fluor 647 anti-mouse, Alexa Fluor 488 anti-rabbit, and Alexa Fluor 568 anti-rabbit) were prepared in the blocking buffer at a concentration of 1:200. DAPI was also included according to the manufacturer’s instructions. Samples were incubated in the dark with secondary antibodies for 1 h with shaking, followed by three washes with 1X PBS for 15 min each at RT, protected from light.

Post-incubation, tissue clearing was performed by adding CUBIC reagent 1 (4.955 M Urea, 1 M Quadrol (*N*,*N*,*N*′,*N*′-Tetrakis(2-Hydroxypropyl)ethylenediamine), and 29.4% Triton X-100 in water) [82] and incubating overnight, protected from light. The clearing solution was then removed, and samples were washed three times with 1X PBS at RT for 5 minutes each before imaging. Images were captured using a laser scanning confocal microscope (Leica SP8) with x10 and x40 objectives. Z-stacks were acquired with a step size of 2 µm, and the pinhole was set to 0.9. For nuclear staining (DAPI), excitation (Ex) was at 357/44 nm and emission (Em) at 447/60 nm. Alexa Fluor 647 was detected with Ex: 635/18 nm and Em: 692/40 nm, Alexa Fluor 488 with Ex: 482/25 nm and Em: 524/24 nm, and Alexa Fluor 568 with Ex: 585/29 nm and Em: 628/32 nm. Images were processed using ImageJ software, with final presentation after 3D reconstruction based on maximum intensity.

### 2.18 Support bath preparation

A gelatine support bath was prepared as previously described with some modifications [83]. Briefly, 4.5% (m/v) of gelatine was dissolved in 75 mL of sterile ultra-pure water enriched with 11 mM CaCl_2_ in continuous stirring at 60°C in a sterile glass bottle. Once dissolved, the gelatine was incubated at 4°C for 12 h. The gelatine was blended for 120 s in 175 mL of solution A (11 mM CaCl_2_ and 100 mM HEPES in sterile ultra-pure water). The blended gelatine slurry was placed into 50 mL centrifuge tubes and centrifuged for 2 min at 4,200 rpm and 4°C. The supernatant was discarded, and the tubes were refilled with solution A, vortexed for 15 s, and centrifuged again at 4,200 rpm for 2 min at 4°C. This process was repeated at least three times until no foam was observed. Tubes were filled with solution B (solution A enriched with 0.1 U mL^-1^ of Thrombin), vortexed for 15 s, placed on ice for 2 min, the supernatant discarded, and again vortexed for 15 s, placed on ice for 2 min and supernatant discarded. The gelatine slurry was then placed into a 12-well plate and centrifuged for 2 min at 4,200 rpm at 4°C to remove possible bubbles. The support bath was kept on ice until used for bio-printing.

### 2.19 Bio-ink preparation and dECM printing

For printing, a cylinder of 5 mm diameter and 3 mm height was designed using Autodesk Fusion 360. dECM bio-inks (supplemented with 1% alginate) were prepared as described above with or without adding the cells. The hydrogels were then transferred to a 3 mL syringe, locked, and kept on ice until use. The syringes were coupled to the printing cartridge (Cellink), and the hydrogels were thoroughly mixed by passing it between the syringe and the printing cartridge 60 times. The printing cartridge containing the hydrogel was locked and centrifuged at 1,500 rpm for 5 min at 4°C. Meanwhile, the BioX bio-printer (Cellink) was set to cartridge temperature of 20°C and printing bed temperature of 20°C. The cartridge was coupled with a 25G nozzle, and the printing was performed at 14 KPa, 11 mm s^-1^, and 98% infill. After printing, the printed constructs, in the support bath, were incubated at RT for 20 min and placed in a non-humidity controlled incubator at 38.5°C for 45-60 min until the gelatin support bath was completely melted. Constructs were washed twice in DMEM/F12 enriched with 10% FBS and 1% AA. After the incubation, constructs were placed in a 48 well-plate with DMEM/F12 enriched with 10% FBS and 1% AA and immediately used for imaging or cultured at 38.5°C, 5% CO_2_ until further analysis. Constructs were imaged using a 4x magnification to determine diameter and height immediately after printing, and cell viability was evaluated on days 1 and 7 as described above.

### 2.20 Statistical Analysis

All data analysis and visualisation were carried out in R (ver. 4.3.1), and the scripts and packages used for carrying out our analyses are described in supplementary file 1. Visualisations included detailed boxplots with jittered data points overlay to show the distribution and individual variability in DNA/RNA concentration, cleavage rates, blastocyst rates, apoptosis rates, cell proliferation, and porosity across different conditions. Line plots were created to visualise indentation data (E’ and E’’) against frequency for different tissue types, with separate lines and points for each alginate concentration. Scatter plots for diameter and height data from 3D printing experiments highlighted the ranges within predefined limits, calculating and reporting the percentage of samples within these ranges for different tissue types.

For continuous outcomes like RNA and DNA concentration, cleavage rates, apoptosis rates, cell death, proliferation and Young’s modulus, we fitted linear models using the lm() function. The response variables included the measured concentrations or rates, while the predictor variable was the combined factor variable representing the different experimental conditions. Each specific dataset was used as the data source for the respective models. After fitting the linear models, we performed pairwise comparisons using Tukey’s post-hoc tests to identify significant differences among groups. This was accomplished using the glht() function from the multcomp package. The linfct argument within glht() was set to mcp(Comparison = “Tukey”), indicating that Tukey’s method should be applied for the pairwise comparisons. The significance level was set at α = 0.05 for all statistical tests.

For binary outcomes like blastocyst formation, we applied Generalized Linear Models (GLM) with a binomial family to model the data. The glm() function was used, specifying the response variable as the proportion of successful formations relative to the total number of attempts, and the predictor variable as the combined factor variable. Pairwise comparisons for these models were also conducted using Tukey’s HSD method within the glht() function, as described above. In addition, we used non-linear least squares (NLS) to model absorbance data from turbidometry experiments. Logistic growth curves were fitted to predict absorbance over time using the nls() function. Parameters for the logistic model, such as the asymptote (L), growth rate (k), and midpoint (Time_0), were estimated. Predicted absorbance values were then used to calculate gelation times (TLag, THalf, TComplete) by identifying the time points corresponding to specific absorbance thresholds.

## 3. Results

### 3.1 Tissue decellularization and dECM bio-ink validation

In a pilot study we observed inadequate ECM preservation, particularly laminin, when decellularization was performed using SDS and Triton X-100 detergents (data not shown). In this study, we opted to use sodium deoxycholate (SDC) as the sole detergent, as it has demonstrated better preservation of ECM components (Fig. 1a shows an overview of the decellularization and hydrogel formation protocol) [84]. Additionally, thorough removal of residual detergents, genetic material, and cellular debris is crucial for ensuring the functionality of hydrogels [85,86]. To assess the efficacy of DNA and RNA removal in the hydrogels, we conducted extraction procedures on native tissues and 10mg mL^-1^ hydrogels. Our analysis revealed significant removal of DNA and RNA, indicating efficient decellularization. Specifically, RNA removal was 99.7%, 99.7%, 99.6% and 99.8% for endometrium, cortex, medulla and oviduct hydrogels, respectively, compared to respective native tissues (Fig. 1b). Similarly, DNA removal was higher than 99% across the different hydrogel types (Fig. 1c). Furthermore, to validate our findings, a commercial collagen sample was included as a control for DNA and RNA measurements, in which neither RNA nor DNA were detected. We next checked for remaining SDC in the processing washes. No remaining SDC detergent was observed in water after 10 washes for endometrium, medulla, and cortex tissues, while a higher number of washes (15-20) were required for complete removal of SDC from oviduct decellularized tissues (Fig. 1d).

**Figure 1.**
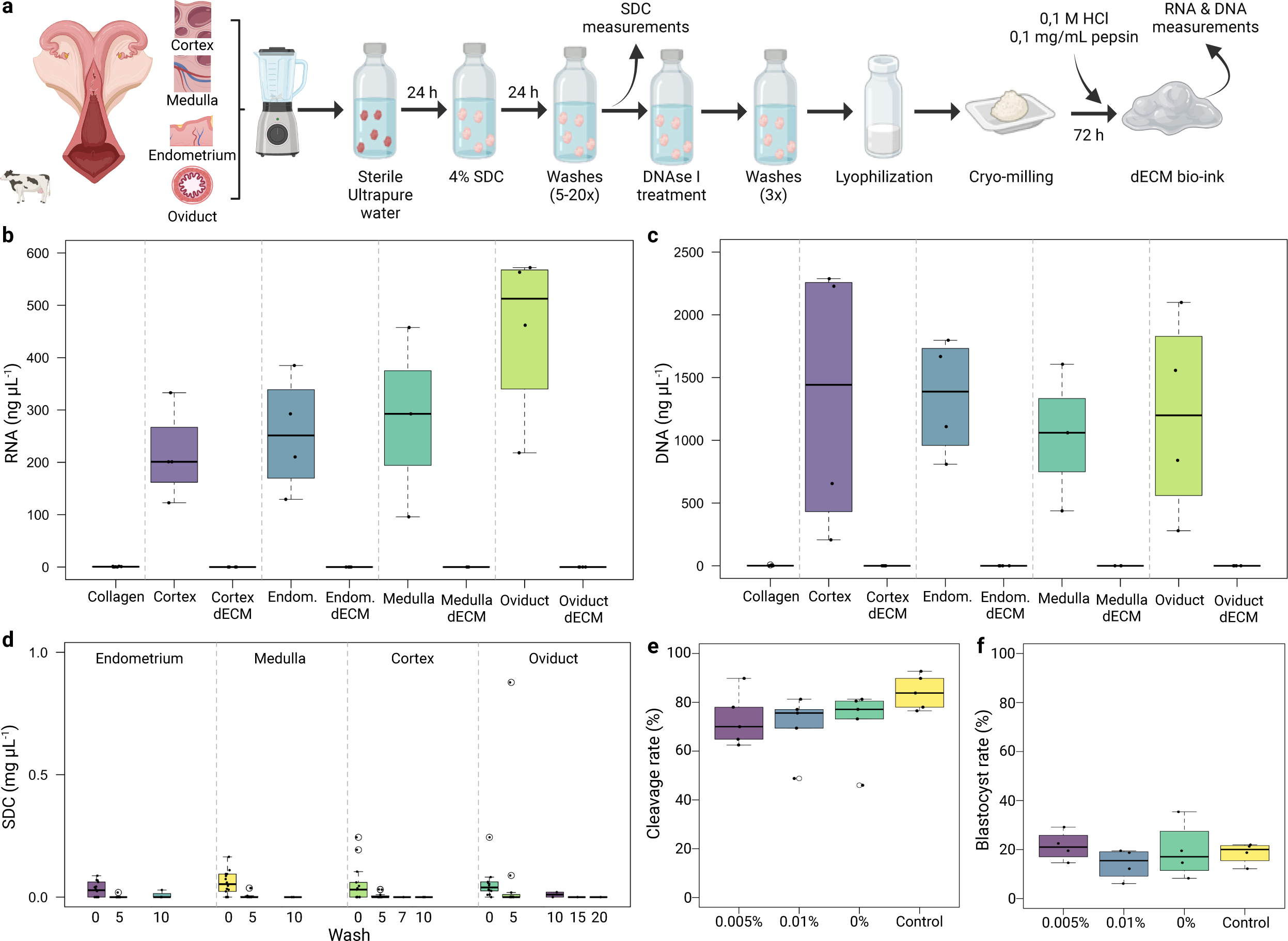
Validation of decellularization protocol and hydrogel formation and characterization. Steps of tissue decellularization and hydrogel formation are shown (a). Decellularization efficiency was validated by RNA (b) and DNA (c) quantification of native tissues and hydrogels. Detection of remaining SDC detergent in water washes was negative after 10 washes for endometrium, medulla and cortex, while 15-20 washes were needed for removal of SDC from oviduct decellularized tissues (d). No effect of hydrogel with or without low amounts of SDC (0, 0.005 and 0.01%, p > 0.05) was observed on embryo cleavage (e) nor blastocyst formation (f). Embryo experiments were performed in 4 replicates, with 30-45 COCs per replicate. Figure was made in Biorender.com.

Assessing embryo sensitivity to biomaterials is crucial in reproductive tract modelling to ensure biocompatibility. To ensure that our dECMs and any residual detergents do not negatively impact embryo development, we investigated the effects of hydrogels and low concentrations of SDC on embryonic outcomes. The presence of low amounts of SDC (0.005% and 0.01%) in hydrogels did not have a significant effect on embryo cleavage or blastocyst formation compared to hydrogels without SDC (Fig. 1e and f). Although no influence on embryo development was observed, the presence of SDC induced cellular apoptosis compared both to the control and the hydrogel without SDC (Suppl. Fig 1). These findings suggest that it is important to detect the presence of SDC in hydrogels previously to use, since low residual levels of SDC can exert a negative effect on embryos.

### 3.2 Proteomics analysis of dECMs bio-inks

We characterised the proteomes of the ECM hydrogels, with a focus on the *Bos taurus* matrisome proteins [76]. For the endometrium, out of a total of 2,182 proteins, 192 matrisome proteins were identified, being 107 core matrisome proteins (collagens, proteoglycans, and glycoproteins) and 85 affiliated proteins (affiliated, regulators and secreted factors; Fig. 2a). A total of 3 matrisome proteins were unique to the endometrium (COL26A1, GDF6 and GDF7, Fig. 2b). For the oviduct, out of 2,790 total proteins, 232 matrisome proteins were identified, being 122 core matrisome proteins and 110 affiliated proteins (Fig. 2a). A total of 13 matrisome proteins were unique to the oviduct (Fig. 2b). Regarding the cortex, out of a total of 1,747 proteins, 210 matrisome proteins were identified, being 119 core matrisome proteins and 91 affiliated proteins (Fig. 2a). A total of 7 matrisome proteins were unique to the cortex (Fig. 1b). Lastly, the medulla, had the highest amount of matrisome proteins detected (250 out of 2,682 total proteins), being 130 core matrisome proteins and 120 affiliated proteins (Fig. 2a), with 14 unique matrisome proteins (Fig. 2b).

**Figure 2.**
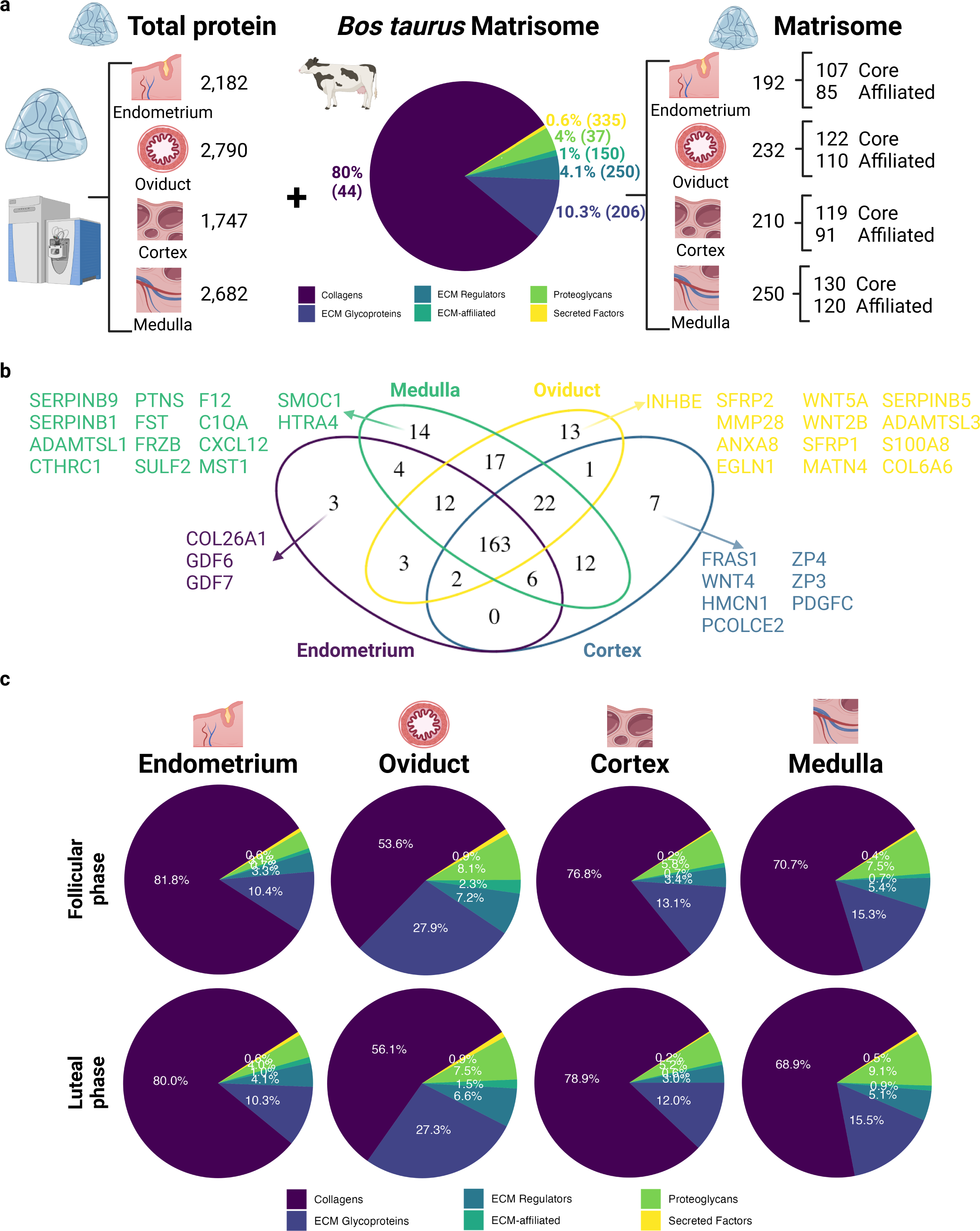
Proteome analysis of endometrium (n=10), oviduct (n=10), cortex (n=11) and medulla (n=10) derived dECMs. Total protein content of each dECM was matched to the Bos taurus matrisome [76], and matrisome components from dECM are shown (a). Venn diagram depicting common and unique matrisome proteins of the different reproductive dECM (b). Estrus stage (follicular and luteal phase) had no effects on dECM matrisome components, which was variable between different tissues dECMS (c). Figure was made in Biorender.com.

We then compared the matrisome composition of hydrogels derived from animals in luteal or follicular phase. Supplementary Table 1 summarises the differentially abundant matrisome proteins for each tissue, being the endometrium and the medulla the tissues that had the highest percentage of differentially abundant matrisome proteins between luteal and follicular phases (3.65 and 2.8%, respectively). When evaluating the proportions of the core and affiliated matrisomes proteins, there were no effects of the estrus stage in any of the tissues (Fig. 2c). Nevertheless, differences between tissues were observed. Specifically, the oviduct was the tissue with the least amount of collagens (54.85, 80.9, 77.8 and 69.8%, for oviduct, endometrium, cortex and medulla, respectively). This reduction in collagens in the oviduct were compensated by an increase in the amount of ECM glycoproteins (27.6, 10.4, 12.5 and 15.4%, for oviduct, endometrium, cortex and medulla, respectively).

Collagens play a pivotal role in bioengineering due to their substantial impact on the mechanical properties of tissues. 12 different collagens were detected in the oviduct (I, II, III, IV, V, VI, XI, XII, XIV, XV, XVI, and XVIII), 14 in the endometrium (I, II, III, IV, V, VI, XI, XII, XIV, XV, XVI, XVIII, XXVI and XXVII), 13 in the cortex (I, III, IV, V, VI, VIII, XI, XII, XIV, XV, XVI, XVIII and XXVII) and 14 in the medulla (I, II, III, IV, V, VI, VIII, XI, XII, XIV, XV, XVI, XVIII and XXVII). Collagen I was the most abundant collagen in all analysed tissues, corresponding to 79.6, 93.9, 89.1 and 86.0% of total collagens for oviduct, endometrium, cortex and medulla, respectively. Collagen VI was the second most abundant collagen in all tissues, corresponding to 13.9, 3.3, 8.7 and 6.1% of total collagens for oviduct, endometrium, cortex and medulla, respectively. As the primary structural proteins in the extracellular matrix, collagens provide crucial tensile strength and elasticity, influencing the overall integrity and function of engineered tissues. Their distribution and abundance are directly instrumental in facilitating cell adhesion, migration, and differentiation, making them indispensable for designing biomaterials and scaffolds [87–90]. In the context of the reproductive tract, understanding the specific types and concentrations of collagen is essential to tailor bio-inks and scaffolds to mimic the unique characteristics of reproductive tissues.

Regarding dECM glycoproteins, 84, 70, 83 and 90 different proteins were identified in the oviduct, endometrium, cortex and medulla, respectively. Dermatopontin (DPT) was the most abundant protein in all tissues, corresponding to 50.3, 36.8, 37.8, and 33.8% of total glycoproteins for oviduct, endometrium, cortex and medulla, respectively. The top five most abundant glycoproteins in the oviduct were DPT, Nidogen 2 (NID2; 6.3%), Nidogen 1 (NID1; 5.2%), Tubulointerstitial nephritis antigen-like 1 (TINAGL1; 4.6%) and Transforming growth factor beta-1 (TGFBI; 2.8%); DPT, Fibrinogen alpha chain (FGA; 7.3%), TINAGL1 (6.6%), Fibrinogen beta chain (FGB; 5.6%) and NID1 (5.2%) for endometrium; DPT, NID1 (8.4%), Cartilage intermediate layer protein 2 (CILP2 (6.9%)), TINAGL1 (6.3%) and Elastin microfibril interfacer 1 (EMILIN1; 4.2%) for cortex; and DPT, TINAGL1 (8.1%), NID1 (6.6%), CILP2 (5.4%) and Microfibril-associated glycoprotein 4 (MFAP4; 4.7%) for medulla. Laminins A, B and C were also detected in all tissues, with their abundance ranging from 0.94 to 1.97% of total glycoproteins. Glycoproteins are essential components of the ECM, contributing significantly to tissue structure and stability by interacting with other matrix elements like collagens and proteoglycans to support the physical architecture of tissues. They play a pivotal role in cell adhesion through specific domains that allow cellular attachment, which is crucial for maintaining tissue integrity and function [91]. Additionally, glycoproteins are involved in critical signalling pathways that influence cellular behaviours such as growth, migration, differentiation, and survival. Glycoproteins also regulate the passage of molecules between tissue compartments and modulate the activity of growth factors and cytokines, affecting their stability and interaction with cellular receptors [91,92]. Their ability to dictate cell migration, shape, and even the cell cycle makes glycoproteins indispensable in tissue engineering, providing essential cues for cell-matrix interactions and facilitating the development of biomimetic materials that replicate the functional architecture of natural tissues.

A total of 15, 13, 12 and 15 proteoglycans were identified in the oviduct, endometrium, cortex and medulla, respectively. Proteoglycans are key constituents of the ECM and play roles in maintaining tissue structure and function. These molecules consist of a core protein linked to glycosaminoglycan (GAG) side chains, which are highly charged and attract water, thereby providing the ECM with hydration and resilience, which enables proteoglycans to contribute significantly to the mechanical properties of tissues, such as compressibility and (visco)elasticity. Beyond their structural roles, proteoglycans are integral to cell signalling processes. They interact with growth factors, cytokines, and chemokines, modulating their availability and activity, thus influencing cell growth, differentiation, and migration [91,92]. Additionally, proteoglycans facilitate cell adhesion by interacting with integrins and other cell surface receptors, supporting cellular communication and the structural assembly of tissue architectures. Given these roles, proteoglycans are vital for designing ECM mimetics in tissue engineering, where recreating the natural biochemical and mechanical environment of tissues is crucial for the successful integration and functionality of engineered tissues. The top five most abundant proteoglycans were Lumican (LUM; 33.7%), Osteoglycin (OGN; 28.6%), Asporin (ASPN; 14.3%), Perlecan (HSPG2; 8.2%) and Prolargin (PRELP; 5.6%) for oviduct; LUM (54.3%), OGN (10.9%), HSPG2 (7.6%), Fibromodulin (FMOD; 7.4%), and Decorin (DCN; 7.1%) for endometrium; Versican (VCAN; 39.8%), LUM (19.7%), FMOD (17.4%), PRELP (6.7%), and OGN (6.4%) for cortex; and LUM (34.4%), FMOD (28.1%), PRELP (9.8%), OGN (7.5%), and ASPN (6.4%) for medulla.

Non-core matrisome proteins, encompassing secreted proteins, affiliated proteins, and ECM regulators, play pivotal roles in the dynamic and regulatory functions of the ECM. Secreted proteins such as thrombospondins, tenascins, and osteopontin primarily modulate cell-matrix interactions, influencing cell adhesion, migration, proliferation, and survival, particularly during tissue repair and in response to environmental stresses [91]. ECM-affiliated regulators, which include a variety of enzymes such as matrix metalloproteinases (MMPs) and their inhibitors (TIMPs), maintain the delicate balance between ECM synthesis and degradation, crucial for tissue homeostasis and adaptation during growth, repair, and pathological conditions [93]. These non-core matrisome components are not merely supportive but are integral to the functional adaptability of the ECM, making them critical targets in the design and development of biomaterials for tissue engineering and regenerative medicine, where recreating a responsive and interactive tissue environment is essential.

Regarding the non-core ECM-affiliated matrisome proteins, 24, 20, 22 and 23 different proteins were identified in the oviduct, endometrium, cortex and medulla, respectively. The top five most abundant ECM-affiliated proteins in the oviduct were Galectin-1 (LGALS1; 41.9%), Annexin A5 (ANXA5; 13.9%), Oviductal Glycoprotein1 (OVGP1; 12.4%), ANXA2 (8.2%) and ANXA1 (6.3%). ANXA2 (36.5%), ANXA1 (13.1%), ANXA5 (11.4%), LGALS1 (6.5%) and ANXA3 (4.7%) for the endometrium.

ANXA5 (32.7%), ANXA2 (15.9%), OVGP1 (5.5%), Tetranectin (CLEC3B; 5.6%) and Hemopexin (HPX; 4.2%) for the cortex. ANXA2 (26%), ANXA5 (14.5%), LGALS1 (10.9%), HPX (7.8%) and CLEC3B; 7.9%) for the medulla.

ECM regulators were presented in the dECM, with 63, 50, 52 and 70 different proteins in the oviduct, endometrium, cortex and medulla, respectively. The top five most abundant ECM regulators in the oviduct were Transglutaminase 2 (TGM2; 76.6%), Collagen-Binding Protein 1 (SERPINH1, 3.9%), Elastase 3B (CELA3B; 2.9%), Alpha-2-macroglobulin (A2M; 1.7%) and Mannan-binding lectin serine protease 2 (MASP; 1.6%). TGM2 (60.2%), SERPINH1 (9.4%), Histidine-rich glycoprotein (HRG; 4.6%), Protein-lysine 6-oxidase (LOX; 3.8%) and Plasminogen (PLG; 3.3%) for the endometrium. TGM2 (59.6%), SERPINH1 (7.7%), LOX (5.5%), CELA3B (2.9%) and Lysyl oxidase homolog 1 (LOXL1; 2.6%) for the cortex. TGM2 (59.9%), LOX (6.1%), SERPINH1 (5.3%), HRG (2.9%) and PLG (2.6%) for the medulla.

Secreted factors were also detected in the dECM with 23, 15, 17 and 27 different secreted factors in the oviduct, endometrium, cortex and medulla, respectively. The top five most abundant ECM regulators in the oviduct were S100 calcium-binding protein A4 (S100A4; 73.7%), S100A11 (10.5%), Protein Wnt-2b (WNT2B; 1.8%), Angiopoietin-related protein 2 (ANGPTL2; 1.7%), and S100A13 (1.1%). S100A4 (68.7%), S100A11 (20.9%), S100A10 (2.1%), ANGPTL2 (1.7%), and Host cell factor 1 (HCFC1; 1.5%) for the endometrium. S100A11 (24.1%), S100A4 (19.7%), Transforming Growth Factor Beta 3 (TGFB3; 11.6%), TGFB2 (9.5%), and ANGPTL2 (4%) for the cortex. S100A4 (42.3%), S100A11 (20.1%), TGFB2 (5.2%), TGFB3 (3.9%), and Pleiotrophin (PTN; 2.6%) for the medulla.

### 3.3 Crosslink ability and biomechanics of dECM bio-inks

To characterize the crosslinking profile in the different dECM bio-inks, a turbidimetric gelation kinetics was performed (Fig. 3), while the calculated kinetic parameters are shown in Suppl. Table 2. All the turbidimetric gelation kinetics, independent of alginate supplementation, showed a sigmoidal shape (Fig. 3 a-d). In general, the lag phase (T_lag_) and the time required to reach half the final turbidity (T_1/2_) were greater in the endometrial dECMs than in the other dECMs (Suppl. Table 2). The presence of alginate in different concentrations did not interfere with the gelation kinetics of the dECMs and the dECMs needed between 32 to 55 min for successful crosslinking (time to complete gelation - T_1_; Suppl. Table 2).

**Figure 3.**
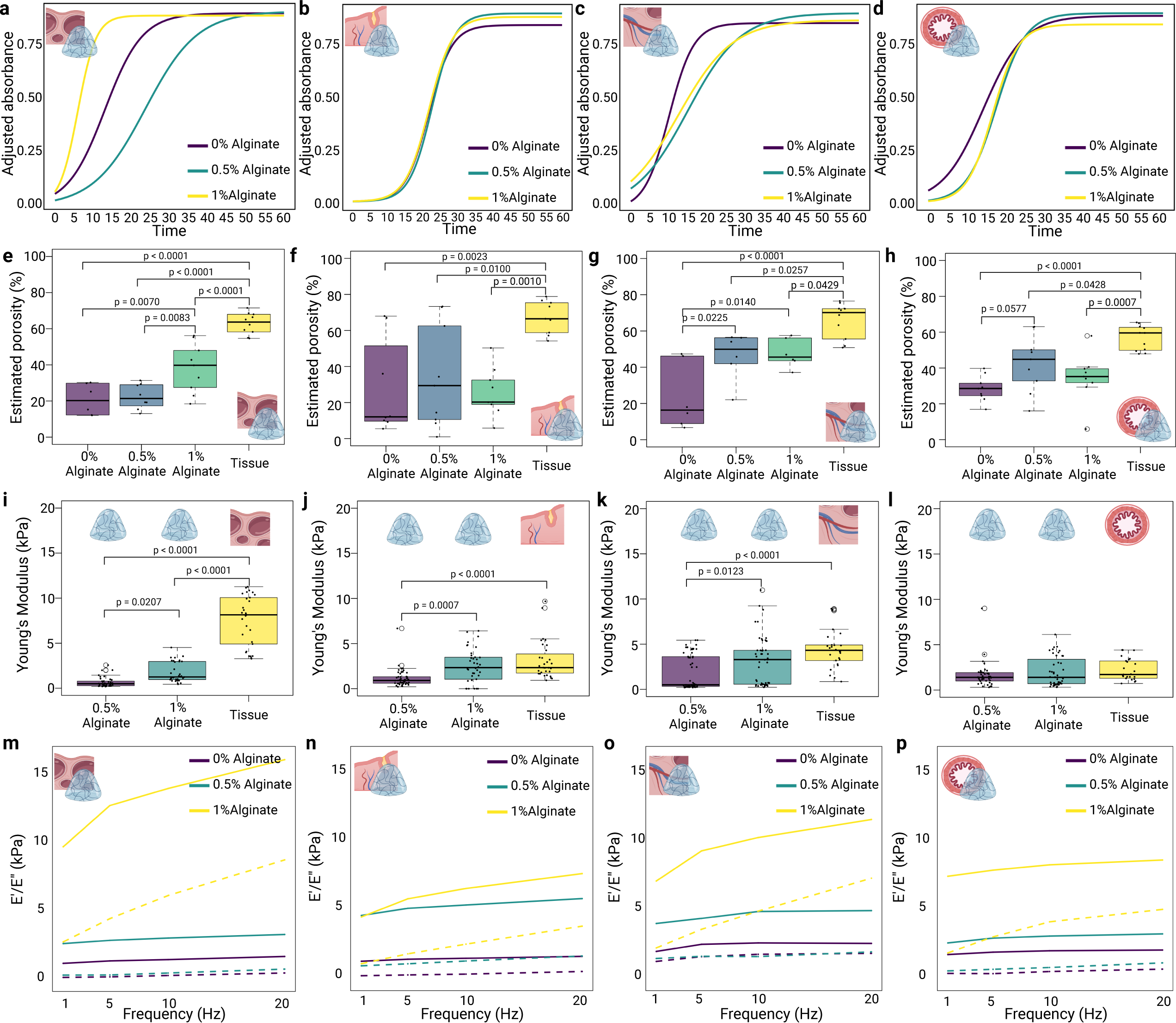
Crosslink ability and mechanical properties of dECM bio-inks supplemented or not with 0.5 and 1% alginate. Gelation kinetic was assessed by turbidimetry assay for CorECM (a), EndoECM (b), OvaECM (c), and OviECM (d). dECM porosity was shown for CorECM (e), EndoECM (f), OvaECM (g), and OviECM (h). Younǵs modulus (KPa) was measured by nanoindentation for CorECM (i), EndoECM (j), OvaECM (k), and OviECM (l). E’/E’’ (Pa) was measured for CorECM (m), EndoECM (n), OvaECM (o), and OviECM (p) by nanoindentation. Native tissues data from our previous work was used for comparison [71]. All experiments were performed in 3 replicates, with 3 hydrogels from each tissue being analysed per replicate. Figure was made in Biorender.com.

The crosslinking process not only determines the mechanical strength, elasticity, and degradation rate of the hydrogels but also affects their porosity Indeed, we observed that all dECMs had reduced estimated porosity when compared to the original tissue, which was also influenced by alginate supplementation (Fig. 3 e-h). Regarding the mechanical properties (stiffness, elasticity, and viscoelasticity), all dECM hydrogels (at a concentration of 10 mg mL^-1^) supplemented with 1% alginate had similar stiffness to the native tissue, with exception of the cortex dECM, for which the Young’s Modulus (YM) was lower than the tissue, independent of the presence of alginate (Fig. 3 i-l). Although the cortex was the stiffest of all tissues analysed, the cortex dECM had similar stiffness to the other dECMs, both with 0.5 and 1% alginate supplementation (p>0.05). This suggests that alginate’s presence could be the key factor influencing hydrogel stiffness, since all hydrogels had the same dECM concentration (10 mg mL^-1^). Indeed, although attempts to measure the YM of the dECM hydrogels without alginate supplementation were partially unsuccessful, due to the hydrogels being more prone to breakage and adherence, the YM values were much lower than the alginate supplemented dECM (Suppl. Fig. 2; p>0.05). All dECMs had similar elastic (E’) and viscoelastic (E”) behaviour, when compared to the native tissues (Figure 3 m-p), demonstrating their suitability for mimicking the female reproductive tract properties.

### 3.4 Cell-ECM interaction: Viability, proliferation, and phenotyping

To assess the cytocompatibility of the dECMs, we initially encapsulated fibroblasts derived from each tissue type examined (cortex, endometrium, medulla, and oviduct) in their respective dECM, supplemented with 0, 0.5, or 1% alginate. Subsequently, we evaluated cell viability on days 1, 7, and 14, and measured cell proliferation on day 7. Supplementing the dECMs with alginate played a crucial role in preventing the matrix from being digested by cells. This is because mammalian cells, which lack the ability to produce alginase, cannot degrade alginate. As seen in Suppl. Fig. 3, the dECMs without alginate supplementation or with 0.5% alginate had reduced area on day 14 in the presence of cells when compared to the same dECM in absence of cells (p<0.05). For the cortex and oviduct dECMs, the use of 1% alginate reduced the degradability of the dECM, which had comparable area to the non-cell dECMs at all time points (p>0.05). At day 14, a significant 70 and 22% reduction in dECMs supplemented with 1% alginate areas were seen for endometrium and medulla, respectively, in the presence of cells, compared to same dECMs in the absence of cells. This reduction was even more significant in the absence of alginate, in which all dECMs had more than 92% area reduction.

The percentage of dead cells in the cortex dECM demonstrated a variation from 0.4 to 5.9%, showing a notable decrease over time, especially in samples supplemented with 0.5% alginate, with significant reductions observed between days 1 and 7, and days 1 and 14 (Fig. 4a). In the case of endometrial cells, there was a decrease in the average percentage of dead cells over the period from days 1 to 14 for both 0% and 0.5% alginate supplements, and from days 1 to 7 and 1 to 14 for the 1% alginate group, where the mean percentage of dead cells ranged from 2.1 to 17.4% (Fig. 4b). For the medulla dECM, no significant change in the percentage of dead cells was observed across different alginate concentrations or culture days, with the mean percentage fluctuating between 1.1 and 7.3% (Fig. 4c). For the oviduct dECM, the average percentage of dead cells varied between 4.9 and 12.2% (Fig. 4d) and, at day 14, dECMs without alginate supplementation had lower cell viability than the ones with alginate (Fig. 4d). Across all groups, the mean viability of cells was above 90% by day 14, indicating the dECM hydrogels’ biocompatibility and non-toxic nature.

**Figure 4.**
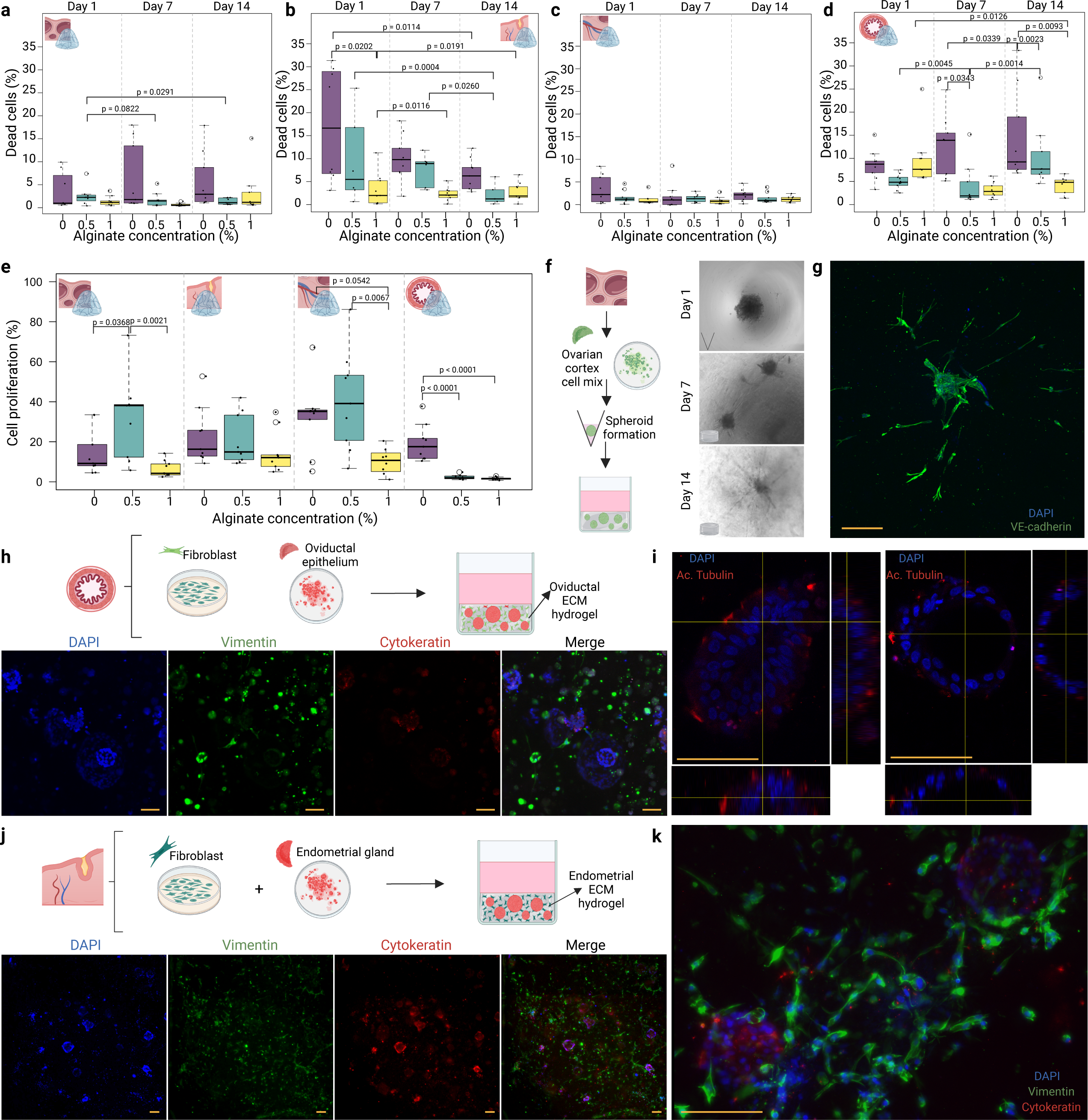
Ability of dECM (enriched or not with 0.5 and 1% alginate) to support cell viability and function. Dead cell percentage was assessed for CorECM (a), EndoECM (b), OvaECM (c), and OviECM (d). Cell proliferation at Day 7 was quantified for the different dECMs (e). Steps of ovarian cortex spheroid formation together with images of day 1 (well), day 7 (well) and day 14 (dECM) of culture is shown in (f). Ovarian cortex spheroids embedded in CorECM developed microvasculature, as seen by VE-cadherin positive staining at Day 14 of culture (g). Oviductal epithelium (cytokeratin + cells) co-cultured with stromal cells (vimentin + cells) in OviECM formed spheroids (non-lumen vesicles) and organoids (presence of lumen) (h). Confocal microscopy images and orthogonal view images of spheroids (left) and organoids (right) depicting ciliation (acetylated alpha tubulin - red) to the outward side (i). Steps of endometrial epithelium glands co-cultured with endometrial stromal cells are shown together with the immunofluorescence imaging of stromal cells (vimentin + cells) and organoids (cytokeratin + cells) co-cultured in EndoECM (j). Confocal microscopy image showing endometrial glands organoids (cytokeratin - red) and stromal cells (vimentin - green) interaction (i). All experiments were performed in 3 replicates, with 3 hydrogels being analysed per replicate per tissue. Scale bars = 50 µm. Figure was made in Biorender.com.

Cell proliferation, a key indicator of cytocompatibility and scaffold efficacy in tissue engineering, was observed across different groups (varying from 1.6 to 40.6%), with notable variations attributed to alginate concentration. Specifically, the 0.5% alginate-supplemented group exhibited higher cell proliferation rates in the cortex dECM compared to both the 0 and 1% alginate groups, and also outperformed the 1% alginate group in the medulla dECM. For the oviduct, the presence of alginate reduced the cell proliferation when compared to no alginate (Fig. 4e). However, in the case of the endometrium dECM, alginate concentration did not significantly affect cell proliferation.

Next, we investigated the ability of the dECM to support ovarian spheroid culture. To that end, we digested the ovarian cortex and cultured the resulting cells in V-shaped wells. Within seven days, the cells formed spheroids which were then transferred to the cortex dECM (supplemented with 1% alginate). In this environment, cell viability exceeded 90%, and the cells began to form microvasculature structures within the dECM hydrogel (Fig. 4f and g).

For the oviduct functionality testing, epithelial cells were isolated and allowed to rearrange into epithelial vesicles for 24 h. These vesicles were then embedded in a co-culture system with stromal oviductal fibroblasts in the oviductal dECM (supplemented with 1% alginate). This co-culture system resulted in the formation of a mixture of spheroids, which lacked internal lumens, and organoids, which possessed internal lumens. The stromal cells were characterised by the expression of vimentin, while the epithelial components of the spheroids and organoids were identified by cytokeratin staining (Fig. 4h). Notably, the polarisation of both spheroids and organoids were directed outward, with ciliation observed on the outer surface (Figure 4i).

Next, we evaluated the endometrial dECM (supplemented with 1% alginate) efficacy in supporting endometrial gland organoid growth in co-culture with endometrial fibroblasts. Using standard embryo culture media [80], we were able to grow endometrial organoids without including the conventional complex growth factor supplementation (Fig. 4 j and k) [16]. Contrasting with other models, especially those reliant on Matrigel that necessitate regular passaging because of hydrogel degradation, our endometrial model demonstrated stability for, at least, 14 days (when experiments were terminated).

### 3.5 Printability and construct integrity

To evaluate the printability of the dECMs, FRESH (Freeform Reversible Embedding of Suspended Hydrogels) 3D printing was applied to print cylinders measuring 3 mm in height and 5 mm in diameter using the four different types of dECMs (Fig. 5a). The size accuracy of these printed constructs was assessed by measuring the height and diameter immediately after printing (Fig. 5b and c). Among the tested dECMs, the cortex and medulla dECM exhibited the highest printing accuracy, with 91.7% of the printed hydrogels varying within ±10% of the intended size and 100% of the cortex constructs within ±15% (Fig. 5d). In contrast, the oviduct dECM demonstrated the lowest accuracy, with only 50% of the printed constructs falling within a ±15% variance from the designed size (Fig. 5d). Despite the successful 3D printing of hydrogels using dECMs, these printed constructs exhibited lower porosity compared to the original tissues (Fig. 5e). This observation aligns with our findings for non-printed dECM constructs (Fig. 3e-h).

**Figure 5.**
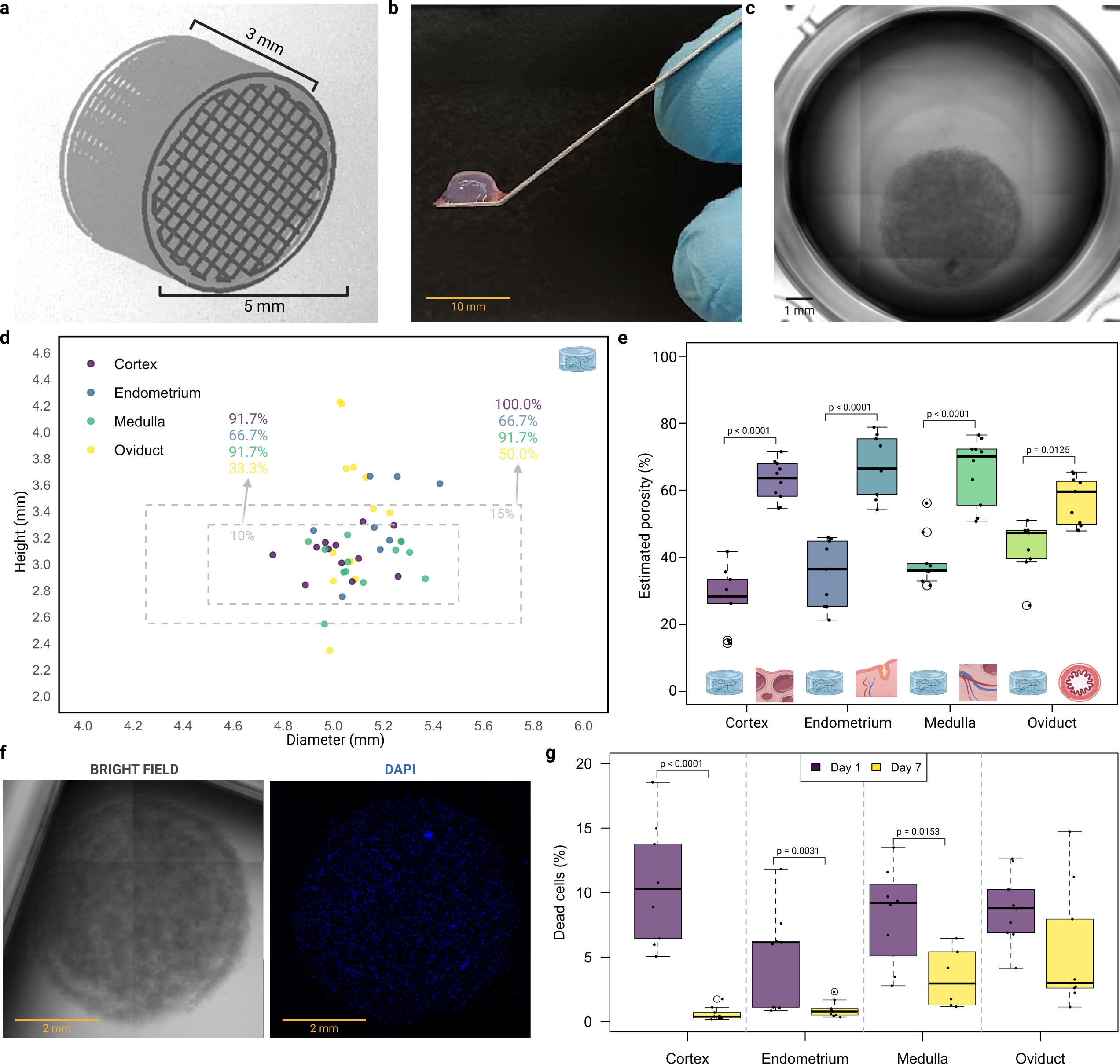
dECM printability, construct integrity, and cell viability. Construct design used to print dECM (a) and image of the printed construct (b). Brightfield image showing the diameter of the printed construct (c). Height and diameter of printed constructs are shown with dashed boxes representing ±10% and ±15% of the designed size (3x5 mm), along with the percentage of each tissue dECM within these variances (d). Estimated porosity of the printed construct at an infill of 99% (e). Image showing the brightfield view of a printed construct with embedded cells in the bio-ink, compared to an immunofluorescence image with DAPI-positive cells on Day 1 (f). Percentage of dead cells from printed constructs embedded in the dECM bio-ink (g). All experiments were performed in 3 replicates, with 3 hydrogels being analysed per replicate per tissue. Figure was made in Biorender.com.

Cell viability was assessed on days 1 and 7 post-printing to evaluate the effects of the printing process on cell health over time. On day 1, the percentage of dead cells was higher, ranging from 5 to 18.5%, indicating that the initial printing process may have caused stress or damage to the cells. By day 7, the percentage of dead cells decreased markedly for endometrium, cortex and medulla, ranging from 0.3 to 6.4% (Fig. 5f and g), while no changes were observed for the oviduct dECM.

## 4. Discussion

This study presented a comprehensive investigation into the development and characterization of dECM hydrogels specifically engineered for the bio-fabrication of female reproductive tissues, including the ovarian cortex and medulla, endometrium, and oviduct. Our findings demonstrated that these dECMs exhibit high biocompatibility, supporting essential cellular processes such as viability, proliferation, and differentiation. Notably, the biomechanical properties of these hydrogels closely mimicked native tissues, underscoring their potential for maintaining functional integrity in bio-fabricated constructs. These results highlight the promising role of dECM-based hydrogels in advancing reproductive tissue engineering and regenerative medicine.

Decellularization is a critical process in tissue engineering, involving the removal of cellular components to create ECM-based scaffolds or hydrogels. Understanding the alterations in bio-mechanical properties post-decellularization is essential for optimising tissue engineering strategies. Notably, detergents like sodium dodecyl sulphate (SDS) and Triton X-100, commonly used in decellularization, can denature collagen and not preserve laminin molecules, potentially influencing cellular behaviour within the ECM hydrogel [84,85]. Additionally, nucleoside and detergent removal is crucial for ensuring the safety and efficacy of following tissue engineering applications and avoid immune response or cytotoxic effects [85,86]. Therefore, an accurate detection and quantification of residual DNA, RNA and SDC are essential to confirm the thoroughness of the washing process and to minimise any potential adverse effects on subsequent biological processes, such as cell proliferation, differentiation, and function. Here, efficient DNA and RNA removal from hydrogels confirmed the effectiveness of the decellularization process and thorough washing minimised residual SDC, crucial for preventing adverse effects in tissue engineering applications.

We then assessed the dECM hydrogel biocompatibility with embryo development. Assessing embryo sensitivity to biomaterials is crucial in reproductive tract modelling to ensure biocompatibility. Previous studies, including our own, have shown that embryos are highly sensitive to their environment, and non-biological polymers such as methacrylate and polyethylene glycol can adversely affect embryo development [58,59,61–64,94]. In here, the dECM had no negative effects on blastocyst formation and quality. Nevertheless, low concentrations of SDC did not affect embryo development, but induced cellular apoptosis, underscoring the importance of meticulous residual checks to ensure biocompatibility in female reproductive tract modelling.

To make sure the decellularization process was preserving the ECM matrisome, we performed proteomics in all dECM hydrogels. The matrisome refers to the ensemble of ECM proteins and associated factors that constitute the structural and functional network surrounding cells in tissues. It plays a crucial role in providing structural support, facilitating cell adhesion, signalling, and regulating cellular functions [92,95,96]. The composition and organisation of the matrisome are critical for tissue development, repair, and homeostasis, influencing processes such as cell migration, differentiation, and response to injury [97,98]. Its significance extends to various fields, including tissue engineering and regenerative medicine, where understanding the matrisome’s interactions and functions can enhance the design of biomaterials and scaffolds for tissue repair and regeneration. The proteomics analysis of the dECM bio-inks revealed significant differences in the composition of core and non-core matrisome proteins among the oviduct, endometrium, cortex, and medulla, showing the distinct structural and functional requirements of each tissue.

Notably, while collagens were abundant across all tissues, their lower concentration in the oviduct (approximately 25% less than in the endometrium and cortex) was counterbalanced by a higher presence of proteoglycans and glycoproteins. The glycoprotein profiles also varied significantly, with the oviduct showing a distinct abundance of DPT, Nidogens, and TGFBI, while cortex and medulla tissues had higher levels of CILP2 and MFAP4. These differences highlight the importance of tailoring bio-inks to replicate the specific biochemical and mechanical environments of each tissue. Moreover, the unique profiles of ECM regulators and secreted factors further emphasize the necessity for precise customization in biomaterial design. For instance, Transglutaminase 2 was overwhelmingly dominant across all tissues, but its relative abundance varied (59 to 76%), suggesting different roles in tissue integrity and remodelling. Additionally, the varied presence of secreted factors like S100 proteins and Transforming Growth Factor Betas across tissues indicates their specific roles in cell signalling and tissue homeostasis. Recognizing these variabilities is crucial for developing biomaterials that can optimally support cell adhesion, migration, differentiation, and overall tissue integrity, ultimately enhancing the effectiveness of bio-fabricated reproductive tissues.

Moreover, the distinct abundance of proteins such as collagens, glycoproteins, and proteoglycans across different tissues informs the choice of crosslinking strategies that can best replicate the native biomechanical environment. In this study, we used a combination of chemical, physical, and enzymatic crosslinking methods tailored to these compositional insights. Crosslinking is a critical process in the formation of hydrogels, especially in the fields of bio-printing and bio-fabrication, where precise control over material properties is essential. The crosslinking process involves the formation of covalent bonds or non-covalent interactions between polymer chains, leading to the transformation of liquid precursors into solid, three-dimensional networks [99]. This process not only determines the mechanical strength, elasticity, and degradation rate of the hydrogels but also affects their porosity, which is crucial for cell infiltration, nutrient diffusion, and waste removal in tissue engineering applications [100]. By carefully modulating temperature (38.5 °C), pH (7.4 - 7.6), and ionic conditions (using CaCl_2_, HEPES, and 10X PBS), and incorporating thrombin, we successfully characterized the gelation profile of the hydrogels, with them reaching crosslinking at around 30 min.

Pore architecture, also play an important role in cell behaviour and cell-ECM interactions [101–103]. Previous studies showed that different pore sizes can impact cell regeneration, proliferation, migration, and adaptation in different scaffolds [49,104–106]. Although dECM bio-inks are more biologically and biophysically similar to the native tissue, it is challenging to preserve good mechanical and structural stability [77,107]. Indeed, none of the dECMs produced here had comparable porosity to the native tissue. Thus, our results suggest that further modifications are needed to obtain more biomimetic porosity.

Other key properties to consider in tissue engineering are tissue stiffness, elasticity, and viscoelasticity. These mechanical characteristics of biological tissues play vital roles in influencing cellular activity and overall tissue function [108]. Consequently, strategies for designing biological tissues must aim to mimic and closely match the mechanical behaviours, encompassing these dynamic aspects of the body’s natural tissues. Most of the produced dECMS had similar stiffness and viscoelastic properties to their native tissue. Nevertheless, changes in the formulation of the cortex dECM bio-ink are needed to bio-fabricate models that better preserve the higher stiffness of the ovarian cortex.

Building on the biomechanical characterization of the dECMs, which demonstrated properties closely resembling native tissues, we explored their application in cell culture environments. A critical consideration in tissue engineering is that while cells need to degrade the dECM to remodel and integrate into the scaffold, excessive degradation can compromise the scaffold’s integrity. Indeed, the degradation of hydrogel construct area has been widely observed in 3D endometrial models, both using dECM [77,84] as well as Matrigel-Collagen [109] hydrogels. Our findings revealed that the inclusion of 1% alginate in all the dECM hydrogels effectively mitigates this degradation, allowing the matrix to maintain its structural stability. This modification is essential for tissue engineering applications, as it enables the matrix to serve as a robust scaffold that supports cell adhesion, proliferation, and differentiation without being excessively broken down by cellular activity.

However, it is important to note that maintaining scaffold integrity with 1% alginate supplementation comes with a trade-off: a reduction in cell proliferation compared to the 0% and 0.5% alginate-enriched dECMs. Despite this decrease in proliferation, cell viability remained unaffected, indicating that the cells were still healthy and functional within the scaffold. This finding highlights the delicate balance required in scaffold composition. While alginate supplementation helps preserve the structural and functional properties of the dECM in cell culture, optimizing the concentration is crucial to support both scaffold integrity and cell proliferation.

Next, we investigated the dECMs ability to support spheroid culture from the different tissues. Cortex spheroids were produced and embedded in CorECM, then cultured in plain culture media without the addition of complex growth factor supplements. This culture environment successfully supported microvasculature formation while preserving high cell viability. The inherent angiogenic properties of the dECM likely played a crucial role in this process. The presence of key angiogenic factors within the dECM, such as angiopoietin 4 (ANGPT4), matrix metalloproteinase 2 (MMP2), peroxidasin (PXDN), and transforming growth factor beta-induced (TGFBI), all of which are known contributors to the angiogenesis pathway, underscores the dECM’s natural ability to promote vascularization without the need for external growth factors.

For the oviduct functionality testing, epithelial vesicles were collected and embedded in a co-culture system with oviductal stromal fibroblasts in the oviductal dECM, supplemented with 1% alginate. This co-culture system resulted in the formation of a mixture of spheroids, which lacked internal lumens, and organoids, which possessed internal lumens. Notably, the polarization of both spheroids and organoids was directed outward, with ciliation observed on their outer surfaces. This is in contrast to other organoid models where polarization and ciliation typically occur internally [110,111]. Importantly, these cultures were successfully maintained in plain media without the need for additional supplements, underscoring the capability of our co-culture system to replicate the physiological conditions of the oviduct and support cellular differentiation and organization.

Lastly, we evaluated the efficacy of the endometrial dECM, supplemented with 1% alginate, in supporting the growth of endometrial gland organoids in co-culture with endometrial fibroblasts. Remarkably, we were able to grow endometrial organoids using standard embryo culture media without the need for conventional complex growth factor supplementation [16,20,109]. Unlike other models, particularly those reliant on Matrigel, which often require regular passaging due to hydrogel degradation, our endometrial model demonstrated stability for at least 14 days, the duration of our experiments. This stability, combined with the dECM’s rich composition and its interaction with endometrial fibroblasts, underscores its ability to sustain organoid development. This approach not only simplifies the culture process but also offers a more physiologically relevant simulation of *in vivo* conditions, representing a significant advancement in refining endometrial models for reproductive biology.

Building on the successful use of dECM to support these various cell co-cultures, the next logical step in advancing these models is to create structures with more biomimetic architecture and complexity. Achieving this level of structural complexity can be effectively accomplished through bio-printing techniques. When considering printability, it’s important to recognize that while dECMs have been successfully used as bio-inks for other organ models [67,69,70], they present unique challenges. These challenges include the need for slow crosslinking (at least 30 minutes) and low viscosity, which complicates direct printing on standard platforms by hindering the maintenance of the designed architecture. This limitation was evident in our initial attempts at direct printing, where the desired structural integrity was not achieved. To overcome these challenges, we employed FRESH printing, which uses a gelatin nanoparticle system supplemented with thrombin to facilitate crosslinking and stabilize the printed constructs [83]. This approach allowed us to better preserve the architecture of the bio-printed structures, enabling the creation of stable tissue models. However, the effectiveness of this approach varied across different tissues. For example, while the cortex and medulla dECMs maintained a stable and well-defined architecture post-printing, the oviductal dECM required further adjustments to the printing parameters and did not achieve similar structural integrity.

Additionally, bio-printing reduced the porosity of the printed construct, resulting in a density lower than that of the native tissue. This reduction in porosity can be attributed to the nature of extrusion printing, where the layer-by-layer deposition of material tends to limit the porosity of the printed hydrogels due to the continuous and dense deposition of hydrogel material. By adjusting the infill percentage—dictating the balance between material and void space within the construct—porosity could be modulated. Alternatively, electrospinning, a versatile technique that involves the use of an electric field to draw very fine fibers from a polymer, could significantly enhance the porosity of hydrogel constructs used in tissue engineering. The process can create fibers with diameters ranging from nanometers to micrometers, resulting in a scaffold with a high surface-area-to-volume ratio and interconnected pore structures [112]. However, the slow crosslinking and low viscosity of dECMs make the use of electrospinning more challenging, as these properties can hinder the formation of stable, fibrous structures necessary for effectively producing these scaffolds.

As part of our validation process for biofabricating female reproductive organs, we observed that cells printed within the constructs initially experienced medium levels of damage after printing. However, by day 7, this damage significantly reduced, indicating that while the initial conditions post-printing were harsher, the cells that survived the initial period adapted and the environment within the constructs became more conducive to cell survival. Looking forward, these advancements highlight the potential of bio-printing for developing more sophisticated models of female reproductive tissues, paving the way for future innovations in tissue engineering and regenerative medicine for reproductive health.

## 5. Conclusions

The reproductive dECMs exhibited biomechanical properties that closely resemble native tissues, which is critical for maintaining functional integrity and supporting cellular processes. Importantly, these bio-inks demonstrated significant biocompatibility with embryo development and cell viability, establishing their potential for use in reproductive applications. Specifically, these dECMs facilitated micro vascularization and cell differentiation without the need for supplemental growth factors or other additives in the media, underscoring their bioactivity and suitability for complex tissue engineering.

Despite the inherent challenges posed by their low viscosity, the use of support baths like the FRESH printing system proved effective in maintaining the structural integrity of the bioprinted constructs. The evaluation of printability revealed that certain dECMs, particularly cortex dECM, achieved high accuracy in replicating intended sizes, although they all exhibited lower porosity compared to original tissues. Adjustments in infill percentages and alternative techniques such as electrospinning offer potential solutions to enhance porosity.

The bioprinted constructs supported cell growth with high viability, indicating a favourable environment for long-term cell survival and scaffold performance. Given the significant differences in the biomechanics of various tissues, it is essential to tailor each dECM individually. Future research should focus on refining these bioprinting processes and material properties to further enhance the functionality and applicability of dECM-based hydrogels in reproductive medicine.

## Availability of data and materials

All data produced and analysed in this project are available within the main manuscript and as supplementary files. The mass spectrometry proteomics data have been deposited to the ProteomeXchange Consortium via the PRIDE [113] partner repository with the dataset identifier PXD053248.

## Supplementary material

Supplementary file 1: All R code to replicate the analysis performed in this study. Supplementary file 2: Suppl. Figures 1-3 and Tables 1 and 2.

Supplementary data 1: dECM matrisome count summary.

## Author’s contributions

Conceptualization: MAMMF, ERM; Methodology: ERM, MAMMF, YF, RF, GF, JBS, ER, TF, AEV; Investigation: all; Supervision: MAMMF, TF; Writing—original draft: ER, MAMMF; Writing—review & editing: all authors.

