## Supplementary material for "Developing and characterising decellularized extracellular matrix hydrogels to bio-fabricate female reproductive tissues": Suppl. File 1

### Supplementary File 1 – R code for all analysis performed

### dECM_all_analysis.R

marciaferraz

2024-06-24

```r
setwd("~/Desktop/R/dECM project")

library(reshape2)
library(nlme)
library(multcomp)

## Loading required package: mvtnorm

## Warning: package 'mvtnorm' was built under R version 4.3.3

## Loading required package: survival

## Warning: package 'survival' was built under R version 4.3.3

## Loading required package: TH.data

## Loading required package: MASS

##
## Attaching package: 'TH.data'

## The following object is masked from 'package:MASS':
##
##     geyser

library(viridis)

## Loading required package: viridisLite

library(ggplot2)
library(gridExtra)
library(survival)
library(survminer)

## Loading required package: ggpubr

##
## Attaching package: 'survminer'

## The following object is masked from 'package:survival':
##
##     myeloma

library(lme4)

## Loading required package: Matrix
```

```
##
## Attaching package: 'lme4'

## The following object is masked from 'package:nlme':
##
##     lmList

library(RColorBrewer)
library(tidyr)

##
## Attaching package: 'tidyr'

## The following objects are masked from 'package:Matrix':
##
##     expand, pack, unpack

## The following object is masked from 'package:reshape2':
##
##     smiths

library(purrr)
library(dplyr)

##
## Attaching package: 'dplyr'

## The following object is masked from 'package:gridExtra':
##
##     combine

## The following object is masked from 'package:MASS':
##
##     select

## The following object is masked from 'package:nlme':
##
##     collapse

## The following objects are masked from 'package:stats':
##
##     filter, lag

## The following objects are masked from 'package:base':
##
##     intersect, setdiff, setequal, union

library(MASS)
library(scales)

##
## Attaching package: 'scales'
```

```
## The following object is masked from 'package:purrr':
##
##     discard

## The following object is masked from 'package:viridis':
##
##     viridis_pal

########## SDC ####################################

#data input
data <- read.csv("SDC_final.csv")
data$batch <- as.factor(data$Batch)
data$wash <- as.factor(data$Wash)

####Basic Summary Statistics


# Means
means <- aggregate(SDC ~Tissue + Wash, data = data, FUN = "mean")

# SDs

sds <- aggregate(SDC ~Tissue + Wash, data = data, FUN = "sd")

SUM <- merge(means, sds, by = c("Tissue"))
names(SUM)[3:4] <- c("Mean", "SD")

write.csv(SUM, file = "Summary_SDC_conc.csv")


##### plot


png(filename="SDC.png",
    width = 8.2, height = 5, units = "in",
    res = 600)


boxplot(  SDC ~ Wash*Tissue,
        data = data,
        cex.axis = 0.9,
        col = viridis(7, alpha = 0.7),
        lwd = 0.7,
        ylab = "",
        ylim = c(0,1),
        axes = FALSE,
        frame.plot = TRUE,
        xlab ="")
```

```
TISSUE <- unique(data$Tissue)
groups <- 4
numbox <- 6
total <- groups * numbox
xpoints <- seq(median(1:numbox),total,numbox)

axis(2, pretty(data$SDC), las = 1, cex.axis = 1)
axis(1, labels = TISSUE, at = xpoints,las = 1, cex.axis = 1)
abline(v = 6.5, lty = "dashed", col = "grey70")
abline(v = 12.5, lty = "dashed", col = "grey70")
abline(v = 18.5, lty = "dashed", col = "grey70")



stripchart(data$SDC ~ data$Wash*data$Tissue,               # Data
           method = "jitter", # Random noise
           pch = 19,          # Pch symbols
           col = "black",           # Color of the symbol
           vertical = TRUE,   # Vertical mode
           add = TRUE,
           cex = 0.25)        # Add it over

title(ylab = expression("SDC (mg/mL)"), line = 2.5)

dev.off()

## quartz_off_screen
##                  2

######RNA & DNA ###################################



#data input
data <- read.csv("RNA_DNA_SDC.csv")
data$Batch <- as.factor(data$Batch)

####     Basic Summary Statistics   DNA
# Means
means <- aggregate(DNA ~Tissue, data = data, FUN = "mean")

# SDs

sds <- aggregate(DNA ~Tissue, data = data, FUN = "sd")

SUM <- merge(means, sds, by = c("Tissue"))
names(SUM)[3:2] <- c("Mean", "SD")
```

```
write.csv(SUM, file = "Summary_DNA_conc.csv")

####     Basic Summary Statistics   RNA
# Means
means2 <- aggregate(RNA ~Tissue, data = data, FUN = "mean")

# SDs

sds2 <- aggregate(RNA ~Tissue, data = data, FUN = "sd")

SUM2 <- merge(means2, sds2, by = c("Tissue"))
names(SUM2)[3:2] <- c("Mean", "SD")

write.csv(SUM2, file = "Summary_RNA_conc.csv")

## plot  DNA concentration

png(filename="dECM_DNA_conc.png",
    width = 8.2, height = 6, units = "in",
    res = 600)

boxplot(DNA ~ Tissue,
        data = data,
        cex.axis = 1,
        col = viridis(9, alpha = 0.7),
        lwd = 0.7,
        ylab = "",
        #ylab = expression(Area ~ (mu * m^2)), # "Area (um2)"
        ylim = c(0,2500),
        axes = FALSE,
        frame.plot = TRUE,
        xlab = "")

TISSUE <- unique(data$Tissue)
groups <- 9
numbox <- 1
total <- groups * numbox
xpoints <- seq(median(1:numbox),total,numbox)

axis(2, pretty(data$DNA), las = 1, cex.axis = 1)
axis(1, labels = TISSUE, at = xpoints,las = 1, cex.axis = 0.7)
abline(v = 1.5, lty = "dashed", col = "grey70")
abline(v = 3.5, lty = "dashed", col = "grey70")
abline(v = 5.5, lty = "dashed", col = "grey70")
abline(v = 7.5, lty = "dashed", col = "grey70")


stripchart(data$DNA ~ data$Tissue,               # Data
           method = "jitter", # Random noise
```

```
            pch = 19,           # Pch symbols
            col = "black",            # Color of the symbol
            vertical = TRUE,    # Vertical mode
            add = TRUE,
            cex = 0.4)         # Add it over

#title("All treatments")
title(ylab = expression("DNA (ng/uL)"), line = 2.7)

dev.off()

## quartz_off_screen
##                 2

## plot  RNA concentration

png(filename="dECM_RNA_conc.png",
    width = 8.2, height = 6, units = "in",
    res = 600)

boxplot(RNA ~ Tissue,
        data = data,
        cex.axis = 1,
        col = viridis(9, alpha = 0.7),
        lwd = 0.7,
        ylab = "",
        #ylab = expression(Area ~ (mu * m^2)), # "Area (um2)"
        ylim = c(0,600),
        axes = FALSE,
        frame.plot = TRUE,
        xlab = "")

TISSUE <- unique(data$Tissue)
groups <- 9
numbox <- 1
total <- groups * numbox
xpoints <- seq(median(1:numbox),total,numbox)

axis(2, pretty(data$RNA), las = 1, cex.axis = 1)
axis(1, labels = TISSUE, at = xpoints,las = 1, cex.axis = 0.7)
abline(v = 1.5, lty = "dashed", col = "grey70")
abline(v = 3.5, lty = "dashed", col = "grey70")
abline(v = 5.5, lty = "dashed", col = "grey70")
abline(v = 7.5, lty = "dashed", col = "grey70")


stripchart(data$RNA ~ data$Tissue,               # Data
           method = "jitter", # Random noise
           pch = 19,           # Pch symbols
           col = "black",            # Color of the symbol
```

```
           vertical = TRUE,    # Vertical mode
           add = TRUE,
           cex = 0.4)         # Add it over

#title("All treatments")
title(ylab = expression("RNA (ng/uL)"), line = 2.7)

dev.off()

## quartz_off_screen
##                  2

############ Embryo bio-compatibility #########################
######Cleavage and blastocyst formation
#data input
data <- read.csv("IVF_final.csv")
data$Replicate <- as.factor(data$Replicate)
data$SDC <- as.factor(data$SDC)

#### Basic Summary Statistics


# Summarize mean and SD by SDC group
summary_data <- data %>%
  group_by(SDC) %>%
  summarize(
    mean_cleavage = mean(Cleavage),
    sd_cleavage = sd(Cleavage),
    mean_blastocyst_D7 = mean(Blasto_D7),
    sd_blastocyst_D7 = sd(Blasto_D7),
    mean_blastocyst_D9 = mean(Blasto_D9),
    sd_blastocyst_D9 = sd(Blasto_D9)
  )


# Save summarized data to a CSV file
write.csv(summary_data, "summary_embryo.csv", row.names = FALSE)


###plot  Cleavage


png(filename="cleavage.png",
    width = 5, height = 6, units = "in",
    res = 600)

boxplot(Cleavage ~ SDC,
        data = data,
        cex.axis = 0.9,
```

```
        col = viridis(4, alpha = 0.7),
        lwd = 0.7,
        ylab = "",
        ylim = c(0,100),
        axes = TRUE,
        frame.plot = TRUE,
        xlab = "")


GROUP <- unique(data$SDC)
groups <- 4
numbox <- 1
total <- groups * numbox
xpoints <- seq(median(1:numbox),total,numbox)

stripchart(data$Cleavage ~ data$SDC,               # Data
           method = "jitter", # Random noise
           pch = 19,          # Pch symbols
           col = "black",           # Color of the symbol
           vertical = TRUE,   # Vertical mode
           add = TRUE,
           cex = 0.5)        # Add it over


title(ylab = expression("Cleavage rate (%)"), line = 1.7)

dev.off()

## quartz_off_screen
##                  2

###plot  Blastocyst all


png(filename="blastocyst_all.png",
    width = 5, height = 6, units = "in",
    res = 600)


boxplot(Blasto_D9 ~ SDC,
        data = data,
        cex.axis = 0.9,
        col = viridis(4, alpha = 0.7),
        lwd = 0.7,
        ylab = "",
        ylim = c(0,100),
        axes = TRUE,
        frame.plot = TRUE,
        xlab = "")
```

```
GROUP <- unique(data$SDC)
groups <- 4
numbox <- 1
total <- groups * numbox
xpoints <- seq(median(1:numbox),total,numbox)


stripchart(data$Blasto_D9 ~ data$SDC,               # Data
           method = "jitter", # Random noise
           pch = 19,          # Pch symbols
           col = "black",           # Color of the symbol
           vertical = TRUE,   # Vertical mode
           add = TRUE,
           cex = 0.4)        # Add it over

title(ylab = expression("Blastocyst rate (%)"), line = 1.5)


dev.off()

## quartz_off_screen
##                 2

####regression model on cleavage

#regression model
test1 <- glm(Cleavage/100 ~ SDC, data = data,
             family = binomial)

## Warning in eval(family$initialize): non-integer #successes in a binomial g
lm!

#Return results
summary(test1)

##
## Call:
## glm(formula = Cleavage/100 ~ SDC, family = binomial, data = data)
##
## Coefficients:
##              Estimate Std. Error z value Pr(>|z|)
## (Intercept)  0.99625    1.00771   0.989    0.323
## SDC0.01SDC  -0.12877    1.40558  -0.092    0.927
## SDC0SDC     -0.07134    1.41390  -0.050    0.960
## SDCControl   0.67393    1.58610   0.425    0.671
##
## (Dispersion parameter for binomial family taken to be 1)
##
```

```
##     Null deviance: 1.4713  on 19  degrees of freedom
## Residual deviance: 1.1309  on 16  degrees of freedom
## AIC: 23.299
##
## Number of Fisher Scoring iterations: 4

###Tukey posthoc
summary(glht(test1, linfct=mcp(SDC = "Tukey")))

##
##   Simultaneous Tests for General Linear Hypotheses
##
## Multiple Comparisons of Means: Tukey Contrasts
##
##
## Fit: glm(formula = Cleavage/100 ~ SDC, family = binomial, data = data)
##
## Linear Hypotheses:
##                         Estimate Std. Error z value Pr(>|z|)
## 0.01SDC - 0.005SDC == 0 -0.12877    1.40558  -0.092    1.000
## 0SDC - 0.005SDC == 0    -0.07134    1.41390  -0.050    1.000
## Control - 0.005SDC == 0  0.67393    1.58610   0.425    0.974
## 0SDC - 0.01SDC == 0      0.05743    1.39421   0.041    1.000
## Control - 0.01SDC == 0   0.80271    1.56857   0.512    0.956
## Control - 0SDC == 0      0.74528    1.57603   0.473    0.965
## (Adjusted p values reported -- single-step method)

#### regression model on Blastocyst ALL

#regression model
test1 <- glm(Blasto_D9/100 ~ SDC, data = data,
             family = binomial)

## Warning in eval(family$initialize): non-integer #successes in a binomial g
lm!

#Return results
summary(test1)

##
## Call:
## glm(formula = Blasto_D9/100 ~ SDC, family = binomial, data = data)
##
## Coefficients:
##             Estimate Std. Error z value Pr(>|z|)
## (Intercept)  -1.2983     1.2182  -1.066    0.287
## SDC0.01SDC   -0.5054     1.8823  -0.269    0.788
## SDC0SDC      -0.1222     1.7548  -0.070    0.944
## SDCControl   -0.1817     1.7718  -0.103    0.918
##
## (Dispersion parameter for binomial family taken to be 1)
```

```
##
##     Null deviance: 0.54370  on 15  degrees of freedom
## Residual deviance: 0.46533  on 12  degrees of freedom
##   (4 observations deleted due to missingness)
## AIC: 14.522
##
## Number of Fisher Scoring iterations: 4
```

```
###Tukey posthoc
summary(glht(test1, linfct=mcp(SDC = "Tukey")))
```

```
##
##   Simultaneous Tests for General Linear Hypotheses
##
## Multiple Comparisons of Means: Tukey Contrasts
##
##
## Fit: glm(formula = Blasto_D9/100 ~ SDC, family = binomial, data = data)
##
## Linear Hypotheses:
##                          Estimate Std. Error z value Pr(>|z|)
## 0.01SDC - 0.005SDC == 0 -0.50540    1.88231  -0.269    0.993
## 0SDC - 0.005SDC == 0    -0.12224    1.75478  -0.070    1.000
## Control - 0.005SDC == 0 -0.18171    1.77177  -0.103    1.000
## 0SDC - 0.01SDC == 0      0.38316    1.91162   0.200    0.997
## Control - 0.01SDC == 0   0.32369    1.92723   0.168    0.998
## Control - 0SDC == 0     -0.05946    1.80289  -0.033    1.000
## (Adjusted p values reported -- single-step method)
```

```
###### Apoptosis and total cell number

#data input
data <- read.csv("IVF_apoptosis_final.csv")
data$Replicate <- as.factor(data$Replicate)
data$SDC <- as.factor(data$SDC)

####     Basic Summary Statistics


# Summarize mean and SD by SDC group
summary_data <- data %>%
  group_by(SDC) %>%
  summarize(
    mean_apoptosis = mean(Apoptosis_rate),
    sd_cleavage = sd(Apoptosis_rate),
    mean_cellnumber = mean(Total_cells),
    sd_cellnumber = sd(Total_cells)
  )
```

```r
# Save summarized data to a CSV file
write.csv(summary_data, "summary_data_IVF_apoptosis.csv", row.names = FALSE)


## plot  Apoptosis


png(filename="IVF_apoptosis.png",
    width = 5, height = 6, units = "in",
    res = 600)

boxplot(Apoptosis_rate ~ SDC,
        data = data,
        cex.axis = 0.9,
        col = viridis(4, alpha = 0.7),
        lwd = 0.7,
        ylab = "",
        ylim = c(0,100),
        axes = TRUE,
        frame.plot = TRUE,
        xlab = "")


GROUP <- unique(data$SDC)
groups <- 4
numbox <- 1
total <- groups * numbox
xpoints <- seq(median(1:numbox),total,numbox)

stripchart(data$Apoptosis_rate ~ data$SDC,               # Data
           method = "jitter", # Random noise
           pch = 19,          # Pch symbols
           col = "black",           # Color of the symbol
           vertical = TRUE,   # Vertical mode
           add = TRUE,
           cex = 0.5)        # Add it over


title(ylab = expression("Apoptotic cells (%)"), line = 1.7)

dev.off()

## quartz_off_screen
##                 2

####regression model on Apoptosis


#regression model
```

```r
test1 <- lm(Apoptosis_rate ~ SDC, data = data)



#Return results
summary(test1)

##
## Call:
## lm(formula = Apoptosis_rate ~ SDC, data = data)
##
## Residuals:
##     Min      1Q  Median      3Q     Max
## -13.548  -6.043  -2.966   5.066  37.804
##
## Coefficients:
##             Estimate Std. Error t value Pr(>|t|)
## (Intercept)  13.5478     1.7825   7.600  1.7e-11 ***
## SDC0.01SDC   -0.3097     2.8049  -0.110  0.91229
## SDC0SDC      -7.5044     2.5875  -2.900  0.00459 **
## SDCControl   -7.8809     2.7313  -2.885  0.00480 **
## ---
## Signif. codes:  0 '***' 0.001 '**' 0.01 '*' 0.05 '.' 0.1 ' ' 1
##
## Residual standard error: 9.925 on 99 degrees of freedom
## Multiple R-squared:  0.131,  Adjusted R-squared:  0.1047
## F-statistic: 4.976 on 3 and 99 DF,  p-value: 0.002942

###Tukey posthoc
summary(glht(test1, linfct=mcp(SDC = "Tukey")))

##
##   Simultaneous Tests for General Linear Hypotheses
##
## Multiple Comparisons of Means: Tukey Contrasts
##
##
## Fit: lm(formula = Apoptosis_rate ~ SDC, data = data)
##
## Linear Hypotheses:
##                         Estimate Std. Error t value Pr(>|t|)
## 0.01SDC - 0.005SDC == 0  -0.3097     2.8049  -0.110   0.9995
## 0SDC - 0.005SDC == 0     -7.5044     2.5875  -2.900   0.0233 *
## Control - 0.005SDC == 0  -7.8809     2.7313  -2.885   0.0242 *
## 0SDC - 0.01SDC == 0      -7.1946     2.8650  -2.511   0.0640 .
## Control - 0.01SDC == 0   -7.5711     2.9955  -2.528   0.0616 .
## Control - 0SDC == 0      -0.3765     2.7929  -0.135   0.9991
## ---
## Signif. codes:  0 '***' 0.001 '**' 0.01 '*' 0.05 '.' 0.1 ' ' 1
## (Adjusted p values reported -- single-step method)
```

```r
## plot  EMbryo total cells


png(filename="IVF_cell_number.png",
    width = 5, height = 6, units = "in",
    res = 600)

boxplot(Total_cells ~ SDC,
        data = data,
        cex.axis = 0.9,
        col = viridis(4, alpha = 0.7),
        lwd = 0.7,
        ylab = "",
        ylim = c(0,140),
        axes = TRUE,
        frame.plot = TRUE,
        xlab = "")


GROUP <- unique(data$SDC)
groups <- 4
numbox <- 1
total <- groups * numbox
xpoints <- seq(median(1:numbox),total,numbox)

stripchart(data$Total_cells ~ data$SDC,               # Data
           method = "jitter", # Random noise
           pch = 19,          # Pch symbols
           col = "black",           # Color of the symbol
           vertical = TRUE,   # Vertical mode
           add = TRUE,
           cex = 0.5)        # Add it over


title(ylab = expression("Total cell number"), line = 1.7)

dev.off()

## quartz_off_screen
##                 2

####regression model on total cell number


#regression model
test1 <- lm(Total_cells ~ SDC, data = data)
```

```r
#Return results
summary(test1)

##
## Call:
## lm(formula = Total_cells ~ SDC, data = data)
##
## Residuals:
##    Min     1Q Median     3Q    Max
## -36.96 -16.76  -2.00  15.00  54.00
##
## Coefficients:
##             Estimate Std. Error t value Pr(>|t|)
## (Intercept)   57.258      4.071  14.065  < 2e-16 ***
## SDC0.01SDC     1.171      6.406   0.183  0.85539
## SDC0SDC       -6.258      5.909  -1.059  0.29218
## SDCControl    19.698      6.238   3.158  0.00211 **
## ---
## Signif. codes:  0 '***' 0.001 '**' 0.01 '*' 0.05 '.' 0.1 ' ' 1
##
## Residual standard error: 22.67 on 99 degrees of freedom
## Multiple R-squared:  0.1526, Adjusted R-squared:  0.127
## F-statistic: 5.944 on 3 and 99 DF,  p-value: 0.0009045

###Tukey posthoc
summary(glht(test1, linfct=mcp(SDC = "Tukey")))

##
##   Simultaneous Tests for General Linear Hypotheses
##
## Multiple Comparisons of Means: Tukey Contrasts
##
##
## Fit: lm(formula = Total_cells ~ SDC, data = data)
##
## Linear Hypotheses:
##                         Estimate Std. Error t value Pr(>|t|)
## 0.01SDC - 0.005SDC == 0    1.171      6.406   0.183   0.9978
## 0SDC - 0.005SDC == 0      -6.258      5.909  -1.059   0.7146
## Control - 0.005SDC == 0   19.698      6.238   3.158   0.0111 *
## 0SDC - 0.01SDC == 0       -7.429      6.543  -1.135   0.6681
## Control - 0.01SDC == 0    18.528      6.841   2.708   0.0390 *
## Control - 0SDC == 0       25.957      6.379   4.069   <0.001 ***
## ---
## Signif. codes:  0 '***' 0.001 '**' 0.01 '*' 0.05 '.' 0.1 ' ' 1
## (Adjusted p values reported -- single-step method)

#######Porosity  All ####################################################



### Cortex
```

```r
#data input
data <- read.csv("Porosity_ctx.csv")


####     Basic Summary Statistics

# Means
means <- aggregate(Porosity ~ Percentage_Alginate, data = data, FUN = "mean")

# SDs

sds <- aggregate(Porosity ~ Percentage_Alginate, data = data, FUN = "sd")

SUM <- merge(means, sds, by = c("Percentage_Alginate"))
names(SUM)[2:3] <- c("Mean", "SD")

write.csv(SUM, file = "Summary_Porosity_ctx.csv")

### Plot

png(filename="Porosity_ctx.png",
    width = 5, height = 5, units = "in",
    res = 600)


boxplot(Porosity ~ Percentage_Alginate,
        data = data,
        cex.axis = 0.9,
        col = viridis(4, alpha = 0.7),
        lwd = 0.7,
        ylab = "",
        ylim = c(0,100),
        axes = TRUE,
        frame.plot = TRUE,
        xlab ="")

stripchart(data$Porosity ~ data$Percentage_Alginate,               # Data
           method = "jitter", # Random noise
           pch = 19,          # Pch symbols
           col = "black",           # Color of the symbol
           vertical = TRUE,   # Vertical mode
           add = TRUE,
           cex = 0.25)        # Add it over

title(ylab = expression("Estimated Porosity (%)"), line = 2.2)


dev.off()
```

```
## quartz_off_screen
##                 2

##Stats

# Fit a general linear model
model <- lm(Porosity ~ Percentage_Alginate, data = data)

# Perform Tukey post-hoc test
tukey_results <- TukeyHSD(aov(model))

# View the results
print(tukey_results)

##   Tukey multiple comparisons of means
##     95% family-wise confidence level
##
## Fit: aov(formula = model)
##
## $Percentage_Alginate
##                 diff        lwr      upr     p adj
## 0.5-0       1.673708 -12.031377 15.37879 0.9870347
## 1-0        17.396000   4.021206 30.77079 0.0070293
## Tissue-0   42.091732  28.987164 55.19630 0.0000000
## 1-0.5      15.722292   3.391341 28.05324 0.0083893
## Tissue-0.5 40.418024  28.380710 52.45534 0.0000000
## Tissue-1   24.695732  13.035857 36.35561 0.0000171

### Medulla
#data input
data <- read.csv("Porosity_med.csv")


####     Basic Summary Statistics

# Means
means <- aggregate(Porosity ~ Percentage_Alginate, data = data, FUN = "mean")

# SDs

sds <- aggregate(Porosity ~ Percentage_Alginate, data = data, FUN = "sd")

SUM <- merge(means, sds, by = c("Percentage_Alginate"))
names(SUM)[2:3] <- c("Mean", "SD")

write.csv(SUM, file = "Summary_Porosity_med.csv")

### Plot
```

```r
png(filename="Porosity_med.png",
    width = 5, height = 5, units = "in",
    res = 600)


boxplot(Porosity ~ Percentage_Alginate,
        data = data,
        cex.axis = 0.9,
        col = viridis(4, alpha = 0.7),
        lwd = 0.7,
        ylab = "",
        ylim = c(0,100),
        axes = TRUE,
        frame.plot = TRUE,
        xlab ="")

stripchart(data$Porosity ~ data$Percentage_Alginate,               # Data
           method = "jitter", # Random noise
           pch = 19,          # Pch symbols
           col = "black",           # Color of the symbol
           vertical = TRUE,   # Vertical mode
           add = TRUE,
           cex = 0.25)        # Add it over

title(ylab = expression("Estimated Porosity (%)"), line = 2.2)


dev.off()

## quartz_off_screen
##                  2

##Stats

# Fit a general linear model
model <- lm(Porosity ~ Percentage_Alginate, data = data)

# Perform Tukey post-hoc test
tukey_results <- TukeyHSD(aov(model))

# View the results
print(tukey_results)

##   Tukey multiple comparisons of means
##     95% family-wise confidence level
##
## Fit: aov(formula = model)
##
## $Percentage_Alginate
##                  diff         lwr       upr      p adj
```

```
## 0.5-0      22.479667   2.6001838 42.35915 0.0225600
## 1-0        23.970833   4.0913505 43.85032 0.0140068
## Tissue-0   42.205929  24.4251787 59.98668 0.0000051
## 1-0.5       1.491167 -18.3883162 21.37065 0.9967780
## Tissue-0.5 19.726262   1.9455120 37.50701 0.0257794
## Tissue-1   18.235095   0.4543453 36.01585 0.0429743

### Oviduct
#data input
data <- read.csv("Porosity_ovi.csv")


####     Basic Summary Statistics

# Means
means <- aggregate(Porosity ~ Percentage_Alginate, data = data, FUN = "mean")

# SDs

sds <- aggregate(Porosity ~ Percentage_Alginate, data = data, FUN = "sd")

SUM <- merge(means, sds, by = c("Percentage_Alginate"))
names(SUM)[2:3] <- c("Mean", "SD")

write.csv(SUM, file = "Summary_Porosity_ovi.csv")

### Plot

png(filename="Porosity_ovi.png",
    width = 5, height = 5, units = "in",
    res = 600)


boxplot(Porosity ~ Percentage_Alginate,
        data = data,
        cex.axis = 0.9,
        col = viridis(4, alpha = 0.7),
        lwd = 0.7,
        ylab = "",
        ylim = c(0,100),
        axes = TRUE,
        frame.plot = TRUE,
        xlab ="")

stripchart(data$Porosity ~ data$Percentage_Alginate,               # Data
           method = "jitter", # Random noise
           pch = 19,          # Pch symbols
           col = "black",           # Color of the symbol
           vertical = TRUE,   # Vertical mode
```

```
           add = TRUE,
           cex = 0.25)        # Add it over

title(ylab = expression("Estimated Porosity (%)"), line = 2.2)


dev.off()

## quartz_off_screen
##                  2

##Stats

# Fit a general linear model
model <- lm(Porosity ~ Percentage_Alginate, data = data)

# Perform Tukey post-hoc test
tukey_results <- TukeyHSD(aov(model))

# View the results
print(tukey_results)

##   Tukey multiple comparisons of means
##     95% family-wise confidence level
##
## Fit: aov(formula = model)
##
## $Percentage_Alginate
##                  diff         lwr       upr     p adj
## 0.5-0      14.139889  -0.3431492 28.622927 0.0577554
## 1-0         6.206000  -8.2770381 20.689038 0.6572511
## Tissue-0   28.291719  14.4826843 42.100754 0.0000200
## 1-0.5      -7.933889 -22.4169270  6.549149 0.4605947
## Tissue-0.5 14.151830   0.3427954 27.960865 0.0428762
## Tissue-1   22.085719   8.2766843 35.894754 0.0007109

### Endo
#data input
data <- read.csv("Porosity_endo.csv")


####     Basic Summary Statistics

# Means
means <- aggregate(Porosity ~ Percentage_Alginate, data = data, FUN = "mean")

# SDs

sds <- aggregate(Porosity ~ Percentage_Alginate, data = data, FUN = "sd")
```

```r
SUM <- merge(means, sds, by = c("Percentage_Alginate"))
names(SUM)[2:3] <- c("Mean", "SD")

write.csv(SUM, file = "Summary_Porosity_endo.csv")

### Plot

png(filename="Porosity_endo.png",
    width = 5, height = 5, units = "in",
    res = 600)


boxplot(Porosity ~ Percentage_Alginate,
        data = data,
        cex.axis = 0.9,
        col = viridis(4, alpha = 0.7),
        lwd = 0.7,
        ylab = "",
        ylim = c(0,100),
        axes = TRUE,
        frame.plot = TRUE,
        xlab ="")

stripchart(data$Porosity ~ data$Percentage_Alginate,               # Data
           method = "jitter", # Random noise
           pch = 19,          # Pch symbols
           col = "black",           # Color of the symbol
           vertical = TRUE,   # Vertical mode
           add = TRUE,
           cex = 0.25)        # Add it over

title(ylab = expression("Estimated Porosity (%)"), line = 2.2)


dev.off()

## quartz_off_screen
##                 2

##Stats

# Fit a general linear model
model <- lm(Porosity ~ Percentage_Alginate, data = data)

# Perform Tukey post-hoc test
tukey_results <- TukeyHSD(aov(model))

# View the results
print(tukey_results)
```

```
##   Tukey multiple comparisons of means
##     95% family-wise confidence level
##
## Fit: aov(formula = model)
##
## $Percentage_Alginate
##                 diff        lwr      upr     p adj
## 0.5-0       6.629736 -20.908726 34.16820 0.9135528
## 1-0        -1.892375 -29.430837 25.64609 0.9976479
## Tissue-0   39.907390  12.368928 67.44585 0.0023621
## 1-0.5      -8.522111 -35.238343 18.19412 0.8223268
## Tissue-0.5 33.277654   6.561422 59.99389 0.0100839
## Tissue-1   41.799765  15.083533 68.51600 0.0010038

#######Porosity  ECM ####################################################

#data input
data <- read.csv("Porosity.csv")

# Create a combined group variable
data$TissueAlginate <- with(data, paste(Tissue, Alginate_concentration, sep =
"_"))


####     Basic Summary Statistics

# Means
means <- aggregate(Porosity ~ Tissue + Alginate_concentration, data = data, F
UN = "mean")

# SDs

sds <- aggregate(Porosity ~ Tissue + Alginate_concentration, data = data, FUN
= "sd")

SUM <- merge(means, sds, by = c("Tissue"))
names(SUM)[3:4] <- c("Mean", "SD")

write.csv(SUM, file = "Summary_porosity.csv")


## plot  Porosity

png(filename="porosity.png",
    width = 10, height = 6.5, units = "in",
    res = 600)
```

```
boxplot(Porosity ~ Tissue + Alginate_concentration,
        data = data,
        cex.axis = 0.9,
        col = viridis(4, alpha = 0.7),
        lwd = 0.7,
        ylab = "",
        ylim = c(0,100),
        axes = TRUE,
        frame.plot = TRUE,
        las = 2,
        xlab = "")


GROUP <- unique(data$Tissue)
groups <- 4
numbox <- 3
total <- groups * numbox
xpoints <- seq(median(1:numbox),total,numbox)

abline(v = 4.5, lty = "dashed", col = "grey70")
abline(v = 8.5, lty = "dashed", col = "grey70")

stripchart(data$Porosity ~ data$Tissue + data$Alginate_concentration,
# Data
           method = "jitter", # Random noise
           pch = 19,          # Pch symbols
           col = "black",           # Color of the symbol
           vertical = TRUE,   # Vertical mode
           add = TRUE,
           cex = 0.4)        # Add it over

#title("All treatments")
title(ylab = expression("Porosity (%)"), line = 1.8)


dev.off()

## quartz_off_screen
##                 2

####### Porosity  Tissue ##########################################

#data input
data <- read.csv("porosity2.csv")


####     Basic Summary Statistics

# Means
```

```
means <- aggregate(Porosity ~ Tissue , data = data, FUN = "mean")

# SDs

sds <- aggregate(Porosity ~ Tissue , data = data, FUN = "sd")

SUM <- merge(means, sds, by = c("Tissue"))
names(SUM)[3:2] <- c("Mean", "SD")

write.csv(SUM, file = "Summary_porosity_tissue.csv")


## plot  Porosity

png(filename="porosity_tissue.png",
    width = 10, height = 6.5, units = "in",
    res = 600)



boxplot(Porosity ~ Tissue,
        data = data,
        cex.axis = 0.9,
        col = viridis(8, alpha = 0.7),
        lwd = 0.7,
        ylab = "",
        ylim = c(0,100),
        axes = TRUE,
        frame.plot = TRUE,
        las = 2,
        xlab = "")


GROUP <- unique(data$Tissue)
groups <- 8
numbox <- 1
total <- groups * numbox
xpoints <- seq(median(1:numbox),total,numbox)

abline(v = 2.5, lty = "dashed", col = "grey70")
abline(v = 4.5, lty = "dashed", col = "grey70")
abline(v = 6.5, lty = "dashed", col = "grey70")


stripchart(data$Porosity ~ data$Tissue,               # Data
           method = "jitter", # Random noise
           pch = 19,          # Pch symbols
           col = "black",           # Color of the symbol
```

```
           vertical = TRUE,    # Vertical mode
           add = TRUE,
           cex = 0.4)          # Add it over


title(ylab = expression("Porosity (%)"), line = 1.8)





dev.off()

## quartz_off_screen
##                 2

########## Turbidometry ############################################

# Read the dataset
data <- read.csv("Turbi.csv")

# Convert variables to appropriate types
data$Time <- as.numeric(data$Time)
data$Abs <- as.numeric(data$Abs)
data$Tissue <- as.factor(data$Tissue)
data$Alginate_concentration <- as.factor(data$Alginate_concentration)

# Create a combined group variable
data$TissueAlginate <- with(data, paste(Tissue, Alginate_concentration, sep =
"_"))

# Get unique combinations of TissueAlginate
selected_tissues <- unique(data$TissueAlginate)

# Initialize an empty data frame with column names
combined_data <- data.frame(Abs = numeric(0), Time = numeric(0), TissueAlgina
te = character(0))

# Loop through each TissueAlginate combination
for (TissueAlginate in selected_tissues) {
  # Fit the model
  FIT <- nls(Abs ~ L / (1 + exp(-k * (Time - Time_0))),
             start = list(L = 1, k = 0.1, Time_0 = 10),
             data = data[which(data$TissueAlginate == TissueAlginate),])

  # Predict values
  PRED <- predict(FIT, newdata = data.frame(Time = seq(0, 60, 0.1)), type = "
response")

  # Create a data frame for the predicted values
```

```
  tissue_pred <- data.frame(Abs = PRED, Time = seq(0, 60, 0.1), TissueAlginat
e = TissueAlginate)

  # Combine with existing data
  combined_data <- rbind(combined_data, tissue_pred)
}

# Create the plot
p <- ggplot(data = combined_data, aes(x = Time, y = Abs, color = TissueAlgina
te)) +
  geom_line(size = 1) +
  ylab(expression("Adjusted absorbance")) +
  scale_color_viridis_d() +
  scale_x_continuous(breaks = seq(0, 60, 5)) +
  labs(y = "Adjusted Absorbance", x = "Time", title = "Adjusted Absorbance fo
r Different Tissues") +
  theme_minimal() +
  theme(panel.grid.major = element_blank(),
        panel.grid.minor = element_blank(),
        panel.border = element_rect(color = "black", fill = NA, linewidth = 1
),
        panel.background = element_rect(fill = NA),
        plot.background = element_rect(fill = NA))

## Warning: Using `size` aesthetic for lines was deprecated in ggplot2 3.4.0.
## ℹ Please use `linewidth` instead.
## This warning is displayed once every 8 hours.
## Call `lifecycle::last_lifecycle_warnings()` to see where this warning was
## generated.

# Save the plot
ggsave(filename = "Adjusted_Absorbance.png", plot = p, width = 15, height = 1
0, units = "cm", dpi = 300)




# Read the dataset
data <- read.csv("Turbi.csv")

# Convert variables to appropriate types
data$Time <- as.numeric(data$Time)
data$Abs <- as.numeric(data$Abs)
data$Tissue <- as.factor(data$Tissue)
data$Alginate_concentration <- as.factor(data$Alginate_concentration)

# Create a combined group variable
data$TissueAlginate <- with(data, paste(Tissue, Alginate_concentration, sep =
"_"))
```

```
# Get unique Tissues
selected_tissues <- unique(data$Tissue)

# Loop through each Tissue
for (Tissue in selected_tissues) {
  # Filter data for the current Tissue
  subset_data <- data[which(data$Tissue == Tissue),]

  # Get unique combinations of TissueAlginate within the current Tissue
  selected_tissue_alginate <- unique(subset_data$TissueAlginate)

  # Initialize an empty data frame with column names for the current Tissue
  combined_data <- data.frame(Abs = numeric(0), Time = numeric(0), TissueAlgi
nate = character(0))

  # Loop through each TissueAlginate combination within the current Tissue
  for (TissueAlginate in selected_tissue_alginate) {
    # Fit the model for the current TissueAlginate
    FIT <- nls(Abs ~ L / (1 + exp(-k * (Time - Time_0))),
               start = list(L = 1, k = 0.1, Time_0 = 10),
               data = subset_data[which(subset_data$TissueAlginate == TissueA
lginate),])

    # Predict values
    PRED <- predict(FIT, newdata = data.frame(Time = seq(0, 60, 0.1)), type =
"response")

    # Create a data frame for the predicted values for the current TissueAlgi
nate
    tissue_pred <- data.frame(Abs = PRED, Time = seq(0, 60, 0.1), TissueAlgin
ate = TissueAlginate)

    # Combine with existing data for the current Tissue
    combined_data <- rbind(combined_data, tissue_pred)
  }

  # Create the plot for the current Tissue
  p <- ggplot(data = combined_data, aes(x = Time, y = Abs, color = TissueAlgi
nate)) +
    geom_line(size = 1) +
    ylab(expression("Adjusted absorbance")) +
    scale_color_viridis_d() +
    scale_x_continuous(breaks = seq(0, 60, 5)) +
    labs(y = "Adjusted Absorbance", x = "Time", title = paste("Adjusted Absor
bance for", Tissue)) +
    theme_minimal() +
    theme(panel.grid.major = element_blank(),
          panel.grid.minor = element_blank(),
```

```
          panel.border = element_rect(color = "black", fill = NA, linewidth =
1),
          panel.background = element_rect(fill = NA),
          plot.background = element_rect(fill = NA))

  # Save the plot with a filename based on Tissue
  filename <- paste0("Adjusted_Absorbance_", gsub(" ", "_", Tissue), ".png")
  ggsave(filename = filename, plot = p, width = 15, height = 10, units = "cm"
, dpi = 300)
}


# Calculate the maximum absorbance for each TissueAlginate group
max_abs <- combined_data %>%
  group_by(TissueAlginate) %>%
  summarize(MaxAbs = max(Abs, na.rm = TRUE), .groups = 'drop')

# Join the max_abs back into combined_data for calculating relative absorbanc
e levels
combined_data <- left_join(combined_data, max_abs, by = "TissueAlginate")

# Function to find nearest time for given absorbance thresholds
find_nearest_time <- function(data, target_abs) {
  abs_diff <- abs(data$Abs - target_abs)
  nearest_index <- which.min(abs_diff)
  return(data$Time[nearest_index])
}

# Initialize a dataframe to store the gelation times for each TissueAlginate
gelation_times <- data.frame(TissueAlginate = character(),
                             TLag = numeric(),
                             THalf = numeric(),
                             TComplete = numeric())

# Loop through each unique TissueAlginate to calculate TLag, T1/2, and T1
for (ta in unique(combined_data$TissueAlginate)) {
  # Filter data for the current TissueAlginate
  ta_data <- combined_data %>% filter(TissueAlginate == ta)

  # Assuming Abs starts at 0 or near 0 for TLag calculation
  # Adjust this line if you have a specific Abs threshold for TLag
  t_lag <- min(ta_data$Time[ta_data$Abs > 0])

  # Calculate T1/2 and T1 using 50% and 100% of the maximum absorbance
  t_half <- find_nearest_time(ta_data, ta_data$MaxAbs[1] * 0.5)
  t_complete <- find_nearest_time(ta_data, ta_data$MaxAbs[1])

  # Append the results to gelation_times
  gelation_times <- rbind(gelation_times, data.frame(TissueAlginate = ta, TLa
```

```
g = t_lag, THalf = t_half, TComplete = t_complete))
}

# View the results
print(gelation_times)

##   TissueAlginate TLag THalf TComplete
## 1       Cortex_0    0  13.1        60
## 2     Cortex_0.5    0  23.6        60
## 3       Cortex_1    0   6.0        60

# save this dataframe to a CSV file
write.csv(gelation_times, "gelation_times.csv", row.names = FALSE)

### stats kinetic

##Tlag

# Read the dataset
data <- read.csv("Kinetic.csv")

# Convert variables to appropriate types
data$TissueAlginate <- as.factor(data$TissueAlginate)


# Fit a general linear model
model <- lm(Tlag ~ TissueAlginate, data = data)

# Perform Tukey post-hoc test
tukey_results <- TukeyHSD(aov(model))

# View the results
print(tukey_results)

##   Tukey multiple comparisons of means
##     95% family-wise confidence level
##
## Fit: aov(formula = model)
##
## $TissueAlginate
##                            diff        lwr        upr     p adj
## Ctx_0.5-Ctx_0    -6.666667e-01  -6.735845  5.4025117 0.9999995
## Ctx_1-Ctx_0       2.220446e-15  -6.069178  6.0691784 1.0000000
## Endo_0-Ctx_0      9.000000e+00   2.930822 15.0691784 0.0008504
## Endo_0.5-Ctx_0    9.333333e+00   3.264155 15.4025117 0.0005273
## Endo_1-Ctx_0      7.333333e+00   1.264155 13.4025117 0.0091158
## Med_0-Ctx_0      -6.666667e-01  -6.735845  5.4025117 0.9999995
## Med_0.5-Ctx_0     2.664535e-15  -6.069178  6.0691784 1.0000000
## Med_1-Ctx_0      -6.666667e-01  -6.735845  5.4025117 0.9999995
## Ovi_0-Ctx_0      -6.666667e-01  -6.735845  5.4025117 0.9999995
```

```
## Ovi_0.5-Ctx_0      2.666667e+00  -3.402512  8.7358451 0.8986902
## Ovi_1-Ctx_0        2.666667e+00  -3.402512  8.7358451 0.8986902
## Ctx_1-Ctx_0.5      6.666667e-01  -5.402512  6.7358451 0.9999995
## Endo_0-Ctx_0.5     9.666667e+00   3.597488 15.7358451 0.0003273
## Endo_0.5-Ctx_0.5   1.000000e+01   3.930822 16.0691784 0.0002035
## Endo_1-Ctx_0.5     8.000000e+00   1.930822 14.0691784 0.0035580
## Med_0-Ctx_0.5     -5.329071e-15  -6.069178  6.0691784 1.0000000
## Med_0.5-Ctx_0.5    6.666667e-01  -5.402512  6.7358451 0.9999995
## Med_1-Ctx_0.5     -3.996803e-15  -6.069178  6.0691784 1.0000000
## Ovi_0-Ctx_0.5     -4.440892e-15  -6.069178  6.0691784 1.0000000
## Ovi_0.5-Ctx_0.5    3.333333e+00  -2.735845  9.4025117 0.7012525
## Ovi_1-Ctx_0.5      3.333333e+00  -2.735845  9.4025117 0.7012525
## Endo_0-Ctx_1       9.000000e+00   2.930822 15.0691784 0.0008504
## Endo_0.5-Ctx_1     9.333333e+00   3.264155 15.4025117 0.0005273
## Endo_1-Ctx_1       7.333333e+00   1.264155 13.4025117 0.0091158
## Med_0-Ctx_1       -6.666667e-01  -6.735845  5.4025117 0.9999995
## Med_0.5-Ctx_1      4.440892e-16  -6.069178  6.0691784 1.0000000
## Med_1-Ctx_1       -6.666667e-01  -6.735845  5.4025117 0.9999995
## Ovi_0-Ctx_1       -6.666667e-01  -6.735845  5.4025117 0.9999995
## Ovi_0.5-Ctx_1      2.666667e+00  -3.402512  8.7358451 0.8986902
## Ovi_1-Ctx_1        2.666667e+00  -3.402512  8.7358451 0.8986902
## Endo_0.5-Endo_0    3.333333e-01  -5.735845  6.4025117 1.0000000
## Endo_1-Endo_0     -1.666667e+00  -7.735845  4.4025117 0.9964985
## Med_0-Endo_0      -9.666667e+00 -15.735845 -3.5974883 0.0003273
## Med_0.5-Endo_0    -9.000000e+00 -15.069178 -2.9308216 0.0008504
## Med_1-Endo_0      -9.666667e+00 -15.735845 -3.5974883 0.0003273
## Ovi_0-Endo_0      -9.666667e+00 -15.735845 -3.5974883 0.0003273
## Ovi_0.5-Endo_0    -6.333333e+00 -12.402512 -0.2641549 0.0354989
## Ovi_1-Endo_0      -6.333333e+00 -12.402512 -0.2641549 0.0354989
## Endo_1-Endo_0.5   -2.000000e+00  -8.069178  4.0691784 0.9849910
## Med_0-Endo_0.5    -1.000000e+01 -16.069178 -3.9308216 0.0002035
## Med_0.5-Endo_0.5  -9.333333e+00 -15.402512 -3.2641549 0.0005273
## Med_1-Endo_0.5    -1.000000e+01 -16.069178 -3.9308216 0.0002035
## Ovi_0-Endo_0.5    -1.000000e+01 -16.069178 -3.9308216 0.0002035
## Ovi_0.5-Endo_0.5  -6.666667e+00 -12.735845 -0.5974883 0.0227807
## Ovi_1-Endo_0.5    -6.666667e+00 -12.735845 -0.5974883 0.0227807
## Med_0-Endo_1      -8.000000e+00 -14.069178 -1.9308216 0.0035580
## Med_0.5-Endo_1    -7.333333e+00 -13.402512 -1.2641549 0.0091158
## Med_1-Endo_1      -8.000000e+00 -14.069178 -1.9308216 0.0035580
## Ovi_0-Endo_1      -8.000000e+00 -14.069178 -1.9308216 0.0035580
## Ovi_0.5-Endo_1    -4.666667e+00 -10.735845  1.4025117 0.2520694
## Ovi_1-Endo_1      -4.666667e+00 -10.735845  1.4025117 0.2520694
## Med_0.5-Med_0      6.666667e-01  -5.402512  6.7358451 0.9999995
## Med_1-Med_0        1.332268e-15  -6.069178  6.0691784 1.0000000
## Ovi_0-Med_0        8.881784e-16  -6.069178  6.0691784 1.0000000
## Ovi_0.5-Med_0      3.333333e+00  -2.735845  9.4025117 0.7012525
## Ovi_1-Med_0        3.333333e+00  -2.735845  9.4025117 0.7012525
## Med_1-Med_0.5     -6.666667e-01  -6.735845  5.4025117 0.9999995
## Ovi_0-Med_0.5     -6.666667e-01  -6.735845  5.4025117 0.9999995
## Ovi_0.5-Med_0.5    2.666667e+00  -3.402512  8.7358451 0.8986902
```

```
## Ovi_1-Med_0.5     2.666667e+00  -3.402512  8.7358451 0.8986902
## Ovi_0-Med_1      -4.440892e-16  -6.069178  6.0691784 1.0000000
## Ovi_0.5-Med_1     3.333333e+00  -2.735845  9.4025117 0.7012525
## Ovi_1-Med_1       3.333333e+00  -2.735845  9.4025117 0.7012525
## Ovi_0.5-Ovi_0     3.333333e+00  -2.735845  9.4025117 0.7012525
## Ovi_1-Ovi_0       3.333333e+00  -2.735845  9.4025117 0.7012525
## Ovi_1-Ovi_0.5     2.664535e-15  -6.069178  6.0691784 1.0000000
```

```
##T1/2


# Fit a general linear model
model <- lm(t1.2 ~ TissueAlginate, data = data)

# Perform Tukey post-hoc test
tukey_results <- TukeyHSD(aov(model))

# View the results
print(tukey_results)
```

```
##   Tukey multiple comparisons of means
##     95% family-wise confidence level
##
## Fit: aov(formula = model)
##
## $TissueAlginate
##                            diff        lwr         upr     p adj
## Ctx_0.5-Ctx_0     1.000000e+01  -2.027563 22.02756259 0.1680874
## Ctx_1-Ctx_0      -7.666667e+00 -21.113890  5.78055711 0.6497617
## Endo_0-Ctx_0      4.000000e+00  -8.027563 16.02756259 0.9831766
## Endo_0.5-Ctx_0    7.000000e+00  -5.027563 19.02756259 0.6230311
## Endo_1-Ctx_0      7.000000e+00  -5.027563 19.02756259 0.6230311
## Med_0-Ctx_0      -3.666667e+00 -15.694229  8.36089593 0.9913533
## Med_0.5-Ctx_0    -3.333333e-01 -12.360896 11.69422926 1.0000000
## Med_1-Ctx_0      -1.333333e+00 -13.360896 10.69422926 0.9999994
## Ovi_0-Ctx_0      -2.000000e+00 -14.027563 10.02756259 0.9999643
## Ovi_0.5-Ctx_0     1.000000e+00 -11.027563 13.02756259 1.0000000
## Ovi_1-Ctx_0       2.000000e+00 -10.027563 14.02756259 0.9999643
## Ctx_1-Ctx_0.5    -1.766667e+01 -31.113890 -4.21944289 0.0038329
## Endo_0-Ctx_0.5   -6.000000e+00 -18.027563  6.02756259 0.7991836
## Endo_0.5-Ctx_0.5 -3.000000e+00 -15.027563  9.02756259 0.9983795
## Endo_1-Ctx_0.5   -3.000000e+00 -15.027563  9.02756259 0.9983795
## Med_0-Ctx_0.5    -1.366667e+01 -25.694229 -1.63910407 0.0169011
## Med_0.5-Ctx_0.5  -1.033333e+01 -22.360896  1.69422926 0.1395750
## Med_1-Ctx_0.5    -1.133333e+01 -23.360896  0.69422926 0.0772635
## Ovi_0-Ctx_0.5    -1.200000e+01 -24.027563  0.02756259 0.0508880
## Ovi_0.5-Ctx_0.5  -9.000000e+00 -21.027563  3.02756259 0.2817048
## Ovi_1-Ctx_0.5    -8.000000e+00 -20.027563  4.02756259 0.4383026
## Endo_0-Ctx_1      1.166667e+01  -1.780557 25.11389045 0.1317238
## Endo_0.5-Ctx_1    1.466667e+01   1.219443 28.11389045 0.0244722
```

```
## Endo_1-Ctx_1       1.466667e+01   1.219443 28.11389045 0.0244722
## Med_0-Ctx_1        4.000000e+00  -9.447224 17.44722378 0.9928852
## Med_0.5-Ctx_1      7.333333e+00  -6.113890 20.78055711 0.7044749
## Med_1-Ctx_1        6.333333e+00  -7.113890 19.78055711 0.8484573
## Ovi_0-Ctx_1        5.666667e+00  -7.780557 19.11389045 0.9179709
## Ovi_0.5-Ctx_1      8.666667e+00  -4.780557 22.11389045 0.4826312
## Ovi_1-Ctx_1        9.666667e+00  -3.780557 23.11389045 0.3324971
## Endo_0.5-Endo_0    3.000000e+00  -9.027563 15.02756259 0.9983795
## Endo_1-Endo_0      3.000000e+00  -9.027563 15.02756259 0.9983795
## Med_0-Endo_0      -7.666667e+00 -19.694229  4.36089593 0.4981602
## Med_0.5-Endo_0    -4.333333e+00 -16.360896  7.69422926 0.9701058
## Med_1-Endo_0      -5.333333e+00 -17.360896  6.69422926 0.8901266
## Ovi_0-Endo_0      -6.000000e+00 -18.027563  6.02756259 0.7991836
## Ovi_0.5-Endo_0    -3.000000e+00 -15.027563  9.02756259 0.9983795
## Ovi_1-Endo_0      -2.000000e+00 -14.027563 10.02756259 0.9999643
## Endo_1-Endo_0.5   -3.552714e-15 -12.027563 12.02756259 1.0000000
## Med_0-Endo_0.5    -1.066667e+01 -22.694229  1.36089593 0.1152105
## Med_0.5-Endo_0.5 -7.333333e+00 -19.360896  4.69422926 0.5602112
## Med_1-Endo_0.5    -8.333333e+00 -20.360896  3.69422926 0.3817748
## Ovi_0-Endo_0.5    -9.000000e+00 -21.027563  3.02756259 0.2817048
## Ovi_0.5-Endo_0.5 -6.000000e+00 -18.027563  6.02756259 0.7991836
## Ovi_1-Endo_0.5    -5.000000e+00 -17.027563  7.02756259 0.9243575
## Med_0-Endo_1      -1.066667e+01 -22.694229  1.36089593 0.1152105
## Med_0.5-Endo_1    -7.333333e+00 -19.360896  4.69422926 0.5602112
## Med_1-Endo_1      -8.333333e+00 -20.360896  3.69422926 0.3817748
## Ovi_0-Endo_1      -9.000000e+00 -21.027563  3.02756259 0.2817048
## Ovi_0.5-Endo_1    -6.000000e+00 -18.027563  6.02756259 0.7991836
## Ovi_1-Endo_1      -5.000000e+00 -17.027563  7.02756259 0.9243575
## Med_0.5-Med_0      3.333333e+00  -8.694229 15.36089593 0.9960081
## Med_1-Med_0        2.333333e+00  -9.694229 14.36089593 0.9998394
## Ovi_0-Med_0        1.666667e+00 -10.360896 13.69422926 0.9999943
## Ovi_0.5-Med_0      4.666667e+00  -7.360896 16.69422926 0.9508375
## Ovi_1-Med_0        5.666667e+00  -6.360896 17.69422926 0.8481890
## Med_1-Med_0.5     -1.000000e+00 -13.027563 11.02756259 1.0000000
## Ovi_0-Med_0.5     -1.666667e+00 -13.694229 10.36089593 0.9999943
## Ovi_0.5-Med_0.5    1.333333e+00 -10.694229 13.36089593 0.9999994
## Ovi_1-Med_0.5      2.333333e+00  -9.694229 14.36089593 0.9998394
## Ovi_0-Med_1       -6.666667e-01 -12.694229 11.36089593 1.0000000
## Ovi_0.5-Med_1      2.333333e+00  -9.694229 14.36089593 0.9998394
## Ovi_1-Med_1        3.333333e+00  -8.694229 15.36089593 0.9960081
## Ovi_0.5-Ovi_0      3.000000e+00  -9.027563 15.02756259 0.9983795
## Ovi_1-Ovi_0        4.000000e+00  -8.027563 16.02756259 0.9831766
## Ovi_1-Ovi_0.5      1.000000e+00 -11.027563 13.02756259 1.0000000
```

```
##T1


# Fit a general linear model
model <- lm(T1 ~ TissueAlginate, data = data)
```

```r
# Perform Tukey post-hoc test
tukey_results <- TukeyHSD(aov(model))

# View the results
print(tukey_results)
```

```
##   Tukey multiple comparisons of means
##     95% family-wise confidence level
##
## Fit: aov(formula = model)
##
## $TissueAlginate
##                         diff        lwr       upr     p adj
## Ctx_0.5-Ctx_0     12.0000000 -17.639517 41.639517 0.9378192
## Ctx_1-Ctx_0       -8.6666667 -38.306184 20.972850 0.9941139
## Endo_0-Ctx_0       1.0000000 -28.639517 30.639517 1.0000000
## Endo_0.5-Ctx_0     3.3333333 -26.306184 32.972850 0.9999994
## Endo_1-Ctx_0      14.0000000 -15.639517 43.639517 0.8501065
## Med_0-Ctx_0       -8.0000000 -37.639517 21.639517 0.9969741
## Med_0.5-Ctx_0     13.3333333 -16.306184 42.972850 0.8844240
## Med_1-Ctx_0       15.3333333 -14.306184 44.972850 0.7679608
## Ovi_0-Ctx_0        4.0000000 -25.639517 33.639517 0.9999960
## Ovi_0.5-Ctx_0      2.6666667 -26.972850 32.306184 0.9999999
## Ovi_1-Ctx_0        1.3333333 -28.306184 30.972850 1.0000000
## Ctx_1-Ctx_0.5    -20.6666667 -50.306184  8.972850 0.3772186
## Endo_0-Ctx_0.5   -11.0000000 -40.639517 18.639517 0.9647911
## Endo_0.5-Ctx_0.5  -8.6666667 -38.306184 20.972850 0.9941139
## Endo_1-Ctx_0.5     2.0000000 -27.639517 31.639517 1.0000000
## Med_0-Ctx_0.5    -20.0000000 -49.639517  9.639517 0.4227033
## Med_0.5-Ctx_0.5    1.3333333 -28.306184 30.972850 1.0000000
## Med_1-Ctx_0.5      3.3333333 -26.306184 32.972850 0.9999994
## Ovi_0-Ctx_0.5     -8.0000000 -37.639517 21.639517 0.9969741
## Ovi_0.5-Ctx_0.5   -9.3333333 -38.972850 20.306184 0.9893927
## Ovi_1-Ctx_0.5    -10.6666667 -40.306184 18.972850 0.9715365
## Endo_0-Ctx_1       9.6666667 -19.972850 39.306184 0.9861182
## Endo_0.5-Ctx_1    12.0000000 -17.639517 41.639517 0.9378192
## Endo_1-Ctx_1      22.6666667  -6.972850 52.306184 0.2584553
## Med_0-Ctx_1        0.6666667 -28.972850 30.306184 1.0000000
## Med_0.5-Ctx_1     22.0000000  -7.639517 51.639517 0.2948684
## Med_1-Ctx_1       24.0000000  -5.639517 53.639517 0.1954684
## Ovi_0-Ctx_1       12.6666667 -16.972850 42.306184 0.9136932
## Ovi_0.5-Ctx_1     11.3333333 -18.306184 40.972850 0.9569670
## Ovi_1-Ctx_1       10.0000000 -19.639517 39.639517 0.9821121
## Endo_0.5-Endo_0    2.3333333 -27.306184 31.972850 1.0000000
## Endo_1-Endo_0     13.0000000 -16.639517 42.639517 0.8997017
## Med_0-Endo_0      -9.0000000 -38.639517 20.639517 0.9920279
## Med_0.5-Endo_0    12.3333333 -17.306184 41.972850 0.9263947
## Med_1-Endo_0      14.3333333 -15.306184 43.972850 0.8311551
## Ovi_0-Endo_0       3.0000000 -26.639517 32.639517 0.9999998
## Ovi_0.5-Endo_0     1.6666667 -27.972850 31.306184 1.0000000
```

```
## Ovi_1-Endo_0       0.3333333 -29.306184 29.972850 1.0000000
## Endo_1-Endo_0.5   10.6666667 -18.972850 40.306184 0.9715365
## Med_0-Endo_0.5   -11.3333333 -40.972850 18.306184 0.9569670
## Med_0.5-Endo_0.5  10.0000000 -19.639517 39.639517 0.9821121
## Med_1-Endo_0.5    12.0000000 -17.639517 41.639517 0.9378192
## Ovi_0-Endo_0.5     0.6666667 -28.972850 30.306184 1.0000000
## Ovi_0.5-Endo_0.5  -0.6666667 -30.306184 28.972850 1.0000000
## Ovi_1-Endo_0.5    -2.0000000 -31.639517 27.639517 1.0000000
## Med_0-Endo_1     -22.0000000 -51.639517  7.639517 0.2948684
## Med_0.5-Endo_1    -0.6666667 -30.306184 28.972850 1.0000000
## Med_1-Endo_1       1.3333333 -28.306184 30.972850 1.0000000
## Ovi_0-Endo_1     -10.0000000 -39.639517 19.639517 0.9821121
## Ovi_0.5-Endo_1   -11.3333333 -40.972850 18.306184 0.9569670
## Ovi_1-Endo_1     -12.6666667 -42.306184 16.972850 0.9136932
## Med_0.5-Med_0     21.3333333  -8.306184 50.972850 0.3345114
## Med_1-Med_0       23.3333333  -6.306184 52.972850 0.2253318
## Ovi_0-Med_0       12.0000000 -17.639517 41.639517 0.9378192
## Ovi_0.5-Med_0     10.6666667 -18.972850 40.306184 0.9715365
## Ovi_1-Med_0        9.3333333 -20.306184 38.972850 0.9893927
## Med_1-Med_0.5      2.0000000 -27.639517 31.639517 1.0000000
## Ovi_0-Med_0.5     -9.3333333 -38.972850 20.306184 0.9893927
## Ovi_0.5-Med_0.5  -10.6666667 -40.306184 18.972850 0.9715365
## Ovi_1-Med_0.5    -12.0000000 -41.639517 17.639517 0.9378192
## Ovi_0-Med_1      -11.3333333 -40.972850 18.306184 0.9569670
## Ovi_0.5-Med_1    -12.6666667 -42.306184 16.972850 0.9136932
## Ovi_1-Med_1      -14.0000000 -43.639517 15.639517 0.8501065
## Ovi_0.5-Ovi_0     -1.3333333 -30.972850 28.306184 1.0000000
## Ovi_1-Ovi_0       -2.6666667 -32.306184 26.972850 0.9999999
## Ovi_1-Ovi_0.5     -1.3333333 -30.972850 28.306184 1.0000000

####### Nanoindentation ###################################################

################### E' E" ##############################################


# Preprocess and aggregate data
data <- read.csv("E1_E2.csv")
data$Indentation <- as.numeric(data$Indentation)

# Aggregate data to calculate mean indentation
data_summary <- aggregate(Indentation ~ Hz + Tissue + Percentage_Alginate + M
odule, data, mean)
names(data_summary)[which(names(data_summary) == "Indentation")] <- "mean_Ind
entation"

# Loop through each unique tissue type in the aggregated summary data
for(tissue in unique(data_summary$Tissue)) {
  # Subset aggregated data for the specific tissue
  tissue_data <- subset(data_summary, Tissue == tissue)
```

```
  # Create the plot for the tissue using the aggregated data
  p <- ggplot(tissue_data, aes(x = Hz, y = mean_Indentation, color = as.facto
r(Percentage_Alginate))) +
    geom_line(aes(linetype = Module), linewidth = 1.5, alpha = 1) +  # Use 'M
odule' for linetype
    geom_point(size = 1, alpha = 1) +
    scale_color_viridis_d() +  # Use viridis discrete color scale
    scale_linetype_manual(values = c("E1" = "solid", "E2" = "dashed")) +  # D
efine linetypes for 'E1' and 'E2'
    scale_x_continuous(breaks = c(1, 5, 10, 20)) +
    scale_y_continuous(labels = scientific_format(), limits = c(0, 16000)) +
    labs(title = paste("Indentation vs. Hz for", tissue),
         y = expression(paste("E'/E'' (Pa)")),
         x = "Frequency (Hz)",
         color = "Percentage Alginate",
         linetype = "Module") +  # Correct label for linetype
    theme_minimal() +
    theme(panel.grid.major = element_blank(),
          panel.grid.minor = element_blank(),
          panel.border = element_rect(colour = "black", fill = NA, linewidth
= 1),
          axis.text = element_text(size = 16))

  # Save the plot
  ggsave(filename = paste0("plot_", tissue, ".png"), plot = p, width = 10, he
ight = 8)
}



################### YM #####################################################
###

#data input
data <- read.csv("YM_final.csv")


####     Basic Summary Statistics

# Means
means <- aggregate(EfYM ~ Tissue + Percentage_Alginate, data = data, FUN = "m
ean")

# SDs

sds <- aggregate(EfYM ~ Tissue + Percentage_Alginate, data = data, FUN = "sd"
)
```

```
SUM <- merge(means, sds, by = c("Tissue"))
names(SUM)[3:4] <- c("Mean", "SD")

write.csv(SUM, file = "Summary_YM.csv")

### Plot YM

png(filename="YM_final.png",
    width = 5, height = 5, units = "in",
    res = 600)


boxplot(EfYM/1000 ~ Percentage_Alginate*Tissue,
        data = data,
        cex.axis = 0.9,
        col = viridis(2, alpha = 0.7),
        lwd = 0.7,
        ylab = "",
        ylim = c(0,11),
        axes = FALSE,
        frame.plot = TRUE,
        xlab ="")

TISSUE <- unique(data$Tissue)
groups <- 4
numbox <- 2
total <- groups * numbox
xpoints <- seq(median(1:numbox),total,numbox)

axis(2, pretty(data$EfYM/1000), las = 1, cex.axis = 1)
axis(1, labels = TISSUE, at = xpoints,las = 2, cex.axis = 1)
abline(v = 2.5, lty = "dashed", col = "grey70")
abline(v = 4.5, lty = "dashed", col = "grey70")
abline(v = 6.5, lty = "dashed", col = "grey70")



stripchart(data$EfYM/1000 ~ data$Percentage_Alginate*data$Tissue,
# Data
           method = "jitter", # Random noise
           pch = 19,          # Pch symbols
           col = "black",           # Color of the symbol
           vertical = TRUE,   # Vertical mode
           add = TRUE,
           cex = 0.25)        # Add it over

title(ylab = expression("Young's modulud (KPa)"), line = 2.2)
legend("top", fill = viridis(2), legend = c("0.5% alginate", "1.0% alginate")
```

```
, horiz = TRUE, cex = 0.7)

dev.off()

## quartz_off_screen
##                  2

### Cortex
#data input
data <- read.csv("YM_final_ctx.csv")


####     Basic Summary Statistics

# Means
means <- aggregate(EfYM ~ Percentage_Alginate, data = data, FUN = "mean")

# SDs

sds <- aggregate(EfYM ~ Percentage_Alginate, data = data, FUN = "sd")

SUM <- merge(means, sds, by = c("Percentage_Alginate"))
names(SUM)[2:3] <- c("Mean", "SD")

write.csv(SUM, file = "Summary_YM_ctx.csv")

### Plot YM

png(filename="YM_final_ctx.png",
    width = 5, height = 5, units = "in",
    res = 600)


boxplot(EfYM/1000 ~ Percentage_Alginate,
        data = data,
        cex.axis = 0.9,
        col = viridis(3, alpha = 0.7),
        lwd = 0.7,
        ylab = "",
        ylim = c(0,20),
        axes = TRUE,
        frame.plot = TRUE,
        xlab ="")

stripchart(data$EfYM/1000 ~ data$Percentage_Alginate,               # Data
           method = "jitter", # Random noise
           pch = 19,          # Pch symbols
           col = "black",           # Color of the symbol
           vertical = TRUE,   # Vertical mode
```

```
           add = TRUE,
           cex = 0.25)        # Add it over

title(ylab = expression("Young's modulud (KPa)"), line = 2.2)


dev.off()

## quartz_off_screen
##                 2

##Stats

# Fit a general linear model
model <- lm(EfYM ~ Percentage_Alginate, data = data)

# Perform Tukey post-hoc test
tukey_results <- TukeyHSD(aov(model))

# View the results
print(tukey_results)

##   Tukey multiple comparisons of means
##     95% family-wise confidence level
##
## Fit: aov(formula = model)
##
## $Percentage_Alginate
##                 diff       lwr      upr     p adj
## 1-0.5      1070.951  134.8825 2007.020 0.0207533
## Tissue-0.5 6846.478 5868.7469 7824.209 0.0000000
## Tissue-1   5775.527 4773.2107 6777.843 0.0000000

### Medulla
#data input
data <- read.csv("YM_final_med.csv")


####     Basic Summary Statistics

# Means
means <- aggregate(EfYM ~ Percentage_Alginate, data = data, FUN = "mean")

# SDs

sds <- aggregate(EfYM ~ Percentage_Alginate, data = data, FUN = "sd")

SUM <- merge(means, sds, by = c("Percentage_Alginate"))
names(SUM)[2:3] <- c("Mean", "SD")
```

```r
write.csv(SUM, file = "Summary_YM_med.csv")

### Plot YM

png(filename="YM_final_med.png",
    width = 5, height = 5, units = "in",
    res = 600)


boxplot(EfYM/1000 ~ Percentage_Alginate,
        data = data,
        cex.axis = 0.9,
        col = viridis(3, alpha = 0.7),
        lwd = 0.7,
        ylab = "",
        ylim = c(0,20),
        axes = TRUE,
        frame.plot = TRUE,
        xlab ="")

stripchart(data$EfYM/1000 ~ data$Percentage_Alginate,                # Data
           method = "jitter", # Random noise
           pch = 19,          # Pch symbols
           col = "black",           # Color of the symbol
           vertical = TRUE,   # Vertical mode
           add = TRUE,
           cex = 0.25)        # Add it over

title(ylab = expression("Young's modulud (KPa)"), line = 2.2)


dev.off()

## quartz_off_screen
##                  2

##Stats

# Fit a general linear model
model <- lm(EfYM ~ Percentage_Alginate, data = data)

# Perform Tukey post-hoc test
tukey_results <- TukeyHSD(aov(model))

# View the results
print(tukey_results)

##   Tukey multiple comparisons of means
##     95% family-wise confidence level
```

```
##
## Fit: aov(formula = model)
##
## $Percentage_Alginate
##                 diff        lwr      upr     p adj
## 1-0.5      1287.100  232.78614 2341.414 0.0123198
## Tissue-0.5 2534.657 1253.89811 3815.416 0.0000202
## Tissue-1   1247.557  -65.54452 2560.659 0.0663602

### Oviduct
#data input
data <- read.csv("YM_final_ovi.csv")


####     Basic Summary Statistics

# Means
means <- aggregate(EfYM ~ Percentage_Alginate, data = data, FUN = "mean")

# SDs

sds <- aggregate(EfYM ~ Percentage_Alginate, data = data, FUN = "sd")

SUM <- merge(means, sds, by = c("Percentage_Alginate"))
names(SUM)[2:3] <- c("Mean", "SD")

write.csv(SUM, file = "Summary_YM_ovi.csv")

### Plot YM

png(filename="YM_final_ovi.png",
    width = 5, height = 5, units = "in",
    res = 600)


boxplot(EfYM/1000 ~ Percentage_Alginate,
        data = data,
        cex.axis = 0.9,
        col = viridis(3, alpha = 0.7),
        lwd = 0.7,
        ylab = "",
        ylim = c(0,20),
        axes = TRUE,
        frame.plot = TRUE,
        xlab ="")

stripchart(data$EfYM/1000 ~ data$Percentage_Alginate,               # Data
           method = "jitter", # Random noise
           pch = 19,          # Pch symbols
```

```
           col = "black",            # Color of the symbol
           vertical = TRUE,    # Vertical mode
           add = TRUE,
           cex = 0.25)         # Add it over

title(ylab = expression("Young's modulud (KPa)"), line = 2.2)


dev.off()

## quartz_off_screen
##                 2

##Stats

# Fit a general linear model
model <- lm(EfYM ~ Percentage_Alginate, data = data)

# Perform Tukey post-hoc test
tukey_results <- TukeyHSD(aov(model))

# View the results
print(tukey_results)

##   Tukey multiple comparisons of means
##     95% family-wise confidence level
##
## Fit: aov(formula = model)
##
## $Percentage_Alginate
##                 diff       lwr      upr     p adj
## 1-0.5      326.8711 -411.5045 1065.247 0.5459376
## Tissue-0.5 583.5801 -291.5110 1458.671 0.2566099
## Tissue-1   256.7091 -592.9485 1106.367 0.7534396

### Endo
#data input
data <- read.csv("YM_final_endo.csv")


####     Basic Summary Statistics

# Means
means <- aggregate(EfYM ~ Percentage_Alginate, data = data, FUN = "mean")

# SDs

sds <- aggregate(EfYM ~ Percentage_Alginate, data = data, FUN = "sd")

SUM <- merge(means, sds, by = c("Percentage_Alginate"))
```

```r
names(SUM)[2:3] <- c("Mean", "SD")

write.csv(SUM, file = "Summary_YM_endo.csv")

### Plot YM

png(filename="YM_final_endo.png",
    width = 5, height = 5, units = "in",
    res = 600)


boxplot(EfYM/1000 ~ Percentage_Alginate,
        data = data,
        cex.axis = 0.9,
        col = viridis(3, alpha = 0.7),
        lwd = 0.7,
        ylab = "",
        ylim = c(0,20),
        axes = TRUE,
        frame.plot = TRUE,
        xlab ="")

stripchart(data$EfYM/1000 ~ data$Percentage_Alginate,               # Data
           method = "jitter", # Random noise
           pch = 19,          # Pch symbols
           col = "black",           # Color of the symbol
           vertical = TRUE,   # Vertical mode
           add = TRUE,
           cex = 0.25)        # Add it over

title(ylab = expression("Young's modulud (KPa)"), line = 2.2)


dev.off()

## quartz_off_screen
##                 2

##Stats

# Fit a general linear model
model <- lm(EfYM ~ Percentage_Alginate, data = data)

# Perform Tukey post-hoc test
tukey_results <- TukeyHSD(aov(model))

# View the results
print(tukey_results)
```

```
##   Tukey multiple comparisons of means
##     95% family-wise confidence level
##
## Fit: aov(formula = model)
##
## $Percentage_Alginate
##                 diff       lwr      upr     p adj
## 1-0.5      1355.8107  502.5186 2209.103 0.0007477
## Tissue-0.5 1957.9194 1061.4847 2854.354 0.0000028
## Tissue-1    602.1087 -336.3415 1540.559 0.2837201

############## Cell Proliferation#########################################

#data input
data <- read.csv("Proliferation.csv")
data$Day <- factor(data$Day)
data$Alginate_content <- factor(data$Alginate)

####Basic Summary Statistics


# Means
means <- aggregate(Prolif ~Tissue + Alginate, data = data, FUN = "mean")

# SDs

sds <- aggregate(Prolif ~Tissue + Alginate, data = data, FUN = "sd")

SUM <- merge(means, sds, by = c("Tissue"))
names(SUM)[3:4] <- c("Mean", "SD")

write.csv(SUM, file = "Summary_Proliferation.csv")


##### plot


png(filename="Proliferation.png",
    width = 8.2, height = 5, units = "in",
    res = 600)


boxplot(Prolif ~ Alginate*Tissue,
          data = data,
          cex.axis = 0.9,
          col = viridis(3, alpha = 0.7),
          lwd = 0.7,
          ylab = "",
          ylim = c(0,100),
```

```
          axes = FALSE,
          frame.plot = TRUE,
          xlab ="")

TISSUE <- unique(data$Tissue)
groups <- 4
numbox <- 3
total <- groups * numbox
xpoints <- seq(median(1:numbox),total,numbox)

axis(2, pretty(data$Prolif), las = 1, cex.axis = 1)
#axis(1, labels = Alginate, at = xpoints,las = 1, cex.axis = 1)
abline(v = 3.5, lty = "dashed", col = "grey70")
abline(v = 6.5, lty = "dashed", col = "grey70")
abline(v = 9.5, lty = "dashed", col = "grey70")



stripchart(data$Prolif ~ data$Alginate*data$Tissue,                # Data
           method = "jitter", # Random noise
           pch = 19,          # Pch symbols
           col = "black",           # Color of the symbol
           vertical = TRUE,   # Vertical mode
           add = TRUE,
           cex = 0.25)        # Add it over

title(ylab = expression("Cell proliferation (%)"), line = 2.5)

dev.off()

## quartz_off_screen
##                  2

##Stat
subset_ctx <- subset(data, Tissue == "Cortex")
subset_endo <- subset(data, Tissue == "Endo")
subset_med <- subset(data, Tissue == "Medulla")
subset_ovi <- subset(data, Tissue == "Oviduct")

## Cortex

# Fit a general linear model
model <- lm(Prolif ~ Alginate_content, data = subset_ctx)

# Perform Tukey post-hoc test
tukey_results <- TukeyHSD(aov(model))

# View the results
print(tukey_results)
```

```
##   Tukey multiple comparisons of means
##     95% family-wise confidence level
##
## Fit: aov(formula = model)
##
## $Alginate_content
##              diff         lwr       upr     p adj
## 0.5-0  17.323732   0.9353182 33.712146 0.0368633
## 1-0    -7.984961 -24.3733746  8.403453 0.4550704
## 1-0.5 -25.308693 -41.6971068 -8.920279 0.0021064
```

```
## Endo

# Fit a general linear model
model <- lm(Prolif ~ Alginate_content, data = subset_endo)

# Perform Tukey post-hoc test
tukey_results <- TukeyHSD(aov(model))

# View the results
print(tukey_results)
```

```
##   Tukey multiple comparisons of means
##     95% family-wise confidence level
##
## Fit: aov(formula = model)
##
## $Alginate_content
##             diff       lwr       upr     p adj
## 0.5-0 -2.047717 -16.81336 12.717923 0.9361981
## 1-0   -8.255357 -23.02100  6.510283 0.3586549
## 1-0.5 -6.207639 -20.97328  8.558001 0.5535220
```

```
## Medulla

# Fit a general linear model
model <- lm(Prolif ~ Alginate_content, data = subset_med)

# Perform Tukey post-hoc test
tukey_results <- TukeyHSD(aov(model))

# View the results
print(tukey_results)
```

```
##   Tukey multiple comparisons of means
##     95% family-wise confidence level
##
## Fit: aov(formula = model)
##
## $Alginate_content
```

```
##              diff       lwr        upr     p adj
## 0.5-0   8.300837 -13.40029 30.0019631 0.6100989
## 1-0   -22.020568 -44.38958  0.3484401 0.0542304
## 1-0.5 -30.321406 -52.69041 -7.9523972 0.0067674

## Oviduct

# Fit a general linear model
model <- lm(Prolif ~ Alginate_content, data = subset_ovi)

# Perform Tukey post-hoc test
tukey_results <- TukeyHSD(aov(model))

# View the results
print(tukey_results)

##   Tukey multiple comparisons of means
##     95% family-wise confidence level
##
## Fit: aov(formula = model)
##
## $Alginate_content
##              diff        lwr        upr     p adj
## 0.5-0 -17.0844389 -23.344562 -10.824316 0.0000014
## 1-0   -17.7486355 -24.008758 -11.488513 0.0000007
## 1-0.5  -0.6641966  -6.924319   5.595926 0.9620996

## YM NO-Alginate samples

#data input
data <- read.csv("YM_0alg.csv")


####     Basic Summary Statistics

# Means
means <- aggregate(EfYM ~ Tissue, data = data, FUN = "mean")

# SDs

sds <- aggregate(EfYM ~ Tissue, data = data, FUN = "sd")

SUM <- merge(means, sds, by = c("Tissue"))
names(SUM)[2:3] <- c("Mean", "SD")

write.csv(SUM, file = "Summary_YM_0alg.csv")

### Plot YM
```

```r
png(filename="YM_no_alg.png",
    width = 5, height = 5, units = "in",
    res = 600)


boxplot(EfYM/1000 ~ Tissue,
        data = data,
        cex.axis = 0.9,
        col = viridis(3, alpha = 0.7),
        lwd = 0.7,
        ylab = "",
        ylim = c(0,0.3),
        axes = TRUE,
        frame.plot = TRUE,
        xlab ="")

stripchart(data$EfYM/1000 ~ data$Tissue,               # Data
           method = "jitter", # Random noise
           pch = 19,          # Pch symbols
           col = "black",           # Color of the symbol
           vertical = TRUE,   # Vertical mode
           add = TRUE,
           cex = 0.25)        # Add it over

title(ylab = expression("Young's modulus (KPa)"), line = 2.2)


dev.off()

## quartz_off_screen
##                 2

############## Cell Viability#################################################
####

### Endometrium
#data input
data <- read.csv("Live_Dead_endo.csv")
data$Day <- factor(data$Day)
data$Alginate <- factor(data$Alginate)
data$Alginate_content <- factor(data$Alginate)

####Basic Summary Statistics


# Means
means <- aggregate(Dead ~Day + Alginate, data = data, FUN = "mean")

# SDs
```

```r
sds <- aggregate(Dead ~Day + Alginate, data = data, FUN = "sd")

SUM <- merge(means, sds, by = c("Day"))
names(SUM)[3:4] <- c("Mean", "SD")

write.csv(SUM, file = "Summary_Viability_endo.csv")


##### plot


png(filename="Cell_viability_endo.png",
    width = 6, height = 6, units = "in",
    res = 600)


boxplot(Dead ~ Day*Alginate,
        data = data,
        cex.axis = 0.9,
        col = viridis(3, alpha = 0.7),
        lwd = 0.7,
        ylab = "",
        ylim = c(0,40),
        axes = FALSE,
        frame.plot = TRUE,
        xlab ="")

TISSUE <- unique(data$Alginate)
groups <- 4
numbox <- 3
total <- groups * numbox
xpoints <- seq(median(1:numbox),total,numbox)

axis(2, pretty(data$Dead), las = 1, cex.axis = 1)
#axis(1, labels = Alginate, at = xpoints,las = 1, cex.axis = 1)
abline(v = 3.5, lty = "dashed", col = "grey70")
abline(v = 6.5, lty = "dashed", col = "grey70")


stripchart(data$Dead ~ data$Day*data$Alginate,               # Data
           method = "jitter", # Random noise
           pch = 19,          # Pch symbols
           col = "black",           # Color of the symbol
           vertical = TRUE,   # Vertical mode
           add = TRUE,
           cex = 0.25)        # Add it over

title(ylab = expression("Dead cells (%)"), line = 2)
```

```r
dev.off()

## quartz_off_screen
##                 2

### LM individual days by alginate

##Stat
subset_1 <- subset(data, Day == "1")
subset_7 <- subset(data, Day == "7")
subset_14 <- subset(data, Day == "14")

##Day 1

# Fit a general linear model
model <- lm(Dead ~ Alginate, data = subset_1)

# Perform Tukey post-hoc test
tukey_results <- TukeyHSD(aov(model))

# View the results
print(tukey_results)

##   Tukey multiple comparisons of means
##     95% family-wise confidence level
##
## Fit: aov(formula = model)
##
## $Alginate
##              diff       lwr       upr     p adj
## 0.5-0  -7.303238 -16.60784  2.001364 0.1434837
## 1-0   -10.849925 -20.15453 -1.545323 0.0202543
## 1-0.5  -3.546687 -12.57348  5.480103 0.5940998

##Day 7

# Fit a general linear model
model <- lm(Dead ~ Alginate, data = subset_7)

# Perform Tukey post-hoc test
tukey_results <- TukeyHSD(aov(model))

# View the results
print(tukey_results)

##   Tukey multiple comparisons of means
##     95% family-wise confidence level
##
## Fit: aov(formula = model)
```

```
##
## $Alginate
##              diff       lwr      upr     p adj
## 0.5-0 -1.834702 -10.86792 7.198519 0.8591598
## 1-0   -7.387673 -16.42089 1.645549 0.1183407
## 1-0.5 -5.552971 -14.58619 3.480251 0.2773474

## Day 14

# Fit a general linear model
model <- lm(Dead ~ Alginate, data = subset_14)

# Perform Tukey post-hoc test
tukey_results <- TukeyHSD(aov(model))

# View the results
print(tukey_results)

##   Tukey multiple comparisons of means
##     95% family-wise confidence level
##
## Fit: aov(formula = model)
##
## $Alginate
##              diff       lwr      upr     p adj
## 0.5-0 -1.0247825 -3.986149 1.936584 0.6675199
## 1-0   -0.4606072 -3.421973 2.500759 0.9204746
## 1-0.5  0.5641752 -2.397191 3.525541 0.8833041

### LM individual alginate by days

##Stat
subset_0 <- subset(data, Alginate == "0")
subset_0.5 <- subset(data, Alginate == "0.5")
subset_1 <- subset(data, Alginate == "1")

##0%

# Fit a general linear model
model <- lm(Dead ~ Day, data = subset_0)

# Perform Tukey post-hoc test
tukey_results <- TukeyHSD(aov(model))

# View the results
print(tukey_results)

##   Tukey multiple comparisons of means
##     95% family-wise confidence level
##
```

```
## Fit: aov(formula = model)
##
## $Day
##              diff       lwr       upr     p adj
## 7-1   -7.907336 -20.21341  4.398741 0.2583113
## 14-1 -14.085915 -25.15813 -3.013702 0.0114755
## 14-7  -6.178580 -18.18808  5.830922 0.4106700
```

```
##0.5%

# Fit a general linear model
model <- lm(Dead ~ Day, data = subset_0.5)

# Perform Tukey post-hoc test
tukey_results <- TukeyHSD(aov(model))

# View the results
print(tukey_results)
```

```
##   Tukey multiple comparisons of means
##     95% family-wise confidence level
##
## Fit: aov(formula = model)
##
## $Day
##          diff        lwr        upr     p adj
## 7-1  -2.43880  -7.220256  2.3426561 0.4185427
## 14-1 -7.80746 -12.084124 -3.5307956 0.0004357
## 14-7 -5.36866 -10.150116 -0.5872038 0.0260239
```

```
##1%

# Fit a general linear model
model <- lm(Dead ~ Day, data = subset_1)

# Perform Tukey post-hoc test
tukey_results <- TukeyHSD(aov(model))

# View the results
print(tukey_results)
```

```
##   Tukey multiple comparisons of means
##     95% family-wise confidence level
##
## Fit: aov(formula = model)
##
## $Day
##              diff       lwr        upr     p adj
## 7-1  -4.4450837 -7.952637 -0.9375305 0.0116352
```

```
## 14-1 -3.6965975 -6.833848 -0.5593465 0.0191812
## 14-7  0.7484862 -2.759067  4.2560394 0.8536791

### Cortex
#data input
data <- read.csv("Live_Dead_ctx.csv")
data$Day <- factor(data$Day)
data$Alginate <- factor(data$Alginate)
data$Alginate_content <- factor(data$Alginate)

####Basic Summary Statistics


# Means
means <- aggregate(Dead ~Day + Alginate, data = data, FUN = "mean")

# SDs

sds <- aggregate(Dead ~Day + Alginate, data = data, FUN = "sd")

SUM <- merge(means, sds, by = c("Day"))
names(SUM)[3:4] <- c("Mean", "SD")

write.csv(SUM, file = "Summary_Viability_Cortex.csv")


##### plot


png(filename="Cell_viability_Cortex.png",
    width = 6, height = 6, units = "in",
    res = 600)


boxplot(Dead ~ Day*Alginate,
        data = data,
        cex.axis = 0.9,
        col = viridis(3, alpha = 0.7),
        lwd = 0.7,
        ylab = "",
        ylim = c(0,40),
        axes = FALSE,
        frame.plot = TRUE,
        xlab ="")

TISSUE <- unique(data$Alginate)
groups <- 4
numbox <- 3
total <- groups * numbox
```

```
xpoints <- seq(median(1:numbox),total,numbox)

axis(2, pretty(data$Dead), las = 1, cex.axis = 1)
#axis(1, labels = Alginate, at = xpoints,las = 1, cex.axis = 1)
abline(v = 3.5, lty = "dashed", col = "grey70")
abline(v = 6.5, lty = "dashed", col = "grey70")


stripchart(data$Dead ~ data$Day*data$Alginate,               # Data
           method = "jitter", # Random noise
           pch = 19,          # Pch symbols
           col = "black",           # Color of the symbol
           vertical = TRUE,   # Vertical mode
           add = TRUE,
           cex = 0.25)        # Add it over

title(ylab = expression("Dead cells (%)"), line = 2)

dev.off()

## quartz_off_screen
##                 2

### LM individual days by alginate

##Stat
subset_1 <- subset(data, Day == "1")
subset_7 <- subset(data, Day == "7")
subset_14 <- subset(data, Day == "14")

##Day 1

# Fit a general linear model
model <- lm(Dead ~ Alginate, data = subset_1)

# Perform Tukey post-hoc test
tukey_results <- TukeyHSD(aov(model))

# View the results
print(tukey_results)

##   Tukey multiple comparisons of means
##     95% family-wise confidence level
##
## Fit: aov(formula = model)
##
## $Alginate
##             diff       lwr      upr     p adj
## 0.5-0  2.7282486 -4.483659 9.940156 0.6165831
```

```
## 1-0    2.1748781 -5.037029 9.386785 0.7335567
## 1-0.5 -0.5533705 -7.549948 6.443207 0.9786240
```

```r
##Day 7

# Fit a general linear model
model <- lm(Dead ~ Alginate, data = subset_7)

# Perform Tukey post-hoc test
tukey_results <- TukeyHSD(aov(model))

# View the results
print(tukey_results)
```

```
##   Tukey multiple comparisons of means
##     95% family-wise confidence level
##
## Fit: aov(formula = model)
##
## $Alginate
##              diff       lwr       upr     p adj
## 0.5-0 -0.9180497 -2.784811 0.9487112 0.4471821
## 1-0   -1.2076976 -3.074459 0.6590633 0.2575635
## 1-0.5 -0.2896479 -2.100672 1.5213762 0.9156898
```

```r
## Day 14

# Fit a general linear model
model <- lm(Dead ~ Alginate, data = subset_14)

# Perform Tukey post-hoc test
tukey_results <- TukeyHSD(aov(model))

# View the results
print(tukey_results)
```

```
##   Tukey multiple comparisons of means
##     95% family-wise confidence level
##
## Fit: aov(formula = model)
##
## $Alginate
##              diff        lwr      upr     p adj
## 0.5-0 -0.7715299 -4.0193274 2.476268 0.8250343
## 1-0    1.7130020 -1.5347954 4.960799 0.3996729
## 1-0.5  2.4845319 -0.7632655 5.732329 0.1574841
```

```r
### LM individual alginate by days

##Stat
```

```r
subset_0 <- subset(data, Alginate == "0")
subset_0.5 <- subset(data, Alginate == "0.5")
subset_1 <- subset(data, Alginate == "1")

##0%

# Fit a general linear model
model <- lm(Dead ~ Day, data = subset_0)

# Perform Tukey post-hoc test
tukey_results <- TukeyHSD(aov(model))

# View the results
print(tukey_results)

##   Tukey multiple comparisons of means
##     95% family-wise confidence level
##
## Fit: aov(formula = model)
##
## $Day
##             diff       lwr      upr     p adj
## 7-1  -0.9040777 -4.135231 2.327076 0.7643231
## 14-1 -2.0454984 -5.185615 1.094618 0.2519662
## 14-7 -1.1414208 -4.281538 1.998696 0.6379174

##0.5%

# Fit a general linear model
model <- lm(Dead ~ Day, data = subset_0.5)

# Perform Tukey post-hoc test
tukey_results <- TukeyHSD(aov(model))

# View the results
print(tukey_results)

##   Tukey multiple comparisons of means
##     95% family-wise confidence level
##
## Fit: aov(formula = model)
##
## $Day
##             diff        lwr        upr     p adj
## 7-1  -4.5503760  -9.590649  0.4898967 0.0822137
## 14-1 -5.5452769 -10.585550 -0.5050042 0.0291170
## 14-7 -0.9949009  -6.035174  4.0453717 0.8753316

##1%
```

```r
# Fit a general linear model
model <- lm(Dead ~ Day, data = subset_1)

# Perform Tukey post-hoc test
tukey_results <- TukeyHSD(aov(model))

# View the results
print(tukey_results)

##   Tukey multiple comparisons of means
##     95% family-wise confidence level
##
## Fit: aov(formula = model)
##
## $Day
##            diff       lwr       upr     p adj
## 7-1  -4.286653 -9.435522 0.8622157 0.1155241
## 14-1 -2.507374 -7.656243 2.6414946 0.4554343
## 14-7  1.779279 -3.369590 6.9281479 0.6682603

### Medulla
#data input
data <- read.csv("Live_Dead_med.csv")
data$Day <- factor(data$Day)
data$Alginate <- factor(data$Alginate)
data$Alginate_content <- factor(data$Alginate)

####Basic Summary Statistics


# Means
means <- aggregate(Dead ~Day + Alginate, data = data, FUN = "mean")

# SDs

sds <- aggregate(Dead ~Day + Alginate, data = data, FUN = "sd")

SUM <- merge(means, sds, by = c("Day"))
names(SUM)[3:4] <- c("Mean", "SD")

write.csv(SUM, file = "Summary_Viability_Medulla.csv")


##### plot


png(filename="Cell_viability_Medulla.png",
    width = 6, height = 6, units = "in",
    res = 600)
```

```r
boxplot(Dead ~ Day*Alginate,
        data = data,
        cex.axis = 0.9,
        col = viridis(3, alpha = 0.7),
        lwd = 0.7,
        ylab = "",
        ylim = c(0,40),
        axes = FALSE,
        frame.plot = TRUE,
        xlab ="")

TISSUE <- unique(data$Alginate)
groups <- 4
numbox <- 3
total <- groups * numbox
xpoints <- seq(median(1:numbox),total,numbox)

axis(2, pretty(data$Dead), las = 1, cex.axis = 1)
#axis(1, labels = Alginate, at = xpoints,las = 1, cex.axis = 1)
abline(v = 3.5, lty = "dashed", col = "grey70")
abline(v = 6.5, lty = "dashed", col = "grey70")


stripchart(data$Dead ~ data$Day*data$Alginate,              # Data
           method = "jitter", # Random noise
           pch = 19,          # Pch symbols
           col = "black",           # Color of the symbol
           vertical = TRUE,   # Vertical mode
           add = TRUE,
           cex = 0.25)        # Add it over

title(ylab = expression("Dead cells (%)"), line = 2)

dev.off()

## quartz_off_screen
##                  2

### LM individual days by alginate

##Stat
subset_1 <- subset(data, Day == "1")
subset_7 <- subset(data, Day == "7")
subset_14 <- subset(data, Day == "14")

##Day 1
```

```r
# Fit a general linear model
model <- lm(Dead ~ Alginate, data = subset_1)

# Perform Tukey post-hoc test
tukey_results <- TukeyHSD(aov(model))

# View the results
print(tukey_results)
```

```
##   Tukey multiple comparisons of means
##     95% family-wise confidence level
##
## Fit: aov(formula = model)
##
## $Alginate
##              diff       lwr      upr     p adj
## 0.5-0 -1.5965547 -4.504647 1.311537 0.3714585
## 1-0   -1.2306043 -4.138696 1.677488 0.5493490
## 1-0.5  0.3659504 -2.542141 3.274042 0.9471366
```

```r
##Day 7

# Fit a general linear model
model <- lm(Dead ~ Alginate, data = subset_7)

# Perform Tukey post-hoc test
tukey_results <- TukeyHSD(aov(model))

# View the results
print(tukey_results)
```

```
##   Tukey multiple comparisons of means
##     95% family-wise confidence level
##
## Fit: aov(formula = model)
##
## $Alginate
##              diff       lwr      upr     p adj
## 0.5-0 -0.2784576 -1.666070 1.109155 0.8714547
## 1-0   -0.1747119 -1.562324 1.212901 0.9470799
## 1-0.5  0.1037457 -1.283867 1.491358 0.9809802
```

```r
## Day 14

# Fit a general linear model
model <- lm(Dead ~ Alginate, data = subset_14)

# Perform Tukey post-hoc test
tukey_results <- TukeyHSD(aov(model))
```

```r
# View the results
print(tukey_results)
```

```
##   Tukey multiple comparisons of means
##     95% family-wise confidence level
##
## Fit: aov(formula = model)
##
## $Alginate
##              diff       lwr       upr     p adj
## 0.5-0 -6.2146083 -18.51482  6.085603 0.4298261
## 1-0   -5.9359593 -18.23617  6.364252 0.4616797
## 1-0.5  0.2786489 -12.02156 12.578860 0.9982371
```

```r
### LM individual alginate by days

##Stat
subset_0 <- subset(data, Alginate == "0")
subset_0.5 <- subset(data, Alginate == "0.5")
subset_1 <- subset(data, Alginate == "1")

##0%

# Fit a general linear model
model <- lm(Dead ~ Day, data = subset_0)

# Perform Tukey post-hoc test
tukey_results <- TukeyHSD(aov(model))

# View the results
print(tukey_results)
```

```
##   Tukey multiple comparisons of means
##     95% family-wise confidence level
##
## Fit: aov(formula = model)
##
## $Day
##            diff        lwr      upr     p adj
## 7-1   -1.712535 -14.198051 10.77298 0.9375402
## 14-1   3.824850  -8.660666 16.31037 0.7275991
## 14-7   5.537385  -6.948130 18.02290 0.5187864
```

```r
##0.5%

# Fit a general linear model
model <- lm(Dead ~ Day, data = subset_0.5)

# Perform Tukey post-hoc test
tukey_results <- TukeyHSD(aov(model))
```

```
# View the results
print(tukey_results)

##   Tukey multiple comparisons of means
##     95% family-wise confidence level
##
## Fit: aov(formula = model)
##
## $Day
##             diff       lwr      upr     p adj
## 7-1  -0.3944382 -2.430209 1.641333 0.8795719
## 14-1 -0.7932035 -2.828975 1.242567 0.6005396
## 14-7 -0.3987653 -2.434536 1.637006 0.8771004

##1%

# Fit a general linear model
model <- lm(Dead ~ Day, data = subset_1)

# Perform Tukey post-hoc test
tukey_results <- TukeyHSD(aov(model))

# View the results
print(tukey_results)

##   Tukey multiple comparisons of means
##     95% family-wise confidence level
##
## Fit: aov(formula = model)
##
## $Day
##             diff       lwr       upr     p adj
## 7-1  -0.6566429 -1.939298 0.6260125 0.4206127
## 14-1 -0.8805050 -2.163160 0.4021504 0.2205362
## 14-7 -0.2238621 -1.506517 1.0587933 0.9010134

### Oviduct
#data input
data <- read.csv("Live_Dead_ovi.csv")
data$Day <- factor(data$Day)
data$Alginate <- factor(data$Alginate)
data$Alginate_content <- factor(data$Alginate)

####Basic Summary Statistics


# Means
means <- aggregate(Dead ~Day + Alginate, data = data, FUN = "mean")
```

```r
# SDs

sds <- aggregate(Dead ~Day + Alginate, data = data, FUN = "sd")

SUM <- merge(means, sds, by = c("Day"))
names(SUM)[3:4] <- c("Mean", "SD")

write.csv(SUM, file = "Summary_Viability_Oviduct.csv")


##### plot


png(filename="Cell_viability_Oviduct.png",
    width = 6, height = 6, units = "in",
    res = 600)


boxplot(Dead ~ Day*Alginate,
        data = data,
        cex.axis = 0.9,
        col = viridis(3, alpha = 0.7),
        lwd = 0.7,
        ylab = "",
        ylim = c(0,40),
        axes = FALSE,
        frame.plot = TRUE,
        xlab ="")

TISSUE <- unique(data$Alginate)
groups <- 4
numbox <- 3
total <- groups * numbox
xpoints <- seq(median(1:numbox),total,numbox)

axis(2, pretty(data$Dead), las = 1, cex.axis = 1)
#axis(1, labels = Alginate, at = xpoints,las = 1, cex.axis = 1)
abline(v = 3.5, lty = "dashed", col = "grey70")
abline(v = 6.5, lty = "dashed", col = "grey70")


stripchart(data$Dead ~ data$Day*data$Alginate,               # Data
           method = "jitter", # Random noise
           pch = 19,          # Pch symbols
           col = "black",           # Color of the symbol
           vertical = TRUE,   # Vertical mode
           add = TRUE,
           cex = 0.25)        # Add it over
```

```r
title(ylab = expression("Dead cells (%)"), line = 2)

dev.off()

## quartz_off_screen
##                 2

### LM individual days by alginate

##Stat
subset_1 <- subset(data, Day == "1")
subset_7 <- subset(data, Day == "7")
subset_14 <- subset(data, Day == "14")

##Day 1

# Fit a general linear model
model <- lm(Dead ~ Alginate, data = subset_1)

# Perform Tukey post-hoc test
tukey_results <- TukeyHSD(aov(model))

# View the results
print(tukey_results)

##   Tukey multiple comparisons of means
##     95% family-wise confidence level
##
## Fit: aov(formula = model)
##
## $Alginate
##           diff       lwr      upr     p adj
## 0.5-0 3.610720 -4.742606 11.96404 0.5356411
## 1-0   5.871582 -2.481743 14.22491 0.2059819
## 1-0.5 2.260862 -6.092463 10.61419 0.7795702

##Day 7

# Fit a general linear model
model <- lm(Dead ~ Alginate, data = subset_7)

# Perform Tukey post-hoc test
tukey_results <- TukeyHSD(aov(model))

# View the results
print(tukey_results)

##   Tukey multiple comparisons of means
##     95% family-wise confidence level
##
```

```
## Fit: aov(formula = model)
##
## $Alginate
##               diff        lwr       upr     p adj
## 0.5-0 -0.8090039 -6.4403107  4.822303 0.9320421
## 1-0    5.3712160 -0.2600908 11.002523 0.0636250
## 1-0.5  6.1802199  0.4026210 11.957819 0.0343544

## Day 14

# Fit a general linear model
model <- lm(Dead ~ Alginate, data = subset_14)

# Perform Tukey post-hoc test
tukey_results <- TukeyHSD(aov(model))

# View the results
print(tukey_results)

##   Tukey multiple comparisons of means
##     95% family-wise confidence level
##
## Fit: aov(formula = model)
##
## $Alginate
##               diff        lwr       upr     p adj
## 0.5-0 -6.7439282 -11.168797 -2.319059 0.0023865
## 1-0   -5.7493970 -10.174266 -1.324528 0.0093022
## 1-0.5  0.9945312  -3.430338  5.419400 0.8417036

### LM individual alginate by days

##Stat
subset_0 <- subset(data, Alginate == "0")
subset_0.5 <- subset(data, Alginate == "0.5")
subset_1 <- subset(data, Alginate == "1")

##0%

# Fit a general linear model
model <- lm(Dead ~ Day, data = subset_0)

# Perform Tukey post-hoc test
tukey_results <- TukeyHSD(aov(model))

# View the results
print(tukey_results)

##   Tukey multiple comparisons of means
##     95% family-wise confidence level
```

```
##
## Fit: aov(formula = model)
##
## $Day
##            diff        lwr       upr     p adj
## 7-1  -3.645073 -8.2216983 0.9315531 0.1370644
## 14-1  1.260345 -3.4351741 5.9558636 0.7836432
## 14-7  4.905417  0.3287917 9.4820430 0.0339509
```

```
##0.5%

# Fit a general linear model
model <- lm(Dead ~ Day, data = subset_0.5)

# Perform Tukey post-hoc test
tukey_results <- TukeyHSD(aov(model))

# View the results
print(tukey_results)
```

```
##   Tukey multiple comparisons of means
##     95% family-wise confidence level
##
## Fit: aov(formula = model)
##
## $Day
##            diff        lwr       upr     p adj
## 7-1  -8.064796 -13.743765 -2.385827 0.0045102
## 14-1 -9.094303 -14.773272 -3.415334 0.0014781
## 14-7 -1.029507  -6.708476  4.649462 0.8936720
```

```
##1%

# Fit a general linear model
model <- lm(Dead ~ Day, data = subset_1)

# Perform Tukey post-hoc test
tukey_results <- TukeyHSD(aov(model))

# View the results
print(tukey_results)
```

```
##   Tukey multiple comparisons of means
##     95% family-wise confidence level
##
## Fit: aov(formula = model)
##
## $Day
##             diff       lwr       upr     p adj
## 7-1   -4.145439 -12.45044  4.159563 0.4382939
```

```
## 14-1 -10.360634 -18.66564 -2.055633 0.0126079
## 14-7  -6.215196 -14.52020  2.089806 0.1694888

############## ECM area #####################################################
##

### Endometrium
#data input
data <- read.csv("Area_endo.csv")
data$Day <- factor(data$Day)
data$Alginate <- factor(data$Alginate)

# Combine 'Alginate' and 'Cells_content' into a new column
data$Combined = paste(data$Alginate, data$Cells_content, sep = "_")


####Basic Summary Statistics


# Means
means <- aggregate(Area ~Day + Combined, data = data, FUN = "mean")

# SDs

sds <- aggregate(Area ~Day + Combined, data = data, FUN = "sd")

SUM <- merge(means, sds, by = c("Day"))
names(SUM)[3:4] <- c("Mean", "SD")

write.csv(SUM, file = "Summary_Area_endo.csv")


##### plot


png(filename="Area_endo.png",
    width = 12, height = 8, units = "in",
    res = 600)


boxplot(Area ~ Combined * Day,
        data = data,
        cex.axis = 0.9,
        col = viridis(6, alpha = 0.7),  # Ensure 'viridis' library is loaded
        lwd = 0.7,
        ylab = "",
        ylim = c(0,25000000),
        axes = FALSE,
        frame.plot = TRUE,
```

```
        xlab = "")

DAYS <- unique(data$Day)
groups <- 3
numbox <- 6
total <- groups * numbox
xpoints <- seq(median(1:numbox),total,numbox)

# Re-add axes with the correct labels and positions
axis(2, pretty(data$Area), las = 1, cex.axis = 1)
axis(1, labels = unique(data$Day), at = xpoints, las = 1, cex.axis = 1)
abline(v = seq(7, total, by = numbox) - 0.5, lty = "dashed", col = "grey70")

# Adjust 'stripchart' to use 'Combined'
stripchart(Area ~ Combined * Day,  # Update to use Combined
           data = data,
           method = "jitter",
           pch = 19,
           col = "black",
           vertical = TRUE,
           add = TRUE,
           cex = 0.25)

# Title can remain unchanged
title(ylab = expression("Area (um^2)"), line = 3.9)

legend("top", fill = viridis(6), legend = c("0% Cells", "0% No_Cells", "0.5%
Cells", "0.5% No_Cells","1% Cells", "1% No_Cells"), horiz = T, cex = 0.8)

dev.off()

## quartz_off_screen
##                 2

### LM area

##Stat
subset_0NC <- subset(data, Combined == "0_No_cells")
subset_0C <- subset(data, Combined == "0_Cells")
subset_0.5NC <- subset(data, Combined == "0.5_No_cells")
subset_0.5C <- subset(data, Combined == "0.5_Cells")
subset_1NC <- subset(data, Combined == "1_No_cells")
subset_1C <- subset(data, Combined == "1_Cells")

##0NC

# Fit a general linear model
model <- lm(Area ~ Day, data = subset_0NC)
```

```r
# Perform Tukey post-hoc test
tukey_results <- TukeyHSD(aov(model))

# View the results
print(tukey_results)
```

```
##   Tukey multiple comparisons of means
##     95% family-wise confidence level
##
## Fit: aov(formula = model)
##
## $Day
##            diff      lwr     upr     p adj
## 7-1  -504236.1 -2300923 1292451 0.7630943
## 14-1 -203664.7 -2189977 1782647 0.9641477
## 14-7  300571.4 -1496115 2097258 0.9076448
```

```r
##0C

# Fit a general linear model
model <- lm(Area ~ Day, data = subset_0C)

# Perform Tukey post-hoc test
tukey_results <- TukeyHSD(aov(model))

# View the results
print(tukey_results)
```

```
##   Tukey multiple comparisons of means
##     95% family-wise confidence level
##
## Fit: aov(formula = model)
##
## $Day
##            diff       lwr          upr     p adj
## 7-1  -10562869 -12715652  -8410087.03 0.0000000
## 14-1 -12687242 -14910626 -10463857.43 0.0000000
## 14-7  -2124372  -4277154     28410.29 0.0534725
```

```r
##0.5NC

# Fit a general linear model
model <- lm(Area ~ Day, data = subset_0.5NC)

# Perform Tukey post-hoc test
tukey_results <- TukeyHSD(aov(model))

# View the results
print(tukey_results)
```

```
##   Tukey multiple comparisons of means
##     95% family-wise confidence level
##
## Fit: aov(formula = model)
##
## $Day
##           diff        lwr     upr     p adj
## 7-1  475748.7  -749945.0 1701442 0.5998828
## 14-1 634131.7  -720923.6 1989187 0.4796878
## 14-7 158383.0 -1067310.7 1384077 0.9437194
```

```
##0.5C

# Fit a general linear model
model <- lm(Area ~ Day, data = subset_0.5C)

# Perform Tukey post-hoc test
tukey_results <- TukeyHSD(aov(model))

# View the results
print(tukey_results)
```

```
##   Tukey multiple comparisons of means
##     95% family-wise confidence level
##
## Fit: aov(formula = model)
##
## $Day
##           diff       lwr      upr     p adj
## 7-1  -6237048 -10141386 -2332709 0.0018509
## 14-1 -9052625 -13085008 -5020242 0.0000484
## 14-7 -2815577  -6719916  1088761 0.1862042
```

```
##1NC

# Fit a general linear model
model <- lm(Area ~ Day, data = subset_1NC)

# Perform Tukey post-hoc test
tukey_results <- TukeyHSD(aov(model))

# View the results
print(tukey_results)
```

```
##   Tukey multiple comparisons of means
##     95% family-wise confidence level
##
## Fit: aov(formula = model)
##
## $Day
```

```
##            diff        lwr     upr     p adj
## 7-1   855767.2  -927321.7 2638856 0.4624376
## 14-1  696864.2  -916000.5 2309729 0.5328630
## 14-7 -158903.1 -1771767.7 1453962 0.9668477

##1C

# Fit a general linear model
model <- lm(Area ~ Day, data = subset_1C)

# Perform Tukey post-hoc test
tukey_results <- TukeyHSD(aov(model))

# View the results
print(tukey_results)

##   Tukey multiple comparisons of means
##     95% family-wise confidence level
##
## Fit: aov(formula = model)
##
## $Day
##           diff       lwr        upr     p adj
## 7-1   -2752418  -8248888  2744051.5 0.4273906
## 14-1 -8402022 -13723956 -3080088.1 0.0020598
## 14-7 -5649604 -10971538  -327669.7 0.0363698

#### day for different Combined
##Stat
subset_1 <- subset(data, Day == "1")
subset_7 <- subset(data, Day == "7")
subset_14 <- subset(data, Day == "14")


## day 1

# Fit a general linear model
model <- lm(Area ~ Combined, data = subset_1)

# Perform Tukey post-hoc test
tukey_results <- TukeyHSD(aov(model))

# View the results
print(tukey_results)

##   Tukey multiple comparisons of means
##     95% family-wise confidence level
##
## Fit: aov(formula = model)
##
```

```
## $Combined
##                                diff      lwr       upr     p adj
## 0_No_cells-0_Cells       1722568.34 -1393070 4838206.3 0.5637740
## 0.5_Cells-0_Cells        1854174.46 -1261463 4969812.4 0.4843134
## 0.5_No_cells-0_Cells     1257852.40 -1857786 4373490.3 0.8268812
## 1_Cells-0_Cells          -681111.33 -3796749 2434526.6 0.9854118
## 1_No_cells-0_Cells       1192168.21 -1923470 4307806.1 0.8563917
## 0.5_Cells-0_No_cells      131606.11 -2984032 3247244.0 0.9999949
## 0.5_No_cells-0_No_cells  -464715.94 -3580354 2650922.0 0.9975160
## 1_Cells-0_No_cells      -2403679.67 -5519318  711958.3 0.2122223
## 1_No_cells-0_No_cells    -530400.13 -3646038 2585237.8 0.9953665
## 0.5_No_cells-0.5_Cells   -596322.06 -3711960 2519315.9 0.9920219
## 1_Cells-0.5_Cells       -2535285.79 -5650924  580352.1 0.1671544
## 1_No_cells-0.5_Cells     -662006.24 -3777644 2453631.7 0.9871621
## 1_Cells-0.5_No_cells    -1938963.73 -5054602 1176674.2 0.4348588
## 1_No_cells-0.5_No_cells   -65684.19 -3181322 3049953.7 0.9999998
## 1_No_cells-1_Cells       1873279.54 -1242358 4988917.5 0.4730145
```

```
## day 7

# Fit a general linear model
model <- lm(Area ~ Combined, data = subset_7)

# Perform Tukey post-hoc test
tukey_results <- TukeyHSD(aov(model))

# View the results
print(tukey_results)
```

```
##   Tukey multiple comparisons of means
##     95% family-wise confidence level
##
## Fit: aov(formula = model)
##
## $Combined
##                               diff      lwr        upr     p adj
## 0_No_cells-0_Cells      11781201.6  8085404 15476999.0 0.0000000
## 0.5_Cells-0_Cells        6179996.2  2203114 10156878.2 0.0004245
## 0.5_No_cells-0_Cells    12296470.5  8600673 15992267.9 0.0000000
## 1_Cells-0_Cells          7129339.7  3012876 11245803.6 0.0000748
## 1_No_cells-0_Cells      12610804.9  8494341 16727268.7 0.0000000
## 0.5_Cells-0_No_cells    -5601205.4 -9297003 -1905408.0 0.0006121
## 0.5_No_cells-0_No_cells   515268.9 -2876227  3906765.2 0.9974828
## 1_Cells-0_No_cells      -4651861.9 -8497457  -806266.6 0.0095998
## 1_No_cells-0_No_cells     829603.2 -3015992  4675198.6 0.9871613
## 0.5_No_cells-0.5_Cells   6116474.3  2420677  9812271.7 0.0001596
## 1_Cells-0.5_Cells         949343.5 -3167120  5065807.3 0.9826833
## 1_No_cells-0.5_Cells     6430808.6  2314345 10547272.5 0.0003922
## 1_Cells-0.5_No_cells    -5167130.8 -9012726 -1321535.5 0.0029938
```

```
## 1_No_cells-0.5_No_cells   314334.4 -3531261  4159929.7 0.9998761
## 1_No_cells-1_Cells       5481465.2  1230000  9732930.8 0.0048443

## day 14

# Fit a general linear model
#model <- lm(Area ~ Combined, Combined = subset_14)

# Perform Tukey post-hoc test
#tukey_results <- TukeyHSD(aov(model))

# View the results
#print(tukey_results)




### Cortex
#data input
data <- read.csv("Area_ctx.csv")
data$Day <- factor(data$Day)
data$Alginate <- factor(data$Alginate)

# Combine 'Alginate' and 'Cells_content' into a new column
data$Combined = paste(data$Alginate, data$Cells_content, sep = "_")


####Basic Summary Statistics


# Means
means <- aggregate(Area ~Day + Combined, data = data, FUN = "mean")

# SDs

sds <- aggregate(Area ~Day + Combined, data = data, FUN = "sd")

SUM <- merge(means, sds, by = c("Day"))
names(SUM)[3:4] <- c("Mean", "SD")

write.csv(SUM, file = "Summary_Area_ctx.csv")


##### plot


png(filename="Area_ctx.png",
    width = 12, height = 8, units = "in",
```

```
    res = 600)


boxplot(Area ~ Combined * Day,
        data = data,
        cex.axis = 0.9,
        col = viridis(6, alpha = 0.7),  # Ensure 'viridis' library is loaded
        lwd = 0.7,
        ylab = "",
        ylim = c(0,27000000),
        axes = FALSE,
        frame.plot = TRUE,
        xlab = "")

DAYS <- unique(data$Day)
groups <- 3
numbox <- 6
total <- groups * numbox
xpoints <- seq(median(1:numbox),total,numbox)

# Re-add axes with the correct labels and positions
axis(2, pretty(data$Area), las = 1, cex.axis = 1)
axis(1, labels = unique(data$Day), at = xpoints, las = 1, cex.axis = 1)
abline(v = seq(7, total, by = numbox) - 0.5, lty = "dashed", col = "grey70")

# Adjust 'stripchart' to use 'Combined'
stripchart(Area ~ Combined * Day,  # Update to use Combined
           data = data,
           method = "jitter",
           pch = 19,
           col = "black",
           vertical = TRUE,
           add = TRUE,
           cex = 0.25)

# Title can remain unchanged
title(ylab = expression("Area (um^2)"), line = 3.9)

legend("top", fill = viridis(6), legend = c("0% Cells", "0% No_Cells", "0.5%
Cells", "0.5% No_Cells","1% Cells", "1% No_Cells"), horiz = T, cex = 0.8)

dev.off()

## quartz_off_screen
##                 2

### LM area

##Stat
```

```r
subset_0NC <- subset(data, Combined == "0_No_cells")
subset_0C <- subset(data, Combined == "0_Cells")
subset_0.5NC <- subset(data, Combined == "0.5_No_cells")
subset_0.5C <- subset(data, Combined == "0.5_Cells")
subset_1NC <- subset(data, Combined == "1_No_cells")
subset_1C <- subset(data, Combined == "1_Cells")

##0NC

# Fit a general linear model
model <- lm(Area ~ Day, data = subset_0NC)

# Perform Tukey post-hoc test
tukey_results <- TukeyHSD(aov(model))

# View the results
print(tukey_results)

##   Tukey multiple comparisons of means
##     95% family-wise confidence level
##
## Fit: aov(formula = model)
##
## $Day
##           diff      lwr       upr     p adj
## 7-1   -2849525 -6852595 1153545.5 0.1958144
## 14-1 -3753775 -7756845  249295.5 0.0686627
## 14-7  -904250 -4907320 3098820.5 0.8376982

##0C

# Fit a general linear model
model <- lm(Area ~ Day, data = subset_0C)

# Perform Tukey post-hoc test
tukey_results <- TukeyHSD(aov(model))

# View the results
print(tukey_results)

##   Tukey multiple comparisons of means
##     95% family-wise confidence level
##
## Fit: aov(formula = model)
##
## $Day
##             diff      lwr      upr     p adj
## 7-1   -4936666.7 -7878324 -1995009 0.0009152
## 14-1 -5147555.6 -8089213 -2205898 0.0005838
## 14-7  -210888.9 -3152546  2730768 0.9824979
```

```
##0.5NC

# Fit a general linear model
model <- lm(Area ~ Day, data = subset_0.5NC)

# Perform Tukey post-hoc test
tukey_results <- TukeyHSD(aov(model))

# View the results
print(tukey_results)

##   Tukey multiple comparisons of means
##     95% family-wise confidence level
##
## Fit: aov(formula = model)
##
## $Day
##             diff      lwr     upr     p adj
## 7-1  -151666.67 -2168789 1865455 0.9807661
## 14-1 -127111.11 -2144233 1890011 0.9864475
## 14-7   24555.56 -1992567 2041678 0.9994906

##0.5C

# Fit a general linear model
model <- lm(Area ~ Day, data = subset_0.5C)

# Perform Tukey post-hoc test
tukey_results <- TukeyHSD(aov(model))

# View the results
print(tukey_results)

##   Tukey multiple comparisons of means
##     95% family-wise confidence level
##
## Fit: aov(formula = model)
##
## $Day
##           diff       lwr        upr     p adj
## 7-1  -7659000  -9747004 -5570995.6 0.0000000
## 14-1 -9857222 -11945227 -7769217.8 0.0000000
## 14-7 -2198222  -4286227  -110217.8 0.0377282

##1NC

# Fit a general linear model
model <- lm(Area ~ Day, data = subset_1NC)

# Perform Tukey post-hoc test
```

```
tukey_results <- TukeyHSD(aov(model))

# View the results
print(tukey_results)

##   Tukey multiple comparisons of means
##     95% family-wise confidence level
##
## Fit: aov(formula = model)
##
## $Day
##             diff      lwr     upr     p adj
## 7-1  -283888.9 -3491053 2923275 0.9734517
## 14-1 -431222.2 -3638386 2775941 0.9398986
## 14-7 -147333.3 -3354497 3059830 0.9927725

##1C

# Fit a general linear model
model <- lm(Area ~ Day, data = subset_1C)

# Perform Tukey post-hoc test
tukey_results <- TukeyHSD(aov(model))

# View the results
print(tukey_results)

##   Tukey multiple comparisons of means
##     95% family-wise confidence level
##
## Fit: aov(formula = model)
##
## $Day
##              diff      lwr     upr     p adj
## 7-1  -2111333.3 -6190575 1967909 0.4129288
## 14-1 -2657444.4 -6736687 1421798 0.2540332
## 14-7  -546111.1 -4625353 3533131 0.9403992

#### day for different Combined
##Stat
subset_1 <- subset(data, Day == "1")
subset_7 <- subset(data, Day == "7")
subset_14 <- subset(data, Day == "14")


## day 1

# Fit a general linear model
model <- lm(Area ~ Combined, data = subset_1)
```

```
# Perform Tukey post-hoc test
tukey_results <- TukeyHSD(aov(model))

# View the results
print(tukey_results)

##   Tukey multiple comparisons of means
##     95% family-wise confidence level
##
## Fit: aov(formula = model)
##
## $Combined
##                                  diff      lwr      upr     p adj
## 0_No_cells-0_Cells       9083108.33  4849995 13316221 0.0000011
## 0.5_Cells-0_Cells        9059000.00  4952277 13165723 0.0000006
## 0.5_No_cells-0_Cells     9465333.33  5358611 13572056 0.0000002
## 1_Cells-0_Cells         11652333.33  7545611 15759056 0.0000000
## 1_No_cells-0_Cells      11253444.44  7146722 15360167 0.0000000
## 0.5_Cells-0_No_cells      -24108.33 -4257221  4209005 1.0000000
## 0.5_No_cells-0_No_cells   382225.00 -3850888  4615338 0.9997990
## 1_Cells-0_No_cells       2569225.00 -1663888  6802338 0.4738171
## 1_No_cells-0_No_cells    2170336.11 -2062777  6403449 0.6515428
## 0.5_No_cells-0.5_Cells    406333.33 -3700389  4513056 0.9996852
## 1_Cells-0.5_Cells        2593333.33 -1513389  6700056 0.4293166
## 1_No_cells-0.5_Cells     2194444.44 -1912278  6301167 0.6107656
## 1_Cells-0.5_No_cells     2187000.00 -1919723  6293723 0.6142007
## 1_No_cells-0.5_No_cells  1788111.11 -2318612  5894834 0.7871017
## 1_No_cells-1_Cells       -398888.89 -4505612  3707834 0.9997125

## day 7

# Fit a general linear model
model <- lm(Area ~ Combined, data = subset_7)

# Perform Tukey post-hoc test
tukey_results <- TukeyHSD(aov(model))

# View the results
print(tukey_results)

##   Tukey multiple comparisons of means
##     95% family-wise confidence level
##
## Fit: aov(formula = model)
##
## $Combined
##                                diff        lwr        upr     p adj
## 0_No_cells-0_Cells       11170250.0  7203317.2 15137182.8 0.0000000
## 0.5_Cells-0_Cells         6336666.7  2488176.5 10185156.8 0.0001691
## 0.5_No_cells-0_Cells     14250333.3 10401843.2 18098823.5 0.0000000
```

```
## 1_Cells-0_Cells          14477666.7 10629176.5 18326156.8 0.0000000
## 1_No_cells-0_Cells       15906222.2 12057732.1 19754712.4 0.0000000
## 0.5_Cells-0_No_cells     -4833583.3 -8800516.2  -866650.5 0.0088685
## 0.5_No_cells-0_No_cells   3080083.3  -886849.5  7047016.2 0.2119975
## 1_Cells-0_No_cells        3307416.7  -659516.2  7274349.5 0.1520112
## 1_No_cells-0_No_cells     4735972.2   769039.4  8702905.1 0.0109118
## 0.5_No_cells-0.5_Cells    7913666.7  4065176.5 11762156.8 0.0000027
## 1_Cells-0.5_Cells         8141000.0  4292509.9 11989490.1 0.0000015
## 1_No_cells-0.5_Cells      9569555.6  5721065.4 13418045.7 0.0000000
## 1_Cells-0.5_No_cells       227333.3 -3621156.8  4075823.5 0.9999753
## 1_No_cells-0.5_No_cells   1655888.9 -2192601.3  5504379.0 0.7952449
## 1_No_cells-1_Cells        1428555.6 -2419934.6  5277045.7 0.8778295

## day 14

# Fit a general linear model
#model <- lm(Area ~ Combined, Combined = subset_14)

# Perform Tukey post-hoc test
#tukey_results <- TukeyHSD(aov(model))

# View the results
#print(tukey_results)




### Medulla
#data input
data <- read.csv("Area_med.csv")
data$Day <- factor(data$Day)
data$Alginate <- factor(data$Alginate)

# Combine 'Alginate' and 'Cells_content' into a new column
data$Combined = paste(data$Alginate, data$Cells_content, sep = "_")


####Basic Summary Statistics


# Means
means <- aggregate(Area ~Day + Combined, data = data, FUN = "mean")

# SDs

sds <- aggregate(Area ~Day + Combined, data = data, FUN = "sd")

SUM <- merge(means, sds, by = c("Day"))
names(SUM)[3:4] <- c("Mean", "SD")
```

```r
write.csv(SUM, file = "Summary_Area_med.csv")


##### plot


png(filename="Area_med.png",
    width = 12, height = 8, units = "in",
    res = 600)


boxplot(Area ~ Combined * Day,
        data = data,
        cex.axis = 0.9,
        col = viridis(6, alpha = 0.7),  # Ensure 'viridis' library is loaded
        lwd = 0.7,
        ylab = "",
        ylim = c(0,25000000),
        axes = FALSE,
        frame.plot = TRUE,
        xlab = "")

DAYS <- unique(data$Day)
groups <- 3
numbox <- 6
total <- groups * numbox
xpoints <- seq(median(1:numbox),total,numbox)

# Re-add axes with the correct labels and positions
axis(2, pretty(data$Area), las = 1, cex.axis = 1)
axis(1, labels = unique(data$Day), at = xpoints, las = 1, cex.axis = 1)
abline(v = seq(7, total, by = numbox) - 0.5, lty = "dashed", col = "grey70")

# Adjust 'stripchart' to use 'Combined'
stripchart(Area ~ Combined * Day,  # Update to use Combined
           data = data,
           method = "jitter",
           pch = 19,
           col = "black",
           vertical = TRUE,
           add = TRUE,
           cex = 0.25)

# Title can remain unchanged
title(ylab = expression("Area (um^2)"), line = 3.9)

legend("top", fill = viridis(6), legend = c("0% Cells", "0% No_Cells", "0.5%
Cells", "0.5% No_Cells","1% Cells", "1% No_Cells"), horiz = T, cex = 0.8)
```

```r
dev.off()

## quartz_off_screen
##                 2

### LM area

##Stat
subset_0NC <- subset(data, Combined == "0_No_cells")
subset_0C <- subset(data, Combined == "0_Cells")
subset_0.5NC <- subset(data, Combined == "0.5_No_cells")
subset_0.5C <- subset(data, Combined == "0.5_Cells")
subset_1NC <- subset(data, Combined == "1_No_cells")
subset_1C <- subset(data, Combined == "1_Cells")

##0NC

# Fit a general linear model
model <- lm(Area ~ Day, data = subset_0NC)

# Perform Tukey post-hoc test
tukey_results <- TukeyHSD(aov(model))

# View the results
print(tukey_results)

##   Tukey multiple comparisons of means
##     95% family-wise confidence level
##
## Fit: aov(formula = model)
##
## $Day
##           diff      lwr        upr     p adj
## 7-1  -2490111 -4628042  -352180.4 0.0202908
## 14-1 -3813000 -5950931 -1675069.3 0.0004726
## 14-7 -1322889 -3460820   815041.8 0.2883832

##0C

# Fit a general linear model
model <- lm(Area ~ Day, data = subset_0C)

# Perform Tukey post-hoc test
tukey_results <- TukeyHSD(aov(model))

# View the results
print(tukey_results)
```

```
##   Tukey multiple comparisons of means
##     95% family-wise confidence level
##
## Fit: aov(formula = model)
##
## $Day
##           diff       lwr      upr     p adj
## 7-1  -7850889 -10780364 -4921413 0.0000019
## 14-1 -9265667 -12195142 -6336191 0.0000001
## 14-7 -1414778  -4344253  1514698 0.4611717
```

```
##0.5NC

# Fit a general linear model
model <- lm(Area ~ Day, data = subset_0.5NC)

# Perform Tukey post-hoc test
tukey_results <- TukeyHSD(aov(model))

# View the results
print(tukey_results)
```

```
##   Tukey multiple comparisons of means
##     95% family-wise confidence level
##
## Fit: aov(formula = model)
##
## $Day
##             diff      lwr     upr     p adj
## 7-1   -63111.11 -1553567 1427345 0.9938559
## 14-1 -299444.44 -1789901 1191012 0.8711767
## 14-7 -236333.33 -1726789 1254123 0.9174947
```

```
##0.5C

# Fit a general linear model
model <- lm(Area ~ Day, data = subset_0.5C)

# Perform Tukey post-hoc test
tukey_results <- TukeyHSD(aov(model))

# View the results
print(tukey_results)
```

```
##   Tukey multiple comparisons of means
##     95% family-wise confidence level
##
## Fit: aov(formula = model)
##
## $Day
```

```
##            diff       lwr         upr      p adj
## 7-1   -3922333  -7318722  -525944.9 0.0214564
## 14-1 -11139111 -14535500 -7742722.6 0.0000001
## 14-7  -7216778 -10613166 -3820389.3 0.0000553
```

```r
##1NC

# Fit a general linear model
model <- lm(Area ~ Day, data = subset_1NC)

# Perform Tukey post-hoc test
tukey_results <- TukeyHSD(aov(model))

# View the results
print(tukey_results)
```

```
##   Tukey multiple comparisons of means
##     95% family-wise confidence level
##
## Fit: aov(formula = model)
##
## $Day
##            diff      lwr     upr     p adj
## 7-1   387444.4 -1864481 2639370 0.9031570
## 14-1 1044013.9 -1277218 3365245 0.5080138
## 14-7  656569.4 -1664662 2977801 0.7610421
```

```r
##1C

# Fit a general linear model
model <- lm(Area ~ Day, data = subset_1C)

# Perform Tukey post-hoc test
tukey_results <- TukeyHSD(aov(model))

# View the results
print(tukey_results)
```

```
##   Tukey multiple comparisons of means
##     95% family-wise confidence level
##
## Fit: aov(formula = model)
##
## $Day
##             diff      lwr      upr     p adj
## 7-1    239777.8 -1127313  1606869 0.9000877
## 14-1 -2486333.3 -3853424 -1119242 0.0003787
## 14-7 -2726111.1 -4093202 -1359020 0.0001255
```

```
#### day for different Combined
##Stat
subset_1 <- subset(data, Day == "1")
subset_7 <- subset(data, Day == "7")
subset_14 <- subset(data, Day == "14")


## day 1

# Fit a general linear model
model <- lm(Area ~ Combined, data = subset_1)

# Perform Tukey post-hoc test
tukey_results <- TukeyHSD(aov(model))

# View the results
print(tukey_results)

##   Tukey multiple comparisons of means
##     95% family-wise confidence level
##
## Fit: aov(formula = model)
##
## $Combined
##                                diff      lwr      upr     p adj
## 0_No_cells-0_Cells       5705888.89  2667861  8743917 0.0000160
## 0.5_Cells-0_Cells        7154444.44  4116417 10192472 0.0000001
## 0.5_No_cells-0_Cells     6789222.22  3751194  9827250 0.0000004
## 1_Cells-0_Cells          5587888.89  2549861  8625917 0.0000237
## 1_No_cells-0_Cells       5784555.56  2746528  8822583 0.0000122
## 0.5_Cells-0_No_cells     1448555.56 -1589472  4486583 0.7178155
## 0.5_No_cells-0_No_cells  1083333.33 -1954694  4121361 0.8952003
## 1_Cells-0_No_cells       -118000.00 -3156028  2920028 0.9999969
## 1_No_cells-0_No_cells      78666.67 -2959361  3116694 0.9999996
## 0.5_No_cells-0.5_Cells   -365222.22 -3403250  2672806 0.9991916
## 1_Cells-0.5_Cells       -1566555.56 -4604583  1471472 0.6468256
## 1_No_cells-0.5_Cells    -1369888.89 -4407917  1668139 0.7624297
## 1_Cells-0.5_No_cells    -1201333.33 -4239361  1836694 0.8470667
## 1_No_cells-0.5_No_cells -1004666.67 -4042694  2033361 0.9216181
## 1_No_cells-1_Cells        196666.67 -2841361  3234694 0.9999613

## day 7

# Fit a general linear model
model <- lm(Area ~ Combined, data = subset_7)

# Perform Tukey post-hoc test
tukey_results <- TukeyHSD(aov(model))
```

```
# View the results
print(tukey_results)

##   Tukey multiple comparisons of means
##     95% family-wise confidence level
##
## Fit: aov(formula = model)
##
## $Combined
##                                diff         lwr      upr     p adj
## 0_No_cells-0_Cells      11066666.67  8013198.52 14120135 0.0000000
## 0.5_Cells-0_Cells       11083000.00  8029531.85 14136468 0.0000000
## 0.5_No_cells-0_Cells    14577000.00 11523531.85 17630468 0.0000000
## 1_Cells-0_Cells         13678555.56 10625087.41 16732024 0.0000000
## 1_No_cells-0_Cells      14022888.89 10969420.74 17076357 0.0000000
## 0.5_Cells-0_No_cells       16333.33 -3037134.81  3069801 1.0000000
## 0.5_No_cells-0_No_cells  3510333.33   456865.19  6563801 0.0156795
## 1_Cells-0_No_cells       2611888.89  -441579.26  5665357 0.1333042
## 1_No_cells-0_No_cells    2956222.22   -97245.92  6009690 0.0628594
## 0.5_No_cells-0.5_Cells   3494000.00   440531.85  6547468 0.0163790
## 1_Cells-0.5_Cells        2595555.56  -457912.59  5649024 0.1378065
## 1_No_cells-0.5_Cells     2939888.89  -113579.26  5993357 0.0652786
## 1_Cells-0.5_No_cells     -898444.44 -3951912.59  2155024 0.9510767
## 1_No_cells-0.5_No_cells  -554111.11 -3607579.26  2499357 0.9942543
## 1_No_cells-1_Cells        344333.33 -2709134.81  3397801 0.9994075

## day 14

# Fit a general linear model
#model <- lm(Area ~ Combined, Combined = subset_14)

# Perform Tukey post-hoc test
#tukey_results <- TukeyHSD(aov(model))

# View the results
#print(tukey_results)



### Oviduct
#data input
data <- read.csv("Area_ovi.csv")
data$Day <- factor(data$Day)
data$Alginate <- factor(data$Alginate)

# Combine 'Alginate' and 'Cells_content' into a new column
data$Combined = paste(data$Alginate, data$Cells_content, sep = "_")
```

```r
####Basic Summary Statistics


# Means
means <- aggregate(Area ~Day + Combined, data = data, FUN = "mean")

# SDs

sds <- aggregate(Area ~Day + Combined, data = data, FUN = "sd")

SUM <- merge(means, sds, by = c("Day"))
names(SUM)[3:4] <- c("Mean", "SD")

write.csv(SUM, file = "Summary_Area_ovi.csv")


##### plot


png(filename="Area_ovi.png",
    width = 12, height = 8, units = "in",
    res = 600)


boxplot(Area ~ Combined * Day,
        data = data,
        cex.axis = 0.9,
        col = viridis(6, alpha = 0.7),  # Ensure 'viridis' library is loaded
        lwd = 0.7,
        ylab = "",
        ylim = c(0,25000000),
        axes = FALSE,
        frame.plot = TRUE,
        xlab = "")

DAYS <- unique(data$Day)
groups <- 3
numbox <- 6
total <- groups * numbox
xpoints <- seq(median(1:numbox),total,numbox)

# Re-add axes with the correct labels and positions
axis(2, pretty(data$Area), las = 1, cex.axis = 1)
axis(1, labels = unique(data$Day), at = xpoints, las = 1, cex.axis = 1)
abline(v = seq(7, total, by = numbox) - 0.5, lty = "dashed", col = "grey70")

# Adjust 'stripchart' to use 'Combined'
stripchart(Area ~ Combined * Day,  # Update to use Combined
```

```
           data = data,
           method = "jitter",
           pch = 19,
           col = "black",
           vertical = TRUE,
           add = TRUE,
           cex = 0.25)

# Title can remain unchanged
title(ylab = expression("Area (um^2)"), line = 3.9)

legend("top", fill = viridis(6), legend = c("0% Cells", "0% No_Cells", "0.5%
Cells", "0.5% No_Cells","1% Cells", "1% No_Cells"), horiz = T, cex = 0.8)

dev.off()

## quartz_off_screen
##                 2

### LM area

##Stat
subset_0NC <- subset(data, Combined == "0_No_cells")
subset_0C <- subset(data, Combined == "0_Cells")
subset_0.5NC <- subset(data, Combined == "0.5_No_cells")
subset_0.5C <- subset(data, Combined == "0.5_Cells")
subset_1NC <- subset(data, Combined == "1_No_cells")
subset_1C <- subset(data, Combined == "1_Cells")

##0NC

# Fit a general linear model
model <- lm(Area ~ Day, data = subset_0NC)

# Perform Tukey post-hoc test
tukey_results <- TukeyHSD(aov(model))

# View the results
print(tukey_results)

##   Tukey multiple comparisons of means
##     95% family-wise confidence level
##
## Fit: aov(formula = model)
##
## $Day
##             diff      lwr        upr     p adj
## 7-1   -800916.7 -2645548 1043714.64 0.5327273
## 14-1 -1762663.6 -3607295   81967.75 0.0629903
## 14-7  -961746.9 -2806378  882884.42 0.4077962
```

```r
##0C

# Fit a general linear model
model <- lm(Area ~ Day, data = subset_0C)

# Perform Tukey post-hoc test
tukey_results <- TukeyHSD(aov(model))

# View the results
print(tukey_results)

##   Tukey multiple comparisons of means
##     95% family-wise confidence level
##
## Fit: aov(formula = model)
##
## $Day
##             diff      lwr        upr     p adj
## 7-1  -3225401.8 -3976211 -2474592.1 0.0000000
## 14-1 -3624766.3 -4375576 -2873956.7 0.0000000
## 14-7  -399364.6 -1150174   351445.1 0.3937253

##0.5NC

# Fit a general linear model
model <- lm(Area ~ Day, data = subset_0.5NC)

# Perform Tukey post-hoc test
tukey_results <- TukeyHSD(aov(model))

# View the results
print(tukey_results)

##   Tukey multiple comparisons of means
##     95% family-wise confidence level
##
## Fit: aov(formula = model)
##
## $Day
##            diff      lwr     upr     p adj
## 7-1  -236456.1 -3054484 2581572 0.9761089
## 14-1 -446847.2 -3264875 2371181 0.9174918
## 14-7 -210391.1 -3028419 2607637 0.9810338

##0.5C

# Fit a general linear model
model <- lm(Area ~ Day, data = subset_0.5C)

# Perform Tukey post-hoc test
```

```
tukey_results <- TukeyHSD(aov(model))

# View the results
print(tukey_results)

##   Tukey multiple comparisons of means
##     95% family-wise confidence level
##
## Fit: aov(formula = model)
##
## $Day
##           diff      lwr         upr     p adj
## 7-1   -1647904 -3522869   227060.7 0.0924797
## 14-1 -5627357 -7502322 -3752392.0 0.0000003
## 14-7 -3979453 -5854418 -2104487.7 0.0000561

##1NC

# Fit a general linear model
model <- lm(Area ~ Day, data = subset_1NC)

# Perform Tukey post-hoc test
tukey_results <- TukeyHSD(aov(model))

# View the results
print(tukey_results)

##   Tukey multiple comparisons of means
##     95% family-wise confidence level
##
## Fit: aov(formula = model)
##
## $Day
##            diff      lwr     upr     p adj
## 7-1   -332342.1 -2650007 1985323 0.9319541
## 14-1 -201033.2 -2518698 2116632 0.9744924
## 14-7  131308.9 -2186356 2448974 0.9890296

##1C

# Fit a general linear model
model <- lm(Area ~ Day, data = subset_1C)

# Perform Tukey post-hoc test
tukey_results <- TukeyHSD(aov(model))

# View the results
print(tukey_results)
```

```
##   Tukey multiple comparisons of means
##     95% family-wise confidence level
##
## Fit: aov(formula = model)
##
## $Day
##             diff      lwr     upr     p adj
## 7-1  -137154.89 -2990064 2715754 0.9920875
## 14-1 -218116.56 -3071025 2634792 0.9801209
## 14-7  -80961.67 -2933870 2771947 0.9972351
```

```
#### day for different Combined
##Stat
subset_1 <- subset(data, Day == "1")
subset_7 <- subset(data, Day == "7")
subset_14 <- subset(data, Day == "14")


## day 1

# Fit a general linear model
model <- lm(Area ~ Combined, data = subset_1)

# Perform Tukey post-hoc test
tukey_results <- TukeyHSD(aov(model))

# View the results
print(tukey_results)
```

```
##   Tukey multiple comparisons of means
##     95% family-wise confidence level
##
## Fit: aov(formula = model)
##
## $Combined
##                                  diff         lwr         upr     p adj
## 0_No_cells-0_Cells         9009950.44   6403781.9 11616118.9 0.0000000
## 0.5_Cells-0_Cells         11138781.22   8532612.7 13744949.7 0.0000000
## 0.5_No_cells-0_Cells       9836059.67   7229891.2 12442228.2 0.0000000
## 1_Cells-0_Cells           13527019.33  10920850.8 16133187.8 0.0000000
## 1_No_cells-0_Cells        11209239.56   8603071.1 13815408.1 0.0000000
## 0.5_Cells-0_No_cells       2128830.78   -477337.7  4734999.3 0.1684551
## 0.5_No_cells-0_No_cells     826109.22  -1780059.3  3432277.7 0.9337465
## 1_Cells-0_No_cells         4517068.89   1910900.4  7123237.4 0.0000695
## 1_No_cells-0_No_cells      2199289.11   -406879.4  4805457.6 0.1431358
## 0.5_No_cells-0.5_Cells    -1302721.56  -3908890.1  1303446.9 0.6761385
## 1_Cells-0.5_Cells          2388238.11   -217930.4  4994406.6 0.0898246
## 1_No_cells-0.5_Cells         70458.33  -2535710.2  2676626.8 0.9999995
## 1_Cells-0.5_No_cells       3690959.67   1084791.2  6297128.2 0.0015111
```

```
## 1_No_cells-0.5_No_cells  1373179.89 -1232988.6  3979348.4 0.6256775
## 1_No_cells-1_Cells      -2317779.78 -4923948.3   288388.7 0.1073835

## day 7

# Fit a general linear model
model <- lm(Area ~ Combined, data = subset_7)

# Perform Tukey post-hoc test
tukey_results <- TukeyHSD(aov(model))

# View the results
print(tukey_results)

##   Tukey multiple comparisons of means
##     95% family-wise confidence level
##
## Fit: aov(formula = model)
##
## $Combined
##                                diff         lwr        upr     p adj
## 0_No_cells-0_Cells      11434435.6  8680227.81 14188643.3 0.0000000
## 0.5_Cells-0_Cells       12716278.7  9962070.92 15470486.4 0.0000000
## 0.5_No_cells-0_Cells    12825005.3 10070797.58 15579213.1 0.0000000
## 1_Cells-0_Cells         16615266.2 13861058.47 19369474.0 0.0000000
## 1_No_cells-0_Cells      14102299.2 11348091.47 16856507.0 0.0000000
## 0.5_Cells-0_No_cells     1281843.1 -1472364.64  4036050.9 0.7377765
## 0.5_No_cells-0_No_cells  1390569.8 -1363637.97  4144777.5 0.6668590
## 1_Cells-0_No_cells       5180830.7  2426622.92  7935038.4 0.0000155
## 1_No_cells-0_No_cells    2667863.7   -86344.08  5422071.4 0.0626383
## 0.5_No_cells-0.5_Cells    108726.7 -2645481.08  2862934.4 0.9999967
## 1_Cells-0.5_Cells        3898987.6  1144779.81  6653195.3 0.0015194
## 1_No_cells-0.5_Cells     1386020.6 -1368187.19  4140228.3 0.6699135
## 1_Cells-0.5_No_cells     3790260.9  1036053.14  6544468.6 0.0021866
## 1_No_cells-0.5_No_cells  1277293.9 -1476913.86  4031501.6 0.7406290
## 1_No_cells-1_Cells      -2512967.0 -5267174.75   241240.7 0.0922421

## day 14

# Fit a general linear model
#model <- lm(Area ~ Combined, Combined = subset_14)

# Perform Tukey post-hoc test
#tukey_results <- TukeyHSD(aov(model))

# View the results
#print(tukey_results)

#######Porosity  Printing ###################################################
#
```

```
#data input
data <- read.csv("Porosity_printing.csv")


####     Basic Summary Statistics

# Means
means <- aggregate(porosity ~ Tissue, data = data, FUN = "mean")

# SDs

sds <- aggregate(porosity ~ Tissue, data = data, FUN = "sd")

SUM <- merge(means, sds, by = c("Tissue"))
names(SUM)[2:3] <- c("Mean", "SD")

write.csv(SUM, file = "Summary_Porosity_printing.csv")

### Plot




png(filename="Porosity_printing.png",
    width = 5, height = 5, units = "in",
    res = 600)


boxplot(porosity ~ Tissue,
        data = data,
        cex.axis = 0.9,
        col = viridis(8, alpha = 0.7),
        lwd = 0.7,
        ylab = "",
        ylim = c(0,100),
        axes = TRUE,
        frame.plot = TRUE,
        xlab ="")

stripchart(data$porosity ~ data$Tissue,               # Data
           method = "jitter", # Random noise
           pch = 19,          # Pch symbols
           col = "black",           # Color of the symbol
           vertical = TRUE,   # Vertical mode
           add = TRUE,
```

```
          cex = 0.25)        # Add it over

title(ylab = expression("Estimated Porosity (%)"), line = 2.2)


dev.off()

## quartz_off_screen
##                 2

##Stats

# Fit a general linear model
model <- lm(porosity ~ Tissue, data = data)

# Perform Tukey post-hoc test
tukey_results <- TukeyHSD(aov(model))

# View the results
print(tukey_results)

##   Tukey multiple comparisons of means
##     95% family-wise confidence level
##
## Fit: aov(formula = model)
##
## $Tissue
##                                          diff        lwr        upr     p a
dj
## Cortex tissue-Cortex                37.688179  22.671237  46.705122 0.00000
00
## Endometrium-Cortex                   6.931850  -5.397273  19.260973 0.64966
64
## Endometrium tissue-Cortex           39.185546  26.856423  51.514669 0.00000
00
## Medulla-Cortex                      10.861972  -1.467151  23.191094 0.12417
83
## Medulla tissue-Cortex               37.613042  25.596100  49.629985 0.00000
00
## Oviduct-Cortex                      14.939689   2.610567  27.268812 0.00738
02
## Oviduct tissue-Cortex               28.531055  16.775698  40.286413 0.00000
00
## Endometrium-Cortex tissue          -27.756330 -39.773272 -15.739387 0.00000
00
## Endometrium tissue-Cortex tissue     4.497367  -7.519576  16.514309 0.93755
51
## Medulla-Cortex tissue              -23.826208 -35.843150 -11.809265 0.00000
10
## Medulla tissue-Cortex tissue         2.924863  -8.771570  14.621296 0.99353
```

```
09
## Oviduct-Cortex tissue             -19.748490 -31.765432  -7.731548 0.00006
63
## Oviduct tissue-Cortex tissue       -6.157124 -17.584637   5.270389 0.69670
88
## Endometrium tissue-Endometrium     32.253696  19.924573  44.582819 0.00000
00
## Medulla-Endometrium                 3.930122  -8.399001  16.259244 0.97338
62
## Medulla tissue-Endometrium         30.681193  18.664250  42.698135 0.00000
00
## Oviduct-Endometrium                 8.007840  -4.321283  20.336962 0.46926
82
## Oviduct tissue-Endometrium         21.599205   9.843848  33.354563 0.00000
63
## Medulla-Endometrium tissue        -28.323574 -40.652697 -15.994452 0.00000
00
## Medulla tissue-Endometrium tissue  -1.572503 -13.589446  10.444439 0.99990
19
## Oviduct-Endometrium tissue        -24.245857 -36.574979 -11.916734 0.00000
13
## Oviduct tissue-Endometrium tissue -10.654491 -22.409848   1.100867 0.10348
44
## Medulla tissue-Medulla             26.751071  14.734129  38.768013 0.00000
00
## Oviduct-Medulla                     4.077718  -8.251405  16.406840 0.96740
87
## Oviduct tissue-Medulla             17.669084   5.913726  29.424441 0.00033
61
## Oviduct-Medulla tissue            -22.673353 -34.690295 -10.656411 0.00000
34
## Oviduct tissue-Medulla tissue      -9.081987 -20.509500   2.345526 0.21939
01
## Oviduct tissue-Oviduct             13.591366   1.836009  25.346723 0.01258
66

#######Size  Printing ####################################################



#data input
data <- read.csv("Diameter_printing.csv")


####     Basic Summary Statistics

## Diameter
# Means
means <- aggregate(Diameter ~ Tissue, data = data, FUN = "mean")
```

```
# SDs

sds <- aggregate(Diameter ~ Tissue, data = data, FUN = "sd")

SUM <- merge(means, sds, by = c("Tissue"))
names(SUM)[2:3] <- c("Mean", "SD")

write.csv(SUM, file = "Summary_Diameter_printing.csv")

## Height
# Means
means <- aggregate(Height ~ Tissue, data = data, FUN = "mean")

# SDs

sds <- aggregate(Height ~ Tissue, data = data, FUN = "sd")

SUM <- merge(means, sds, by = c("Tissue"))
names(SUM)[2:3] <- c("Mean", "SD")

write.csv(SUM, file = "Summary_Height_printing.csv")


### Plot

png(filename="Size_printing.png",
    width = 7, height = 4, units = "in",
    res = 600)




# Generate the plot
ggplot(data, aes(x = Diameter, y = Height, color = Tissue)) +
  geom_point() +  # Add points
  geom_rect(aes(xmin = 4.25, xmax = 5.75, ymin = 2.55, ymax = 3.45), color =
"grey", fill = NA, linetype = "dashed") +  # Add dashed rectangle for 15% ran
ge
  geom_rect(aes(xmin = 4.5, xmax = 5.5, ymin = 2.7, ymax = 3.3), color = "gre
y", fill = NA, linetype = "dashed") +  # Add dashed rectangle for 10% range
  scale_color_viridis(discrete = TRUE, option = "D", alpha = 0.8) +  # Use vi
ridis colors with 80% opacity
  theme_minimal() +  # Minimal theme
  theme(panel.grid.major = element_blank(),  # Remove major grid
        panel.grid.minor = element_blank(),  # Remove minor grid
        panel.background = element_blank(),  # No background
        panel.border = element_rect(colour = "black", fill = NA, size = 1)) +
```

```
# Add plot border
  scale_x_continuous(breaks = seq(4, 6, by = 0.2), limits = c(4, 6)) +  # X-a
xis breaks and limits
  scale_y_continuous(breaks = seq(2, 4.6, by = 0.2), limits = c(2, 4.6))   #
Y-axis breaks and limits

## Warning: The `size` argument of `element_rect()` is deprecated as of ggplo
t2 3.4.0.
## ℹ Please use the `linewidth` argument instead.
## This warning is displayed once every 8 hours.
## Call `lifecycle::last_lifecycle_warnings()` to see where this warning was
## generated.

dev.off()

## quartz_off_screen
##                 2

# Calculate the percentage of samples within the specified range for each tis
sue type
range_data <- data %>%
  filter(Diameter >= 4.25, Diameter <= 5.75, Height >= 2.55, Height <= 3.45)
%>%
  group_by(Tissue) %>%
  summarise(Count_in_range = n())

total_data <- data %>%
  group_by(Tissue) %>%
  summarise(Total_count = n())

percentage_data15 <- left_join(range_data, total_data, by = "Tissue") %>%
  mutate(Percentage = (Count_in_range / Total_count) * 100)

print(percentage_data15)

## # A tibble: 4 × 4
##   Tissue      Count_in_range Total_count Percentage
##   <chr>                <int>       <int>      <dbl>
## 1 Cortex                  12          12      100
## 2 Endometrium              6           9       66.7
## 3 Medulla                 11          12       91.7
## 4 Oviduct                  6          12       50

# Calculate the percentage of samples within the specified range for each tis
sue type
range_data <- data %>%
  filter(Diameter >= 4.5, Diameter <= 5.5, Height >= 2.7, Height <= 3.3) %>%
  group_by(Tissue) %>%
  summarise(Count_in_range = n())

total_data <- data %>%
```

```
  group_by(Tissue) %>%
  summarise(Total_count = n())

percentage_data30 <- left_join(range_data, total_data, by = "Tissue") %>%
  mutate(Percentage = (Count_in_range / Total_count) * 100)

print(percentage_data30)

## # A tibble: 4 × 4
##   Tissue      Count_in_range Total_count Percentage
##   <chr>                <int>       <int>      <dbl>
## 1 Cortex                  11          12       91.7
## 2 Endometrium              6           9       66.7
## 3 Medulla                 11          12       91.7
## 4 Oviduct                  4          12       33.3

#######Viability  Printing ##################################################
##



#data input

#data input
data <- read.csv("Live_Dead_printing.csv")
data$Day <- factor(data$Day)
data$Alginate <- factor(data$Tissue)


####Basic Summary Statistics


# Means
means <- aggregate(Dead ~Day + Tissue, data = data, FUN = "mean")

# SDs

sds <- aggregate(Dead ~Day + Tissue, data = data, FUN = "sd")

SUM <- merge(means, sds, by = c("Day"))
names(SUM)[3:4] <- c("Mean", "SD")

write.csv(SUM, file = "Summary_Viability_printing.csv")


##### plot


png(filename="Cell_viability_printing.png",
```

```
    width = 7, height = 5, units = "in",
    res = 600)


boxplot(Dead ~ Day*Tissue,
        data = data,
        cex.axis = 0.9,
        col = viridis(2, alpha = 0.7),
        lwd = 0.7,
        ylab = "",
        ylim = c(0,20),
        axes = FALSE,
        frame.plot = TRUE,
        xlab ="")

TISSUE <- unique(data$Tissue)
groups <- 4
numbox <- 2
total <- groups * numbox
xpoints <- seq(median(1:numbox),total,numbox)

axis(2, pretty(data$Dead), las = 1, cex.axis = 1)
axis(1, labels = TISSUE, at = xpoints,las = 1, cex.axis = 1)
abline(v = 2.5, lty = "dashed", col = "grey70")
abline(v = 4.5, lty = "dashed", col = "grey70")
abline(v = 6.5, lty = "dashed", col = "grey70")


stripchart(data$Dead ~ data$Day*data$Tissue,               # Data
           method = "jitter", # Random noise
           pch = 19,          # Pch symbols
           col = "black",           # Color of the symbol
           vertical = TRUE,   # Vertical mode
           add = TRUE,
           cex = 0.25)        # Add it over


title(ylab = expression("Dead cells (%)"), line = 2.2)
legend("top", fill = viridis(2), legend = c("Day 1", "Day 7"), horiz = T, cex
= 0.8)
dev.off()

## quartz_off_screen
##                 2

### LM individual days by Tissue

##Stat
subset_Ovi <- subset(data, Tissue == "Oviduct")
```

```
subset_Endo <- subset(data, Tissue == "Endometrium")
subset_Cort <- subset(data, Tissue == "Cortex")
subset_Medu <- subset(data, Tissue == "Medulla")


##Ovi

# Fit a general linear model
model <- lm(Dead ~ Day, data = subset_Ovi)

# Perform Tukey post-hoc test
tukey_results <- TukeyHSD(aov(model))

# View the results
print(tukey_results)
```

```
##   Tukey multiple comparisons of means
##     95% family-wise confidence level
##
## Fit: aov(formula = model)
##
## $Day
##          diff       lwr       upr     p adj
## 7-1 -3.309939 -7.190609 0.5707313 0.0894197
```

```
##Endo

# Fit a general linear model
model <- lm(Dead ~ Day, data = subset_Endo)

# Perform Tukey post-hoc test
tukey_results <- TukeyHSD(aov(model))

# View the results
print(tukey_results)
```

```
##   Tukey multiple comparisons of means
##     95% family-wise confidence level
##
## Fit: aov(formula = model)
##
## $Day
##          diff       lwr       upr     p adj
## 7-1 -4.266144 -6.875665 -1.656623 0.0031862
```

```
##Cortex

# Fit a general linear model
model <- lm(Dead ~ Day, data = subset_Cort)
```

```
# Perform Tukey post-hoc test
tukey_results <- TukeyHSD(aov(model))

# View the results
print(tukey_results)

##   Tukey multiple comparisons of means
##     95% family-wise confidence level
##
## Fit: aov(formula = model)
##
## $Day
##          diff       lwr       upr    p adj
## 7-1 -9.912491 -13.13958 -6.685399 7.2e-06

##Medulla

# Fit a general linear model
model <- lm(Dead ~ Day, data = subset_Medu)

# Perform Tukey post-hoc test
tukey_results <- TukeyHSD(aov(model))

# View the results
print(tukey_results)

##   Tukey multiple comparisons of means
##     95% family-wise confidence level
##
## Fit: aov(formula = model)
##
## $Day
##          diff       lwr       upr     p adj
## 7-1 -4.897521 -8.675528 -1.119515 0.0153329

################## Proteomics #################################################

setwd("~/Desktop/LMU/Students/Eduardo Ribes/Chapter 1 manuscript/Proteomics")

# load the required packages
library(VennDiagram)

## Loading required package: grid

## Loading required package: futile.logger

##
## Attaching package: 'VennDiagram'
```

```
## The following object is masked from 'package:ggpubr':
##
##     rotate
```

```
library(dplyr)
library(tidyr)
library(ggplot2)
library(viridis)
library(multcomp)


################## All analysis

# Read the file
gene_data <- read.csv("Venn_dia.csv", header = TRUE)

# Extract the genes from each column
endometrium_genes <- gene_data$Endometrium
cortex_genes <- gene_data$Cortex
medulla_genes <- gene_data$Medulla
oviduct_genes <- gene_data$Oviduct

# Create the Venn diagram with adjusted parameters
venn.plot <- venn.diagram(
  x = list(Endometrium = endometrium_genes,
           Cortex = cortex_genes,
           Medulla = medulla_genes,
           Oviduct = oviduct_genes),
  category.names = c("Endometrium", "Cortex", "Medulla", "Oviduct"),
  filename = NULL,
  col = viridis(4),
  cat.cex = 0.5,  # Adjust category label size
  cat.fontface = "bold",  # Make category labels bold
  cat.dist = 0.4,  # Adjust distance between category labels and diagram
  margin = 0.1,  # Adjust margin around the plot
  cex = 0.6,  # Adjust overall text size
  category.cex = 0.6,  # Adjust category label text size
  lwd = 1  # Adjust line width for borders
)

# Save the Venn diagram as a high-resolution PNG file with adjusted parameter
s
png("venn_diagram.png", width = 800, height = 600, res = 300)  # Adjust width
, height, and resolution as needed
grid.draw(venn.plot)
dev.off()
```

```
## quartz_off_screen
##                 2
```

```
############## OVIDUCT

# Clean data

# Read your data into a data frame or data table
data <- read.csv("Oviduct_all_count.csv")

# Remove rows with more than 5 NA values
clean_data <- data %>%
  filter(rowSums(is.na(.)) <= 5)

# Replace remaining NA values with 0 in all columns
clean_data <- clean_data %>%
  mutate_all(funs(replace(., is.na(.), 0)))

## Warning: `funs()` was deprecated in dplyr 0.8.0.
## ℹ Please use a list of either functions or lambdas:
##
## # Simple named list: list(mean = mean, median = median)
##
## # Auto named with `tibble::lst()`: tibble::lst(mean, median)
##
## # Using lambdas list(~ mean(., trim = .2), ~ median(., na.rm = TRUE))
## Call `lifecycle::last_lifecycle_warnings()` to see where this warning was
## generated.

# Save the cleaned data to a new file
write.csv(clean_data, "cleaned_data_oviduct.csv", row.names = FALSE)

### Matrisome categorization

# Read your first data file into a data frame or data table
data <- read.csv("cleaned_data_oviduct.csv")

# Read your second dataset into another data frame or data table
matrisome_info <- read.csv("Bt_Matrisome.csv")

# Perform a left join based on the "Genes" column
joined_data <- left_join(data, matrisome_info, by = "Genes")

# Save the joined data to a new file
write.csv(joined_data, "oviduct_matrisome.csv", row.names = FALSE)

# Remove rows where either Matrisome_Division or Matrisome_Category have NA v
alues
cleaned_joined_data <- joined_data %>%
  filter(!is.na(Matrisome_Division) & !is.na(Matrisome_Category))

# Save the cleaned data to a new file
```

```r
write.csv(cleaned_joined_data, "cleaned_matrisome_oviduct.csv", row.names = F
ALSE)


#######Ploting


#### Collagens
# Read your cleaned joined data file into a data frame
cleaned_joined_data <- read.csv("cleaned_matrisome_oviduct.csv")

# Filter rows with Matrisome_Category "Collagens"
collagen_data <- cleaned_joined_data %>%
  filter(Matrisome_Category == "Collagens")

# Reshape the data to have separate columns for each replicate and each tissu
e stage
organized_data <- collagen_data %>%
  pivot_longer(cols = starts_with("OF") | starts_with("OL"),
               names_to = c("Tissue", "Stage"),
               names_sep = "\\.",
               values_to = "Protein_Count")

## Warning: Expected 2 pieces. Missing pieces filled with `NA` in 2 rows [1,
5].

# Save the reshaped data to a new file

write.csv(organized_data, "Collagens_matrisome_oviduct_to_clean.csv", row.nam
es = FALSE)

data2 <- read.csv("Collagens_matrisome_oviduct_clean.csv")

png(filename="ovi_collagens.png",
    width = 12, height = 6, units = "in",
    res = 600)

boxplot(Protein_Count ~ Tissue*Genes,
        data = data2,
        cex.axis = 0.8,
        col = viridis(2, alpha = 0.7),
        lwd = 0.7,
        ylab = "",
        ylim = c(0, 2500000000),
        axes = FALSE,
        frame.plot = TRUE,
        xlab = "")
```

```
GROUP <- unique(data2$Genes)
groups <- 23
numbox <- 2
total <- groups * numbox
xpoints <- seq(median(1:numbox), total, numbox)

axis(2, seq(0, 2500000000, by = 200000000), las = 1, cex.axis = 0.7)
axis(1, labels = GROUP, at = xpoints, las = 2, cex.axis = 0.7)

stripchart(data2$Protein_Count ~ data2$Tissue*data2$Genes,
           method = "jitter",
           pch = 19,
           col = "black",
           vertical = TRUE,
           add = TRUE,
           cex = 0.3)

title(ylab = expression("Protein abundance (RU)"), line = 3.0)
legend("top", fill = viridis(2), legend = c("Oviduct Follicular", "Oviduct Lu
teal"), horiz = T, cex = 0.7)

dev.off()

## quartz_off_screen
##                  2

# Convert Tissue to factor
data2$Tissue <- as.factor(data2$Tissue)

# Create an empty list to store Tukey results
tukey_results_list <- list()

# Unique genes
unique_genes <- unique(data2$Genes)

# Loop over each gene
for (gene in unique_genes) {
  # Subset data for the current gene
  gene_data <- subset(data2, Genes == gene)

  # Fit the generalized linear model
  glm_model <- glm(Protein_Count ~ Tissue, data = gene_data, family = gaussia
n)

  # Perform Tukey test
  contrast_matrix <- glht(glm_model, linfct = mcp(Tissue = "Tukey"))
  tukey_results <- summary(contrast_matrix)

  # Store the results
```

```
  tukey_results_list[[gene]] <- tukey_results
}

# Print the results for each gene
for (gene in unique_genes) {
  cat("Tukey results for gene:", gene, "\n")
  print(tukey_results_list[[gene]])
  cat("\n")
}

## Tukey results for gene: COL11A1
##
##   Simultaneous Tests for General Linear Hypotheses
##
## Multiple Comparisons of Means: Tukey Contrasts
##
##
## Fit: glm(formula = Protein_Count ~ Tissue, family = gaussian, data = gene_
data)
##
## Linear Hypotheses:
##                Estimate Std. Error z value Pr(>|z|)
## OL - OF == 0  1003191    1639081   0.612    0.541
## (Adjusted p values reported -- single-step method)
##
##
## Tukey results for gene: COL12A1
##
##   Simultaneous Tests for General Linear Hypotheses
##
## Multiple Comparisons of Means: Tukey Contrasts
##
##
## Fit: glm(formula = Protein_Count ~ Tissue, family = gaussian, data = gene_
data)
##
## Linear Hypotheses:
##                Estimate Std. Error z value Pr(>|z|)
## OL - OF == 0 -4379758    4757084  -0.921    0.357
## (Adjusted p values reported -- single-step method)
##
##
## Tukey results for gene: COL14A1
##
##   Simultaneous Tests for General Linear Hypotheses
##
## Multiple Comparisons of Means: Tukey Contrasts
##
##
## Fit: glm(formula = Protein_Count ~ Tissue, family = gaussian, data = gene_
```

```
data)
##
## Linear Hypotheses:
##              Estimate Std. Error z value Pr(>|z|)
## OL - OF == 0 -1789033    5324197  -0.336    0.737
## (Adjusted p values reported -- single-step method)
##
##
## Tukey results for gene: COL15A1
##
##   Simultaneous Tests for General Linear Hypotheses
##
## Multiple Comparisons of Means: Tukey Contrasts
##
##
## Fit: glm(formula = Protein_Count ~ Tissue, family = gaussian, data = gene_
data)
##
## Linear Hypotheses:
##              Estimate Std. Error z value Pr(>|z|)
## OL - OF == 0  -413835    2806186  -0.147    0.883
## (Adjusted p values reported -- single-step method)
##
##
## Tukey results for gene: COL16A1
##
##   Simultaneous Tests for General Linear Hypotheses
##
## Multiple Comparisons of Means: Tukey Contrasts
##
##
## Fit: glm(formula = Protein_Count ~ Tissue, family = gaussian, data = gene_
data)
##
## Linear Hypotheses:
##              Estimate Std. Error z value Pr(>|z|)
## OL - OF == 0   269518     810343   0.333    0.739
## (Adjusted p values reported -- single-step method)
##
##
## Tukey results for gene: COL18A1
##
##   Simultaneous Tests for General Linear Hypotheses
##
## Multiple Comparisons of Means: Tukey Contrasts
##
##
## Fit: glm(formula = Protein_Count ~ Tissue, family = gaussian, data = gene_
data)
##
```

```
## Linear Hypotheses:
##              Estimate Std. Error z value Pr(>|z|)
## OL - OF == 0   480710    1348200   0.357    0.721
## (Adjusted p values reported -- single-step method)
##
##
## Tukey results for gene: COL1A1
##
##   Simultaneous Tests for General Linear Hypotheses
##
## Multiple Comparisons of Means: Tukey Contrasts
##
##
## Fit: glm(formula = Protein_Count ~ Tissue, family = gaussian, data = gene_
data)
##
## Linear Hypotheses:
##               Estimate Std. Error z value Pr(>|z|)
## OL - OF == 0 140663833  407617490   0.345     0.73
## (Adjusted p values reported -- single-step method)
##
##
## Tukey results for gene: COL1A2
##
##   Simultaneous Tests for General Linear Hypotheses
##
## Multiple Comparisons of Means: Tukey Contrasts
##
##
## Fit: glm(formula = Protein_Count ~ Tissue, family = gaussian, data = gene_
data)
##
## Linear Hypotheses:
##               Estimate Std. Error z value Pr(>|z|)
## OL - OF == 0  72662333  230925594   0.315    0.753
## (Adjusted p values reported -- single-step method)
##
##
## Tukey results for gene: COL2A1
##
##   Simultaneous Tests for General Linear Hypotheses
##
## Multiple Comparisons of Means: Tukey Contrasts
##
##
## Fit: glm(formula = Protein_Count ~ Tissue, family = gaussian, data = gene_
data)
##
## Linear Hypotheses:
##              Estimate Std. Error z value Pr(>|z|)
```

```
## OL - OF == 0   197017     217794   0.905    0.366
## (Adjusted p values reported -- single-step method)
##
##
## Tukey results for gene: COL3A1
##
##   Simultaneous Tests for General Linear Hypotheses
##
## Multiple Comparisons of Means: Tukey Contrasts
##
##
## Fit: glm(formula = Protein_Count ~ Tissue, family = gaussian, data = gene_
data)
##
## Linear Hypotheses:
##              Estimate Std. Error z value Pr(>|z|)
## OL - OF == 0 -5555448    2098417  -2.647  0.00811 **
## ---
## Signif. codes:  0 '***' 0.001 '**' 0.01 '*' 0.05 '.' 0.1 ' ' 1
## (Adjusted p values reported -- single-step method)
##
##
## Tukey results for gene: COL4A1
##
##   Simultaneous Tests for General Linear Hypotheses
##
## Multiple Comparisons of Means: Tukey Contrasts
##
##
## Fit: glm(formula = Protein_Count ~ Tissue, family = gaussian, data = gene_
data)
##
## Linear Hypotheses:
##              Estimate Std. Error z value Pr(>|z|)
## OL - OF == 0   943797    2425123   0.389    0.697
## (Adjusted p values reported -- single-step method)
##
##
## Tukey results for gene: COL4A2
##
##   Simultaneous Tests for General Linear Hypotheses
##
## Multiple Comparisons of Means: Tukey Contrasts
##
##
## Fit: glm(formula = Protein_Count ~ Tissue, family = gaussian, data = gene_
data)
##
## Linear Hypotheses:
##              Estimate Std. Error z value Pr(>|z|)
```

```
## OL - OF == 0     3812    1127816   0.003    0.997
## (Adjusted p values reported -- single-step method)
##
##
## Tukey results for gene: COL4A3
##
##   Simultaneous Tests for General Linear Hypotheses
##
## Multiple Comparisons of Means: Tukey Contrasts
##
##
## Fit: glm(formula = Protein_Count ~ Tissue, family = gaussian, data = gene_
data)
##
## Linear Hypotheses:
##              Estimate Std. Error z value Pr(>|z|)
## OL - OF == 0   691036     948199   0.729    0.466
## (Adjusted p values reported -- single-step method)
##
##
## Tukey results for gene: COL4A5
##
##   Simultaneous Tests for General Linear Hypotheses
##
## Multiple Comparisons of Means: Tukey Contrasts
##
##
## Fit: glm(formula = Protein_Count ~ Tissue, family = gaussian, data = gene_
data)
##
## Linear Hypotheses:
##              Estimate Std. Error z value Pr(>|z|)
## OL - OF == 0   239678    2089850   0.115    0.909
## (Adjusted p values reported -- single-step method)
##
##
## Tukey results for gene: COL4A6
##
##   Simultaneous Tests for General Linear Hypotheses
##
## Multiple Comparisons of Means: Tukey Contrasts
##
##
## Fit: glm(formula = Protein_Count ~ Tissue, family = gaussian, data = gene_
data)
##
## Linear Hypotheses:
##              Estimate Std. Error z value Pr(>|z|)
## OL - OF == 0   -93043     797034  -0.117    0.907
## (Adjusted p values reported -- single-step method)
```

```
##
##
## Tukey results for gene: COL5A1
##
##   Simultaneous Tests for General Linear Hypotheses
##
## Multiple Comparisons of Means: Tukey Contrasts
##
##
## Fit: glm(formula = Protein_Count ~ Tissue, family = gaussian, data = gene_
data)
##
## Linear Hypotheses:
##              Estimate Std. Error z value Pr(>|z|)
## OL - OF == 0    -9011    2160252  -0.004    0.997
## (Adjusted p values reported -- single-step method)
##
##
## Tukey results for gene: COL5A2
##
##   Simultaneous Tests for General Linear Hypotheses
##
## Multiple Comparisons of Means: Tukey Contrasts
##
##
## Fit: glm(formula = Protein_Count ~ Tissue, family = gaussian, data = gene_
data)
##
## Linear Hypotheses:
##              Estimate Std. Error z value Pr(>|z|)
## OL - OF == 0  1457530    5317120   0.274    0.784
## (Adjusted p values reported -- single-step method)
##
##
## Tukey results for gene: COL5A3
##
##   Simultaneous Tests for General Linear Hypotheses
##
## Multiple Comparisons of Means: Tukey Contrasts
##
##
## Fit: glm(formula = Protein_Count ~ Tissue, family = gaussian, data = gene_
data)
##
## Linear Hypotheses:
##              Estimate Std. Error z value Pr(>|z|)
## OL - OF == 0   202374    4508151   0.045    0.964
## (Adjusted p values reported -- single-step method)
##
##
```

```
## Tukey results for gene: COL6A1
##
##   Simultaneous Tests for General Linear Hypotheses
##
## Multiple Comparisons of Means: Tukey Contrasts
##
##
## Fit: glm(formula = Protein_Count ~ Tissue, family = gaussian, data = gene_
data)
##
## Linear Hypotheses:
##              Estimate Std. Error z value Pr(>|z|)
## OL - OF == 0 16726150   29555094   0.566    0.571
## (Adjusted p values reported -- single-step method)
##
##
## Tukey results for gene: COL6A2
##
##   Simultaneous Tests for General Linear Hypotheses
##
## Multiple Comparisons of Means: Tukey Contrasts
##
##
## Fit: glm(formula = Protein_Count ~ Tissue, family = gaussian, data = gene_
data)
##
## Linear Hypotheses:
##              Estimate Std. Error z value Pr(>|z|)
## OL - OF == 0 10504217   11852593   0.886    0.375
## (Adjusted p values reported -- single-step method)
##
##
## Tukey results for gene: COL6A3
##
##   Simultaneous Tests for General Linear Hypotheses
##
## Multiple Comparisons of Means: Tukey Contrasts
##
##
## Fit: glm(formula = Protein_Count ~ Tissue, family = gaussian, data = gene_
data)
##
## Linear Hypotheses:
##              Estimate Std. Error z value Pr(>|z|)
## OL - OF == 0 -6567883   47681976  -0.138     0.89
## (Adjusted p values reported -- single-step method)
##
##
## Tukey results for gene: COL6A5
##
```

```
##   Simultaneous Tests for General Linear Hypotheses
##
## Multiple Comparisons of Means: Tukey Contrasts
##
##
## Fit: glm(formula = Protein_Count ~ Tissue, family = gaussian, data = gene_
data)
##
## Linear Hypotheses:
##              Estimate Std. Error z value Pr(>|z|)
## OL - OF == 0  -350679     824435  -0.425    0.671
## (Adjusted p values reported -- single-step method)
##
##
## Tukey results for gene: COL6A6
##
##   Simultaneous Tests for General Linear Hypotheses
##
## Multiple Comparisons of Means: Tukey Contrasts
##
##
## Fit: glm(formula = Protein_Count ~ Tissue, family = gaussian, data = gene_
data)
##
## Linear Hypotheses:
##              Estimate Std. Error z value Pr(>|z|)
## OL - OF == 0   203654     735928   0.277    0.782
## (Adjusted p values reported -- single-step method)
```

```
#### ECM Glycoproteins
# Read your cleaned joined data file into a data frame
cleaned_joined_data <- read.csv("cleaned_matrisome_oviduct.csv")

# Filter rows with Matrisome_Category "ECM Glycoproteins"
Glycoproteins_data <- cleaned_joined_data %>%
  filter(Matrisome_Category == "ECM Glycoproteins")

# Reshape the data to have separate columns for each replicate and each tissu
e stage
organized_data <- Glycoproteins_data %>%
  pivot_longer(cols = starts_with("OF") | starts_with("OL"),
               names_to = c("Tissue", "Stage"),
               names_sep = "\\.",
               values_to = "Protein_Count")
```

```
## Warning: Expected 2 pieces. Missing pieces filled with `NA` in 2 rows [1,
5].
```

```
# Save the reshaped data to a new file
```

```
write.csv(organized_data, "Glycoproteins_data_matrisome_oviduct_to_clean.csv"
, row.names = FALSE)

data2 <- read.csv("Glycoproteins_data_matrisome_oviduct_clean.csv")

png(filename="ovi_ECM Glycoproteins.png",
    width = 12, height = 6, units = "in",
    res = 600)

boxplot(Protein_Count ~ Tissue*Genes,
        data = data2,
        cex.axis = 0.8,
        col = viridis(2, alpha = 0.7),
        lwd = 0.7,
        ylab = "",
        ylim = c(0, 100000000),
        axes = FALSE,
        frame.plot = TRUE,
        xlab = "")

GROUP <- unique(data2$Genes)
groups <- 84
numbox <- 2
total <- groups * numbox
xpoints <- seq(median(1:numbox), total, numbox)

axis(2, seq(0, 100000000, by = 10000000), las = 1, cex.axis = 0.7)
axis(1, labels = GROUP, at = xpoints, las = 2, cex.axis = 0.7)

stripchart(data2$Protein_Count ~ data2$Tissue*data2$Genes,
           method = "jitter",
           pch = 19,
           col = "black",
           vertical = TRUE,
           add = TRUE,
           cex = 0.3)

title(ylab = expression("Protein abundance (RU)"), line = 3.0)
legend("top", fill = viridis(2), legend = c("Oviduct Follicular", "Oviduct Lu
teal"), horiz = T, cex = 0.7)

dev.off()

## quartz_off_screen
##                 2

# Convert Tissue to factor
data2$Tissue <- as.factor(data2$Tissue)
```

```r
# Create an empty list to store Tukey results
tukey_results_list <- list()

# Unique genes
unique_genes <- unique(data2$Genes)

# Loop over each gene
for (gene in unique_genes) {
  # Subset data for the current gene
  gene_data <- subset(data2, Genes == gene)

  # Fit the generalized linear model
  glm_model <- glm(Protein_Count ~ Tissue, data = gene_data, family = gaussia
n)

  # Perform Tukey test
  contrast_matrix <- glht(glm_model, linfct = mcp(Tissue = "Tukey"))
  tukey_results <- summary(contrast_matrix)

  # Store the results
  tukey_results_list[[gene]] <- tukey_results
}

# Print the results for each gene
for (gene in unique_genes) {
  cat("Tukey results for gene:", gene, "\n")
  print(tukey_results_list[[gene]])
  cat("\n")
}
```

```
## Tukey results for gene: ABI3BP
##
##   Simultaneous Tests for General Linear Hypotheses
##
## Multiple Comparisons of Means: Tukey Contrasts
##
##
## Fit: glm(formula = Protein_Count ~ Tissue, family = gaussian, data = gene_
data)
##
## Linear Hypotheses:
##                Estimate Std. Error z value Pr(>|z|)
## OL - OF == 0  -267526     282807   -0.946    0.344
## (Adjusted p values reported -- single-step method)
##
##
## Tukey results for gene: AEBP1
##
##   Simultaneous Tests for General Linear Hypotheses
```

```
##
## Multiple Comparisons of Means: Tukey Contrasts
##
##
## Fit: glm(formula = Protein_Count ~ Tissue, family = gaussian, data = gene_
data)
##
## Linear Hypotheses:
##              Estimate Std. Error z value Pr(>|z|)
## OL - OF == 0   -15592     403458  -0.039    0.969
## (Adjusted p values reported -- single-step method)
##
##
## Tukey results for gene: AGRN
##
##   Simultaneous Tests for General Linear Hypotheses
##
## Multiple Comparisons of Means: Tukey Contrasts
##
##
## Fit: glm(formula = Protein_Count ~ Tissue, family = gaussian, data = gene_
data)
##
## Linear Hypotheses:
##              Estimate Std. Error z value Pr(>|z|)
## OL - OF == 0   164236     278740   0.589    0.556
## (Adjusted p values reported -- single-step method)
##
##
## Tukey results for gene: AHSG
##
##   Simultaneous Tests for General Linear Hypotheses
##
## Multiple Comparisons of Means: Tukey Contrasts
##
##
## Fit: glm(formula = Protein_Count ~ Tissue, family = gaussian, data = gene_
data)
##
## Linear Hypotheses:
##              Estimate Std. Error z value Pr(>|z|)
## OL - OF == 0   -29713     299478  -0.099    0.921
## (Adjusted p values reported -- single-step method)
##
##
## Tukey results for gene: CILP2
##
##   Simultaneous Tests for General Linear Hypotheses
##
## Multiple Comparisons of Means: Tukey Contrasts
```

```
##
##
## Fit: glm(formula = Protein_Count ~ Tissue, family = gaussian, data = gene_
data)
##
## Linear Hypotheses:
##                Estimate Std. Error z value Pr(>|z|)
## OL - OF == 0 -3954781    5968816  -0.663    0.508
## (Adjusted p values reported -- single-step method)
##
##
## Tukey results for gene: CRELD2
##
##   Simultaneous Tests for General Linear Hypotheses
##
## Multiple Comparisons of Means: Tukey Contrasts
##
##
## Fit: glm(formula = Protein_Count ~ Tissue, family = gaussian, data = gene_
data)
##
## Linear Hypotheses:
##                Estimate Std. Error z value Pr(>|z|)
## OL - OF == 0   -21900      62576   -0.35    0.726
## (Adjusted p values reported -- single-step method)
##
##
## Tukey results for gene: DPT
##
##   Simultaneous Tests for General Linear Hypotheses
##
## Multiple Comparisons of Means: Tukey Contrasts
##
##
## Fit: glm(formula = Protein_Count ~ Tissue, family = gaussian, data = gene_
data)
##
## Linear Hypotheses:
##                 Estimate Std. Error z value Pr(>|z|)
## OL - OF == 0    648417  119494494   0.005    0.996
## (Adjusted p values reported -- single-step method)
##
##
## Tukey results for gene: ECM1
##
##   Simultaneous Tests for General Linear Hypotheses
##
## Multiple Comparisons of Means: Tukey Contrasts
##
##
```

```
## Fit: glm(formula = Protein_Count ~ Tissue, family = gaussian, data = gene_
data)
##
## Linear Hypotheses:
##              Estimate Std. Error z value Pr(>|z|)
## OL - OF == 0   -88292    1118613  -0.079    0.937
## (Adjusted p values reported -- single-step method)
##
##
## Tukey results for gene: ECM2
##
##   Simultaneous Tests for General Linear Hypotheses
##
## Multiple Comparisons of Means: Tukey Contrasts
##
##
## Fit: glm(formula = Protein_Count ~ Tissue, family = gaussian, data = gene_
data)
##
## Linear Hypotheses:
##              Estimate Std. Error z value Pr(>|z|)
## OL - OF == 0   -14773     157010  -0.094    0.925
## (Adjusted p values reported -- single-step method)
##
##
## Tukey results for gene: EFEMP1
##
##   Simultaneous Tests for General Linear Hypotheses
##
## Multiple Comparisons of Means: Tukey Contrasts
##
##
## Fit: glm(formula = Protein_Count ~ Tissue, family = gaussian, data = gene_
data)
##
## Linear Hypotheses:
##              Estimate Std. Error z value Pr(>|z|)
## OL - OF == 0   197463     342962   0.576    0.565
## (Adjusted p values reported -- single-step method)
##
##
## Tukey results for gene: EFEMP2
##
##   Simultaneous Tests for General Linear Hypotheses
##
## Multiple Comparisons of Means: Tukey Contrasts
##
##
## Fit: glm(formula = Protein_Count ~ Tissue, family = gaussian, data = gene_
data)
```

```
##
## Linear Hypotheses:
##               Estimate Std. Error z value Pr(>|z|)
## OL - OF == 0   386695     416812   0.928    0.354
## (Adjusted p values reported -- single-step method)
##
##
## Tukey results for gene: EMID1
##
##   Simultaneous Tests for General Linear Hypotheses
##
## Multiple Comparisons of Means: Tukey Contrasts
##
##
## Fit: glm(formula = Protein_Count ~ Tissue, family = gaussian, data = gene_
data)
##
## Linear Hypotheses:
##               Estimate Std. Error z value Pr(>|z|)
## OL - OF == 0   137939     124787   1.105    0.269
## (Adjusted p values reported -- single-step method)
##
##
## Tukey results for gene: EMILIN1
##
##   Simultaneous Tests for General Linear Hypotheses
##
## Multiple Comparisons of Means: Tukey Contrasts
##
##
## Fit: glm(formula = Protein_Count ~ Tissue, family = gaussian, data = gene_
data)
##
## Linear Hypotheses:
##               Estimate Std. Error z value Pr(>|z|)
## OL - OF == 0  3699799    8131341   0.455    0.649
## (Adjusted p values reported -- single-step method)
##
##
## Tukey results for gene: EMILIN2
##
##   Simultaneous Tests for General Linear Hypotheses
##
## Multiple Comparisons of Means: Tukey Contrasts
##
##
## Fit: glm(formula = Protein_Count ~ Tissue, family = gaussian, data = gene_
data)
##
## Linear Hypotheses:
```

```
##               Estimate Std. Error z value Pr(>|z|)
## OL - OF == 0    97512     162874   0.599    0.549
## (Adjusted p values reported -- single-step method)
##
##
## Tukey results for gene: EMILIN3
##
##   Simultaneous Tests for General Linear Hypotheses
##
## Multiple Comparisons of Means: Tukey Contrasts
##
##
## Fit: glm(formula = Protein_Count ~ Tissue, family = gaussian, data = gene_
data)
##
## Linear Hypotheses:
##               Estimate Std. Error z value Pr(>|z|)
## OL - OF == 0   407727     577578   0.706     0.48
## (Adjusted p values reported -- single-step method)
##
##
## Tukey results for gene: FBLN1
##
##   Simultaneous Tests for General Linear Hypotheses
##
## Multiple Comparisons of Means: Tukey Contrasts
##
##
## Fit: glm(formula = Protein_Count ~ Tissue, family = gaussian, data = gene_
data)
##
## Linear Hypotheses:
##               Estimate Std. Error z value Pr(>|z|)
## OL - OF == 0   460509    1645021    0.28     0.78
## (Adjusted p values reported -- single-step method)
##
##
## Tukey results for gene: FBLN2
##
##   Simultaneous Tests for General Linear Hypotheses
##
## Multiple Comparisons of Means: Tukey Contrasts
##
##
## Fit: glm(formula = Protein_Count ~ Tissue, family = gaussian, data = gene_
data)
##
## Linear Hypotheses:
##               Estimate Std. Error z value Pr(>|z|)
## OL - OF == 0    67107      72033   0.932    0.352
```

```
## (Adjusted p values reported -- single-step method)
##
##
## Tukey results for gene: FBLN5
##
##   Simultaneous Tests for General Linear Hypotheses
##
## Multiple Comparisons of Means: Tukey Contrasts
##
##
## Fit: glm(formula = Protein_Count ~ Tissue, family = gaussian, data = gene_
data)
##
## Linear Hypotheses:
##              Estimate Std. Error z value Pr(>|z|)
## OL - OF == 0   745574     787954   0.946    0.344
## (Adjusted p values reported -- single-step method)
##
##
## Tukey results for gene: FBN1
##
##   Simultaneous Tests for General Linear Hypotheses
##
## Multiple Comparisons of Means: Tukey Contrasts
##
##
## Fit: glm(formula = Protein_Count ~ Tissue, family = gaussian, data = gene_
data)
##
## Linear Hypotheses:
##              Estimate Std. Error z value Pr(>|z|)
## OL - OF == 0  1556192    1601720   0.972    0.331
## (Adjusted p values reported -- single-step method)
##
##
## Tukey results for gene: FBN2
##
##   Simultaneous Tests for General Linear Hypotheses
##
## Multiple Comparisons of Means: Tukey Contrasts
##
##
## Fit: glm(formula = Protein_Count ~ Tissue, family = gaussian, data = gene_
data)
##
## Linear Hypotheses:
##              Estimate Std. Error z value Pr(>|z|)
## OL - OF == 0   194169     138002   1.407    0.159
## (Adjusted p values reported -- single-step method)
##
```

```
##
## Tukey results for gene: FBN3
##
##   Simultaneous Tests for General Linear Hypotheses
##
## Multiple Comparisons of Means: Tukey Contrasts
##
##
## Fit: glm(formula = Protein_Count ~ Tissue, family = gaussian, data = gene_
data)
##
## Linear Hypotheses:
##              Estimate Std. Error z value Pr(>|z|)
## OL - OF == 0   346156     457681   0.756    0.449
## (Adjusted p values reported -- single-step method)
##
##
## Tukey results for gene: FGA
##
##   Simultaneous Tests for General Linear Hypotheses
##
## Multiple Comparisons of Means: Tukey Contrasts
##
##
## Fit: glm(formula = Protein_Count ~ Tissue, family = gaussian, data = gene_
data)
##
## Linear Hypotheses:
##              Estimate Std. Error z value Pr(>|z|)
## OL - OF == 0 -5352865    5264522  -1.017    0.309
## (Adjusted p values reported -- single-step method)
##
##
## Tukey results for gene: FGB
##
##   Simultaneous Tests for General Linear Hypotheses
##
## Multiple Comparisons of Means: Tukey Contrasts
##
##
## Fit: glm(formula = Protein_Count ~ Tissue, family = gaussian, data = gene_
data)
##
## Linear Hypotheses:
##              Estimate Std. Error z value Pr(>|z|)
## OL - OF == 0 -8921127    9229021  -0.967    0.334
## (Adjusted p values reported -- single-step method)
##
##
## Tukey results for gene: FGG
```

```
##
##   Simultaneous Tests for General Linear Hypotheses
##
## Multiple Comparisons of Means: Tukey Contrasts
##
##
## Fit: glm(formula = Protein_Count ~ Tissue, family = gaussian, data = gene_
data)
##
## Linear Hypotheses:
##               Estimate Std. Error z value Pr(>|z|)
## OL - OF == 0 -3741212    3414184  -1.096    0.273
## (Adjusted p values reported -- single-step method)
##
##
## Tukey results for gene: FGL2
##
##   Simultaneous Tests for General Linear Hypotheses
##
## Multiple Comparisons of Means: Tukey Contrasts
##
##
## Fit: glm(formula = Protein_Count ~ Tissue, family = gaussian, data = gene_
data)
##
## Linear Hypotheses:
##               Estimate Std. Error z value Pr(>|z|)
## OL - OF == 0   -420.3   143473.9  -0.003    0.998
## (Adjusted p values reported -- single-step method)
##
##
## Tukey results for gene: FN1
##
##   Simultaneous Tests for General Linear Hypotheses
##
## Multiple Comparisons of Means: Tukey Contrasts
##
##
## Fit: glm(formula = Protein_Count ~ Tissue, family = gaussian, data = gene_
data)
##
## Linear Hypotheses:
##               Estimate Std. Error z value Pr(>|z|)
## OL - OF == 0 -1443922    3186191  -0.453     0.65
## (Adjusted p values reported -- single-step method)
##
##
## Tukey results for gene: IGFALS
##
##   Simultaneous Tests for General Linear Hypotheses
```

```
##
## Multiple Comparisons of Means: Tukey Contrasts
##
##
## Fit: glm(formula = Protein_Count ~ Tissue, family = gaussian, data = gene_
data)
##
## Linear Hypotheses:
##              Estimate Std. Error z value Pr(>|z|)
## OL - OF == 0    36820      52620     0.7    0.484
## (Adjusted p values reported -- single-step method)
##
##
## Tukey results for gene: IGFBP5
##
##   Simultaneous Tests for General Linear Hypotheses
##
## Multiple Comparisons of Means: Tukey Contrasts
##
##
## Fit: glm(formula = Protein_Count ~ Tissue, family = gaussian, data = gene_
data)
##
## Linear Hypotheses:
##              Estimate Std. Error z value Pr(>|z|)
## OL - OF == 0    -8900      58518  -0.152    0.879
## (Adjusted p values reported -- single-step method)
##
##
## Tukey results for gene: IGFBP6
##
##   Simultaneous Tests for General Linear Hypotheses
##
## Multiple Comparisons of Means: Tukey Contrasts
##
##
## Fit: glm(formula = Protein_Count ~ Tissue, family = gaussian, data = gene_
data)
##
## Linear Hypotheses:
##              Estimate Std. Error z value Pr(>|z|)
## OL - OF == 0    46178      80538   0.573    0.566
## (Adjusted p values reported -- single-step method)
##
##
## Tukey results for gene: IGFBP7
##
##   Simultaneous Tests for General Linear Hypotheses
##
## Multiple Comparisons of Means: Tukey Contrasts
```

```
##
##
## Fit: glm(formula = Protein_Count ~ Tissue, family = gaussian, data = gene_
data)
##
## Linear Hypotheses:
##              Estimate Std. Error z value Pr(>|z|)
## OL - OF == 0    12411     240881   0.052    0.959
## (Adjusted p values reported -- single-step method)
##
##
## Tukey results for gene: IGSF10
##
##   Simultaneous Tests for General Linear Hypotheses
##
## Multiple Comparisons of Means: Tukey Contrasts
##
##
## Fit: glm(formula = Protein_Count ~ Tissue, family = gaussian, data = gene_
data)
##
## Linear Hypotheses:
##              Estimate Std. Error z value Pr(>|z|)
## OL - OF == 0    10992      26924   0.408    0.683
## (Adjusted p values reported -- single-step method)
##
##
## Tukey results for gene: LAMA1
##
##   Simultaneous Tests for General Linear Hypotheses
##
## Multiple Comparisons of Means: Tukey Contrasts
##
##
## Fit: glm(formula = Protein_Count ~ Tissue, family = gaussian, data = gene_
data)
##
## Linear Hypotheses:
##              Estimate Std. Error z value Pr(>|z|)
## OL - OF == 0   172010     409823    0.42    0.675
## (Adjusted p values reported -- single-step method)
##
##
## Tukey results for gene: LAMA2
##
##   Simultaneous Tests for General Linear Hypotheses
##
## Multiple Comparisons of Means: Tukey Contrasts
##
##
```

```
## Fit: glm(formula = Protein_Count ~ Tissue, family = gaussian, data = gene_
data)
##
## Linear Hypotheses:
##              Estimate Std. Error z value Pr(>|z|)
## OL - OF == 0    25522     125547   0.203    0.839
## (Adjusted p values reported -- single-step method)
##
##
## Tukey results for gene: LAMA3
##
##   Simultaneous Tests for General Linear Hypotheses
##
## Multiple Comparisons of Means: Tukey Contrasts
##
##
## Fit: glm(formula = Protein_Count ~ Tissue, family = gaussian, data = gene_
data)
##
## Linear Hypotheses:
##              Estimate Std. Error z value Pr(>|z|)
## OL - OF == 0   112511     343326   0.328    0.743
## (Adjusted p values reported -- single-step method)
##
##
## Tukey results for gene: LAMA4
##
##   Simultaneous Tests for General Linear Hypotheses
##
## Multiple Comparisons of Means: Tukey Contrasts
##
##
## Fit: glm(formula = Protein_Count ~ Tissue, family = gaussian, data = gene_
data)
##
## Linear Hypotheses:
##              Estimate Std. Error z value Pr(>|z|)
## OL - OF == 0  -287142    1053199  -0.273    0.785
## (Adjusted p values reported -- single-step method)
##
##
## Tukey results for gene: LAMA5
##
##   Simultaneous Tests for General Linear Hypotheses
##
## Multiple Comparisons of Means: Tukey Contrasts
##
##
## Fit: glm(formula = Protein_Count ~ Tissue, family = gaussian, data = gene_
data)
```

```
##
## Linear Hypotheses:
##              Estimate Std. Error z value Pr(>|z|)
## OL - OF == 0   252257    3056912   0.083    0.934
## (Adjusted p values reported -- single-step method)
##
##
## Tukey results for gene: LAMB1
##
##   Simultaneous Tests for General Linear Hypotheses
##
## Multiple Comparisons of Means: Tukey Contrasts
##
##
## Fit: glm(formula = Protein_Count ~ Tissue, family = gaussian, data = gene_
data)
##
## Linear Hypotheses:
##              Estimate Std. Error z value Pr(>|z|)
## OL - OF == 0   -73428    1839503   -0.04    0.968
## (Adjusted p values reported -- single-step method)
##
##
## Tukey results for gene: LAMB2
##
##   Simultaneous Tests for General Linear Hypotheses
##
## Multiple Comparisons of Means: Tukey Contrasts
##
##
## Fit: glm(formula = Protein_Count ~ Tissue, family = gaussian, data = gene_
data)
##
## Linear Hypotheses:
##              Estimate Std. Error z value Pr(>|z|)
## OL - OF == 0   -54625    3300360  -0.017    0.987
## (Adjusted p values reported -- single-step method)
##
##
## Tukey results for gene: LAMB3
##
##   Simultaneous Tests for General Linear Hypotheses
##
## Multiple Comparisons of Means: Tukey Contrasts
##
##
## Fit: glm(formula = Protein_Count ~ Tissue, family = gaussian, data = gene_
data)
##
## Linear Hypotheses:
```

```
##               Estimate Std. Error z value Pr(>|z|)
## OL - OF == 0   -78275     152012  -0.515    0.607
## (Adjusted p values reported -- single-step method)
##
##
## Tukey results for gene: LAMC1
##
##   Simultaneous Tests for General Linear Hypotheses
##
## Multiple Comparisons of Means: Tukey Contrasts
##
##
## Fit: glm(formula = Protein_Count ~ Tissue, family = gaussian, data = gene_
data)
##
## Linear Hypotheses:
##               Estimate Std. Error z value Pr(>|z|)
## OL - OF == 0  -535718    4424732  -0.121    0.904
## (Adjusted p values reported -- single-step method)
##
##
## Tukey results for gene: LAMC2
##
##   Simultaneous Tests for General Linear Hypotheses
##
## Multiple Comparisons of Means: Tukey Contrasts
##
##
## Fit: glm(formula = Protein_Count ~ Tissue, family = gaussian, data = gene_
data)
##
## Linear Hypotheses:
##               Estimate Std. Error z value Pr(>|z|)
## OL - OF == 0   -17700     193784  -0.091    0.927
## (Adjusted p values reported -- single-step method)
##
##
## Tukey results for gene: LAMC3
##
##   Simultaneous Tests for General Linear Hypotheses
##
## Multiple Comparisons of Means: Tukey Contrasts
##
##
## Fit: glm(formula = Protein_Count ~ Tissue, family = gaussian, data = gene_
data)
##
## Linear Hypotheses:
##               Estimate Std. Error z value Pr(>|z|)
## OL - OF == 0    74533      83744    0.89    0.373
```

```
## (Adjusted p values reported -- single-step method)
##
##
## Tukey results for gene: LRG1
##
##   Simultaneous Tests for General Linear Hypotheses
##
## Multiple Comparisons of Means: Tukey Contrasts
##
##
## Fit: glm(formula = Protein_Count ~ Tissue, family = gaussian, data = gene_
data)
##
## Linear Hypotheses:
##              Estimate Std. Error z value Pr(>|z|)
## OL - OF == 0     6699      46552   0.144    0.886
## (Adjusted p values reported -- single-step method)
##
##
## Tukey results for gene: LTBP1
##
##   Simultaneous Tests for General Linear Hypotheses
##
## Multiple Comparisons of Means: Tukey Contrasts
##
##
## Fit: glm(formula = Protein_Count ~ Tissue, family = gaussian, data = gene_
data)
##
## Linear Hypotheses:
##              Estimate Std. Error z value Pr(>|z|)
## OL - OF == 0    38597     212448   0.182    0.856
## (Adjusted p values reported -- single-step method)
##
##
## Tukey results for gene: LTBP2
##
##   Simultaneous Tests for General Linear Hypotheses
##
## Multiple Comparisons of Means: Tukey Contrasts
##
##
## Fit: glm(formula = Protein_Count ~ Tissue, family = gaussian, data = gene_
data)
##
## Linear Hypotheses:
##              Estimate Std. Error z value Pr(>|z|)
## OL - OF == 0   -27564      82292  -0.335    0.738
## (Adjusted p values reported -- single-step method)
##
```

```
##
## Tukey results for gene: LTBP3
##
##   Simultaneous Tests for General Linear Hypotheses
##
## Multiple Comparisons of Means: Tukey Contrasts
##
##
## Fit: glm(formula = Protein_Count ~ Tissue, family = gaussian, data = gene_
data)
##
## Linear Hypotheses:
##              Estimate Std. Error z value Pr(>|z|)
## OL - OF == 0   -25711      28663  -0.897     0.37
## (Adjusted p values reported -- single-step method)
##
##
## Tukey results for gene: LTBP4
##
##   Simultaneous Tests for General Linear Hypotheses
##
## Multiple Comparisons of Means: Tukey Contrasts
##
##
## Fit: glm(formula = Protein_Count ~ Tissue, family = gaussian, data = gene_
data)
##
## Linear Hypotheses:
##              Estimate Std. Error z value Pr(>|z|)
## OL - OF == 0    25293    1324539   0.019    0.985
## (Adjusted p values reported -- single-step method)
##
##
## Tukey results for gene: MATN2
##
##   Simultaneous Tests for General Linear Hypotheses
##
## Multiple Comparisons of Means: Tukey Contrasts
##
##
## Fit: glm(formula = Protein_Count ~ Tissue, family = gaussian, data = gene_
data)
##
## Linear Hypotheses:
##              Estimate Std. Error z value Pr(>|z|)
## OL - OF == 0   487314     296807   1.642    0.101
## (Adjusted p values reported -- single-step method)
##
##
## Tukey results for gene: MATN4
```

```
##
##   Simultaneous Tests for General Linear Hypotheses
##
## Multiple Comparisons of Means: Tukey Contrasts
##
##
## Fit: glm(formula = Protein_Count ~ Tissue, family = gaussian, data = gene_
data)
##
## Linear Hypotheses:
##              Estimate Std. Error z value Pr(>|z|)
## OL - OF == 0    14189      59412   0.239    0.811
## (Adjusted p values reported -- single-step method)
##
##
## Tukey results for gene: MFAP2
##
##   Simultaneous Tests for General Linear Hypotheses
##
## Multiple Comparisons of Means: Tukey Contrasts
##
##
## Fit: glm(formula = Protein_Count ~ Tissue, family = gaussian, data = gene_
data)
##
## Linear Hypotheses:
##              Estimate Std. Error z value Pr(>|z|)
## OL - OF == 0  6163011    5598153   1.101    0.271
## (Adjusted p values reported -- single-step method)
##
##
## Tukey results for gene: MFAP4
##
##   Simultaneous Tests for General Linear Hypotheses
##
## Multiple Comparisons of Means: Tukey Contrasts
##
##
## Fit: glm(formula = Protein_Count ~ Tissue, family = gaussian, data = gene_
data)
##
## Linear Hypotheses:
##              Estimate Std. Error z value Pr(>|z|)
## OL - OF == 0  3386800    6327882   0.535    0.592
## (Adjusted p values reported -- single-step method)
##
##
## Tukey results for gene: MFAP5
##
##   Simultaneous Tests for General Linear Hypotheses
```

```
##
## Multiple Comparisons of Means: Tukey Contrasts
##
##
## Fit: glm(formula = Protein_Count ~ Tissue, family = gaussian, data = gene_
data)
##
## Linear Hypotheses:
##               Estimate Std. Error z value Pr(>|z|)
## OL - OF == 0  3322647    3357433    0.99    0.322
## (Adjusted p values reported -- single-step method)
##
##
## Tukey results for gene: MFGE8
##
##   Simultaneous Tests for General Linear Hypotheses
##
## Multiple Comparisons of Means: Tukey Contrasts
##
##
## Fit: glm(formula = Protein_Count ~ Tissue, family = gaussian, data = gene_
data)
##
## Linear Hypotheses:
##               Estimate Std. Error z value Pr(>|z|)
## OL - OF == 0   -30867     196006  -0.157    0.875
## (Adjusted p values reported -- single-step method)
##
##
## Tukey results for gene: MGP
##
##   Simultaneous Tests for General Linear Hypotheses
##
## Multiple Comparisons of Means: Tukey Contrasts
##
##
## Fit: glm(formula = Protein_Count ~ Tissue, family = gaussian, data = gene_
data)
##
## Linear Hypotheses:
##               Estimate Std. Error z value Pr(>|z|)
## OL - OF == 0  1699730    1448911   1.173    0.241
## (Adjusted p values reported -- single-step method)
##
##
## Tukey results for gene: MMRN2
##
##   Simultaneous Tests for General Linear Hypotheses
##
## Multiple Comparisons of Means: Tukey Contrasts
```

```
##
##
## Fit: glm(formula = Protein_Count ~ Tissue, family = gaussian, data = gene_
data)
##
## Linear Hypotheses:
##              Estimate Std. Error z value Pr(>|z|)
## OL - OF == 0   -33779      92633  -0.365    0.715
## (Adjusted p values reported -- single-step method)
##
##
## Tukey results for gene: MXRA5
##
##   Simultaneous Tests for General Linear Hypotheses
##
## Multiple Comparisons of Means: Tukey Contrasts
##
##
## Fit: glm(formula = Protein_Count ~ Tissue, family = gaussian, data = gene_
data)
##
## Linear Hypotheses:
##              Estimate Std. Error z value Pr(>|z|)
## OL - OF == 0    28321      50770   0.558    0.577
## (Adjusted p values reported -- single-step method)
##
##
## Tukey results for gene: NID1
##
##   Simultaneous Tests for General Linear Hypotheses
##
## Multiple Comparisons of Means: Tukey Contrasts
##
##
## Fit: glm(formula = Protein_Count ~ Tissue, family = gaussian, data = gene_
data)
##
## Linear Hypotheses:
##              Estimate Std. Error z value Pr(>|z|)
## OL - OF == 0 15394208   13227788   1.164    0.245
## (Adjusted p values reported -- single-step method)
##
##
## Tukey results for gene: NID2
##
##   Simultaneous Tests for General Linear Hypotheses
##
## Multiple Comparisons of Means: Tukey Contrasts
##
##
```

```
## Fit: glm(formula = Protein_Count ~ Tissue, family = gaussian, data = gene_
data)
##
## Linear Hypotheses:
##              Estimate Std. Error z value Pr(>|z|)
## OL - OF == 0  4454333    8189727   0.544    0.587
## (Adjusted p values reported -- single-step method)
##
##
## Tukey results for gene: NPNT
##
##   Simultaneous Tests for General Linear Hypotheses
##
## Multiple Comparisons of Means: Tukey Contrasts
##
##
## Fit: glm(formula = Protein_Count ~ Tissue, family = gaussian, data = gene_
data)
##
## Linear Hypotheses:
##              Estimate Std. Error z value Pr(>|z|)
## OL - OF == 0    -7895      49294   -0.16    0.873
## (Adjusted p values reported -- single-step method)
##
##
## Tukey results for gene: NTN4
##
##   Simultaneous Tests for General Linear Hypotheses
##
## Multiple Comparisons of Means: Tukey Contrasts
##
##
## Fit: glm(formula = Protein_Count ~ Tissue, family = gaussian, data = gene_
data)
##
## Linear Hypotheses:
##              Estimate Std. Error z value Pr(>|z|)
## OL - OF == 0     6853      26169   0.262    0.793
## (Adjusted p values reported -- single-step method)
##
##
## Tukey results for gene: PAPLN
##
##   Simultaneous Tests for General Linear Hypotheses
##
## Multiple Comparisons of Means: Tukey Contrasts
##
##
## Fit: glm(formula = Protein_Count ~ Tissue, family = gaussian, data = gene_
data)
```

```
##
## Linear Hypotheses:
##               Estimate Std. Error z value Pr(>|z|)
## OL - OF == 0    13394      59844   0.224    0.823
## (Adjusted p values reported -- single-step method)
##
##
## Tukey results for gene: PCOLCE
##
##   Simultaneous Tests for General Linear Hypotheses
##
## Multiple Comparisons of Means: Tukey Contrasts
##
##
## Fit: glm(formula = Protein_Count ~ Tissue, family = gaussian, data = gene_
data)
##
## Linear Hypotheses:
##               Estimate Std. Error z value Pr(>|z|)
## OL - OF == 0    13579      73918   0.184    0.854
## (Adjusted p values reported -- single-step method)
##
##
## Tukey results for gene: POSTN
##
##   Simultaneous Tests for General Linear Hypotheses
##
## Multiple Comparisons of Means: Tukey Contrasts
##
##
## Fit: glm(formula = Protein_Count ~ Tissue, family = gaussian, data = gene_
data)
##
## Linear Hypotheses:
##               Estimate Std. Error z value Pr(>|z|)
## OL - OF == 0  5910392    9567220   0.618    0.537
## (Adjusted p values reported -- single-step method)
##
##
## Tukey results for gene: PXDN
##
##   Simultaneous Tests for General Linear Hypotheses
##
## Multiple Comparisons of Means: Tukey Contrasts
##
##
## Fit: glm(formula = Protein_Count ~ Tissue, family = gaussian, data = gene_
data)
##
## Linear Hypotheses:
```

```
##               Estimate Std. Error z value Pr(>|z|)
## OL - OF == 0   -97132      58260  -1.667   0.0955 .
## ---
## Signif. codes:  0 '***' 0.001 '**' 0.01 '*' 0.05 '.' 0.1 ' ' 1
## (Adjusted p values reported -- single-step method)
##
##
## Tukey results for gene: SBSPON
##
##   Simultaneous Tests for General Linear Hypotheses
##
## Multiple Comparisons of Means: Tukey Contrasts
##
##
## Fit: glm(formula = Protein_Count ~ Tissue, family = gaussian, data = gene_
data)
##
## Linear Hypotheses:
##               Estimate Std. Error z value Pr(>|z|)
## OL - OF == 0  3594265    6831926   0.526    0.599
## (Adjusted p values reported -- single-step method)
##
##
## Tukey results for gene: SPARC
##
##   Simultaneous Tests for General Linear Hypotheses
##
## Multiple Comparisons of Means: Tukey Contrasts
##
##
## Fit: glm(formula = Protein_Count ~ Tissue, family = gaussian, data = gene_
data)
##
## Linear Hypotheses:
##               Estimate Std. Error z value Pr(>|z|)
## OL - OF == 0 -1516815    1301615  -1.165    0.244
## (Adjusted p values reported -- single-step method)
##
##
## Tukey results for gene: SPARCL1
##
##   Simultaneous Tests for General Linear Hypotheses
##
## Multiple Comparisons of Means: Tukey Contrasts
##
##
## Fit: glm(formula = Protein_Count ~ Tissue, family = gaussian, data = gene_
data)
##
## Linear Hypotheses:
```

```
##               Estimate Std. Error z value Pr(>|z|)
## OL - OF == 0  -104410     106933  -0.976    0.329
## (Adjusted p values reported -- single-step method)
##
##
## Tukey results for gene: SPON1
##
##   Simultaneous Tests for General Linear Hypotheses
##
## Multiple Comparisons of Means: Tukey Contrasts
##
##
## Fit: glm(formula = Protein_Count ~ Tissue, family = gaussian, data = gene_
data)
##
## Linear Hypotheses:
##               Estimate Std. Error z value Pr(>|z|)
## OL - OF == 0    59354     214456   0.277    0.782
## (Adjusted p values reported -- single-step method)
##
##
## Tukey results for gene: SPON2
##
##   Simultaneous Tests for General Linear Hypotheses
##
## Multiple Comparisons of Means: Tukey Contrasts
##
##
## Fit: glm(formula = Protein_Count ~ Tissue, family = gaussian, data = gene_
data)
##
## Linear Hypotheses:
##               Estimate Std. Error z value Pr(>|z|)
## OL - OF == 0   -49489      61013  -0.811    0.417
## (Adjusted p values reported -- single-step method)
##
##
## Tukey results for gene: SRPX2
##
##   Simultaneous Tests for General Linear Hypotheses
##
## Multiple Comparisons of Means: Tukey Contrasts
##
##
## Fit: glm(formula = Protein_Count ~ Tissue, family = gaussian, data = gene_
data)
##
## Linear Hypotheses:
##               Estimate Std. Error z value Pr(>|z|)
## OL - OF == 0  -162565     109749  -1.481    0.139
```

```
## (Adjusted p values reported -- single-step method)
##
##
## Tukey results for gene: SVEP1
##
##   Simultaneous Tests for General Linear Hypotheses
##
## Multiple Comparisons of Means: Tukey Contrasts
##
##
## Fit: glm(formula = Protein_Count ~ Tissue, family = gaussian, data = gene_
data)
##
## Linear Hypotheses:
##                Estimate Std. Error z value Pr(>|z|)
## OL - OF == 0     4766      39295   0.121    0.903
## (Adjusted p values reported -- single-step method)
##
##
## Tukey results for gene: TGFBI
##
##   Simultaneous Tests for General Linear Hypotheses
##
## Multiple Comparisons of Means: Tukey Contrasts
##
##
## Fit: glm(formula = Protein_Count ~ Tissue, family = gaussian, data = gene_
data)
##
## Linear Hypotheses:
##                Estimate Std. Error z value Pr(>|z|)
## OL - OF == 0 -2736575    5272014  -0.519    0.604
## (Adjusted p values reported -- single-step method)
##
##
## Tukey results for gene: THBS1
##
##   Simultaneous Tests for General Linear Hypotheses
##
## Multiple Comparisons of Means: Tukey Contrasts
##
##
## Fit: glm(formula = Protein_Count ~ Tissue, family = gaussian, data = gene_
data)
##
## Linear Hypotheses:
##                Estimate Std. Error z value Pr(>|z|)
## OL - OF == 0  -510454     175649  -2.906  0.00366 **
## ---
## Signif. codes:  0 '***' 0.001 '**' 0.01 '*' 0.05 '.' 0.1 ' ' 1
```

```
## (Adjusted p values reported -- single-step method)
##
##
## Tukey results for gene: THBS2
##
##   Simultaneous Tests for General Linear Hypotheses
##
## Multiple Comparisons of Means: Tukey Contrasts
##
##
## Fit: glm(formula = Protein_Count ~ Tissue, family = gaussian, data = gene_
data)
##
## Linear Hypotheses:
##               Estimate Std. Error z value Pr(>|z|)
## OL - OF == 0  -244093     208199  -1.172    0.241
## (Adjusted p values reported -- single-step method)
##
##
## Tukey results for gene: THBS3
##
##   Simultaneous Tests for General Linear Hypotheses
##
## Multiple Comparisons of Means: Tukey Contrasts
##
##
## Fit: glm(formula = Protein_Count ~ Tissue, family = gaussian, data = gene_
data)
##
## Linear Hypotheses:
##               Estimate Std. Error z value Pr(>|z|)
## OL - OF == 0    16284      36772   0.443    0.658
## (Adjusted p values reported -- single-step method)
##
##
## Tukey results for gene: THSD4
##
##   Simultaneous Tests for General Linear Hypotheses
##
## Multiple Comparisons of Means: Tukey Contrasts
##
##
## Fit: glm(formula = Protein_Count ~ Tissue, family = gaussian, data = gene_
data)
##
## Linear Hypotheses:
##               Estimate Std. Error z value Pr(>|z|)
## OL - OF == 0   345473     971263   0.356    0.722
## (Adjusted p values reported -- single-step method)
##
```

```
##
## Tukey results for gene: TINAGL1
##
##   Simultaneous Tests for General Linear Hypotheses
##
## Multiple Comparisons of Means: Tukey Contrasts
##
##
## Fit: glm(formula = Protein_Count ~ Tissue, family = gaussian, data = gene_
data)
##
## Linear Hypotheses:
##              Estimate Std. Error z value Pr(>|z|)
## OL - OF == 0  7414717   14325547   0.518    0.605
## (Adjusted p values reported -- single-step method)
##
##
## Tukey results for gene: TNC
##
##   Simultaneous Tests for General Linear Hypotheses
##
## Multiple Comparisons of Means: Tukey Contrasts
##
##
## Fit: glm(formula = Protein_Count ~ Tissue, family = gaussian, data = gene_
data)
##
## Linear Hypotheses:
##              Estimate Std. Error z value Pr(>|z|)
## OL - OF == 0   291525     819780   0.356    0.722
## (Adjusted p values reported -- single-step method)
##
##
## Tukey results for gene: TNXB
##
##   Simultaneous Tests for General Linear Hypotheses
##
## Multiple Comparisons of Means: Tukey Contrasts
##
##
## Fit: glm(formula = Protein_Count ~ Tissue, family = gaussian, data = gene_
data)
##
## Linear Hypotheses:
##              Estimate Std. Error z value Pr(>|z|)
## OL - OF == 0  -136878    1038606  -0.132    0.895
## (Adjusted p values reported -- single-step method)
##
##
## Tukey results for gene: TSKU
```

```
##
##   Simultaneous Tests for General Linear Hypotheses
##
## Multiple Comparisons of Means: Tukey Contrasts
##
##
## Fit: glm(formula = Protein_Count ~ Tissue, family = gaussian, data = gene_
data)
##
## Linear Hypotheses:
##              Estimate Std. Error z value Pr(>|z|)
## OL - OF == 0   -40240      81915  -0.491    0.623
## (Adjusted p values reported -- single-step method)
##
##
## Tukey results for gene: VTN
##
##   Simultaneous Tests for General Linear Hypotheses
##
## Multiple Comparisons of Means: Tukey Contrasts
##
##
## Fit: glm(formula = Protein_Count ~ Tissue, family = gaussian, data = gene_
data)
##
## Linear Hypotheses:
##              Estimate Std. Error z value Pr(>|z|)
## OL - OF == 0   946075    6138454   0.154    0.878
## (Adjusted p values reported -- single-step method)
##
##
## Tukey results for gene: VWA1
##
##   Simultaneous Tests for General Linear Hypotheses
##
## Multiple Comparisons of Means: Tukey Contrasts
##
##
## Fit: glm(formula = Protein_Count ~ Tissue, family = gaussian, data = gene_
data)
##
## Linear Hypotheses:
##              Estimate Std. Error z value Pr(>|z|)
## OL - OF == 0   720877    2637120   0.273    0.785
## (Adjusted p values reported -- single-step method)
##
##
## Tukey results for gene: VWA5A
##
##   Simultaneous Tests for General Linear Hypotheses
```

```
##
## Multiple Comparisons of Means: Tukey Contrasts
##
##
## Fit: glm(formula = Protein_Count ~ Tissue, family = gaussian, data = gene_
data)
##
## Linear Hypotheses:
##                Estimate Std. Error z value Pr(>|z|)
## OL - OF == 0   -25311      80253  -0.315    0.752
## (Adjusted p values reported -- single-step method)
##
##
## Tukey results for gene: VWF
##
##   Simultaneous Tests for General Linear Hypotheses
##
## Multiple Comparisons of Means: Tukey Contrasts
##
##
## Fit: glm(formula = Protein_Count ~ Tissue, family = gaussian, data = gene_
data)
##
## Linear Hypotheses:
##                Estimate Std. Error z value Pr(>|z|)
## OL - OF == 0  -136581     107994  -1.265    0.206
## (Adjusted p values reported -- single-step method)
```

```
#### Proteoglycans

# Read your cleaned joined data file into a data frame
cleaned_joined_data <- read.csv("cleaned_matrisome_oviduct.csv")

# Filter rows with Matrisome_Category "Proteoglycans"
Proteoglycans_data <- cleaned_joined_data %>%
  filter(Matrisome_Category == "Proteoglycans")

# Reshape the data to have separate columns for each replicate and each tissu
e stage
organized_data <- Proteoglycans_data %>%
  pivot_longer(cols = starts_with("OF") | starts_with("OL"),
               names_to = c("Tissue", "Stage"),
               names_sep = "\\.",
               values_to = "Protein_Count")
```

```
## Warning: Expected 2 pieces. Missing pieces filled with `NA` in 2 rows [1,
5].
```

```
# Save the reshaped data to a new file
```

```
write.csv(organized_data, "Proteoglycans_data_matrisome_oviduct_to_clean.csv"
, row.names = FALSE)

data2 <- read.csv("Proteoglycans_data_matrisome_oviduct_clean.csv")

png(filename="ovi_Proteoglycans.png",
    width = 12, height = 6, units = "in",
    res = 600)

boxplot(Protein_Count ~ Tissue*Genes,
        data = data2,
        cex.axis = 0.8,
        col = viridis(2, alpha = 0.7),
        lwd = 0.7,
        ylab = "",
        ylim = c(0, 200000000),
        axes = FALSE,
        frame.plot = TRUE,
        xlab = "")

GROUP <- unique(data2$Genes)
groups <- 15
numbox <- 2
total <- groups * numbox
xpoints <- seq(median(1:numbox), total, numbox)

axis(2, seq(0, 200000000, by = 10000000), las = 1, cex.axis = 0.7)
axis(1, labels = GROUP, at = xpoints, las = 2, cex.axis = 0.7)

stripchart(data2$Protein_Count ~ data2$Tissue*data2$Genes,
           method = "jitter",
           pch = 19,
           col = "black",
           vertical = TRUE,
           add = TRUE,
           cex = 0.3)

title(ylab = expression("Protein abundance (RU)"), line = 3.0)
legend("top", fill = viridis(2), legend = c("Oviduct Follicular", "Oviduct Lu
teal"), horiz = T, cex = 0.7)

dev.off()

## quartz_off_screen
##                  2

# Convert Tissue to factor
data2$Tissue <- as.factor(data2$Tissue)
```

```r
# Create an empty list to store Tukey results
tukey_results_list <- list()

# Unique genes
unique_genes <- unique(data2$Genes)

# Loop over each gene
for (gene in unique_genes) {
  # Subset data for the current gene
  gene_data <- subset(data2, Genes == gene)

  # Fit the generalized linear model
  glm_model <- glm(Protein_Count ~ Tissue, data = gene_data, family = gaussia
n)

  # Perform Tukey test
  contrast_matrix <- glht(glm_model, linfct = mcp(Tissue = "Tukey"))
  tukey_results <- summary(contrast_matrix)

  # Store the results
  tukey_results_list[[gene]] <- tukey_results
}

# Print the results for each gene
for (gene in unique_genes) {
  cat("Tukey results for gene:", gene, "\n")
  print(tukey_results_list[[gene]])
  cat("\n")
}

## Tukey results for gene: ACAN
##
##   Simultaneous Tests for General Linear Hypotheses
##
## Multiple Comparisons of Means: Tukey Contrasts
##
##
## Fit: glm(formula = Protein_Count ~ Tissue, family = gaussian, data = gene_
data)
##
## Linear Hypotheses:
##              Estimate Std. Error z value Pr(>|z|)
## OL - OF == 0   -71217     204424  -0.348    0.728
## (Adjusted p values reported -- single-step method)
##
##
## Tukey results for gene: ASPN
##
##   Simultaneous Tests for General Linear Hypotheses
```

```
##
## Multiple Comparisons of Means: Tukey Contrasts
##
##
## Fit: glm(formula = Protein_Count ~ Tissue, family = gaussian, data = gene_
data)
##
## Linear Hypotheses:
##               Estimate Std. Error z value Pr(>|z|)
## OL - OF == 0  -817333   10222653   -0.08    0.936
## (Adjusted p values reported -- single-step method)
##
##
## Tukey results for gene: BGN
##
##   Simultaneous Tests for General Linear Hypotheses
##
## Multiple Comparisons of Means: Tukey Contrasts
##
##
## Fit: glm(formula = Protein_Count ~ Tissue, family = gaussian, data = gene_
data)
##
## Linear Hypotheses:
##               Estimate Std. Error z value Pr(>|z|)
## OL - OF == 0  -632108    1972105  -0.321    0.749
## (Adjusted p values reported -- single-step method)
##
##
## Tukey results for gene: DCN
##
##   Simultaneous Tests for General Linear Hypotheses
##
## Multiple Comparisons of Means: Tukey Contrasts
##
##
## Fit: glm(formula = Protein_Count ~ Tissue, family = gaussian, data = gene_
data)
##
## Linear Hypotheses:
##               Estimate Std. Error z value Pr(>|z|)
## OL - OF == 0   750862    2262125   0.332     0.74
## (Adjusted p values reported -- single-step method)
##
##
## Tukey results for gene: FMOD
##
##   Simultaneous Tests for General Linear Hypotheses
##
## Multiple Comparisons of Means: Tukey Contrasts
```

```
##
##
## Fit: glm(formula = Protein_Count ~ Tissue, family = gaussian, data = gene_
data)
##
## Linear Hypotheses:
##              Estimate Std. Error z value Pr(>|z|)
## OL - OF == 0   566002    1667949   0.339    0.734
## (Adjusted p values reported -- single-step method)
##
##
## Tukey results for gene: HAPLN1
##
##   Simultaneous Tests for General Linear Hypotheses
##
## Multiple Comparisons of Means: Tukey Contrasts
##
##
## Fit: glm(formula = Protein_Count ~ Tissue, family = gaussian, data = gene_
data)
##
## Linear Hypotheses:
##              Estimate Std. Error z value Pr(>|z|)
## OL - OF == 0  -412201     168417  -2.448   0.0144 *
## ---
## Signif. codes:  0 '***' 0.001 '**' 0.01 '*' 0.05 '.' 0.1 ' ' 1
## (Adjusted p values reported -- single-step method)
##
##
## Tukey results for gene: HAPLN3
##
##   Simultaneous Tests for General Linear Hypotheses
##
## Multiple Comparisons of Means: Tukey Contrasts
##
##
## Fit: glm(formula = Protein_Count ~ Tissue, family = gaussian, data = gene_
data)
##
## Linear Hypotheses:
##              Estimate Std. Error z value Pr(>|z|)
## OL - OF == 0   -14378      59273  -0.243    0.808
## (Adjusted p values reported -- single-step method)
##
##
## Tukey results for gene: HSPG2
##
##   Simultaneous Tests for General Linear Hypotheses
##
## Multiple Comparisons of Means: Tukey Contrasts
```

```
##
##
## Fit: glm(formula = Protein_Count ~ Tissue, family = gaussian, data = gene_
data)
##
## Linear Hypotheses:
##               Estimate Std. Error z value Pr(>|z|)
## OL - OF == 0  2856317    4832503   0.591    0.554
## (Adjusted p values reported -- single-step method)
##
##
## Tukey results for gene: KERA
##
##   Simultaneous Tests for General Linear Hypotheses
##
## Multiple Comparisons of Means: Tukey Contrasts
##
##
## Fit: glm(formula = Protein_Count ~ Tissue, family = gaussian, data = gene_
data)
##
## Linear Hypotheses:
##               Estimate Std. Error z value Pr(>|z|)
## OL - OF == 0    22182     100038   0.222    0.825
## (Adjusted p values reported -- single-step method)
##
##
## Tukey results for gene: LUM
##
##   Simultaneous Tests for General Linear Hypotheses
##
## Multiple Comparisons of Means: Tukey Contrasts
##
##
## Fit: glm(formula = Protein_Count ~ Tissue, family = gaussian, data = gene_
data)
##
## Linear Hypotheses:
##               Estimate Std. Error z value Pr(>|z|)
## OL - OF == 0  -933525   28410280  -0.033    0.974
## (Adjusted p values reported -- single-step method)
##
##
## Tukey results for gene: OGN
##
##   Simultaneous Tests for General Linear Hypotheses
##
## Multiple Comparisons of Means: Tukey Contrasts
##
##
```

```
## Fit: glm(formula = Protein_Count ~ Tissue, family = gaussian, data = gene_
data)
##
## Linear Hypotheses:
##              Estimate Std. Error z value Pr(>|z|)
## OL - OF == 0 -7881133   23233921  -0.339    0.734
## (Adjusted p values reported -- single-step method)
##
##
## Tukey results for gene: PODN
##
##   Simultaneous Tests for General Linear Hypotheses
##
## Multiple Comparisons of Means: Tukey Contrasts
##
##
## Fit: glm(formula = Protein_Count ~ Tissue, family = gaussian, data = gene_
data)
##
## Linear Hypotheses:
##              Estimate Std. Error z value Pr(>|z|)
## OL - OF == 0  -101849     337730  -0.302    0.763
## (Adjusted p values reported -- single-step method)
##
##
## Tukey results for gene: PODNL1
##
##   Simultaneous Tests for General Linear Hypotheses
##
## Multiple Comparisons of Means: Tukey Contrasts
##
##
## Fit: glm(formula = Protein_Count ~ Tissue, family = gaussian, data = gene_
data)
##
## Linear Hypotheses:
##              Estimate Std. Error z value Pr(>|z|)
## OL - OF == 0   -27996     117925  -0.237    0.812
## (Adjusted p values reported -- single-step method)
##
##
## Tukey results for gene: PRELP
##
##   Simultaneous Tests for General Linear Hypotheses
##
## Multiple Comparisons of Means: Tukey Contrasts
##
##
## Fit: glm(formula = Protein_Count ~ Tissue, family = gaussian, data = gene_
data)
```

```
##
## Linear Hypotheses:
##              Estimate Std. Error z value Pr(>|z|)
## OL - OF == 0  1238284    6599541   0.188    0.851
## (Adjusted p values reported -- single-step method)
##
##
## Tukey results for gene: VCAN
##
##   Simultaneous Tests for General Linear Hypotheses
##
## Multiple Comparisons of Means: Tukey Contrasts
##
##
## Fit: glm(formula = Protein_Count ~ Tissue, family = gaussian, data = gene_
data)
##
## Linear Hypotheses:
##              Estimate Std. Error z value Pr(>|z|)
## OL - OF == 0 -1273503    3025296  -0.421    0.674
## (Adjusted p values reported -- single-step method)

#### ECM regulators

# Read your cleaned joined data file into a data frame
cleaned_joined_data <- read.csv("cleaned_matrisome_oviduct.csv")

# Filter rows with Matrisome_Category "ECM regulators"
ECM_regulators_data <- cleaned_joined_data %>%
  filter(Matrisome_Category == "ECM Regulators")

# Reshape the data to have separate columns for each replicate and each tissu
e stage
organized_data <- ECM_regulators_data %>%
  pivot_longer(cols = starts_with("OF") | starts_with("OL"),
               names_to = c("Tissue", "Stage"),
               names_sep = "\\.",
               values_to = "Protein_Count")

## Warning: Expected 2 pieces. Missing pieces filled with `NA` in 2 rows [1,
5].

# Save the reshaped data to a new file

write.csv(organized_data, "ECM regulators_data_matrisome_oviduct_to_clean.csv
", row.names = FALSE)

data2 <- read.csv("ECM regulators_data_matrisome_oviduct_clean.csv")

png(filename="ovi_ECM regulators.png",
```

```
    width = 12, height = 6, units = "in",
    res = 600)

boxplot(Protein_Count ~ Tissue*Genes,
        data = data2,
        cex.axis = 0.8,
        col = viridis(2, alpha = 0.7),
        lwd = 0.7,
        ylab = "",
        ylim = c(0, 18000000),
        axes = FALSE,
        frame.plot = TRUE,
        xlab = "")

GROUP <- unique(data2$Genes)
groups <- 63
numbox <- 2
total <- groups * numbox
xpoints <- seq(median(1:numbox), total, numbox)

axis(2, seq(0, 18000000, by = 2000000), las = 1, cex.axis = 0.7)
axis(1, labels = GROUP, at = xpoints, las = 2, cex.axis = 0.7)

stripchart(data2$Protein_Count ~ data2$Tissue*data2$Genes,
           method = "jitter",
           pch = 19,
           col = "black",
           vertical = TRUE,
           add = TRUE,
           cex = 0.3)

title(ylab = expression("Protein abundance (RU)"), line = 3.0)
legend("top", fill = viridis(2), legend = c("Oviduct Follicular", "Oviduct Lu
teal"), horiz = T, cex = 0.7)

dev.off()

## quartz_off_screen
##                  2

# Convert Tissue to factor
data2$Tissue <- as.factor(data2$Tissue)

# Create an empty list to store Tukey results
tukey_results_list <- list()

# Unique genes
unique_genes <- unique(data2$Genes)
```

```
# Loop over each gene
for (gene in unique_genes) {
  # Subset data for the current gene
  gene_data <- subset(data2, Genes == gene)

  # Fit the generalized linear model
  glm_model <- glm(Protein_Count ~ Tissue, data = gene_data, family = gaussia
n)

  # Perform Tukey test
  contrast_matrix <- glht(glm_model, linfct = mcp(Tissue = "Tukey"))
  tukey_results <- summary(contrast_matrix)

  # Store the results
  tukey_results_list[[gene]] <- tukey_results
}

# Print the results for each gene
for (gene in unique_genes) {
  cat("Tukey results for gene:", gene, "\n")
  print(tukey_results_list[[gene]])
  cat("\n")
}

## Tukey results for gene: A2M
##
##   Simultaneous Tests for General Linear Hypotheses
##
## Multiple Comparisons of Means: Tukey Contrasts
##
##
## Fit: glm(formula = Protein_Count ~ Tissue, family = gaussian, data = gene_
data)
##
## Linear Hypotheses:
##                Estimate Std. Error z value Pr(>|z|)
## OL - OF == 0  -314780     748293  -0.421    0.674
## (Adjusted p values reported -- single-step method)
##
##
## Tukey results for gene: ADAMTS5
##
##   Simultaneous Tests for General Linear Hypotheses
##
## Multiple Comparisons of Means: Tukey Contrasts
##
##
## Fit: glm(formula = Protein_Count ~ Tissue, family = gaussian, data = gene_
data)
##
```

```
## Linear Hypotheses:
##               Estimate Std. Error z value Pr(>|z|)
## OL - OF == 0     6541      97478   0.067    0.946
## (Adjusted p values reported -- single-step method)
##
##
## Tukey results for gene: ADAMTSL3
##
##   Simultaneous Tests for General Linear Hypotheses
##
## Multiple Comparisons of Means: Tukey Contrasts
##
##
## Fit: glm(formula = Protein_Count ~ Tissue, family = gaussian, data = gene_
data)
##
## Linear Hypotheses:
##               Estimate Std. Error z value Pr(>|z|)
## OL - OF == 0   -49268      96507  -0.511     0.61
## (Adjusted p values reported -- single-step method)
##
##
## Tukey results for gene: ADAMTSL4
##
##   Simultaneous Tests for General Linear Hypotheses
##
## Multiple Comparisons of Means: Tukey Contrasts
##
##
## Fit: glm(formula = Protein_Count ~ Tissue, family = gaussian, data = gene_
data)
##
## Linear Hypotheses:
##               Estimate Std. Error z value Pr(>|z|)
## OL - OF == 0    57674      79427   0.726    0.468
## (Adjusted p values reported -- single-step method)
##
##
## Tukey results for gene: ADAMTSL5
##
##   Simultaneous Tests for General Linear Hypotheses
##
## Multiple Comparisons of Means: Tukey Contrasts
##
##
## Fit: glm(formula = Protein_Count ~ Tissue, family = gaussian, data = gene_
data)
##
## Linear Hypotheses:
##               Estimate Std. Error z value Pr(>|z|)
```

```
## OL - OF == 0    14625     106391   0.137    0.891
## (Adjusted p values reported -- single-step method)
##
##
## Tukey results for gene: AGT
##
##   Simultaneous Tests for General Linear Hypotheses
##
## Multiple Comparisons of Means: Tukey Contrasts
##
##
## Fit: glm(formula = Protein_Count ~ Tissue, family = gaussian, data = gene_
data)
##
## Linear Hypotheses:
##              Estimate Std. Error z value Pr(>|z|)
## OL - OF == 0    20503     142965   0.143    0.886
## (Adjusted p values reported -- single-step method)
##
##
## Tukey results for gene: CD109
##
##   Simultaneous Tests for General Linear Hypotheses
##
## Multiple Comparisons of Means: Tukey Contrasts
##
##
## Fit: glm(formula = Protein_Count ~ Tissue, family = gaussian, data = gene_
data)
##
## Linear Hypotheses:
##              Estimate Std. Error z value Pr(>|z|)
## OL - OF == 0    20472     108866   0.188    0.851
## (Adjusted p values reported -- single-step method)
##
##
## Tukey results for gene: CELA1
##
##   Simultaneous Tests for General Linear Hypotheses
##
## Multiple Comparisons of Means: Tukey Contrasts
##
##
## Fit: glm(formula = Protein_Count ~ Tissue, family = gaussian, data = gene_
data)
##
## Linear Hypotheses:
##              Estimate Std. Error z value Pr(>|z|)
## OL - OF == 0   -36371      92878  -0.392    0.695
## (Adjusted p values reported -- single-step method)
```

```
##
##
## Tukey results for gene: CELA3B
##
##   Simultaneous Tests for General Linear Hypotheses
##
## Multiple Comparisons of Means: Tukey Contrasts
##
##
## Fit: glm(formula = Protein_Count ~ Tissue, family = gaussian, data = gene_
data)
##
## Linear Hypotheses:
##              Estimate Std. Error z value Pr(>|z|)
## OL - OF == 0  1426345    3257725   0.438    0.662
## (Adjusted p values reported -- single-step method)
##
##
## Tukey results for gene: CST6
##
##   Simultaneous Tests for General Linear Hypotheses
##
## Multiple Comparisons of Means: Tukey Contrasts
##
##
## Fit: glm(formula = Protein_Count ~ Tissue, family = gaussian, data = gene_
data)
##
## Linear Hypotheses:
##              Estimate Std. Error z value Pr(>|z|)
## OL - OF == 0   -24746      76920  -0.322    0.748
## (Adjusted p values reported -- single-step method)
##
##
## Tukey results for gene: CSTB
##
##   Simultaneous Tests for General Linear Hypotheses
##
## Multiple Comparisons of Means: Tukey Contrasts
##
##
## Fit: glm(formula = Protein_Count ~ Tissue, family = gaussian, data = gene_
data)
##
## Linear Hypotheses:
##              Estimate Std. Error z value Pr(>|z|)
## OL - OF == 0   -79822     296772  -0.269    0.788
## (Adjusted p values reported -- single-step method)
##
##
```

```
## Tukey results for gene: CTSC
##
##   Simultaneous Tests for General Linear Hypotheses
##
## Multiple Comparisons of Means: Tukey Contrasts
##
##
## Fit: glm(formula = Protein_Count ~ Tissue, family = gaussian, data = gene_
data)
##
## Linear Hypotheses:
##              Estimate Std. Error z value Pr(>|z|)
## OL - OF == 0    232.2    59387.3   0.004    0.997
## (Adjusted p values reported -- single-step method)
##
##
## Tukey results for gene: CTSD
##
##   Simultaneous Tests for General Linear Hypotheses
##
## Multiple Comparisons of Means: Tukey Contrasts
##
##
## Fit: glm(formula = Protein_Count ~ Tissue, family = gaussian, data = gene_
data)
##
## Linear Hypotheses:
##              Estimate Std. Error z value Pr(>|z|)
## OL - OF == 0   -25236      50251  -0.502    0.616
## (Adjusted p values reported -- single-step method)
##
##
## Tukey results for gene: CTSF
##
##   Simultaneous Tests for General Linear Hypotheses
##
## Multiple Comparisons of Means: Tukey Contrasts
##
##
## Fit: glm(formula = Protein_Count ~ Tissue, family = gaussian, data = gene_
data)
##
## Linear Hypotheses:
##              Estimate Std. Error z value Pr(>|z|)
## OL - OF == 0  -133331     207277  -0.643     0.52
## (Adjusted p values reported -- single-step method)
##
##
## Tukey results for gene: CTSH
##
```

```
##   Simultaneous Tests for General Linear Hypotheses
##
## Multiple Comparisons of Means: Tukey Contrasts
##
##
## Fit: glm(formula = Protein_Count ~ Tissue, family = gaussian, data = gene_
data)
##
## Linear Hypotheses:
##              Estimate Std. Error z value Pr(>|z|)
## OL - OF == 0    45167     107789   0.419    0.675
## (Adjusted p values reported -- single-step method)
##
##
## Tukey results for gene: CTSS
##
##   Simultaneous Tests for General Linear Hypotheses
##
## Multiple Comparisons of Means: Tukey Contrasts
##
##
## Fit: glm(formula = Protein_Count ~ Tissue, family = gaussian, data = gene_
data)
##
## Linear Hypotheses:
##              Estimate Std. Error z value Pr(>|z|)
## OL - OF == 0  -117717      56561  -2.081   0.0374 *
## ---
## Signif. codes:  0 '***' 0.001 '**' 0.01 '*' 0.05 '.' 0.1 ' ' 1
## (Adjusted p values reported -- single-step method)
##
##
## Tukey results for gene: CTSW
##
##   Simultaneous Tests for General Linear Hypotheses
##
## Multiple Comparisons of Means: Tukey Contrasts
##
##
## Fit: glm(formula = Protein_Count ~ Tissue, family = gaussian, data = gene_
data)
##
## Linear Hypotheses:
##              Estimate Std. Error z value Pr(>|z|)
## OL - OF == 0   -96597     186768  -0.517    0.605
## (Adjusted p values reported -- single-step method)
##
##
## Tukey results for gene: CTSZ
##
```

```
##   Simultaneous Tests for General Linear Hypotheses
##
## Multiple Comparisons of Means: Tukey Contrasts
##
##
## Fit: glm(formula = Protein_Count ~ Tissue, family = gaussian, data = gene_
data)
##
## Linear Hypotheses:
##              Estimate Std. Error z value Pr(>|z|)
## OL - OF == 0   -72220      77321  -0.934     0.35
## (Adjusted p values reported -- single-step method)
##
##
## Tukey results for gene: EGLN1
##
##   Simultaneous Tests for General Linear Hypotheses
##
## Multiple Comparisons of Means: Tukey Contrasts
##
##
## Fit: glm(formula = Protein_Count ~ Tissue, family = gaussian, data = gene_
data)
##
## Linear Hypotheses:
##              Estimate Std. Error z value Pr(>|z|)
## OL - OF == 0     5534      37373   0.148    0.882
## (Adjusted p values reported -- single-step method)
##
##
## Tukey results for gene: F10
##
##   Simultaneous Tests for General Linear Hypotheses
##
## Multiple Comparisons of Means: Tukey Contrasts
##
##
## Fit: glm(formula = Protein_Count ~ Tissue, family = gaussian, data = gene_
data)
##
## Linear Hypotheses:
##              Estimate Std. Error z value Pr(>|z|)
## OL - OF == 0     9510      71430   0.133    0.894
## (Adjusted p values reported -- single-step method)
##
##
## Tukey results for gene: F13B
##
##   Simultaneous Tests for General Linear Hypotheses
##
```

```
## Multiple Comparisons of Means: Tukey Contrasts
##
##
## Fit: glm(formula = Protein_Count ~ Tissue, family = gaussian, data = gene_
data)
##
## Linear Hypotheses:
##              Estimate Std. Error z value Pr(>|z|)
## OL - OF == 0    -5693      80723  -0.071    0.944
## (Adjusted p values reported -- single-step method)
##
##
## Tukey results for gene: F2
##
##   Simultaneous Tests for General Linear Hypotheses
##
## Multiple Comparisons of Means: Tukey Contrasts
##
##
## Fit: glm(formula = Protein_Count ~ Tissue, family = gaussian, data = gene_
data)
##
## Linear Hypotheses:
##              Estimate Std. Error z value Pr(>|z|)
## OL - OF == 0   -57300     148814  -0.385      0.7
## (Adjusted p values reported -- single-step method)
##
##
## Tukey results for gene: F9
##
##   Simultaneous Tests for General Linear Hypotheses
##
## Multiple Comparisons of Means: Tukey Contrasts
##
##
## Fit: glm(formula = Protein_Count ~ Tissue, family = gaussian, data = gene_
data)
##
## Linear Hypotheses:
##              Estimate Std. Error z value Pr(>|z|)
## OL - OF == 0     7553      64475   0.117    0.907
## (Adjusted p values reported -- single-step method)
##
##
## Tukey results for gene: FETUB
##
##   Simultaneous Tests for General Linear Hypotheses
##
## Multiple Comparisons of Means: Tukey Contrasts
##
```

```
##
## Fit: glm(formula = Protein_Count ~ Tissue, family = gaussian, data = gene_
data)
##
## Linear Hypotheses:
##              Estimate Std. Error z value Pr(>|z|)
## OL - OF == 0    27461      29965   0.916    0.359
## (Adjusted p values reported -- single-step method)
##
##
## Tukey results for gene: HRG
##
##   Simultaneous Tests for General Linear Hypotheses
##
## Multiple Comparisons of Means: Tukey Contrasts
##
##
## Fit: glm(formula = Protein_Count ~ Tissue, family = gaussian, data = gene_
data)
##
## Linear Hypotheses:
##              Estimate Std. Error z value Pr(>|z|)
## OL - OF == 0   -67162     588271  -0.114    0.909
## (Adjusted p values reported -- single-step method)
##
##
## Tukey results for gene: HTRA1
##
##   Simultaneous Tests for General Linear Hypotheses
##
## Multiple Comparisons of Means: Tukey Contrasts
##
##
## Fit: glm(formula = Protein_Count ~ Tissue, family = gaussian, data = gene_
data)
##
## Linear Hypotheses:
##              Estimate Std. Error z value Pr(>|z|)
## OL - OF == 0    -9097      48118  -0.189     0.85
## (Adjusted p values reported -- single-step method)
##
##
## Tukey results for gene: HTRA3
##
##   Simultaneous Tests for General Linear Hypotheses
##
## Multiple Comparisons of Means: Tukey Contrasts
##
##
## Fit: glm(formula = Protein_Count ~ Tissue, family = gaussian, data = gene_
```

```
data)
##
## Linear Hypotheses:
##               Estimate Std. Error z value Pr(>|z|)
## OL - OF == 0   -24962      56031  -0.446    0.656
## (Adjusted p values reported -- single-step method)
##
##
## Tukey results for gene: ITIH1
##
##   Simultaneous Tests for General Linear Hypotheses
##
## Multiple Comparisons of Means: Tukey Contrasts
##
##
## Fit: glm(formula = Protein_Count ~ Tissue, family = gaussian, data = gene_
data)
##
## Linear Hypotheses:
##               Estimate Std. Error z value Pr(>|z|)
## OL - OF == 0   315753    1173837   0.269    0.788
## (Adjusted p values reported -- single-step method)
##
##
## Tukey results for gene: ITIH2
##
##   Simultaneous Tests for General Linear Hypotheses
##
## Multiple Comparisons of Means: Tukey Contrasts
##
##
## Fit: glm(formula = Protein_Count ~ Tissue, family = gaussian, data = gene_
data)
##
## Linear Hypotheses:
##               Estimate Std. Error z value Pr(>|z|)
## OL - OF == 0   174137     543803    0.32    0.749
## (Adjusted p values reported -- single-step method)
##
##
## Tukey results for gene: ITIH4
##
##   Simultaneous Tests for General Linear Hypotheses
##
## Multiple Comparisons of Means: Tukey Contrasts
##
##
## Fit: glm(formula = Protein_Count ~ Tissue, family = gaussian, data = gene_
data)
##
```

```
## Linear Hypotheses:
##              Estimate Std. Error z value Pr(>|z|)
## OL - OF == 0   -11297     390929  -0.029    0.977
## (Adjusted p values reported -- single-step method)
##
##
## Tukey results for gene: ITIH5
##
##   Simultaneous Tests for General Linear Hypotheses
##
## Multiple Comparisons of Means: Tukey Contrasts
##
##
## Fit: glm(formula = Protein_Count ~ Tissue, family = gaussian, data = gene_
data)
##
## Linear Hypotheses:
##              Estimate Std. Error z value Pr(>|z|)
## OL - OF == 0    61371     103710   0.592    0.554
## (Adjusted p values reported -- single-step method)
##
##
## Tukey results for gene: KNG1
##
##   Simultaneous Tests for General Linear Hypotheses
##
## Multiple Comparisons of Means: Tukey Contrasts
##
##
## Fit: glm(formula = Protein_Count ~ Tissue, family = gaussian, data = gene_
data)
##
## Linear Hypotheses:
##              Estimate Std. Error z value Pr(>|z|)
## OL - OF == 0  -146144     721069  -0.203    0.839
## (Adjusted p values reported -- single-step method)
##
##
## Tukey results for gene: KNG2
##
##   Simultaneous Tests for General Linear Hypotheses
##
## Multiple Comparisons of Means: Tukey Contrasts
##
##
## Fit: glm(formula = Protein_Count ~ Tissue, family = gaussian, data = gene_
data)
##
## Linear Hypotheses:
##              Estimate Std. Error z value Pr(>|z|)
```

```
## OL - OF == 0   -43584      39033  -1.117    0.264
## (Adjusted p values reported -- single-step method)
##
##
## Tukey results for gene: LOX
##
##   Simultaneous Tests for General Linear Hypotheses
##
## Multiple Comparisons of Means: Tukey Contrasts
##
##
## Fit: glm(formula = Protein_Count ~ Tissue, family = gaussian, data = gene_
data)
##
## Linear Hypotheses:
##              Estimate Std. Error z value Pr(>|z|)
## OL - OF == 0    91158     799180   0.114    0.909
## (Adjusted p values reported -- single-step method)
##
##
## Tukey results for gene: LOXL1
##
##   Simultaneous Tests for General Linear Hypotheses
##
## Multiple Comparisons of Means: Tukey Contrasts
##
##
## Fit: glm(formula = Protein_Count ~ Tissue, family = gaussian, data = gene_
data)
##
## Linear Hypotheses:
##              Estimate Std. Error z value Pr(>|z|)
## OL - OF == 0    90877     455686   0.199    0.842
## (Adjusted p values reported -- single-step method)
##
##
## Tukey results for gene: LOXL3
##
##   Simultaneous Tests for General Linear Hypotheses
##
## Multiple Comparisons of Means: Tukey Contrasts
##
##
## Fit: glm(formula = Protein_Count ~ Tissue, family = gaussian, data = gene_
data)
##
## Linear Hypotheses:
##              Estimate Std. Error z value Pr(>|z|)
## OL - OF == 0    38169      65887   0.579    0.562
## (Adjusted p values reported -- single-step method)
```

```
##
##
## Tukey results for gene: MASP2
##
##   Simultaneous Tests for General Linear Hypotheses
##
## Multiple Comparisons of Means: Tukey Contrasts
##
##
## Fit: glm(formula = Protein_Count ~ Tissue, family = gaussian, data = gene_
data)
##
## Linear Hypotheses:
##              Estimate Std. Error z value Pr(>|z|)
## OL - OF == 0  -358028     344248   -1.04    0.298
## (Adjusted p values reported -- single-step method)
##
##
## Tukey results for gene: MMP2
##
##   Simultaneous Tests for General Linear Hypotheses
##
## Multiple Comparisons of Means: Tukey Contrasts
##
##
## Fit: glm(formula = Protein_Count ~ Tissue, family = gaussian, data = gene_
data)
##
## Linear Hypotheses:
##              Estimate Std. Error z value Pr(>|z|)
## OL - OF == 0   -44514      75988  -0.586    0.558
## (Adjusted p values reported -- single-step method)
##
##
## Tukey results for gene: MMP23
##
##   Simultaneous Tests for General Linear Hypotheses
##
## Multiple Comparisons of Means: Tukey Contrasts
##
##
## Fit: glm(formula = Protein_Count ~ Tissue, family = gaussian, data = gene_
data)
##
## Linear Hypotheses:
##              Estimate Std. Error z value Pr(>|z|)
## OL - OF == 0    32459     125857   0.258    0.796
## (Adjusted p values reported -- single-step method)
##
##
```

```
## Tukey results for gene: MMP28
##
##   Simultaneous Tests for General Linear Hypotheses
##
## Multiple Comparisons of Means: Tukey Contrasts
##
##
## Fit: glm(formula = Protein_Count ~ Tissue, family = gaussian, data = gene_
data)
##
## Linear Hypotheses:
##              Estimate Std. Error z value Pr(>|z|)
## OL - OF == 0   -25748      91547  -0.281    0.779
## (Adjusted p values reported -- single-step method)
##
##
## Tukey results for gene: MMP3
##
##   Simultaneous Tests for General Linear Hypotheses
##
## Multiple Comparisons of Means: Tukey Contrasts
##
##
## Fit: glm(formula = Protein_Count ~ Tissue, family = gaussian, data = gene_
data)
##
## Linear Hypotheses:
##              Estimate Std. Error z value Pr(>|z|)
## OL - OF == 0   -36008      80673  -0.446    0.655
## (Adjusted p values reported -- single-step method)
##
##
## Tukey results for gene: P4HA1
##
##   Simultaneous Tests for General Linear Hypotheses
##
## Multiple Comparisons of Means: Tukey Contrasts
##
##
## Fit: glm(formula = Protein_Count ~ Tissue, family = gaussian, data = gene_
data)
##
## Linear Hypotheses:
##              Estimate Std. Error z value Pr(>|z|)
## OL - OF == 0    68702     119270   0.576    0.565
## (Adjusted p values reported -- single-step method)
##
##
## Tukey results for gene: P4HA2
##
```

```
##   Simultaneous Tests for General Linear Hypotheses
##
## Multiple Comparisons of Means: Tukey Contrasts
##
##
## Fit: glm(formula = Protein_Count ~ Tissue, family = gaussian, data = gene_
data)
##
## Linear Hypotheses:
##              Estimate Std. Error z value Pr(>|z|)
## OL - OF == 0   -26685      28150  -0.948    0.343
## (Adjusted p values reported -- single-step method)
##
##
## Tukey results for gene: PLG
##
##   Simultaneous Tests for General Linear Hypotheses
##
## Multiple Comparisons of Means: Tukey Contrasts
##
##
## Fit: glm(formula = Protein_Count ~ Tissue, family = gaussian, data = gene_
data)
##
## Linear Hypotheses:
##              Estimate Std. Error z value Pr(>|z|)
## OL - OF == 0  -154193     580129  -0.266     0.79
## (Adjusted p values reported -- single-step method)
##
##
## Tukey results for gene: PLOD1
##
##   Simultaneous Tests for General Linear Hypotheses
##
## Multiple Comparisons of Means: Tukey Contrasts
##
##
## Fit: glm(formula = Protein_Count ~ Tissue, family = gaussian, data = gene_
data)
##
## Linear Hypotheses:
##              Estimate Std. Error z value Pr(>|z|)
## OL - OF == 0  -123573     108473  -1.139    0.255
## (Adjusted p values reported -- single-step method)
##
##
## Tukey results for gene: PLOD3
##
##   Simultaneous Tests for General Linear Hypotheses
##
```

```
## Multiple Comparisons of Means: Tukey Contrasts
##
##
## Fit: glm(formula = Protein_Count ~ Tissue, family = gaussian, data = gene_
data)
##
## Linear Hypotheses:
##               Estimate Std. Error z value Pr(>|z|)
## OL - OF == 0   -52877      72408   -0.73    0.465
## (Adjusted p values reported -- single-step method)
##
##
## Tukey results for gene: PRSS1
##
##   Simultaneous Tests for General Linear Hypotheses
##
## Multiple Comparisons of Means: Tukey Contrasts
##
##
## Fit: glm(formula = Protein_Count ~ Tissue, family = gaussian, data = gene_
data)
##
## Linear Hypotheses:
##               Estimate Std. Error z value Pr(>|z|)
## OL - OF == 0    66192    1534614   0.043    0.966
## (Adjusted p values reported -- single-step method)
##
##
## Tukey results for gene: SERPINA1
##
##   Simultaneous Tests for General Linear Hypotheses
##
## Multiple Comparisons of Means: Tukey Contrasts
##
##
## Fit: glm(formula = Protein_Count ~ Tissue, family = gaussian, data = gene_
data)
##
## Linear Hypotheses:
##               Estimate Std. Error z value Pr(>|z|)
## OL - OF == 0    44586      74480   0.599    0.549
## (Adjusted p values reported -- single-step method)
##
##
## Tukey results for gene: SERPINA10
##
##   Simultaneous Tests for General Linear Hypotheses
##
## Multiple Comparisons of Means: Tukey Contrasts
##
```

```
##
## Fit: glm(formula = Protein_Count ~ Tissue, family = gaussian, data = gene_
data)
##
## Linear Hypotheses:
##              Estimate Std. Error z value Pr(>|z|)
## OL - OF == 0   -34092      31861   -1.07    0.285
## (Adjusted p values reported -- single-step method)
##
##
## Tukey results for gene: SERPINA5
##
##   Simultaneous Tests for General Linear Hypotheses
##
## Multiple Comparisons of Means: Tukey Contrasts
##
##
## Fit: glm(formula = Protein_Count ~ Tissue, family = gaussian, data = gene_
data)
##
## Linear Hypotheses:
##              Estimate Std. Error z value Pr(>|z|)
## OL - OF == 0   -27187      67078  -0.405    0.685
## (Adjusted p values reported -- single-step method)
##
##
## Tukey results for gene: SERPINA7
##
##   Simultaneous Tests for General Linear Hypotheses
##
## Multiple Comparisons of Means: Tukey Contrasts
##
##
## Fit: glm(formula = Protein_Count ~ Tissue, family = gaussian, data = gene_
data)
##
## Linear Hypotheses:
##              Estimate Std. Error z value Pr(>|z|)
## OL - OF == 0   -19554      18865  -1.037      0.3
## (Adjusted p values reported -- single-step method)
##
##
## Tukey results for gene: SERPINB5
##
##   Simultaneous Tests for General Linear Hypotheses
##
## Multiple Comparisons of Means: Tukey Contrasts
##
##
## Fit: glm(formula = Protein_Count ~ Tissue, family = gaussian, data = gene_
```

```
data)
##
## Linear Hypotheses:
##                Estimate Std. Error z value Pr(>|z|)
## OL - OF == 0     2798      71520   0.039    0.969
## (Adjusted p values reported -- single-step method)
##
##
## Tukey results for gene: SERPINB6
##
##   Simultaneous Tests for General Linear Hypotheses
##
## Multiple Comparisons of Means: Tukey Contrasts
##
##
## Fit: glm(formula = Protein_Count ~ Tissue, family = gaussian, data = gene_
data)
##
## Linear Hypotheses:
##                Estimate Std. Error z value Pr(>|z|)
## OL - OF == 0   -55436     247330  -0.224    0.823
## (Adjusted p values reported -- single-step method)
##
##
## Tukey results for gene: SERPINC1
##
##   Simultaneous Tests for General Linear Hypotheses
##
## Multiple Comparisons of Means: Tukey Contrasts
##
##
## Fit: glm(formula = Protein_Count ~ Tissue, family = gaussian, data = gene_
data)
##
## Linear Hypotheses:
##                Estimate Std. Error z value Pr(>|z|)
## OL - OF == 0    44838     435192   0.103    0.918
## (Adjusted p values reported -- single-step method)
##
##
## Tukey results for gene: SERPIND1
##
##   Simultaneous Tests for General Linear Hypotheses
##
## Multiple Comparisons of Means: Tukey Contrasts
##
##
## Fit: glm(formula = Protein_Count ~ Tissue, family = gaussian, data = gene_
data)
##
```

```
## Linear Hypotheses:
##              Estimate Std. Error z value Pr(>|z|)
## OL - OF == 0   -18064      65601  -0.275    0.783
## (Adjusted p values reported -- single-step method)
##
##
## Tukey results for gene: SERPINF1
##
##   Simultaneous Tests for General Linear Hypotheses
##
## Multiple Comparisons of Means: Tukey Contrasts
##
##
## Fit: glm(formula = Protein_Count ~ Tissue, family = gaussian, data = gene_
data)
##
## Linear Hypotheses:
##              Estimate Std. Error z value Pr(>|z|)
## OL - OF == 0    83647      98580   0.849    0.396
## (Adjusted p values reported -- single-step method)
##
##
## Tukey results for gene: SERPINF2
##
##   Simultaneous Tests for General Linear Hypotheses
##
## Multiple Comparisons of Means: Tukey Contrasts
##
##
## Fit: glm(formula = Protein_Count ~ Tissue, family = gaussian, data = gene_
data)
##
## Linear Hypotheses:
##              Estimate Std. Error z value Pr(>|z|)
## OL - OF == 0    54872     328546   0.167    0.867
## (Adjusted p values reported -- single-step method)
##
##
## Tukey results for gene: SERPING1
##
##   Simultaneous Tests for General Linear Hypotheses
##
## Multiple Comparisons of Means: Tukey Contrasts
##
##
## Fit: glm(formula = Protein_Count ~ Tissue, family = gaussian, data = gene_
data)
##
## Linear Hypotheses:
##              Estimate Std. Error z value Pr(>|z|)
```

```
## OL - OF == 0   110658      78859   1.403    0.161
## (Adjusted p values reported -- single-step method)
##
##
## Tukey results for gene: SERPINH1
##
##   Simultaneous Tests for General Linear Hypotheses
##
## Multiple Comparisons of Means: Tukey Contrasts
##
##
## Fit: glm(formula = Protein_Count ~ Tissue, family = gaussian, data = gene_
data)
##
## Linear Hypotheses:
##               Estimate Std. Error z value Pr(>|z|)
## OL - OF == 0 -2643573    2040451  -1.296    0.195
## (Adjusted p values reported -- single-step method)
##
##
## Tukey results for gene: SERPINI2
##
##   Simultaneous Tests for General Linear Hypotheses
##
## Multiple Comparisons of Means: Tukey Contrasts
##
##
## Fit: glm(formula = Protein_Count ~ Tissue, family = gaussian, data = gene_
data)
##
## Linear Hypotheses:
##               Estimate Std. Error z value Pr(>|z|)
## OL - OF == 0   107069     368675    0.29    0.771
## (Adjusted p values reported -- single-step method)
##
##
## Tukey results for gene: TGM1
##
##   Simultaneous Tests for General Linear Hypotheses
##
## Multiple Comparisons of Means: Tukey Contrasts
##
##
## Fit: glm(formula = Protein_Count ~ Tissue, family = gaussian, data = gene_
data)
##
## Linear Hypotheses:
##               Estimate Std. Error z value Pr(>|z|)
## OL - OF == 0   -10174     184658  -0.055    0.956
## (Adjusted p values reported -- single-step method)
```

```
##
##
## Tukey results for gene: TGM2
##
##   Simultaneous Tests for General Linear Hypotheses
##
## Multiple Comparisons of Means: Tukey Contrasts
##
##
## Fit: glm(formula = Protein_Count ~ Tissue, family = gaussian, data = gene_
data)
##
## Linear Hypotheses:
##              Estimate Std. Error z value Pr(>|z|)
## OL - OF == 0 -5890583   53312277   -0.11    0.912
## (Adjusted p values reported -- single-step method)
##
##
## Tukey results for gene: TIMP2
##
##   Simultaneous Tests for General Linear Hypotheses
##
## Multiple Comparisons of Means: Tukey Contrasts
##
##
## Fit: glm(formula = Protein_Count ~ Tissue, family = gaussian, data = gene_
data)
##
## Linear Hypotheses:
##              Estimate Std. Error z value Pr(>|z|)
## OL - OF == 0    74671      76717   0.973     0.33
## (Adjusted p values reported -- single-step method)

#### ECM-affiliated

# Read your cleaned joined data file into a data frame
cleaned_joined_data <- read.csv("cleaned_matrisome_oviduct.csv")

# Filter rows with Matrisome_Category "ECM-affiliated"
ECMaffiliated_data <- cleaned_joined_data %>%
  filter(Matrisome_Category == "ECM-affiliated")

# Reshape the data to have separate columns for each replicate and each tissu
e stage
organized_data <- ECMaffiliated_data %>%
  pivot_longer(cols = starts_with("OF") | starts_with("OL"),
               names_to = c("Tissue", "Stage"),
               names_sep = "\\.",
               values_to = "Protein_Count")
```

```
## Warning: Expected 2 pieces. Missing pieces filled with `NA` in 2 rows [1,
5].

# Save the reshaped data to a new file

write.csv(organized_data, "ECM-affiliated_data_matrisome_oviduct_to_clean.csv
", row.names = FALSE)

data2 <- read.csv("ECM-affiliated_data_matrisome_oviduct_clean.csv")

png(filename="ovi_ECM-affiliated.png",
    width = 12, height = 6, units = "in",
    res = 600)

boxplot(Protein_Count ~ Tissue*Genes,
        data = data2,
        cex.axis = 0.8,
        col = viridis(2, alpha = 0.7),
        lwd = 0.7,
        ylab = "",
        ylim = c(0, 150000000),
        axes = FALSE,
        frame.plot = TRUE,
        xlab = "")

GROUP <- unique(data2$Genes)
groups <- 24
numbox <- 2
total <- groups * numbox
xpoints <- seq(median(1:numbox), total, numbox)

axis(2, seq(0, 150000000, by = 5000000), las = 1, cex.axis = 0.7)
axis(1, labels = GROUP, at = xpoints, las = 2, cex.axis = 0.7)

stripchart(data2$Protein_Count ~ data2$Tissue*data2$Genes,
           method = "jitter",
           pch = 19,
           col = "black",
           vertical = TRUE,
           add = TRUE,
           cex = 0.3)

title(ylab = expression("Protein abundance (RU)"), line = 3.0)
legend("top", fill = viridis(2), legend = c("Oviduct Follicular", "Oviduct Lu
teal"), horiz = T, cex = 0.7)

dev.off()
```

```
## quartz_off_screen
##                  2

# Convert Tissue to factor
data2$Tissue <- as.factor(data2$Tissue)

# Create an empty list to store Tukey results
tukey_results_list <- list()

# Unique genes
unique_genes <- unique(data2$Genes)

# Loop over each gene
for (gene in unique_genes) {
  # Subset data for the current gene
  gene_data <- subset(data2, Genes == gene)

  # Fit the generalized linear model
  glm_model <- glm(Protein_Count ~ Tissue, data = gene_data, family = gaussia
n)

  # Perform Tukey test
  contrast_matrix <- glht(glm_model, linfct = mcp(Tissue = "Tukey"))
  tukey_results <- summary(contrast_matrix)

  # Store the results
  tukey_results_list[[gene]] <- tukey_results
}

# Print the results for each gene
for (gene in unique_genes) {
  cat("Tukey results for gene:", gene, "\n")
  print(tukey_results_list[[gene]])
  cat("\n")
}

## Tukey results for gene: ANXA1
##
##   Simultaneous Tests for General Linear Hypotheses
##
## Multiple Comparisons of Means: Tukey Contrasts
##
##
## Fit: glm(formula = Protein_Count ~ Tissue, family = gaussian, data = gene_
data)
##
## Linear Hypotheses:
##              Estimate Std. Error z value Pr(>|z|)
## OL - OF == 0   663529    1012939   0.655    0.512
## (Adjusted p values reported -- single-step method)
```

```
##
##
## Tukey results for gene: ANXA11
##
##   Simultaneous Tests for General Linear Hypotheses
##
## Multiple Comparisons of Means: Tukey Contrasts
##
##
## Fit: glm(formula = Protein_Count ~ Tissue, family = gaussian, data = gene_
data)
##
## Linear Hypotheses:
##              Estimate Std. Error z value Pr(>|z|)
## OL - OF == 0   -10276     215601  -0.048    0.962
## (Adjusted p values reported -- single-step method)
##
##
## Tukey results for gene: ANXA2
##
##   Simultaneous Tests for General Linear Hypotheses
##
## Multiple Comparisons of Means: Tukey Contrasts
##
##
## Fit: glm(formula = Protein_Count ~ Tissue, family = gaussian, data = gene_
data)
##
## Linear Hypotheses:
##              Estimate Std. Error z value Pr(>|z|)
## OL - OF == 0    88382    2324851   0.038     0.97
## (Adjusted p values reported -- single-step method)
##
##
## Tukey results for gene: ANXA3
##
##   Simultaneous Tests for General Linear Hypotheses
##
## Multiple Comparisons of Means: Tukey Contrasts
##
##
## Fit: glm(formula = Protein_Count ~ Tissue, family = gaussian, data = gene_
data)
##
## Linear Hypotheses:
##              Estimate Std. Error z value Pr(>|z|)
## OL - OF == 0    43180     146241   0.295    0.768
## (Adjusted p values reported -- single-step method)
##
##
```

```
## Tukey results for gene: ANXA4
##
##   Simultaneous Tests for General Linear Hypotheses
##
## Multiple Comparisons of Means: Tukey Contrasts
##
##
## Fit: glm(formula = Protein_Count ~ Tissue, family = gaussian, data = gene_
data)
##
## Linear Hypotheses:
##              Estimate Std. Error z value Pr(>|z|)
## OL - OF == 0   206868     214763   0.963    0.335
## (Adjusted p values reported -- single-step method)
##
##
## Tukey results for gene: ANXA5
##
##   Simultaneous Tests for General Linear Hypotheses
##
## Multiple Comparisons of Means: Tukey Contrasts
##
##
## Fit: glm(formula = Protein_Count ~ Tissue, family = gaussian, data = gene_
data)
##
## Linear Hypotheses:
##              Estimate Std. Error z value Pr(>|z|)
## OL - OF == 0 -2290668    5888735  -0.389    0.697
## (Adjusted p values reported -- single-step method)
##
##
## Tukey results for gene: ANXA6
##
##   Simultaneous Tests for General Linear Hypotheses
##
## Multiple Comparisons of Means: Tukey Contrasts
##
##
## Fit: glm(formula = Protein_Count ~ Tissue, family = gaussian, data = gene_
data)
##
## Linear Hypotheses:
##              Estimate Std. Error z value Pr(>|z|)
## OL - OF == 0   287314     602772   0.477    0.634
## (Adjusted p values reported -- single-step method)
##
##
## Tukey results for gene: ANXA7
##
```

```
##   Simultaneous Tests for General Linear Hypotheses
##
## Multiple Comparisons of Means: Tukey Contrasts
##
##
## Fit: glm(formula = Protein_Count ~ Tissue, family = gaussian, data = gene_
data)
##
## Linear Hypotheses:
##               Estimate Std. Error z value Pr(>|z|)
## OL - OF == 0   157028     214061   0.734    0.463
## (Adjusted p values reported -- single-step method)
##
##
## Tukey results for gene: ANXA8
##
##   Simultaneous Tests for General Linear Hypotheses
##
## Multiple Comparisons of Means: Tukey Contrasts
##
##
## Fit: glm(formula = Protein_Count ~ Tissue, family = gaussian, data = gene_
data)
##
## Linear Hypotheses:
##               Estimate Std. Error z value Pr(>|z|)
## OL - OF == 0  -103175     315010  -0.328    0.743
## (Adjusted p values reported -- single-step method)
##
##
## Tukey results for gene: C1QC
##
##   Simultaneous Tests for General Linear Hypotheses
##
## Multiple Comparisons of Means: Tukey Contrasts
##
##
## Fit: glm(formula = Protein_Count ~ Tissue, family = gaussian, data = gene_
data)
##
## Linear Hypotheses:
##               Estimate Std. Error z value Pr(>|z|)
## OL - OF == 0    -5969      66422   -0.09    0.928
## (Adjusted p values reported -- single-step method)
##
##
## Tukey results for gene: C1QTNF3
##
##   Simultaneous Tests for General Linear Hypotheses
##
```

```
## Multiple Comparisons of Means: Tukey Contrasts
##
##
## Fit: glm(formula = Protein_Count ~ Tissue, family = gaussian, data = gene_
data)
##
## Linear Hypotheses:
##               Estimate Std. Error z value Pr(>|z|)
## OL - OF == 0    -1260      30979  -0.041    0.968
## (Adjusted p values reported -- single-step method)
##
##
## Tukey results for gene: C1QTNF5
##
##   Simultaneous Tests for General Linear Hypotheses
##
## Multiple Comparisons of Means: Tukey Contrasts
##
##
## Fit: glm(formula = Protein_Count ~ Tissue, family = gaussian, data = gene_
data)
##
## Linear Hypotheses:
##               Estimate Std. Error z value Pr(>|z|)
## OL - OF == 0     1042      58773   0.018    0.986
## (Adjusted p values reported -- single-step method)
##
##
## Tukey results for gene: C1QTNF7
##
##   Simultaneous Tests for General Linear Hypotheses
##
## Multiple Comparisons of Means: Tukey Contrasts
##
##
## Fit: glm(formula = Protein_Count ~ Tissue, family = gaussian, data = gene_
data)
##
## Linear Hypotheses:
##               Estimate Std. Error z value Pr(>|z|)
## OL - OF == 0   -83536     147127  -0.568     0.57
## (Adjusted p values reported -- single-step method)
##
##
## Tukey results for gene: CLEC11A
##
##   Simultaneous Tests for General Linear Hypotheses
##
## Multiple Comparisons of Means: Tukey Contrasts
##
```

```
##
## Fit: glm(formula = Protein_Count ~ Tissue, family = gaussian, data = gene_
data)
##
## Linear Hypotheses:
##              Estimate Std. Error z value Pr(>|z|)
## OL - OF == 0   -24250      29076  -0.834    0.404
## (Adjusted p values reported -- single-step method)
##
##
## Tukey results for gene: CLEC3B
##
##   Simultaneous Tests for General Linear Hypotheses
##
## Multiple Comparisons of Means: Tukey Contrasts
##
##
## Fit: glm(formula = Protein_Count ~ Tissue, family = gaussian, data = gene_
data)
##
## Linear Hypotheses:
##              Estimate Std. Error z value Pr(>|z|)
## OL - OF == 0  -209062     547669  -0.382    0.703
## (Adjusted p values reported -- single-step method)
##
##
## Tukey results for gene: CSPG4
##
##   Simultaneous Tests for General Linear Hypotheses
##
## Multiple Comparisons of Means: Tukey Contrasts
##
##
## Fit: glm(formula = Protein_Count ~ Tissue, family = gaussian, data = gene_
data)
##
## Linear Hypotheses:
##              Estimate Std. Error z value Pr(>|z|)
## OL - OF == 0     3562      76288   0.047    0.963
## (Adjusted p values reported -- single-step method)
##
##
## Tukey results for gene: HPX
##
##   Simultaneous Tests for General Linear Hypotheses
##
## Multiple Comparisons of Means: Tukey Contrasts
##
##
## Fit: glm(formula = Protein_Count ~ Tissue, family = gaussian, data = gene_
```

```
data)
##
## Linear Hypotheses:
##                Estimate Std. Error z value Pr(>|z|)
## OL - OF == 0   152249     136351   1.117    0.264
## (Adjusted p values reported -- single-step method)
##
##
## Tukey results for gene: LGALS1
##
##   Simultaneous Tests for General Linear Hypotheses
##
## Multiple Comparisons of Means: Tukey Contrasts
##
##
## Fit: glm(formula = Protein_Count ~ Tissue, family = gaussian, data = gene_
data)
##
## Linear Hypotheses:
##                 Estimate Std. Error z value Pr(>|z|)
## OL - OF == 0 -27155048   22997062  -1.181    0.238
## (Adjusted p values reported -- single-step method)
##
##
## Tukey results for gene: LGALS7
##
##   Simultaneous Tests for General Linear Hypotheses
##
## Multiple Comparisons of Means: Tukey Contrasts
##
##
## Fit: glm(formula = Protein_Count ~ Tissue, family = gaussian, data = gene_
data)
##
## Linear Hypotheses:
##                Estimate Std. Error z value Pr(>|z|)
## OL - OF == 0   -45818      63856  -0.718    0.473
## (Adjusted p values reported -- single-step method)
##
##
## Tukey results for gene: LGALS9
##
##   Simultaneous Tests for General Linear Hypotheses
##
## Multiple Comparisons of Means: Tukey Contrasts
##
##
## Fit: glm(formula = Protein_Count ~ Tissue, family = gaussian, data = gene_
data)
##
```

```
## Linear Hypotheses:
##               Estimate Std. Error z value Pr(>|z|)
## OL - OF == 0    47995     356248   0.135    0.893
## (Adjusted p values reported -- single-step method)
##
##
## Tukey results for gene: LMAN1
##
##   Simultaneous Tests for General Linear Hypotheses
##
## Multiple Comparisons of Means: Tukey Contrasts
##
##
## Fit: glm(formula = Protein_Count ~ Tissue, family = gaussian, data = gene_
data)
##
## Linear Hypotheses:
##               Estimate Std. Error z value Pr(>|z|)
## OL - OF == 0   -25270      47101  -0.537    0.592
## (Adjusted p values reported -- single-step method)
##
##
## Tukey results for gene: OVGP1
##
##   Simultaneous Tests for General Linear Hypotheses
##
## Multiple Comparisons of Means: Tukey Contrasts
##
##
## Fit: glm(formula = Protein_Count ~ Tissue, family = gaussian, data = gene_
data)
##
## Linear Hypotheses:
##               Estimate Std. Error z value Pr(>|z|)
## OL - OF == 0 -1488215    3251216  -0.458    0.647
## (Adjusted p values reported -- single-step method)
##
##
## Tukey results for gene: PLXNB1
##
##   Simultaneous Tests for General Linear Hypotheses
##
## Multiple Comparisons of Means: Tukey Contrasts
##
##
## Fit: glm(formula = Protein_Count ~ Tissue, family = gaussian, data = gene_
data)
##
## Linear Hypotheses:
##               Estimate Std. Error z value Pr(>|z|)
```

```
## OL - OF == 0    29872     148738   0.201    0.841
## (Adjusted p values reported -- single-step method)
##
##
## Tukey results for gene: PLXNB2
##
##   Simultaneous Tests for General Linear Hypotheses
##
## Multiple Comparisons of Means: Tukey Contrasts
##
##
## Fit: glm(formula = Protein_Count ~ Tissue, family = gaussian, data = gene_
data)
##
## Linear Hypotheses:
##                Estimate Std. Error z value Pr(>|z|)
## OL - OF == 0    16956     122624   0.138     0.89
## (Adjusted p values reported -- single-step method)
```

```
#### Secreted Factors

# Read your cleaned joined data file into a data frame
cleaned_joined_data <- read.csv("cleaned_matrisome_oviduct.csv")

# Filter rows with Matrisome_Category "Secreted Factors"
Secreted_Factors_data <- cleaned_joined_data %>%
  filter(Matrisome_Category == "Secreted Factors")

# Reshape the data to have separate columns for each replicate and each tissu
e stage
organized_data <- Secreted_Factors_data %>%
  pivot_longer(cols = starts_with("OF") | starts_with("OL"),
               names_to = c("Tissue", "Stage"),
               names_sep = "\\.",
               values_to = "Protein_Count")
```

```
## Warning: Expected 2 pieces. Missing pieces filled with `NA` in 2 rows [1,
5].
```

```
# Save the reshaped data to a new file

write.csv(organized_data, "Secreted_factors_data_matrisome_oviduct_to_clean.c
sv", row.names = FALSE)

data2 <- read.csv("Secreted_factors_data_matrisome_oviduct_clean.csv")

png(filename="ovi_Secreted_factors.png",
    width = 12, height = 6, units = "in",
    res = 600)
```

```
boxplot(Protein_Count ~ Tissue*Genes,
        data = data2,
        cex.axis = 0.8,
        col = viridis(2, alpha = 0.7),
        lwd = 0.7,
        ylab = "",
        ylim = c(0, 6500000),
        axes = FALSE,
        frame.plot = TRUE,
        xlab = "")

GROUP <- unique(data2$Genes)
groups <- 23
numbox <- 2
total <- groups * numbox
xpoints <- seq(median(1:numbox), total, numbox)

axis(2, seq(0, 6500000, by = 1000000), las = 1, cex.axis = 0.7)
axis(1, labels = GROUP, at = xpoints, las = 2, cex.axis = 0.7)

stripchart(data2$Protein_Count ~ data2$Tissue*data2$Genes,
           method = "jitter",
           pch = 19,
           col = "black",
           vertical = TRUE,
           add = TRUE,
           cex = 0.3)

title(ylab = expression("Protein abundance (RU)"), line = 3.0)
legend("top", fill = viridis(2), legend = c("Oviduct Follicular", "Oviduct Lu
teal"), horiz = T, cex = 0.7)

dev.off()

## quartz_off_screen
##                 2

# Convert Tissue to factor
data2$Tissue <- as.factor(data2$Tissue)

# Create an empty list to store Tukey results
tukey_results_list <- list()

# Unique genes
unique_genes <- unique(data2$Genes)

# Loop over each gene
for (gene in unique_genes) {
  # Subset data for the current gene
  gene_data <- subset(data2, Genes == gene)
```

```
  # Fit the generalized linear model
  glm_model <- glm(Protein_Count ~ Tissue, data = gene_data, family = gaussia
n)

  # Perform Tukey test
  contrast_matrix <- glht(glm_model, linfct = mcp(Tissue = "Tukey"))
  tukey_results <- summary(contrast_matrix)

  # Store the results
  tukey_results_list[[gene]] <- tukey_results
}

# Print the results for each gene
for (gene in unique_genes) {
  cat("Tukey results for gene:", gene, "\n")
  print(tukey_results_list[[gene]])
  cat("\n")
}

## Tukey results for gene: ANGPT4
##
##   Simultaneous Tests for General Linear Hypotheses
##
## Multiple Comparisons of Means: Tukey Contrasts
##
##
## Fit: glm(formula = Protein_Count ~ Tissue, family = gaussian, data = gene_
data)
##
## Linear Hypotheses:
##              Estimate Std. Error z value Pr(>|z|)
## OL - OF == 0   -31576      88250  -0.358     0.72
## (Adjusted p values reported -- single-step method)
##
##
## Tukey results for gene: ANGPTL2
##
##   Simultaneous Tests for General Linear Hypotheses
##
## Multiple Comparisons of Means: Tukey Contrasts
##
##
## Fit: glm(formula = Protein_Count ~ Tissue, family = gaussian, data = gene_
data)
##
## Linear Hypotheses:
##              Estimate Std. Error z value Pr(>|z|)
## OL - OF == 0   -56327     171343  -0.329    0.742
## (Adjusted p values reported -- single-step method)
```

```
##
##
## Tukey results for gene: CHRD
##
##   Simultaneous Tests for General Linear Hypotheses
##
## Multiple Comparisons of Means: Tukey Contrasts
##
##
## Fit: glm(formula = Protein_Count ~ Tissue, family = gaussian, data = gene_
data)
##
## Linear Hypotheses:
##              Estimate Std. Error z value Pr(>|z|)
## OL - OF == 0   -22045     162265  -0.136    0.892
## (Adjusted p values reported -- single-step method)
##
##
## Tukey results for gene: HCFC1
##
##   Simultaneous Tests for General Linear Hypotheses
##
## Multiple Comparisons of Means: Tukey Contrasts
##
##
## Fit: glm(formula = Protein_Count ~ Tissue, family = gaussian, data = gene_
data)
##
## Linear Hypotheses:
##              Estimate Std. Error z value Pr(>|z|)
## OL - OF == 0    26036     103069   0.253    0.801
## (Adjusted p values reported -- single-step method)
##
##
## Tukey results for gene: HGFAC
##
##   Simultaneous Tests for General Linear Hypotheses
##
## Multiple Comparisons of Means: Tukey Contrasts
##
##
## Fit: glm(formula = Protein_Count ~ Tissue, family = gaussian, data = gene_
data)
##
## Linear Hypotheses:
##              Estimate Std. Error z value Pr(>|z|)
## OL - OF == 0   -30100      72451  -0.415    0.678
## (Adjusted p values reported -- single-step method)
##
##
```

```
## Tukey results for gene: IGF2
##
##   Simultaneous Tests for General Linear Hypotheses
##
## Multiple Comparisons of Means: Tukey Contrasts
##
##
## Fit: glm(formula = Protein_Count ~ Tissue, family = gaussian, data = gene_
data)
##
## Linear Hypotheses:
##              Estimate Std. Error z value Pr(>|z|)
## OL - OF == 0   -54710      72462  -0.755     0.45
## (Adjusted p values reported -- single-step method)
##
##
## Tukey results for gene: IL17D
##
##   Simultaneous Tests for General Linear Hypotheses
##
## Multiple Comparisons of Means: Tukey Contrasts
##
##
## Fit: glm(formula = Protein_Count ~ Tissue, family = gaussian, data = gene_
data)
##
## Linear Hypotheses:
##              Estimate Std. Error z value Pr(>|z|)
## OL - OF == 0    91913     160976   0.571    0.568
## (Adjusted p values reported -- single-step method)
##
##
## Tukey results for gene: INHBB
##
##   Simultaneous Tests for General Linear Hypotheses
##
## Multiple Comparisons of Means: Tukey Contrasts
##
##
## Fit: glm(formula = Protein_Count ~ Tissue, family = gaussian, data = gene_
data)
##
## Linear Hypotheses:
##              Estimate Std. Error z value Pr(>|z|)
## OL - OF == 0   -14910     128449  -0.116    0.908
## (Adjusted p values reported -- single-step method)
##
##
## Tukey results for gene: INHBE
##
```

```
##   Simultaneous Tests for General Linear Hypotheses
##
## Multiple Comparisons of Means: Tukey Contrasts
##
##
## Fit: glm(formula = Protein_Count ~ Tissue, family = gaussian, data = gene_
data)
##
## Linear Hypotheses:
##              Estimate Std. Error z value Pr(>|z|)
## OL - OF == 0    41431     116116   0.357    0.721
## (Adjusted p values reported -- single-step method)
##
##
## Tukey results for gene: S100A10
##
##   Simultaneous Tests for General Linear Hypotheses
##
## Multiple Comparisons of Means: Tukey Contrasts
##
##
## Fit: glm(formula = Protein_Count ~ Tissue, family = gaussian, data = gene_
data)
##
## Linear Hypotheses:
##              Estimate Std. Error z value Pr(>|z|)
## OL - OF == 0   214341     308998   0.694    0.488
## (Adjusted p values reported -- single-step method)
##
##
## Tukey results for gene: S100A11
##
##   Simultaneous Tests for General Linear Hypotheses
##
## Multiple Comparisons of Means: Tukey Contrasts
##
##
## Fit: glm(formula = Protein_Count ~ Tissue, family = gaussian, data = gene_
data)
##
## Linear Hypotheses:
##              Estimate Std. Error z value Pr(>|z|)
## OL - OF == 0    -7214     673596  -0.011    0.991
## (Adjusted p values reported -- single-step method)
##
##
## Tukey results for gene: S100A12
##
##   Simultaneous Tests for General Linear Hypotheses
##
```

```
## Multiple Comparisons of Means: Tukey Contrasts
##
##
## Fit: glm(formula = Protein_Count ~ Tissue, family = gaussian, data = gene_
data)
##
## Linear Hypotheses:
##              Estimate Std. Error z value Pr(>|z|)
## OL - OF == 0   -197.8    32891.4  -0.006    0.995
## (Adjusted p values reported -- single-step method)
##
##
## Tukey results for gene: S100A13
##
##   Simultaneous Tests for General Linear Hypotheses
##
## Multiple Comparisons of Means: Tukey Contrasts
##
##
## Fit: glm(formula = Protein_Count ~ Tissue, family = gaussian, data = gene_
data)
##
## Linear Hypotheses:
##              Estimate Std. Error z value Pr(>|z|)
## OL - OF == 0   202256     430514    0.47    0.638
## (Adjusted p values reported -- single-step method)
##
##
## Tukey results for gene: S100A14
##
##   Simultaneous Tests for General Linear Hypotheses
##
## Multiple Comparisons of Means: Tukey Contrasts
##
##
## Fit: glm(formula = Protein_Count ~ Tissue, family = gaussian, data = gene_
data)
##
## Linear Hypotheses:
##              Estimate Std. Error z value Pr(>|z|)
## OL - OF == 0    57819     131102   0.441    0.659
## (Adjusted p values reported -- single-step method)
##
##
## Tukey results for gene: S100A16
##
##   Simultaneous Tests for General Linear Hypotheses
##
## Multiple Comparisons of Means: Tukey Contrasts
##
```

```
##
## Fit: glm(formula = Protein_Count ~ Tissue, family = gaussian, data = gene_
data)
##
## Linear Hypotheses:
##              Estimate Std. Error z value Pr(>|z|)
## OL - OF == 0    -8070      96816  -0.083    0.934
## (Adjusted p values reported -- single-step method)
##
##
## Tukey results for gene: S100A4
##
##   Simultaneous Tests for General Linear Hypotheses
##
## Multiple Comparisons of Means: Tukey Contrasts
##
##
## Fit: glm(formula = Protein_Count ~ Tissue, family = gaussian, data = gene_
data)
##
## Linear Hypotheses:
##              Estimate Std. Error z value Pr(>|z|)
## OL - OF == 0  3927542    4976030   0.789     0.43
## (Adjusted p values reported -- single-step method)
##
##
## Tukey results for gene: S100A8
##
##   Simultaneous Tests for General Linear Hypotheses
##
## Multiple Comparisons of Means: Tukey Contrasts
##
##
## Fit: glm(formula = Protein_Count ~ Tissue, family = gaussian, data = gene_
data)
##
## Linear Hypotheses:
##              Estimate Std. Error z value Pr(>|z|)
## OL - OF == 0    50014      33751   1.482    0.138
## (Adjusted p values reported -- single-step method)
##
##
## Tukey results for gene: SFRP1
##
##   Simultaneous Tests for General Linear Hypotheses
##
## Multiple Comparisons of Means: Tukey Contrasts
##
##
## Fit: glm(formula = Protein_Count ~ Tissue, family = gaussian, data = gene_
```

```
data)
##
## Linear Hypotheses:
##              Estimate Std. Error z value Pr(>|z|)
## OL - OF == 0    20726      44451   0.466    0.641
## (Adjusted p values reported -- single-step method)
##
##
## Tukey results for gene: SFRP2
##
##   Simultaneous Tests for General Linear Hypotheses
##
## Multiple Comparisons of Means: Tukey Contrasts
##
##
## Fit: glm(formula = Protein_Count ~ Tissue, family = gaussian, data = gene_
data)
##
## Linear Hypotheses:
##              Estimate Std. Error z value Pr(>|z|)
## OL - OF == 0   165011     267270   0.617    0.537
## (Adjusted p values reported -- single-step method)
##
##
## Tukey results for gene: TGFB2
##
##   Simultaneous Tests for General Linear Hypotheses
##
## Multiple Comparisons of Means: Tukey Contrasts
##
##
## Fit: glm(formula = Protein_Count ~ Tissue, family = gaussian, data = gene_
data)
##
## Linear Hypotheses:
##              Estimate Std. Error z value Pr(>|z|)
## OL - OF == 0   -34642      56628  -0.612    0.541
## (Adjusted p values reported -- single-step method)
##
##
## Tukey results for gene: WNT2B
##
##   Simultaneous Tests for General Linear Hypotheses
##
## Multiple Comparisons of Means: Tukey Contrasts
##
##
## Fit: glm(formula = Protein_Count ~ Tissue, family = gaussian, data = gene_
data)
##
```

```
## Linear Hypotheses:
##                Estimate Std. Error z value Pr(>|z|)
## OL - OF == 0   -75463     184780  -0.408    0.683
## (Adjusted p values reported -- single-step method)
##
##
## Tukey results for gene: WNT5A
##
##   Simultaneous Tests for General Linear Hypotheses
##
## Multiple Comparisons of Means: Tukey Contrasts
##
##
## Fit: glm(formula = Protein_Count ~ Tissue, family = gaussian, data = gene_
data)
##
## Linear Hypotheses:
##                Estimate Std. Error z value Pr(>|z|)
## OL - OF == 0    -1504     121744  -0.012     0.99
## (Adjusted p values reported -- single-step method)
##
##
## Tukey results for gene: WNT6
##
##   Simultaneous Tests for General Linear Hypotheses
##
## Multiple Comparisons of Means: Tukey Contrasts
##
##
## Fit: glm(formula = Protein_Count ~ Tissue, family = gaussian, data = gene_
data)
##
## Linear Hypotheses:
##                Estimate Std. Error z value Pr(>|z|)
## OL - OF == 0   -38991     137640  -0.283    0.777
## (Adjusted p values reported -- single-step method)

####Components proportions

data <- read.csv("Oviduct_matrisome_final.csv")

# Subset the data for each tissue (OF and OL) separately
data_OF <- subset(data, Tissue == "OF")
data_OL <- subset(data, Tissue == "OL")

# Function to calculate proportions
calculate_proportions <- function(data_subset) {
  proportions <- data_subset %>%
    group_by(Matrisome_Category) %>%
    summarize(Proportion = sum(Protein_Count) / sum(data_subset$Protein_Count
```

```
) * 100)
  return(proportions)
}

# Calculate proportions for OF
proportions_OF <- calculate_proportions(data_OF)

# Calculate proportions for OL
proportions_OL <- calculate_proportions(data_OL)

# Print proportions for OF
cat("Proportions for Matrisome categories in tissue OF:\n")

## Proportions for Matrisome categories in tissue OF:

print(proportions_OF)

## # A tibble: 6 × 2
##   Matrisome_Category Proportion
##   <chr>                   <dbl>
## 1 Collagens              53.6
## 2 ECM Glycoproteins      27.9
## 3 ECM Regulators          7.17
## 4 ECM-affiliated          2.33
## 5 Proteoglycans           8.11
## 6 Secreted Factors        0.882

# Print proportions for OL
cat("Proportions for Matrisome categories in tissue OL:\n")

## Proportions for Matrisome categories in tissue OL:

print(proportions_OL)

## # A tibble: 6 × 2
##   Matrisome_Category Proportion
##   <chr>                   <dbl>
## 1 Collagens              56.1
## 2 ECM Glycoproteins      27.3
## 3 ECM Regulators          6.62
## 4 ECM-affiliated          1.52
## 5 Proteoglycans           7.50
## 6 Secreted Factors        0.940

###Pie charts

# Define a function to create a pie chart with values and save it
create_and_save_pie_chart <- function(proportions, title, filename) {

  # Create a color palette using Viridis
  color_palette <- viridis(length(unique(proportions$Matrisome_Category)))
```

```
  # Create a pie chart
  pie_chart <- ggplot(proportions, aes(x = "", y = Proportion, fill = Matriso
me_Category)) +
    geom_bar(stat = "identity") +
    coord_polar("y", start = 1) +
    geom_text(aes(label = sprintf("%.1f%%", Proportion)), position = position
_stack(vjust = 0.5), color = "white", size = 6) +
    labs(title = title, x = NULL, y = NULL, fill = "Matrisome Category") +
    scale_fill_manual(values = color_palette) +  # Set the color palette
    theme_void() +
    theme(legend.position = "right")

  # Save the pie chart
  ggsave(filename, plot = pie_chart, width = 6, height = 6, units = "in", dpi
= 300)
}

# Create and save a pie chart for OF
create_and_save_pie_chart(proportions_OF, "Matrisome Categories Proportion in
Tissue OF", "pie_chart_OF.png")

# Create and save a pie chart for OL
create_and_save_pie_chart(proportions_OL, "Matrisome Categories Proportion in
Tissue OL", "pie_chart_OL.png")

#####Summary data

data <- read.csv("oviduct_matrisome_final.csv")

# Calculate mean and standard deviation for each Gene considering Tissue and
Matrisome_Category
gene_stats <- data %>%
  group_by(Genes, Tissue, Matrisome_Category) %>%
  summarize(
    Mean_Protein_Count = mean(Protein_Count),
    SD_Protein_Count = sd(Protein_Count)
  )

## `summarise()` has grouped output by 'Genes', 'Tissue'. You can override us
ing
## the `.groups` argument.

# Print the results
print(gene_stats)

## # A tibble: 464 × 5
## # Groups:   Genes, Tissue [464]
##    Genes    Tissue Matrisome_Category Mean_Protein_Count SD_Protein_Count
##    <chr>    <chr>  <chr>                           <dbl>            <dbl>
```

```
## 1 A2M OF ECM Regulators 5191380 1300155.
## 2 A2M OL ECM Regulators 4876600 1065803.
## 3 ABI3BP OF ECM Glycoproteins 1149890. 541863.
## 4 ABI3BP OL ECM Glycoproteins 882364 361874.
## 5 ACAN OF Proteoglycans 396077. 370773.
## 6 ACAN OL Proteoglycans 324861. 279260.
## 7 ADAMTS5 OF ECM Regulators 306157. 95016.
## 8 ADAMTS5 OL ECM Regulators 312698. 176269.
## 9 ADAMTSL3 OF ECM Regulators 252108 134813.
## 10 ADAMTSL3 OL ECM Regulators 202840. 157669.
## # ℹ 454 more rows

# Save the summarized data to a CSV file
write.csv(gene_stats, file = "COviduct_Summary.csv", row.names = FALSE)



####ENDOMETRIUM

# Read your data into a data frame or data table
data <- read.csv("Endo_all_count.csv")

# Remove rows with more than 5 NA values
clean_data <- data %>%
  filter(rowSums(is.na(.)) <= 5)

# Replace remaining NA values with 0 in all columns
clean_data <- clean_data %>%
  mutate_all(funs(replace(., is.na(.), 0)))

## Warning: `funs()` was deprecated in dplyr 0.8.0.
## ℹ Please use a list of either functions or lambdas:
##
## # Simple named list: list(mean = mean, median = median)
##
## # Auto named with `tibble::lst()`: tibble::lst(mean, median)
##
## # Using lambdas list(~ mean(., trim = .2), ~ median(., na.rm = TRUE))
## Call `lifecycle::last_lifecycle_warnings()` to see where this warning was
## generated.

# Save the cleaned data to a new file
write.csv(clean_data, "cleaned_data_endo.csv", row.names = FALSE)

### Matrisome categorization

# Read your first data file into a data frame or data table
data <- read.csv("cleaned_data_endo.csv")
```

```
# Read your second dataset into another data frame or data table
matrisome_info <- read.csv("Bt_Matrisome.csv")

# Perform a left join based on the "Genes" column
joined_data <- left_join(data, matrisome_info, by = "Genes")

# Save the joined data to a new file
write.csv(joined_data, "endo_matrisome.csv", row.names = FALSE)

# Remove rows where either Matrisome_Division or Matrisome_Category have NA v
alues
cleaned_joined_data <- joined_data %>%
  filter(!is.na(Matrisome_Division) & !is.na(Matrisome_Category))

# Save the cleaned data to a new file

write.csv(cleaned_joined_data, "cleaned_matrisome_endo.csv", row.names = FALS
E)

#######Ploting


#### Collagens
# Read your cleaned joined data file into a data frame
cleaned_joined_data <- read.csv("cleaned_matrisome_endo.csv")

# Filter rows with Matrisome_Category "Collagens"
collagen_data <- cleaned_joined_data %>%
  filter(Matrisome_Category == "Collagens")

# Reshape the data to have separate columns for each replicate and each tissu
e stage
organized_data <- collagen_data %>%
  pivot_longer(cols = starts_with("EF") | starts_with("EL"),
               names_to = c("Tissue", "Stage"),
               names_sep = "\\.",
               values_to = "Protein_Count")

## Warning: Expected 2 pieces. Missing pieces filled with `NA` in 2 rows [1,
6].

# Save the reshaped data to a new file

write.csv(organized_data, "organized_matrisome_endo_to_clean.csv", row.names
= FALSE)

data2 <- read.csv("Collagens_data_matrisome_endo_clean.csv")

png(filename="endo_collagens.png",
```

```r
    width = 12, height = 6, units = "in",
    res = 600)

boxplot(Protein_Count ~ Tissue*Genes,
        data = data2,
        cex.axis = 0.8,
        col = viridis(2, alpha = 0.7),
        lwd = 0.7,
        ylab = "",
        ylim = c(0, 8000000000),
        axes = FALSE,
        frame.plot = TRUE,
        xlab = "")

GROUP <- unique(data2$Genes)
groups <- 24
numbox <- 2
total <- groups * numbox
xpoints <- seq(median(1:numbox), total, numbox)

axis(2, seq(0, 8000000000, by = 1000000000), las = 1, cex.axis = 0.7)
axis(1, labels = GROUP, at = xpoints, las = 2, cex.axis = 0.7)

stripchart(data2$Protein_Count ~ data2$Tissue*data2$Genes,
           method = "jitter",
           pch = 19,
           col = "black",
           vertical = TRUE,
           add = TRUE,
           cex = 0.3)

title(ylab = expression("Protein abundance (RU)"), line = 3.0)
legend("top", fill = viridis(2), legend = c("Endometrium Follicular", "Endome
trium Luteal"), horiz = T, cex = 0.7)

dev.off()

## quartz_off_screen
##                  2

# Convert Tissue to factor
data2$Tissue <- as.factor(data2$Tissue)

# Create an empty list to store Tukey results
tukey_results_list <- list()

# Unique genes
unique_genes <- unique(data2$Genes)
```

```
# Loop over each gene
for (gene in unique_genes) {
  # Subset data for the current gene
  gene_data <- subset(data2, Genes == gene)

  # Fit the generalized linear model
  glm_model <- glm(Protein_Count ~ Tissue, data = gene_data, family = gaussia
n)

  # Perform Tukey test
  contrast_matrix <- glht(glm_model, linfct = mcp(Tissue = "Tukey"))
  tukey_results <- summary(contrast_matrix)

  # Store the results
  tukey_results_list[[gene]] <- tukey_results
}

# Print the results for each gene
for (gene in unique_genes) {
  cat("Tukey results for gene:", gene, "\n")
  print(tukey_results_list[[gene]])
  cat("\n")
}

## Tukey results for gene: COL11A1
##
##   Simultaneous Tests for General Linear Hypotheses
##
## Multiple Comparisons of Means: Tukey Contrasts
##
##
## Fit: glm(formula = Protein_Count ~ Tissue, family = gaussian, data = gene_
data)
##
## Linear Hypotheses:
##                Estimate Std. Error z value Pr(>|z|)
## EL - EF == 0  -234552     250526  -0.936    0.349
## (Adjusted p values reported -- single-step method)
##
##
## Tukey results for gene: COL12A1
##
##   Simultaneous Tests for General Linear Hypotheses
##
## Multiple Comparisons of Means: Tukey Contrasts
##
##
## Fit: glm(formula = Protein_Count ~ Tissue, family = gaussian, data = gene_
data)
##
```

```
## Linear Hypotheses:
##               Estimate Std. Error z value Pr(>|z|)
## EL - EF == 0 -1209316    2111741  -0.573    0.567
## (Adjusted p values reported -- single-step method)
##
##
## Tukey results for gene: COL14A1
##
##   Simultaneous Tests for General Linear Hypotheses
##
## Multiple Comparisons of Means: Tukey Contrasts
##
##
## Fit: glm(formula = Protein_Count ~ Tissue, family = gaussian, data = gene_
data)
##
## Linear Hypotheses:
##               Estimate Std. Error z value Pr(>|z|)
## EL - EF == 0  -849076    3129545  -0.271    0.786
## (Adjusted p values reported -- single-step method)
##
##
## Tukey results for gene: COL15A1
##
##   Simultaneous Tests for General Linear Hypotheses
##
## Multiple Comparisons of Means: Tukey Contrasts
##
##
## Fit: glm(formula = Protein_Count ~ Tissue, family = gaussian, data = gene_
data)
##
## Linear Hypotheses:
##               Estimate Std. Error z value Pr(>|z|)
## EL - EF == 0 -3503918    2123972   -1.65    0.099 .
## ---
## Signif. codes:  0 '***' 0.001 '**' 0.01 '*' 0.05 '.' 0.1 ' ' 1
## (Adjusted p values reported -- single-step method)
##
##
## Tukey results for gene: COL16A1
##
##   Simultaneous Tests for General Linear Hypotheses
##
## Multiple Comparisons of Means: Tukey Contrasts
##
##
## Fit: glm(formula = Protein_Count ~ Tissue, family = gaussian, data = gene_
data)
##
```

```
## Linear Hypotheses:
##              Estimate Std. Error z value Pr(>|z|)
## EL - EF == 0   -27647    1226635  -0.023    0.982
## (Adjusted p values reported -- single-step method)
##
##
## Tukey results for gene: COL18A1
##
##   Simultaneous Tests for General Linear Hypotheses
##
## Multiple Comparisons of Means: Tukey Contrasts
##
##
## Fit: glm(formula = Protein_Count ~ Tissue, family = gaussian, data = gene_
data)
##
## Linear Hypotheses:
##              Estimate Std. Error z value Pr(>|z|)
## EL - EF == 0   149108    5342956   0.028    0.978
## (Adjusted p values reported -- single-step method)
##
##
## Tukey results for gene: COL1A1
##
##   Simultaneous Tests for General Linear Hypotheses
##
## Multiple Comparisons of Means: Tukey Contrasts
##
##
## Fit: glm(formula = Protein_Count ~ Tissue, family = gaussian, data = gene_
data)
##
## Linear Hypotheses:
##                Estimate Std. Error z value Pr(>|z|)
## EL - EF == 0 -1.309e+09  1.289e+09  -1.016     0.31
## (Adjusted p values reported -- single-step method)
##
##
## Tukey results for gene: COL1A2
##
##   Simultaneous Tests for General Linear Hypotheses
##
## Multiple Comparisons of Means: Tukey Contrasts
##
##
## Fit: glm(formula = Protein_Count ~ Tissue, family = gaussian, data = gene_
data)
##
## Linear Hypotheses:
##                Estimate Std. Error z value Pr(>|z|)
```

```
## EL - EF == 0 -333475000  409159287  -0.815    0.415
## (Adjusted p values reported -- single-step method)
##
##
## Tukey results for gene: COL26A1
##
##   Simultaneous Tests for General Linear Hypotheses
##
## Multiple Comparisons of Means: Tukey Contrasts
##
##
## Fit: glm(formula = Protein_Count ~ Tissue, family = gaussian, data = gene_
data)
##
## Linear Hypotheses:
##              Estimate Std. Error z value Pr(>|z|)
## EL - EF == 0    21819      89421   0.244    0.807
## (Adjusted p values reported -- single-step method)
##
##
## Tukey results for gene: COL27A1
##
##   Simultaneous Tests for General Linear Hypotheses
##
## Multiple Comparisons of Means: Tukey Contrasts
##
##
## Fit: glm(formula = Protein_Count ~ Tissue, family = gaussian, data = gene_
data)
##
## Linear Hypotheses:
##              Estimate Std. Error z value Pr(>|z|)
## EL - EF == 0   -25679      43316  -0.593    0.553
## (Adjusted p values reported -- single-step method)
##
##
## Tukey results for gene: COL2A1
##
##   Simultaneous Tests for General Linear Hypotheses
##
## Multiple Comparisons of Means: Tukey Contrasts
##
##
## Fit: glm(formula = Protein_Count ~ Tissue, family = gaussian, data = gene_
data)
##
## Linear Hypotheses:
##              Estimate Std. Error z value Pr(>|z|)
## EL - EF == 0   490096     248878   1.969   0.0489 *
## ---
```

```
## Signif. codes:  0 '***' 0.001 '**' 0.01 '*' 0.05 '.' 0.1 ' ' 1
## (Adjusted p values reported -- single-step method)
##
##
## Tukey results for gene: COL3A1
##
##   Simultaneous Tests for General Linear Hypotheses
##
## Multiple Comparisons of Means: Tukey Contrasts
##
##
## Fit: glm(formula = Protein_Count ~ Tissue, family = gaussian, data = gene_
data)
##
## Linear Hypotheses:
##               Estimate Std. Error z value Pr(>|z|)
## EL - EF == 0 -10508680   11428983  -0.919    0.358
## (Adjusted p values reported -- single-step method)
##
##
## Tukey results for gene: COL4A1
##
##   Simultaneous Tests for General Linear Hypotheses
##
## Multiple Comparisons of Means: Tukey Contrasts
##
##
## Fit: glm(formula = Protein_Count ~ Tissue, family = gaussian, data = gene_
data)
##
## Linear Hypotheses:
##              Estimate Std. Error z value Pr(>|z|)
## EL - EF == 0 -5711040    2910498  -1.962   0.0497 *
## ---
## Signif. codes:  0 '***' 0.001 '**' 0.01 '*' 0.05 '.' 0.1 ' ' 1
## (Adjusted p values reported -- single-step method)
##
##
## Tukey results for gene: COL4A2
##
##   Simultaneous Tests for General Linear Hypotheses
##
## Multiple Comparisons of Means: Tukey Contrasts
##
##
## Fit: glm(formula = Protein_Count ~ Tissue, family = gaussian, data = gene_
data)
##
## Linear Hypotheses:
##              Estimate Std. Error z value Pr(>|z|)
```

```
## EL - EF == 0 -4767668    2544666  -1.874    0.061 .
## ---
## Signif. codes:  0 '***' 0.001 '**' 0.01 '*' 0.05 '.' 0.1 ' ' 1
## (Adjusted p values reported -- single-step method)
##
##
## Tukey results for gene: COL4A3
##
##   Simultaneous Tests for General Linear Hypotheses
##
## Multiple Comparisons of Means: Tukey Contrasts
##
##
## Fit: glm(formula = Protein_Count ~ Tissue, family = gaussian, data = gene_
data)
##
## Linear Hypotheses:
##              Estimate Std. Error z value Pr(>|z|)
## EL - EF == 0 -1257224    1793422  -0.701    0.483
## (Adjusted p values reported -- single-step method)
##
##
## Tukey results for gene: COL4A5
##
##   Simultaneous Tests for General Linear Hypotheses
##
## Multiple Comparisons of Means: Tukey Contrasts
##
##
## Fit: glm(formula = Protein_Count ~ Tissue, family = gaussian, data = gene_
data)
##
## Linear Hypotheses:
##              Estimate Std. Error z value Pr(>|z|)
## EL - EF == 0  -452053     646763  -0.699    0.485
## (Adjusted p values reported -- single-step method)
##
##
## Tukey results for gene: COL4A6
##
##   Simultaneous Tests for General Linear Hypotheses
##
## Multiple Comparisons of Means: Tukey Contrasts
##
##
## Fit: glm(formula = Protein_Count ~ Tissue, family = gaussian, data = gene_
data)
##
## Linear Hypotheses:
##              Estimate Std. Error z value Pr(>|z|)
```

```
## EL - EF == 0  -839823     645412  -1.301    0.193
## (Adjusted p values reported -- single-step method)
##
##
## Tukey results for gene: COL5A1
##
##   Simultaneous Tests for General Linear Hypotheses
##
## Multiple Comparisons of Means: Tukey Contrasts
##
##
## Fit: glm(formula = Protein_Count ~ Tissue, family = gaussian, data = gene_
data)
##
## Linear Hypotheses:
##              Estimate Std. Error z value Pr(>|z|)
## EL - EF == 0 -1543668    2533499  -0.609    0.542
## (Adjusted p values reported -- single-step method)
##
##
## Tukey results for gene: COL5A2
##
##   Simultaneous Tests for General Linear Hypotheses
##
## Multiple Comparisons of Means: Tukey Contrasts
##
##
## Fit: glm(formula = Protein_Count ~ Tissue, family = gaussian, data = gene_
data)
##
## Linear Hypotheses:
##              Estimate Std. Error z value Pr(>|z|)
## EL - EF == 0 -6069542    7721394  -0.786    0.432
## (Adjusted p values reported -- single-step method)
##
##
## Tukey results for gene: COL5A3
##
##   Simultaneous Tests for General Linear Hypotheses
##
## Multiple Comparisons of Means: Tukey Contrasts
##
##
## Fit: glm(formula = Protein_Count ~ Tissue, family = gaussian, data = gene_
data)
##
## Linear Hypotheses:
##              Estimate Std. Error z value Pr(>|z|)
## EL - EF == 0 -1535246    2092778  -0.734    0.463
## (Adjusted p values reported -- single-step method)
```

```
##
##
## Tukey results for gene: COL6A1
##
##   Simultaneous Tests for General Linear Hypotheses
##
## Multiple Comparisons of Means: Tukey Contrasts
##
##
## Fit: glm(formula = Protein_Count ~ Tissue, family = gaussian, data = gene_
data)
##
## Linear Hypotheses:
##                 Estimate Std. Error z value Pr(>|z|)
## EL - EF == 0 -15371180   14337971  -1.072    0.284
## (Adjusted p values reported -- single-step method)
##
##
## Tukey results for gene: COL6A2
##
##   Simultaneous Tests for General Linear Hypotheses
##
## Multiple Comparisons of Means: Tukey Contrasts
##
##
## Fit: glm(formula = Protein_Count ~ Tissue, family = gaussian, data = gene_
data)
##
## Linear Hypotheses:
##                Estimate Std. Error z value Pr(>|z|)
## EL - EF == 0 -8879574    7449737  -1.192    0.233
## (Adjusted p values reported -- single-step method)
##
##
## Tukey results for gene: COL6A3
##
##   Simultaneous Tests for General Linear Hypotheses
##
## Multiple Comparisons of Means: Tukey Contrasts
##
##
## Fit: glm(formula = Protein_Count ~ Tissue, family = gaussian, data = gene_
data)
##
## Linear Hypotheses:
##                 Estimate Std. Error z value Pr(>|z|)
## EL - EF == 0 -22023085   29387460  -0.749    0.454
## (Adjusted p values reported -- single-step method)
##
##
```

```
## Tukey results for gene: COL6A5
##
##   Simultaneous Tests for General Linear Hypotheses
##
## Multiple Comparisons of Means: Tukey Contrasts
##
##
## Fit: glm(formula = Protein_Count ~ Tissue, family = gaussian, data = gene_
data)
##
## Linear Hypotheses:
##                Estimate Std. Error z value Pr(>|z|)
## EL - EF == 0      3386      39290   0.086    0.931
## (Adjusted p values reported -- single-step method)
```

```
#### ECM Glycoproteins
# Read your cleaned joined data file into a data frame
cleaned_joined_data <- read.csv("cleaned_matrisome_endo.csv")

# Filter rows with Matrisome_Category "ECM Glycoproteins"
Glycoproteins_data <- cleaned_joined_data %>%
  filter(Matrisome_Category == "ECM Glycoproteins")

# Reshape the data to have separate columns for each replicate and each tissu
e stage
organized_data <- Glycoproteins_data %>%
  pivot_longer(cols = starts_with("EF") | starts_with("EL"),
               names_to = c("Tissue", "Stage"),
               names_sep = "\\.",
               values_to = "Protein_Count")
```

```
## Warning: Expected 2 pieces. Missing pieces filled with `NA` in 2 rows [1,
6].
```

```
# Save the reshaped data to a new file

write.csv(organized_data, "Glycoproteins_data_matrisome_endo_to_clean.csv", r
ow.names = FALSE)

data2 <- read.csv("Glycoproteins_data_matrisome_endo_clean.csv")

png(filename="endo_ECM Glycoproteins.png",
    width = 12, height = 6, units = "in",
    res = 600)

boxplot(Protein_Count ~ Tissue*Genes,
        data = data2,
        cex.axis = 0.8,
        col = viridis(2, alpha = 0.7),
        lwd = 0.7,
```

```
        ylab = "",
        ylim = c(0, 500000000),
        axes = FALSE,
        frame.plot = TRUE,
        xlab = "")

GROUP <- unique(data2$Genes)
groups <- 70
numbox <- 2
total <- groups * numbox
xpoints <- seq(median(1:numbox), total, numbox)

axis(2, seq(0, 500000000, by = 100000000), las = 1, cex.axis = 0.7)
axis(1, labels = GROUP, at = xpoints, las = 2, cex.axis = 0.7)

stripchart(data2$Protein_Count ~ data2$Tissue*data2$Genes,
           method = "jitter",
           pch = 19,
           col = "black",
           vertical = TRUE,
           add = TRUE,
           cex = 0.3)

title(ylab = expression("Protein abundance (RU)"), line = 3.0)
legend("top", fill = viridis(2), legend = c("Endometrium Follicular", "Endome
trium Luteal"), horiz = T, cex = 0.7)

dev.off()

## quartz_off_screen
##                  2

# Convert Tissue to factor
data2$Tissue <- as.factor(data2$Tissue)

# Create an empty list to store Tukey results
tukey_results_list <- list()

# Unique genes
unique_genes <- unique(data2$Genes)

# Loop over each gene
for (gene in unique_genes) {
  # Subset data for the current gene
  gene_data <- subset(data2, Genes == gene)

  # Fit the generalized linear model
  glm_model <- glm(Protein_Count ~ Tissue, data = gene_data, family = gaussia
n)
```

```
  # Perform Tukey test
  contrast_matrix <- glht(glm_model, linfct = mcp(Tissue = "Tukey"))
  tukey_results <- summary(contrast_matrix)

  # Store the results
  tukey_results_list[[gene]] <- tukey_results
}

# Print the results for each gene
for (gene in unique_genes) {
  cat("Tukey results for gene:", gene, "\n")
  print(tukey_results_list[[gene]])
  cat("\n")
}

## Tukey results for gene: AEBP1
##
##   Simultaneous Tests for General Linear Hypotheses
##
## Multiple Comparisons of Means: Tukey Contrasts
##
##
## Fit: glm(formula = Protein_Count ~ Tissue, family = gaussian, data = gene_
data)
##
## Linear Hypotheses:
##                Estimate Std. Error z value Pr(>|z|)
## EL - EF == 0   760510     976380   0.779    0.436
## (Adjusted p values reported -- single-step method)
##
##
## Tukey results for gene: AGRN
##
##   Simultaneous Tests for General Linear Hypotheses
##
## Multiple Comparisons of Means: Tukey Contrasts
##
##
## Fit: glm(formula = Protein_Count ~ Tissue, family = gaussian, data = gene_
data)
##
## Linear Hypotheses:
##                Estimate Std. Error z value Pr(>|z|)
## EL - EF == 0   121331     281248   0.431    0.666
## (Adjusted p values reported -- single-step method)
##
##
## Tukey results for gene: AHSG
##
```

```
##   Simultaneous Tests for General Linear Hypotheses
##
## Multiple Comparisons of Means: Tukey Contrasts
##
##
## Fit: glm(formula = Protein_Count ~ Tissue, family = gaussian, data = gene_
data)
##
## Linear Hypotheses:
##               Estimate Std. Error z value Pr(>|z|)
## EL - EF == 0   214686     488588   0.439     0.66
## (Adjusted p values reported -- single-step method)
##
##
## Tukey results for gene: CILP2
##
##   Simultaneous Tests for General Linear Hypotheses
##
## Multiple Comparisons of Means: Tukey Contrasts
##
##
## Fit: glm(formula = Protein_Count ~ Tissue, family = gaussian, data = gene_
data)
##
## Linear Hypotheses:
##               Estimate Std. Error z value Pr(>|z|)
## EL - EF == 0 -8319820    8685757  -0.958    0.338
## (Adjusted p values reported -- single-step method)
##
##
## Tukey results for gene: DPT
##
##   Simultaneous Tests for General Linear Hypotheses
##
## Multiple Comparisons of Means: Tukey Contrasts
##
##
## Fit: glm(formula = Protein_Count ~ Tissue, family = gaussian, data = gene_
data)
##
## Linear Hypotheses:
##                Estimate Std. Error z value Pr(>|z|)
## EL - EF == 0 -73661360   77217433  -0.954     0.34
## (Adjusted p values reported -- single-step method)
##
##
## Tukey results for gene: ECM1
##
##   Simultaneous Tests for General Linear Hypotheses
##
```

```
## Multiple Comparisons of Means: Tukey Contrasts
##
##
## Fit: glm(formula = Protein_Count ~ Tissue, family = gaussian, data = gene_
data)
##
## Linear Hypotheses:
##                Estimate Std. Error z value Pr(>|z|)
## EL - EF == 0 -2058056    2001803  -1.028    0.304
## (Adjusted p values reported -- single-step method)
##
##
## Tukey results for gene: ECM2
##
##   Simultaneous Tests for General Linear Hypotheses
##
## Multiple Comparisons of Means: Tukey Contrasts
##
##
## Fit: glm(formula = Protein_Count ~ Tissue, family = gaussian, data = gene_
data)
##
## Linear Hypotheses:
##                Estimate Std. Error z value Pr(>|z|)
## EL - EF == 0    34577     239010   0.145    0.885
## (Adjusted p values reported -- single-step method)
##
##
## Tukey results for gene: EFEMP1
##
##   Simultaneous Tests for General Linear Hypotheses
##
## Multiple Comparisons of Means: Tukey Contrasts
##
##
## Fit: glm(formula = Protein_Count ~ Tissue, family = gaussian, data = gene_
data)
##
## Linear Hypotheses:
##                Estimate Std. Error z value Pr(>|z|)
## EL - EF == 0  -447188     293022  -1.526    0.127
## (Adjusted p values reported -- single-step method)
##
##
## Tukey results for gene: EFEMP2
##
##   Simultaneous Tests for General Linear Hypotheses
##
## Multiple Comparisons of Means: Tukey Contrasts
##
```

```
##
## Fit: glm(formula = Protein_Count ~ Tissue, family = gaussian, data = gene_
data)
##
## Linear Hypotheses:
##              Estimate Std. Error z value Pr(>|z|)
## EL - EF == 0    95721     573030   0.167    0.867
## (Adjusted p values reported -- single-step method)
##
##
## Tukey results for gene: EMID1
##
##   Simultaneous Tests for General Linear Hypotheses
##
## Multiple Comparisons of Means: Tukey Contrasts
##
##
## Fit: glm(formula = Protein_Count ~ Tissue, family = gaussian, data = gene_
data)
##
## Linear Hypotheses:
##              Estimate Std. Error z value Pr(>|z|)
## EL - EF == 0   -61715     183482  -0.336    0.737
## (Adjusted p values reported -- single-step method)
##
##
## Tukey results for gene: EMILIN1
##
##   Simultaneous Tests for General Linear Hypotheses
##
## Multiple Comparisons of Means: Tukey Contrasts
##
##
## Fit: glm(formula = Protein_Count ~ Tissue, family = gaussian, data = gene_
data)
##
## Linear Hypotheses:
##              Estimate Std. Error z value Pr(>|z|)
## EL - EF == 0 -1856058    3433417  -0.541    0.589
## (Adjusted p values reported -- single-step method)
##
##
## Tukey results for gene: EMILIN2
##
##   Simultaneous Tests for General Linear Hypotheses
##
## Multiple Comparisons of Means: Tukey Contrasts
##
##
## Fit: glm(formula = Protein_Count ~ Tissue, family = gaussian, data = gene_
```

```
data)
##
## Linear Hypotheses:
##                Estimate Std. Error z value Pr(>|z|)
## EL - EF == 0  -125622     302869  -0.415    0.678
## (Adjusted p values reported -- single-step method)
##
##
## Tukey results for gene: EMILIN3
##
##   Simultaneous Tests for General Linear Hypotheses
##
## Multiple Comparisons of Means: Tukey Contrasts
##
##
## Fit: glm(formula = Protein_Count ~ Tissue, family = gaussian, data = gene_
data)
##
## Linear Hypotheses:
##                Estimate Std. Error z value Pr(>|z|)
## EL - EF == 0   426412     529441   0.805    0.421
## (Adjusted p values reported -- single-step method)
##
##
## Tukey results for gene: FBLN1
##
##   Simultaneous Tests for General Linear Hypotheses
##
## Multiple Comparisons of Means: Tukey Contrasts
##
##
## Fit: glm(formula = Protein_Count ~ Tissue, family = gaussian, data = gene_
data)
##
## Linear Hypotheses:
##                Estimate Std. Error z value Pr(>|z|)
## EL - EF == 0 -2760098    1656735  -1.666   0.0957 .
## ---
## Signif. codes:  0 '***' 0.001 '**' 0.01 '*' 0.05 '.' 0.1 ' ' 1
## (Adjusted p values reported -- single-step method)
##
##
## Tukey results for gene: FBLN2
##
##   Simultaneous Tests for General Linear Hypotheses
##
## Multiple Comparisons of Means: Tukey Contrasts
##
##
## Fit: glm(formula = Protein_Count ~ Tissue, family = gaussian, data = gene_
```

```
data)
##
## Linear Hypotheses:
##              Estimate Std. Error z value Pr(>|z|)
## EL - EF == 0  -430862     267679   -1.61    0.107
## (Adjusted p values reported -- single-step method)
##
##
## Tukey results for gene: FBLN5
##
##   Simultaneous Tests for General Linear Hypotheses
##
## Multiple Comparisons of Means: Tukey Contrasts
##
##
## Fit: glm(formula = Protein_Count ~ Tissue, family = gaussian, data = gene_
data)
##
## Linear Hypotheses:
##              Estimate Std. Error z value Pr(>|z|)
## EL - EF == 0  1601884    1576022   1.016    0.309
## (Adjusted p values reported -- single-step method)
##
##
## Tukey results for gene: FBN1
##
##   Simultaneous Tests for General Linear Hypotheses
##
## Multiple Comparisons of Means: Tukey Contrasts
##
##
## Fit: glm(formula = Protein_Count ~ Tissue, family = gaussian, data = gene_
data)
##
## Linear Hypotheses:
##              Estimate Std. Error z value Pr(>|z|)
## EL - EF == 0 -2497531    1268641  -1.969    0.049 *
## ---
## Signif. codes:  0 '***' 0.001 '**' 0.01 '*' 0.05 '.' 0.1 ' ' 1
## (Adjusted p values reported -- single-step method)
##
##
## Tukey results for gene: FBN3
##
##   Simultaneous Tests for General Linear Hypotheses
##
## Multiple Comparisons of Means: Tukey Contrasts
##
##
## Fit: glm(formula = Protein_Count ~ Tissue, family = gaussian, data = gene_
```

```
data)
##
## Linear Hypotheses:
##              Estimate Std. Error z value Pr(>|z|)
## EL - EF == 0   -88260     192760  -0.458    0.647
## (Adjusted p values reported -- single-step method)
##
##
## Tukey results for gene: FGA
##
##   Simultaneous Tests for General Linear Hypotheses
##
## Multiple Comparisons of Means: Tukey Contrasts
##
##
## Fit: glm(formula = Protein_Count ~ Tissue, family = gaussian, data = gene_
data)
##
## Linear Hypotheses:
##               Estimate Std. Error z value Pr(>|z|)
## EL - EF == 0 -19565180   17291696  -1.131    0.258
## (Adjusted p values reported -- single-step method)
##
##
## Tukey results for gene: FGB
##
##   Simultaneous Tests for General Linear Hypotheses
##
## Multiple Comparisons of Means: Tukey Contrasts
##
##
## Fit: glm(formula = Protein_Count ~ Tissue, family = gaussian, data = gene_
data)
##
## Linear Hypotheses:
##               Estimate Std. Error z value Pr(>|z|)
## EL - EF == 0 -14349563   16344193  -0.878     0.38
## (Adjusted p values reported -- single-step method)
##
##
## Tukey results for gene: FGG
##
##   Simultaneous Tests for General Linear Hypotheses
##
## Multiple Comparisons of Means: Tukey Contrasts
##
##
## Fit: glm(formula = Protein_Count ~ Tissue, family = gaussian, data = gene_
data)
##
```

```
## Linear Hypotheses:
##                Estimate Std. Error z value Pr(>|z|)
## EL - EF == 0 -11715098   12010182  -0.975    0.329
## (Adjusted p values reported -- single-step method)
##
##
## Tukey results for gene: FN1
##
##   Simultaneous Tests for General Linear Hypotheses
##
## Multiple Comparisons of Means: Tukey Contrasts
##
##
## Fit: glm(formula = Protein_Count ~ Tissue, family = gaussian, data = gene_
data)
##
## Linear Hypotheses:
##               Estimate Std. Error z value Pr(>|z|)
## EL - EF == 0 -8146150    6616296  -1.231    0.218
## (Adjusted p values reported -- single-step method)
##
##
## Tukey results for gene: FNDC1
##
##   Simultaneous Tests for General Linear Hypotheses
##
## Multiple Comparisons of Means: Tukey Contrasts
##
##
## Fit: glm(formula = Protein_Count ~ Tissue, family = gaussian, data = gene_
data)
##
## Linear Hypotheses:
##               Estimate Std. Error z value Pr(>|z|)
## EL - EF == 0   125658      74804    1.68    0.093 .
## ---
## Signif. codes:  0 '***' 0.001 '**' 0.01 '*' 0.05 '.' 0.1 ' ' 1
## (Adjusted p values reported -- single-step method)
##
##
## Tukey results for gene: IGFALS
##
##   Simultaneous Tests for General Linear Hypotheses
##
## Multiple Comparisons of Means: Tukey Contrasts
##
##
## Fit: glm(formula = Protein_Count ~ Tissue, family = gaussian, data = gene_
data)
##
```

```
## Linear Hypotheses:
##              Estimate Std. Error z value Pr(>|z|)
## EL - EF == 0    45553      63598   0.716    0.474
## (Adjusted p values reported -- single-step method)
##
##
## Tukey results for gene: IGFBP6
##
##   Simultaneous Tests for General Linear Hypotheses
##
## Multiple Comparisons of Means: Tukey Contrasts
##
##
## Fit: glm(formula = Protein_Count ~ Tissue, family = gaussian, data = gene_
data)
##
## Linear Hypotheses:
##              Estimate Std. Error z value Pr(>|z|)
## EL - EF == 0  -150546     103674  -1.452    0.146
## (Adjusted p values reported -- single-step method)
##
##
## Tukey results for gene: IGFBP7
##
##   Simultaneous Tests for General Linear Hypotheses
##
## Multiple Comparisons of Means: Tukey Contrasts
##
##
## Fit: glm(formula = Protein_Count ~ Tissue, family = gaussian, data = gene_
data)
##
## Linear Hypotheses:
##              Estimate Std. Error z value Pr(>|z|)
## EL - EF == 0  -432673     533782  -0.811    0.418
## (Adjusted p values reported -- single-step method)
##
##
## Tukey results for gene: LAMA1
##
##   Simultaneous Tests for General Linear Hypotheses
##
## Multiple Comparisons of Means: Tukey Contrasts
##
##
## Fit: glm(formula = Protein_Count ~ Tissue, family = gaussian, data = gene_
data)
##
## Linear Hypotheses:
##              Estimate Std. Error z value Pr(>|z|)
```

```
## EL - EF == 0 -2599007    2160422  -1.203    0.229
## (Adjusted p values reported -- single-step method)
##
##
## Tukey results for gene: LAMA2
##
##   Simultaneous Tests for General Linear Hypotheses
##
## Multiple Comparisons of Means: Tukey Contrasts
##
##
## Fit: glm(formula = Protein_Count ~ Tissue, family = gaussian, data = gene_
data)
##
## Linear Hypotheses:
##              Estimate Std. Error z value Pr(>|z|)
## EL - EF == 0   -42359     115770  -0.366    0.714
## (Adjusted p values reported -- single-step method)
##
##
## Tukey results for gene: LAMA3
##
##   Simultaneous Tests for General Linear Hypotheses
##
## Multiple Comparisons of Means: Tukey Contrasts
##
##
## Fit: glm(formula = Protein_Count ~ Tissue, family = gaussian, data = gene_
data)
##
## Linear Hypotheses:
##              Estimate Std. Error z value Pr(>|z|)
## EL - EF == 0  -313803     178358  -1.759   0.0785 .
## ---
## Signif. codes:  0 '***' 0.001 '**' 0.01 '*' 0.05 '.' 0.1 ' ' 1
## (Adjusted p values reported -- single-step method)
##
##
## Tukey results for gene: LAMA4
##
##   Simultaneous Tests for General Linear Hypotheses
##
## Multiple Comparisons of Means: Tukey Contrasts
##
##
## Fit: glm(formula = Protein_Count ~ Tissue, family = gaussian, data = gene_
data)
##
## Linear Hypotheses:
##              Estimate Std. Error z value Pr(>|z|)
```

```
## EL - EF == 0  -749402    1055190   -0.71    0.478
## (Adjusted p values reported -- single-step method)
##
##
## Tukey results for gene: LAMA5
##
##   Simultaneous Tests for General Linear Hypotheses
##
## Multiple Comparisons of Means: Tukey Contrasts
##
##
## Fit: glm(formula = Protein_Count ~ Tissue, family = gaussian, data = gene_
data)
##
## Linear Hypotheses:
##              Estimate Std. Error z value Pr(>|z|)
## EL - EF == 0  -800588     793815  -1.009    0.313
## (Adjusted p values reported -- single-step method)
##
##
## Tukey results for gene: LAMB1
##
##   Simultaneous Tests for General Linear Hypotheses
##
## Multiple Comparisons of Means: Tukey Contrasts
##
##
## Fit: glm(formula = Protein_Count ~ Tissue, family = gaussian, data = gene_
data)
##
## Linear Hypotheses:
##              Estimate Std. Error z value Pr(>|z|)
## EL - EF == 0  -476137     568950  -0.837    0.403
## (Adjusted p values reported -- single-step method)
##
##
## Tukey results for gene: LAMB2
##
##   Simultaneous Tests for General Linear Hypotheses
##
## Multiple Comparisons of Means: Tukey Contrasts
##
##
## Fit: glm(formula = Protein_Count ~ Tissue, family = gaussian, data = gene_
data)
##
## Linear Hypotheses:
##              Estimate Std. Error z value Pr(>|z|)
## EL - EF == 0 -2061146    2307888  -0.893    0.372
## (Adjusted p values reported -- single-step method)
```

```
##
##
## Tukey results for gene: LAMB3
##
##   Simultaneous Tests for General Linear Hypotheses
##
## Multiple Comparisons of Means: Tukey Contrasts
##
##
## Fit: glm(formula = Protein_Count ~ Tissue, family = gaussian, data = gene_
data)
##
## Linear Hypotheses:
##               Estimate Std. Error z value Pr(>|z|)
## EL - EF == 0  -163274     137350  -1.189    0.235
## (Adjusted p values reported -- single-step method)
##
##
## Tukey results for gene: LAMC1
##
##   Simultaneous Tests for General Linear Hypotheses
##
## Multiple Comparisons of Means: Tukey Contrasts
##
##
## Fit: glm(formula = Protein_Count ~ Tissue, family = gaussian, data = gene_
data)
##
## Linear Hypotheses:
##               Estimate Std. Error z value Pr(>|z|)
## EL - EF == 0 -1893466    2034630  -0.931    0.352
## (Adjusted p values reported -- single-step method)
##
##
## Tukey results for gene: LAMC2
##
##   Simultaneous Tests for General Linear Hypotheses
##
## Multiple Comparisons of Means: Tukey Contrasts
##
##
## Fit: glm(formula = Protein_Count ~ Tissue, family = gaussian, data = gene_
data)
##
## Linear Hypotheses:
##               Estimate Std. Error z value Pr(>|z|)
## EL - EF == 0   -73359     112824   -0.65    0.516
## (Adjusted p values reported -- single-step method)
##
##
```

```
## Tukey results for gene: LAMC3
##
##   Simultaneous Tests for General Linear Hypotheses
##
## Multiple Comparisons of Means: Tukey Contrasts
##
##
## Fit: glm(formula = Protein_Count ~ Tissue, family = gaussian, data = gene_
data)
##
## Linear Hypotheses:
##              Estimate Std. Error z value Pr(>|z|)
## EL - EF == 0    18129      54767   0.331    0.741
## (Adjusted p values reported -- single-step method)
##
##
## Tukey results for gene: LRG1
##
##   Simultaneous Tests for General Linear Hypotheses
##
## Multiple Comparisons of Means: Tukey Contrasts
##
##
## Fit: glm(formula = Protein_Count ~ Tissue, family = gaussian, data = gene_
data)
##
## Linear Hypotheses:
##              Estimate Std. Error z value Pr(>|z|)
## EL - EF == 0   -15976      66092  -0.242    0.809
## (Adjusted p values reported -- single-step method)
##
##
## Tukey results for gene: LTBP1
##
##   Simultaneous Tests for General Linear Hypotheses
##
## Multiple Comparisons of Means: Tukey Contrasts
##
##
## Fit: glm(formula = Protein_Count ~ Tissue, family = gaussian, data = gene_
data)
##
## Linear Hypotheses:
##              Estimate Std. Error z value Pr(>|z|)
## EL - EF == 0  -111526     140896  -0.792    0.429
## (Adjusted p values reported -- single-step method)
##
##
## Tukey results for gene: LTBP2
##
```

```
##   Simultaneous Tests for General Linear Hypotheses
##
## Multiple Comparisons of Means: Tukey Contrasts
##
##
## Fit: glm(formula = Protein_Count ~ Tissue, family = gaussian, data = gene_
data)
##
## Linear Hypotheses:
##               Estimate Std. Error z value Pr(>|z|)
## EL - EF == 0   402577    2111601   0.191    0.849
## (Adjusted p values reported -- single-step method)
##
##
## Tukey results for gene: LTBP4
##
##   Simultaneous Tests for General Linear Hypotheses
##
## Multiple Comparisons of Means: Tukey Contrasts
##
##
## Fit: glm(formula = Protein_Count ~ Tissue, family = gaussian, data = gene_
data)
##
## Linear Hypotheses:
##               Estimate Std. Error z value Pr(>|z|)
## EL - EF == 0    178.4   395274.2       0        1
## (Adjusted p values reported -- single-step method)
##
##
## Tukey results for gene: MATN2
##
##   Simultaneous Tests for General Linear Hypotheses
##
## Multiple Comparisons of Means: Tukey Contrasts
##
##
## Fit: glm(formula = Protein_Count ~ Tissue, family = gaussian, data = gene_
data)
##
## Linear Hypotheses:
##               Estimate Std. Error z value Pr(>|z|)
## EL - EF == 0   -99542     517406  -0.192    0.847
## (Adjusted p values reported -- single-step method)
##
##
## Tukey results for gene: MFAP2
##
##   Simultaneous Tests for General Linear Hypotheses
##
```

```
## Multiple Comparisons of Means: Tukey Contrasts
##
##
## Fit: glm(formula = Protein_Count ~ Tissue, family = gaussian, data = gene_
data)
##
## Linear Hypotheses:
##              Estimate Std. Error z value Pr(>|z|)
## EL - EF == 0  -734246    2414638  -0.304    0.761
## (Adjusted p values reported -- single-step method)
##
##
## Tukey results for gene: MFAP4
##
##   Simultaneous Tests for General Linear Hypotheses
##
## Multiple Comparisons of Means: Tukey Contrasts
##
##
## Fit: glm(formula = Protein_Count ~ Tissue, family = gaussian, data = gene_
data)
##
## Linear Hypotheses:
##              Estimate Std. Error z value Pr(>|z|)
## EL - EF == 0 -1592704    1939159  -0.821    0.411
## (Adjusted p values reported -- single-step method)
##
##
## Tukey results for gene: MFAP5
##
##   Simultaneous Tests for General Linear Hypotheses
##
## Multiple Comparisons of Means: Tukey Contrasts
##
##
## Fit: glm(formula = Protein_Count ~ Tissue, family = gaussian, data = gene_
data)
##
## Linear Hypotheses:
##              Estimate Std. Error z value Pr(>|z|)
## EL - EF == 0 -1657016    1299972  -1.275    0.202
## (Adjusted p values reported -- single-step method)
##
##
## Tukey results for gene: MFGE8
##
##   Simultaneous Tests for General Linear Hypotheses
##
## Multiple Comparisons of Means: Tukey Contrasts
##
```

```
##
## Fit: glm(formula = Protein_Count ~ Tissue, family = gaussian, data = gene_
data)
##
## Linear Hypotheses:
##                Estimate Std. Error z value Pr(>|z|)
## EL - EF == 0   -47564     126700  -0.375    0.707
## (Adjusted p values reported -- single-step method)
##
##
## Tukey results for gene: MGP
##
##   Simultaneous Tests for General Linear Hypotheses
##
## Multiple Comparisons of Means: Tukey Contrasts
##
##
## Fit: glm(formula = Protein_Count ~ Tissue, family = gaussian, data = gene_
data)
##
## Linear Hypotheses:
##                Estimate Std. Error z value Pr(>|z|)
## EL - EF == 0 -3465614    2603140  -1.331    0.183
## (Adjusted p values reported -- single-step method)
##
##
## Tukey results for gene: MMRN2
##
##   Simultaneous Tests for General Linear Hypotheses
##
## Multiple Comparisons of Means: Tukey Contrasts
##
##
## Fit: glm(formula = Protein_Count ~ Tissue, family = gaussian, data = gene_
data)
##
## Linear Hypotheses:
##                Estimate Std. Error z value Pr(>|z|)
## EL - EF == 0   116400     157174   0.741    0.459
## (Adjusted p values reported -- single-step method)
##
##
## Tukey results for gene: MXRA5
##
##   Simultaneous Tests for General Linear Hypotheses
##
## Multiple Comparisons of Means: Tukey Contrasts
##
##
## Fit: glm(formula = Protein_Count ~ Tissue, family = gaussian, data = gene_
```

```
data)
##
## Linear Hypotheses:
##              Estimate Std. Error z value Pr(>|z|)
## EL - EF == 0    60192     183863   0.327    0.743
## (Adjusted p values reported -- single-step method)
##
##
## Tukey results for gene: NID1
##
##   Simultaneous Tests for General Linear Hypotheses
##
## Multiple Comparisons of Means: Tukey Contrasts
##
##
## Fit: glm(formula = Protein_Count ~ Tissue, family = gaussian, data = gene_
data)
##
## Linear Hypotheses:
##              Estimate Std. Error z value Pr(>|z|)
## EL - EF == 0 -7258840    9950866  -0.729    0.466
## (Adjusted p values reported -- single-step method)
##
##
## Tukey results for gene: NID2
##
##   Simultaneous Tests for General Linear Hypotheses
##
## Multiple Comparisons of Means: Tukey Contrasts
##
##
## Fit: glm(formula = Protein_Count ~ Tissue, family = gaussian, data = gene_
data)
##
## Linear Hypotheses:
##              Estimate Std. Error z value Pr(>|z|)
## EL - EF == 0 -5802740    5149860  -1.127     0.26
## (Adjusted p values reported -- single-step method)
##
##
## Tukey results for gene: PAPLN
##
##   Simultaneous Tests for General Linear Hypotheses
##
## Multiple Comparisons of Means: Tukey Contrasts
##
##
## Fit: glm(formula = Protein_Count ~ Tissue, family = gaussian, data = gene_
data)
##
```

```
## Linear Hypotheses:
##              Estimate Std. Error z value Pr(>|z|)
## EL - EF == 0   113562     101192   1.122    0.262
## (Adjusted p values reported -- single-step method)
##
##
## Tukey results for gene: PCOLCE
##
##   Simultaneous Tests for General Linear Hypotheses
##
## Multiple Comparisons of Means: Tukey Contrasts
##
##
## Fit: glm(formula = Protein_Count ~ Tissue, family = gaussian, data = gene_
data)
##
## Linear Hypotheses:
##              Estimate Std. Error z value Pr(>|z|)
## EL - EF == 0  -137543     426056  -0.323    0.747
## (Adjusted p values reported -- single-step method)
##
##
## Tukey results for gene: POSTN
##
##   Simultaneous Tests for General Linear Hypotheses
##
## Multiple Comparisons of Means: Tukey Contrasts
##
##
## Fit: glm(formula = Protein_Count ~ Tissue, family = gaussian, data = gene_
data)
##
## Linear Hypotheses:
##              Estimate Std. Error z value Pr(>|z|)
## EL - EF == 0  -363900     470087  -0.774    0.439
## (Adjusted p values reported -- single-step method)
##
##
## Tukey results for gene: PXDN
##
##   Simultaneous Tests for General Linear Hypotheses
##
## Multiple Comparisons of Means: Tukey Contrasts
##
##
## Fit: glm(formula = Protein_Count ~ Tissue, family = gaussian, data = gene_
data)
##
## Linear Hypotheses:
##              Estimate Std. Error z value Pr(>|z|)
```

```
## EL - EF == 0   -87628     194381  -0.451    0.652
## (Adjusted p values reported -- single-step method)
##
##
## Tukey results for gene: SBSPON
##
##   Simultaneous Tests for General Linear Hypotheses
##
## Multiple Comparisons of Means: Tukey Contrasts
##
##
## Fit: glm(formula = Protein_Count ~ Tissue, family = gaussian, data = gene_
data)
##
## Linear Hypotheses:
##              Estimate Std. Error z value Pr(>|z|)
## EL - EF == 0 -1168810    1159638  -1.008    0.313
## (Adjusted p values reported -- single-step method)
##
##
## Tukey results for gene: SMOC2
##
##   Simultaneous Tests for General Linear Hypotheses
##
## Multiple Comparisons of Means: Tukey Contrasts
##
##
## Fit: glm(formula = Protein_Count ~ Tissue, family = gaussian, data = gene_
data)
##
## Linear Hypotheses:
##              Estimate Std. Error z value Pr(>|z|)
## EL - EF == 0    46279      47653   0.971    0.331
## (Adjusted p values reported -- single-step method)
##
##
## Tukey results for gene: SPARC
##
##   Simultaneous Tests for General Linear Hypotheses
##
## Multiple Comparisons of Means: Tukey Contrasts
##
##
## Fit: glm(formula = Protein_Count ~ Tissue, family = gaussian, data = gene_
data)
##
## Linear Hypotheses:
##              Estimate Std. Error z value Pr(>|z|)
## EL - EF == 0    29119    1647199   0.018    0.986
## (Adjusted p values reported -- single-step method)
```

```
##
##
## Tukey results for gene: SPARCL1
##
##   Simultaneous Tests for General Linear Hypotheses
##
## Multiple Comparisons of Means: Tukey Contrasts
##
##
## Fit: glm(formula = Protein_Count ~ Tissue, family = gaussian, data = gene_
data)
##
## Linear Hypotheses:
##               Estimate Std. Error z value Pr(>|z|)
## EL - EF == 0    89901     102350   0.878     0.38
## (Adjusted p values reported -- single-step method)
##
##
## Tukey results for gene: SPON1
##
##   Simultaneous Tests for General Linear Hypotheses
##
## Multiple Comparisons of Means: Tukey Contrasts
##
##
## Fit: glm(formula = Protein_Count ~ Tissue, family = gaussian, data = gene_
data)
##
## Linear Hypotheses:
##               Estimate Std. Error z value Pr(>|z|)
## EL - EF == 0   487113     339075   1.437    0.151
## (Adjusted p values reported -- single-step method)
##
##
## Tukey results for gene: SRPX2
##
##   Simultaneous Tests for General Linear Hypotheses
##
## Multiple Comparisons of Means: Tukey Contrasts
##
##
## Fit: glm(formula = Protein_Count ~ Tissue, family = gaussian, data = gene_
data)
##
## Linear Hypotheses:
##               Estimate Std. Error z value Pr(>|z|)
## EL - EF == 0    16219      82786   0.196    0.845
## (Adjusted p values reported -- single-step method)
##
##
```

```
## Tukey results for gene: TGFBI
##
##   Simultaneous Tests for General Linear Hypotheses
##
## Multiple Comparisons of Means: Tukey Contrasts
##
##
## Fit: glm(formula = Protein_Count ~ Tissue, family = gaussian, data = gene_
data)
##
## Linear Hypotheses:
##               Estimate Std. Error z value Pr(>|z|)
## EL - EF == 0 -2568956    4060489  -0.633    0.527
## (Adjusted p values reported -- single-step method)
##
##
## Tukey results for gene: THBS1
##
##   Simultaneous Tests for General Linear Hypotheses
##
## Multiple Comparisons of Means: Tukey Contrasts
##
##
## Fit: glm(formula = Protein_Count ~ Tissue, family = gaussian, data = gene_
data)
##
## Linear Hypotheses:
##               Estimate Std. Error z value Pr(>|z|)
## EL - EF == 0   -69111     174678  -0.396    0.692
## (Adjusted p values reported -- single-step method)
##
##
## Tukey results for gene: THSD4
##
##   Simultaneous Tests for General Linear Hypotheses
##
## Multiple Comparisons of Means: Tukey Contrasts
##
##
## Fit: glm(formula = Protein_Count ~ Tissue, family = gaussian, data = gene_
data)
##
## Linear Hypotheses:
##               Estimate Std. Error z value Pr(>|z|)
## EL - EF == 0    45481     116516    0.39    0.696
## (Adjusted p values reported -- single-step method)
##
##
## Tukey results for gene: TINAGL1
##
```

```
##   Simultaneous Tests for General Linear Hypotheses
##
## Multiple Comparisons of Means: Tukey Contrasts
##
##
## Fit: glm(formula = Protein_Count ~ Tissue, family = gaussian, data = gene_
data)
##
## Linear Hypotheses:
##                 Estimate Std. Error z value Pr(>|z|)
## EL - EF == 0 -15477100   10898649   -1.42    0.156
## (Adjusted p values reported -- single-step method)
##
##
## Tukey results for gene: TNC
##
##   Simultaneous Tests for General Linear Hypotheses
##
## Multiple Comparisons of Means: Tukey Contrasts
##
##
## Fit: glm(formula = Protein_Count ~ Tissue, family = gaussian, data = gene_
data)
##
## Linear Hypotheses:
##                Estimate Std. Error z value Pr(>|z|)
## EL - EF == 0 -1309708    1141813  -1.147    0.251
## (Adjusted p values reported -- single-step method)
##
##
## Tukey results for gene: TNXB
##
##   Simultaneous Tests for General Linear Hypotheses
##
## Multiple Comparisons of Means: Tukey Contrasts
##
##
## Fit: glm(formula = Protein_Count ~ Tissue, family = gaussian, data = gene_
data)
##
## Linear Hypotheses:
##                Estimate Std. Error z value Pr(>|z|)
## EL - EF == 0  -351362     854672  -0.411    0.681
## (Adjusted p values reported -- single-step method)
##
##
## Tukey results for gene: VTN
##
##   Simultaneous Tests for General Linear Hypotheses
##
```

```
## Multiple Comparisons of Means: Tukey Contrasts
##
##
## Fit: glm(formula = Protein_Count ~ Tissue, family = gaussian, data = gene_
data)
##
## Linear Hypotheses:
##              Estimate Std. Error z value Pr(>|z|)
## EL - EF == 0 -2682702    2670440  -1.005    0.315
## (Adjusted p values reported -- single-step method)
##
##
## Tukey results for gene: VWA1
##
##   Simultaneous Tests for General Linear Hypotheses
##
## Multiple Comparisons of Means: Tukey Contrasts
##
##
## Fit: glm(formula = Protein_Count ~ Tissue, family = gaussian, data = gene_
data)
##
## Linear Hypotheses:
##              Estimate Std. Error z value Pr(>|z|)
## EL - EF == 0 -1444660    1810706  -0.798    0.425
## (Adjusted p values reported -- single-step method)
##
##
## Tukey results for gene: VWF
##
##   Simultaneous Tests for General Linear Hypotheses
##
## Multiple Comparisons of Means: Tukey Contrasts
##
##
## Fit: glm(formula = Protein_Count ~ Tissue, family = gaussian, data = gene_
data)
##
## Linear Hypotheses:
##              Estimate Std. Error z value Pr(>|z|)
## EL - EF == 0  -299948     186922  -1.605    0.109
## (Adjusted p values reported -- single-step method)
```

```
#### Proteoglycans

# Read your cleaned joined data file into a data frame
cleaned_joined_data <- read.csv("cleaned_matrisome_endo.csv")

# Filter rows with Matrisome_Category "Proteoglycans"
Proteoglycans_data <- cleaned_joined_data %>%
```

```
  filter(Matrisome_Category == "Proteoglycans")

# Reshape the data to have separate columns for each replicate and each tissu
e stage
organized_data <- Proteoglycans_data %>%
  pivot_longer(cols = starts_with("EF") | starts_with("EL"),
               names_to = c("Tissue", "Stage"),
               names_sep = "\\.",
               values_to = "Protein_Count")

## Warning: Expected 2 pieces. Missing pieces filled with `NA` in 2 rows [1,
6].

# Save the reshaped data to a new file

write.csv(organized_data, "Proteoglycans_data_matrisome_endo_to_clean.csv", r
ow.names = FALSE)

data2 <- read.csv("Proteoglycans_data_matrisome_endo_clean.csv")

png(filename="endo_Proteoglycans.png",
    width = 12, height = 6, units = "in",
    res = 600)

boxplot(Protein_Count ~ Tissue*Genes,
        data = data2,
        cex.axis = 0.8,
        col = viridis(2, alpha = 0.7),
        lwd = 0.7,
        ylab = "",
        ylim = c(0, 200000000),
        axes = FALSE,
        frame.plot = TRUE,
        xlab = "")

GROUP <- unique(data2$Genes)
groups <- 13
numbox <- 2
total <- groups * numbox
xpoints <- seq(median(1:numbox), total, numbox)

axis(2, seq(0, 200000000, by = 10000000), las = 1, cex.axis = 0.7)
axis(1, labels = GROUP, at = xpoints, las = 2, cex.axis = 0.7)

stripchart(data2$Protein_Count ~ data2$Tissue*data2$Genes,
           method = "jitter",
           pch = 19,
           col = "black",
           vertical = TRUE,
           add = TRUE,
```

```r
           cex = 0.3)

title(ylab = expression("Protein abundance (RU)"), line = 3.0)
legend("top", fill = viridis(2), legend = c("Endometrium Follicular", "Endome
trium Luteal"), horiz = T, cex = 0.7)

dev.off()

## quartz_off_screen
##                 2

# Convert Tissue to factor
data2$Tissue <- as.factor(data2$Tissue)

# Create an empty list to store Tukey results
tukey_results_list <- list()

# Unique genes
unique_genes <- unique(data2$Genes)

# Loop over each gene
for (gene in unique_genes) {
  # Subset data for the current gene
  gene_data <- subset(data2, Genes == gene)

  # Fit the generalized linear model
  glm_model <- glm(Protein_Count ~ Tissue, data = gene_data, family = gaussia
n)

  # Perform Tukey test
  contrast_matrix <- glht(glm_model, linfct = mcp(Tissue = "Tukey"))
  tukey_results <- summary(contrast_matrix)

  # Store the results
  tukey_results_list[[gene]] <- tukey_results
}

# Print the results for each gene
for (gene in unique_genes) {
  cat("Tukey results for gene:", gene, "\n")
  print(tukey_results_list[[gene]])
  cat("\n")
}

## Tukey results for gene: ACAN
##
##   Simultaneous Tests for General Linear Hypotheses
##
## Multiple Comparisons of Means: Tukey Contrasts
##
```

```
##
## Fit: glm(formula = Protein_Count ~ Tissue, family = gaussian, data = gene_
data)
##
## Linear Hypotheses:
##              Estimate Std. Error z value Pr(>|z|)
## EL - EF == 0    -6130     160521  -0.038     0.97
## (Adjusted p values reported -- single-step method)
##
##
## Tukey results for gene: ASPN
##
##   Simultaneous Tests for General Linear Hypotheses
##
## Multiple Comparisons of Means: Tukey Contrasts
##
##
## Fit: glm(formula = Protein_Count ~ Tissue, family = gaussian, data = gene_
data)
##
## Linear Hypotheses:
##              Estimate Std. Error z value Pr(>|z|)
## EL - EF == 0   379598    4444235   0.085    0.932
## (Adjusted p values reported -- single-step method)
##
##
## Tukey results for gene: BGN
##
##   Simultaneous Tests for General Linear Hypotheses
##
## Multiple Comparisons of Means: Tukey Contrasts
##
##
## Fit: glm(formula = Protein_Count ~ Tissue, family = gaussian, data = gene_
data)
##
## Linear Hypotheses:
##              Estimate Std. Error z value Pr(>|z|)
## EL - EF == 0   380404     442282    0.86     0.39
## (Adjusted p values reported -- single-step method)
##
##
## Tukey results for gene: DCN
##
##   Simultaneous Tests for General Linear Hypotheses
##
## Multiple Comparisons of Means: Tukey Contrasts
##
##
## Fit: glm(formula = Protein_Count ~ Tissue, family = gaussian, data = gene_
```

```
data)
##
## Linear Hypotheses:
##               Estimate Std. Error z value Pr(>|z|)
## EL - EF == 0  4510502    8756673   0.515    0.606
## (Adjusted p values reported -- single-step method)
##
##
## Tukey results for gene: FMOD
##
##   Simultaneous Tests for General Linear Hypotheses
##
## Multiple Comparisons of Means: Tukey Contrasts
##
##
## Fit: glm(formula = Protein_Count ~ Tissue, family = gaussian, data = gene_
data)
##
## Linear Hypotheses:
##               Estimate Std. Error z value Pr(>|z|)
## EL - EF == 0  4634398    5269343    0.88    0.379
## (Adjusted p values reported -- single-step method)
##
##
## Tukey results for gene: HAPLN1
##
##   Simultaneous Tests for General Linear Hypotheses
##
## Multiple Comparisons of Means: Tukey Contrasts
##
##
## Fit: glm(formula = Protein_Count ~ Tissue, family = gaussian, data = gene_
data)
##
## Linear Hypotheses:
##               Estimate Std. Error z value Pr(>|z|)
## EL - EF == 0  -116907     244432  -0.478    0.632
## (Adjusted p values reported -- single-step method)
##
##
## Tukey results for gene: HSPG2
##
##   Simultaneous Tests for General Linear Hypotheses
##
## Multiple Comparisons of Means: Tukey Contrasts
##
##
## Fit: glm(formula = Protein_Count ~ Tissue, family = gaussian, data = gene_
data)
##
```

```
## Linear Hypotheses:
##              Estimate Std. Error z value Pr(>|z|)
## EL - EF == 0 -1885142    2234424  -0.844    0.399
## (Adjusted p values reported -- single-step method)
##
##
## Tukey results for gene: KERA
##
##   Simultaneous Tests for General Linear Hypotheses
##
## Multiple Comparisons of Means: Tukey Contrasts
##
##
## Fit: glm(formula = Protein_Count ~ Tissue, family = gaussian, data = gene_
data)
##
## Linear Hypotheses:
##              Estimate Std. Error z value Pr(>|z|)
## EL - EF == 0    49186      98761   0.498    0.618
## (Adjusted p values reported -- single-step method)
##
##
## Tukey results for gene: LUM
##
##   Simultaneous Tests for General Linear Hypotheses
##
## Multiple Comparisons of Means: Tukey Contrasts
##
##
## Fit: glm(formula = Protein_Count ~ Tissue, family = gaussian, data = gene_
data)
##
## Linear Hypotheses:
##               Estimate Std. Error z value Pr(>|z|)
## EL - EF == 0 -14036020   26183574  -0.536    0.592
## (Adjusted p values reported -- single-step method)
##
##
## Tukey results for gene: OGN
##
##   Simultaneous Tests for General Linear Hypotheses
##
## Multiple Comparisons of Means: Tukey Contrasts
##
##
## Fit: glm(formula = Protein_Count ~ Tissue, family = gaussian, data = gene_
data)
##
## Linear Hypotheses:
##               Estimate Std. Error z value Pr(>|z|)
```

```
## EL - EF == 0 -10003518   16393746   -0.61    0.542
## (Adjusted p values reported -- single-step method)
##
##
## Tukey results for gene: PODN
##
##   Simultaneous Tests for General Linear Hypotheses
##
## Multiple Comparisons of Means: Tukey Contrasts
##
##
## Fit: glm(formula = Protein_Count ~ Tissue, family = gaussian, data = gene_
data)
##
## Linear Hypotheses:
##              Estimate Std. Error z value Pr(>|z|)
## EL - EF == 0   163117     156429   1.043    0.297
## (Adjusted p values reported -- single-step method)
##
##
## Tukey results for gene: PRELP
##
##   Simultaneous Tests for General Linear Hypotheses
##
## Multiple Comparisons of Means: Tukey Contrasts
##
##
## Fit: glm(formula = Protein_Count ~ Tissue, family = gaussian, data = gene_
data)
##
## Linear Hypotheses:
##              Estimate Std. Error z value Pr(>|z|)
## EL - EF == 0  3162700    6059724   0.522    0.602
## (Adjusted p values reported -- single-step method)
##
##
## Tukey results for gene: VCAN
##
##   Simultaneous Tests for General Linear Hypotheses
##
## Multiple Comparisons of Means: Tukey Contrasts
##
##
## Fit: glm(formula = Protein_Count ~ Tissue, family = gaussian, data = gene_
data)
##
## Linear Hypotheses:
##              Estimate Std. Error z value Pr(>|z|)
## EL - EF == 0 -2382266    1916634  -1.243    0.214
## (Adjusted p values reported -- single-step method)
```

```
#### ECM regulators

# Read your cleaned joined data file into a data frame
cleaned_joined_data <- read.csv("cleaned_matrisome_endo.csv")

# Filter rows with Matrisome_Category "ECM regulators"
ECM_regulators_data <- cleaned_joined_data %>%
  filter(Matrisome_Category == "ECM Regulators")

# Reshape the data to have separate columns for each replicate and each tissu
e stage
organized_data <- ECM_regulators_data %>%
  pivot_longer(cols = starts_with("EF") | starts_with("EL"),
               names_to = c("Tissue", "Stage"),
               names_sep = "\\.",
               values_to = "Protein_Count")

## Warning: Expected 2 pieces. Missing pieces filled with `NA` in 2 rows [1,
6].

# Save the reshaped data to a new file

write.csv(organized_data, "ECM regulators_data_matrisome_endo_to_clean.csv",
row.names = FALSE)

data2 <- read.csv("ECM regulators_data_matrisome_endo_clean.csv")

png(filename="endo_ECM regulators.png",
    width = 12, height = 6, units = "in",
    res = 600)

boxplot(Protein_Count ~ Tissue*Genes,
        data = data2,
        cex.axis = 0.8,
        col = viridis(2, alpha = 0.7),
        lwd = 0.7,
        ylab = "",
        ylim = c(0, 250000000),
        axes = FALSE,
        frame.plot = TRUE,
        xlab = "")

GROUP <- unique(data2$Genes)
groups <- 50
numbox <- 2
total <- groups * numbox
xpoints <- seq(median(1:numbox), total, numbox)

axis(2, seq(0, 250000000, by = 10000000), las = 1, cex.axis = 0.7)
```

```
axis(1, labels = GROUP, at = xpoints, las = 2, cex.axis = 0.7)

stripchart(data2$Protein_Count ~ data2$Tissue*data2$Genes,
           method = "jitter",
           pch = 19,
           col = "black",
           vertical = TRUE,
           add = TRUE,
           cex = 0.3)

title(ylab = expression("Protein abundance (RU)"), line = 3.0)
legend("top", fill = viridis(2), legend = c("Endometrium Follicular", "Endome
trium Luteal"), horiz = T, cex = 0.7)

dev.off()

## quartz_off_screen
##                 2

# Convert Tissue to factor
data2$Tissue <- as.factor(data2$Tissue)

# Create an empty list to store Tukey results
tukey_results_list <- list()

# Unique genes
unique_genes <- unique(data2$Genes)

# Loop over each gene
for (gene in unique_genes) {
  # Subset data for the current gene
  gene_data <- subset(data2, Genes == gene)

  # Fit the generalized linear model
  glm_model <- glm(Protein_Count ~ Tissue, data = gene_data, family = gaussia
n)

  # Perform Tukey test
  contrast_matrix <- glht(glm_model, linfct = mcp(Tissue = "Tukey"))
  tukey_results <- summary(contrast_matrix)

  # Store the results
  tukey_results_list[[gene]] <- tukey_results
}

# Print the results for each gene
for (gene in unique_genes) {
  cat("Tukey results for gene:", gene, "\n")
  print(tukey_results_list[[gene]])
```

```
  cat("\n")
}

## Tukey results for gene: A2M
##
##   Simultaneous Tests for General Linear Hypotheses
##
## Multiple Comparisons of Means: Tukey Contrasts
##
##
## Fit: glm(formula = Protein_Count ~ Tissue, family = gaussian, data = gene_
data)
##
## Linear Hypotheses:
##               Estimate Std. Error z value Pr(>|z|)
## EL - EF == 0  -564470    1296138  -0.436    0.663
## (Adjusted p values reported -- single-step method)
##
##
## Tukey results for gene: ADAM10
##
##   Simultaneous Tests for General Linear Hypotheses
##
## Multiple Comparisons of Means: Tukey Contrasts
##
##
## Fit: glm(formula = Protein_Count ~ Tissue, family = gaussian, data = gene_
data)
##
## Linear Hypotheses:
##               Estimate Std. Error z value Pr(>|z|)
## EL - EF == 0   -99481     308953  -0.322    0.747
## (Adjusted p values reported -- single-step method)
##
##
## Tukey results for gene: ADAMTS5
##
##   Simultaneous Tests for General Linear Hypotheses
##
## Multiple Comparisons of Means: Tukey Contrasts
##
##
## Fit: glm(formula = Protein_Count ~ Tissue, family = gaussian, data = gene_
data)
##
## Linear Hypotheses:
##               Estimate Std. Error z value Pr(>|z|)
## EL - EF == 0  -239796     256425  -0.935     0.35
## (Adjusted p values reported -- single-step method)
##
```

```
##
## Tukey results for gene: ADAMTSL5
##
##   Simultaneous Tests for General Linear Hypotheses
##
## Multiple Comparisons of Means: Tukey Contrasts
##
##
## Fit: glm(formula = Protein_Count ~ Tissue, family = gaussian, data = gene_
data)
##
## Linear Hypotheses:
##              Estimate Std. Error z value Pr(>|z|)
## EL - EF == 0  -261718     343106  -0.763    0.446
## (Adjusted p values reported -- single-step method)
##
##
## Tukey results for gene: CELA3B
##
##   Simultaneous Tests for General Linear Hypotheses
##
## Multiple Comparisons of Means: Tukey Contrasts
##
##
## Fit: glm(formula = Protein_Count ~ Tissue, family = gaussian, data = gene_
data)
##
## Linear Hypotheses:
##              Estimate Std. Error z value Pr(>|z|)
## EL - EF == 0 -2123886    1815118   -1.17    0.242
## (Adjusted p values reported -- single-step method)
##
##
## Tukey results for gene: CSTB
##
##   Simultaneous Tests for General Linear Hypotheses
##
## Multiple Comparisons of Means: Tukey Contrasts
##
##
## Fit: glm(formula = Protein_Count ~ Tissue, family = gaussian, data = gene_
data)
##
## Linear Hypotheses:
##              Estimate Std. Error z value Pr(>|z|)
## EL - EF == 0   -32818      72520  -0.453    0.651
## (Adjusted p values reported -- single-step method)
##
##
## Tukey results for gene: CTSC
```

```
##
##   Simultaneous Tests for General Linear Hypotheses
##
## Multiple Comparisons of Means: Tukey Contrasts
##
##
## Fit: glm(formula = Protein_Count ~ Tissue, family = gaussian, data = gene_
data)
##
## Linear Hypotheses:
##                Estimate Std. Error z value Pr(>|z|)
## EL - EF == 0  -126954     143295  -0.886    0.376
## (Adjusted p values reported -- single-step method)
##
##
## Tukey results for gene: CTSD
##
##   Simultaneous Tests for General Linear Hypotheses
##
## Multiple Comparisons of Means: Tukey Contrasts
##
##
## Fit: glm(formula = Protein_Count ~ Tissue, family = gaussian, data = gene_
data)
##
## Linear Hypotheses:
##                Estimate Std. Error z value Pr(>|z|)
## EL - EF == 0 -1059849     983523  -1.078    0.281
## (Adjusted p values reported -- single-step method)
##
##
## Tukey results for gene: CTSH
##
##   Simultaneous Tests for General Linear Hypotheses
##
## Multiple Comparisons of Means: Tukey Contrasts
##
##
## Fit: glm(formula = Protein_Count ~ Tissue, family = gaussian, data = gene_
data)
##
## Linear Hypotheses:
##                Estimate Std. Error z value Pr(>|z|)
## EL - EF == 0   161743     175021   0.924    0.355
## (Adjusted p values reported -- single-step method)
##
##
## Tukey results for gene: CTSS
##
##   Simultaneous Tests for General Linear Hypotheses
```

```
##
## Multiple Comparisons of Means: Tukey Contrasts
##
##
## Fit: glm(formula = Protein_Count ~ Tissue, family = gaussian, data = gene_
data)
##
## Linear Hypotheses:
##              Estimate Std. Error z value Pr(>|z|)
## EL - EF == 0  -258912     250434  -1.034    0.301
## (Adjusted p values reported -- single-step method)
##
##
## Tukey results for gene: CTSW
##
##   Simultaneous Tests for General Linear Hypotheses
##
## Multiple Comparisons of Means: Tukey Contrasts
##
##
## Fit: glm(formula = Protein_Count ~ Tissue, family = gaussian, data = gene_
data)
##
## Linear Hypotheses:
##              Estimate Std. Error z value Pr(>|z|)
## EL - EF == 0    14037     106975   0.131    0.896
## (Adjusted p values reported -- single-step method)
##
##
## Tukey results for gene: CTSZ
##
##   Simultaneous Tests for General Linear Hypotheses
##
## Multiple Comparisons of Means: Tukey Contrasts
##
##
## Fit: glm(formula = Protein_Count ~ Tissue, family = gaussian, data = gene_
data)
##
## Linear Hypotheses:
##              Estimate Std. Error z value Pr(>|z|)
## EL - EF == 0  -211946     138273  -1.533    0.125
## (Adjusted p values reported -- single-step method)
##
##
## Tukey results for gene: F10
##
##   Simultaneous Tests for General Linear Hypotheses
##
## Multiple Comparisons of Means: Tukey Contrasts
```

```
##
##
## Fit: glm(formula = Protein_Count ~ Tissue, family = gaussian, data = gene_
data)
##
## Linear Hypotheses:
##              Estimate Std. Error z value Pr(>|z|)
## EL - EF == 0   334060     253948   1.315    0.188
## (Adjusted p values reported -- single-step method)
##
##
## Tukey results for gene: F13B
##
##   Simultaneous Tests for General Linear Hypotheses
##
## Multiple Comparisons of Means: Tukey Contrasts
##
##
## Fit: glm(formula = Protein_Count ~ Tissue, family = gaussian, data = gene_
data)
##
## Linear Hypotheses:
##              Estimate Std. Error z value Pr(>|z|)
## EL - EF == 0   -24172     149517  -0.162    0.872
## (Adjusted p values reported -- single-step method)
##
##
## Tukey results for gene: F2
##
##   Simultaneous Tests for General Linear Hypotheses
##
## Multiple Comparisons of Means: Tukey Contrasts
##
##
## Fit: glm(formula = Protein_Count ~ Tissue, family = gaussian, data = gene_
data)
##
## Linear Hypotheses:
##              Estimate Std. Error z value Pr(>|z|)
## EL - EF == 0    25481     251763   0.101    0.919
## (Adjusted p values reported -- single-step method)
##
##
## Tukey results for gene: F9
##
##   Simultaneous Tests for General Linear Hypotheses
##
## Multiple Comparisons of Means: Tukey Contrasts
##
##
```

```
## Fit: glm(formula = Protein_Count ~ Tissue, family = gaussian, data = gene_
data)
##
## Linear Hypotheses:
##              Estimate Std. Error z value Pr(>|z|)
## EL - EF == 0    21031      88492   0.238    0.812
## (Adjusted p values reported -- single-step method)
##
##
## Tukey results for gene: HRG
##
##   Simultaneous Tests for General Linear Hypotheses
##
## Multiple Comparisons of Means: Tukey Contrasts
##
##
## Fit: glm(formula = Protein_Count ~ Tissue, family = gaussian, data = gene_
data)
##
## Linear Hypotheses:
##              Estimate Std. Error z value Pr(>|z|)
## EL - EF == 0  2678392    1697189   1.578    0.115
## (Adjusted p values reported -- single-step method)
##
##
## Tukey results for gene: HTRA1
##
##   Simultaneous Tests for General Linear Hypotheses
##
## Multiple Comparisons of Means: Tukey Contrasts
##
##
## Fit: glm(formula = Protein_Count ~ Tissue, family = gaussian, data = gene_
data)
##
## Linear Hypotheses:
##              Estimate Std. Error z value Pr(>|z|)
## EL - EF == 0   -59304      58197  -1.019    0.308
## (Adjusted p values reported -- single-step method)
##
##
## Tukey results for gene: ITIH1
##
##   Simultaneous Tests for General Linear Hypotheses
##
## Multiple Comparisons of Means: Tukey Contrasts
##
##
## Fit: glm(formula = Protein_Count ~ Tissue, family = gaussian, data = gene_
data)
```

```
##
## Linear Hypotheses:
##                Estimate Std. Error z value Pr(>|z|)
## EL - EF == 0  -498312     303231  -1.643      0.1
## (Adjusted p values reported -- single-step method)
##
##
## Tukey results for gene: ITIH2
##
##   Simultaneous Tests for General Linear Hypotheses
##
## Multiple Comparisons of Means: Tukey Contrasts
##
##
## Fit: glm(formula = Protein_Count ~ Tissue, family = gaussian, data = gene_
data)
##
## Linear Hypotheses:
##                Estimate Std. Error z value Pr(>|z|)
## EL - EF == 0   853732     501350   1.703   0.0886 .
## ---
## Signif. codes:  0 '***' 0.001 '**' 0.01 '*' 0.05 '.' 0.1 ' ' 1
## (Adjusted p values reported -- single-step method)
##
##
## Tukey results for gene: ITIH4
##
##   Simultaneous Tests for General Linear Hypotheses
##
## Multiple Comparisons of Means: Tukey Contrasts
##
##
## Fit: glm(formula = Protein_Count ~ Tissue, family = gaussian, data = gene_
data)
##
## Linear Hypotheses:
##                Estimate Std. Error z value Pr(>|z|)
## EL - EF == 0  -617416    1310877  -0.471    0.638
## (Adjusted p values reported -- single-step method)
##
##
## Tukey results for gene: ITIH5
##
##   Simultaneous Tests for General Linear Hypotheses
##
## Multiple Comparisons of Means: Tukey Contrasts
##
##
## Fit: glm(formula = Protein_Count ~ Tissue, family = gaussian, data = gene_
data)
```

```
##
## Linear Hypotheses:
##                Estimate Std. Error z value Pr(>|z|)
## EL - EF == 0   106897     276176   0.387    0.699
## (Adjusted p values reported -- single-step method)
##
##
## Tukey results for gene: KNG1
##
##   Simultaneous Tests for General Linear Hypotheses
##
## Multiple Comparisons of Means: Tukey Contrasts
##
##
## Fit: glm(formula = Protein_Count ~ Tissue, family = gaussian, data = gene_
data)
##
## Linear Hypotheses:
##                Estimate Std. Error z value Pr(>|z|)
## EL - EF == 0   -76303     110984  -0.688    0.492
## (Adjusted p values reported -- single-step method)
##
##
## Tukey results for gene: KNG2
##
##   Simultaneous Tests for General Linear Hypotheses
##
## Multiple Comparisons of Means: Tukey Contrasts
##
##
## Fit: glm(formula = Protein_Count ~ Tissue, family = gaussian, data = gene_
data)
##
## Linear Hypotheses:
##                Estimate Std. Error z value Pr(>|z|)
## EL - EF == 0  -223511     116576  -1.917   0.0552 .
## ---
## Signif. codes:  0 '***' 0.001 '**' 0.01 '*' 0.05 '.' 0.1 ' ' 1
## (Adjusted p values reported -- single-step method)
##
##
## Tukey results for gene: LOX
##
##   Simultaneous Tests for General Linear Hypotheses
##
## Multiple Comparisons of Means: Tukey Contrasts
##
##
## Fit: glm(formula = Protein_Count ~ Tissue, family = gaussian, data = gene_
data)
```

```
##
## Linear Hypotheses:
##              Estimate Std. Error z value Pr(>|z|)
## EL - EF == 0 -2915994    2452741  -1.189    0.234
## (Adjusted p values reported -- single-step method)
##
##
## Tukey results for gene: LOXL1
##
##   Simultaneous Tests for General Linear Hypotheses
##
## Multiple Comparisons of Means: Tukey Contrasts
##
##
## Fit: glm(formula = Protein_Count ~ Tissue, family = gaussian, data = gene_
data)
##
## Linear Hypotheses:
##              Estimate Std. Error z value Pr(>|z|)
## EL - EF == 0  -351078     322125   -1.09    0.276
## (Adjusted p values reported -- single-step method)
##
##
## Tukey results for gene: MASP2
##
##   Simultaneous Tests for General Linear Hypotheses
##
## Multiple Comparisons of Means: Tukey Contrasts
##
##
## Fit: glm(formula = Protein_Count ~ Tissue, family = gaussian, data = gene_
data)
##
## Linear Hypotheses:
##              Estimate Std. Error z value Pr(>|z|)
## EL - EF == 0   200819     259089   0.775    0.438
## (Adjusted p values reported -- single-step method)
##
##
## Tukey results for gene: MMP14
##
##   Simultaneous Tests for General Linear Hypotheses
##
## Multiple Comparisons of Means: Tukey Contrasts
##
##
## Fit: glm(formula = Protein_Count ~ Tissue, family = gaussian, data = gene_
data)
##
## Linear Hypotheses:
```

```
##               Estimate Std. Error z value Pr(>|z|)
## EL - EF == 0  -168206     207500  -0.811    0.418
## (Adjusted p values reported -- single-step method)
##
##
## Tukey results for gene: MMP2
##
##   Simultaneous Tests for General Linear Hypotheses
##
## Multiple Comparisons of Means: Tukey Contrasts
##
##
## Fit: glm(formula = Protein_Count ~ Tissue, family = gaussian, data = gene_
data)
##
## Linear Hypotheses:
##               Estimate Std. Error z value Pr(>|z|)
## EL - EF == 0    40423     129154   0.313    0.754
## (Adjusted p values reported -- single-step method)
##
##
## Tukey results for gene: MMP3
##
##   Simultaneous Tests for General Linear Hypotheses
##
## Multiple Comparisons of Means: Tukey Contrasts
##
##
## Fit: glm(formula = Protein_Count ~ Tissue, family = gaussian, data = gene_
data)
##
## Linear Hypotheses:
##               Estimate Std. Error z value Pr(>|z|)
## EL - EF == 0 -2093821    1082807  -1.934   0.0532 .
## ---
## Signif. codes:  0 '***' 0.001 '**' 0.01 '*' 0.05 '.' 0.1 ' ' 1
## (Adjusted p values reported -- single-step method)
##
##
## Tukey results for gene: P4HA1
##
##   Simultaneous Tests for General Linear Hypotheses
##
## Multiple Comparisons of Means: Tukey Contrasts
##
##
## Fit: glm(formula = Protein_Count ~ Tissue, family = gaussian, data = gene_
data)
##
## Linear Hypotheses:
```

```
##               Estimate Std. Error z value Pr(>|z|)
## EL - EF == 0   222451     166325   1.337    0.181
## (Adjusted p values reported -- single-step method)
##
##
## Tukey results for gene: P4HA2
##
##   Simultaneous Tests for General Linear Hypotheses
##
## Multiple Comparisons of Means: Tukey Contrasts
##
##
## Fit: glm(formula = Protein_Count ~ Tissue, family = gaussian, data = gene_
data)
##
## Linear Hypotheses:
##               Estimate Std. Error z value Pr(>|z|)
## EL - EF == 0   -69883     193648  -0.361    0.718
## (Adjusted p values reported -- single-step method)
##
##
## Tukey results for gene: PLG
##
##   Simultaneous Tests for General Linear Hypotheses
##
## Multiple Comparisons of Means: Tukey Contrasts
##
##
## Fit: glm(formula = Protein_Count ~ Tissue, family = gaussian, data = gene_
data)
##
## Linear Hypotheses:
##               Estimate Std. Error z value Pr(>|z|)
## EL - EF == 0  -325546     981715  -0.332     0.74
## (Adjusted p values reported -- single-step method)
##
##
## Tukey results for gene: PLOD1
##
##   Simultaneous Tests for General Linear Hypotheses
##
## Multiple Comparisons of Means: Tukey Contrasts
##
##
## Fit: glm(formula = Protein_Count ~ Tissue, family = gaussian, data = gene_
data)
##
## Linear Hypotheses:
##               Estimate Std. Error z value Pr(>|z|)
## EL - EF == 0    14930     129350   0.115    0.908
```

```
## (Adjusted p values reported -- single-step method)
##
##
## Tukey results for gene: PLOD3
##
##   Simultaneous Tests for General Linear Hypotheses
##
## Multiple Comparisons of Means: Tukey Contrasts
##
##
## Fit: glm(formula = Protein_Count ~ Tissue, family = gaussian, data = gene_
data)
##
## Linear Hypotheses:
##              Estimate Std. Error z value Pr(>|z|)
## EL - EF == 0    26871     278794   0.096    0.923
## (Adjusted p values reported -- single-step method)
##
##
## Tukey results for gene: PRSS1
##
##   Simultaneous Tests for General Linear Hypotheses
##
## Multiple Comparisons of Means: Tukey Contrasts
##
##
## Fit: glm(formula = Protein_Count ~ Tissue, family = gaussian, data = gene_
data)
##
## Linear Hypotheses:
##              Estimate Std. Error z value Pr(>|z|)
## EL - EF == 0 -2866984    3494102  -0.821    0.412
## (Adjusted p values reported -- single-step method)
##
##
## Tukey results for gene: SERPINA1
##
##   Simultaneous Tests for General Linear Hypotheses
##
## Multiple Comparisons of Means: Tukey Contrasts
##
##
## Fit: glm(formula = Protein_Count ~ Tissue, family = gaussian, data = gene_
data)
##
## Linear Hypotheses:
##              Estimate Std. Error z value Pr(>|z|)
## EL - EF == 0   -15639      90989  -0.172    0.864
## (Adjusted p values reported -- single-step method)
##
```

```
##
## Tukey results for gene: SERPINA10
##
##   Simultaneous Tests for General Linear Hypotheses
##
## Multiple Comparisons of Means: Tukey Contrasts
##
##
## Fit: glm(formula = Protein_Count ~ Tissue, family = gaussian, data = gene_
data)
##
## Linear Hypotheses:
##              Estimate Std. Error z value Pr(>|z|)
## EL - EF == 0   -34806     102628  -0.339    0.734
## (Adjusted p values reported -- single-step method)
##
##
## Tukey results for gene: SERPINA3-8
##
##   Simultaneous Tests for General Linear Hypotheses
##
## Multiple Comparisons of Means: Tukey Contrasts
##
##
## Fit: glm(formula = Protein_Count ~ Tissue, family = gaussian, data = gene_
data)
##
## Linear Hypotheses:
##              Estimate Std. Error z value Pr(>|z|)
## EL - EF == 0    10023     101540   0.099    0.921
## (Adjusted p values reported -- single-step method)
##
##
## Tukey results for gene: SERPINA5
##
##   Simultaneous Tests for General Linear Hypotheses
##
## Multiple Comparisons of Means: Tukey Contrasts
##
##
## Fit: glm(formula = Protein_Count ~ Tissue, family = gaussian, data = gene_
data)
##
## Linear Hypotheses:
##              Estimate Std. Error z value Pr(>|z|)
## EL - EF == 0    66527     117641   0.566    0.572
## (Adjusted p values reported -- single-step method)
##
##
## Tukey results for gene: SERPINC1
```

```
##
##   Simultaneous Tests for General Linear Hypotheses
##
## Multiple Comparisons of Means: Tukey Contrasts
##
##
## Fit: glm(formula = Protein_Count ~ Tissue, family = gaussian, data = gene_
data)
##
## Linear Hypotheses:
##              Estimate Std. Error z value Pr(>|z|)
## EL - EF == 0  -608833     450335  -1.352    0.176
## (Adjusted p values reported -- single-step method)
##
##
## Tukey results for gene: SERPIND1
##
##   Simultaneous Tests for General Linear Hypotheses
##
## Multiple Comparisons of Means: Tukey Contrasts
##
##
## Fit: glm(formula = Protein_Count ~ Tissue, family = gaussian, data = gene_
data)
##
## Linear Hypotheses:
##              Estimate Std. Error z value Pr(>|z|)
## EL - EF == 0     6667     110816    0.06    0.952
## (Adjusted p values reported -- single-step method)
##
##
## Tukey results for gene: SERPINE2
##
##   Simultaneous Tests for General Linear Hypotheses
##
## Multiple Comparisons of Means: Tukey Contrasts
##
##
## Fit: glm(formula = Protein_Count ~ Tissue, family = gaussian, data = gene_
data)
##
## Linear Hypotheses:
##              Estimate Std. Error z value Pr(>|z|)
## EL - EF == 0  -158203     137846  -1.148    0.251
## (Adjusted p values reported -- single-step method)
##
##
## Tukey results for gene: SERPINF1
##
##   Simultaneous Tests for General Linear Hypotheses
```

```
##
## Multiple Comparisons of Means: Tukey Contrasts
##
##
## Fit: glm(formula = Protein_Count ~ Tissue, family = gaussian, data = gene_
data)
##
## Linear Hypotheses:
##              Estimate Std. Error z value Pr(>|z|)
## EL - EF == 0   181178     557345   0.325    0.745
## (Adjusted p values reported -- single-step method)
##
##
## Tukey results for gene: SERPINF2
##
##   Simultaneous Tests for General Linear Hypotheses
##
## Multiple Comparisons of Means: Tukey Contrasts
##
##
## Fit: glm(formula = Protein_Count ~ Tissue, family = gaussian, data = gene_
data)
##
## Linear Hypotheses:
##              Estimate Std. Error z value Pr(>|z|)
## EL - EF == 0   413900     722652   0.573    0.567
## (Adjusted p values reported -- single-step method)
##
##
## Tukey results for gene: SERPING1
##
##   Simultaneous Tests for General Linear Hypotheses
##
## Multiple Comparisons of Means: Tukey Contrasts
##
##
## Fit: glm(formula = Protein_Count ~ Tissue, family = gaussian, data = gene_
data)
##
## Linear Hypotheses:
##              Estimate Std. Error z value Pr(>|z|)
## EL - EF == 0   159727     177664   0.899    0.369
## (Adjusted p values reported -- single-step method)
##
##
## Tukey results for gene: SERPINH1
##
##   Simultaneous Tests for General Linear Hypotheses
##
## Multiple Comparisons of Means: Tukey Contrasts
```

```
##
##
## Fit: glm(formula = Protein_Count ~ Tissue, family = gaussian, data = gene_
data)
##
## Linear Hypotheses:
##              Estimate Std. Error z value Pr(>|z|)
## EL - EF == 0 -4188914    7692187  -0.545    0.586
## (Adjusted p values reported -- single-step method)
##
##
## Tukey results for gene: TGM1
##
##   Simultaneous Tests for General Linear Hypotheses
##
## Multiple Comparisons of Means: Tukey Contrasts
##
##
## Fit: glm(formula = Protein_Count ~ Tissue, family = gaussian, data = gene_
data)
##
## Linear Hypotheses:
##              Estimate Std. Error z value Pr(>|z|)
## EL - EF == 0  -138966     507455  -0.274    0.784
## (Adjusted p values reported -- single-step method)
##
##
## Tukey results for gene: TGM2
##
##   Simultaneous Tests for General Linear Hypotheses
##
## Multiple Comparisons of Means: Tukey Contrasts
##
##
## Fit: glm(formula = Protein_Count ~ Tissue, family = gaussian, data = gene_
data)
##
## Linear Hypotheses:
##               Estimate Std. Error z value Pr(>|z|)
## EL - EF == 0 -14016900   44499665  -0.315    0.753
## (Adjusted p values reported -- single-step method)
##
##
## Tukey results for gene: TIMP2
##
##   Simultaneous Tests for General Linear Hypotheses
##
## Multiple Comparisons of Means: Tukey Contrasts
##
##
```

```
## Fit: glm(formula = Protein_Count ~ Tissue, family = gaussian, data = gene_
data)
##
## Linear Hypotheses:
##              Estimate Std. Error z value Pr(>|z|)
## EL - EF == 0    36346     103376   0.352    0.725
## (Adjusted p values reported -- single-step method)

#### ECM-affiliated

# Read your cleaned joined data file into a data frame
cleaned_joined_data <- read.csv("cleaned_matrisome_endo.csv")

# Filter rows with Matrisome_Category "ECM-affiliated"
ECMaffiliated_data <- cleaned_joined_data %>%
  filter(Matrisome_Category == "ECM-affiliated")

# Reshape the data to have separate columns for each replicate and each tissu
e stage
organized_data <- ECMaffiliated_data %>%
  pivot_longer(cols = starts_with("EF") | starts_with("EL"),
               names_to = c("Tissue", "Stage"),
               names_sep = "\\.",
               values_to = "Protein_Count")

## Warning: Expected 2 pieces. Missing pieces filled with `NA` in 2 rows [1,
6].

# Save the reshaped data to a new file

write.csv(organized_data, "ECM-affiliated_data_matrisome_endo_to_clean.csv",
row.names = FALSE)

data2 <- read.csv("ECM-affiliated_data_matrisome_endo_clean.csv")

png(filename="endo_ECM-affiliated.png",
    width = 12, height = 6, units = "in",
    res = 600)

boxplot(Protein_Count ~ Tissue*Genes,
        data = data2,
        cex.axis = 0.8,
        col = viridis(2, alpha = 0.7),
        lwd = 0.7,
        ylab = "",
        ylim = c(0, 40000000),
        axes = FALSE,
        frame.plot = TRUE,
        xlab = "")
```

```
GROUP <- unique(data2$Genes)
groups <- 20
numbox <- 2
total <- groups * numbox
xpoints <- seq(median(1:numbox), total, numbox)

axis(2, seq(0, 40000000, by = 1000000), las = 1, cex.axis = 0.7)
axis(1, labels = GROUP, at = xpoints, las = 2, cex.axis = 0.7)

stripchart(data2$Protein_Count ~ data2$Tissue*data2$Genes,
           method = "jitter",
           pch = 19,
           col = "black",
           vertical = TRUE,
           add = TRUE,
           cex = 0.3)

title(ylab = expression("Protein abundance (RU)"), line = 3.0)
legend("top", fill = viridis(2), legend = c("Endometrium Follicular", "Endome
trium Luteal"), horiz = T, cex = 0.7)

dev.off()

## quartz_off_screen
##                 2

# Convert Tissue to factor
data2$Tissue <- as.factor(data2$Tissue)

# Create an empty list to store Tukey results
tukey_results_list <- list()

# Unique genes
unique_genes <- unique(data2$Genes)

# Loop over each gene
for (gene in unique_genes) {
  # Subset data for the current gene
  gene_data <- subset(data2, Genes == gene)

  # Fit the generalized linear model
  glm_model <- glm(Protein_Count ~ Tissue, data = gene_data, family = gaussia
n)

  # Perform Tukey test
  contrast_matrix <- glht(glm_model, linfct = mcp(Tissue = "Tukey"))
  tukey_results <- summary(contrast_matrix)

  # Store the results
```

```
  tukey_results_list[[gene]] <- tukey_results
}

# Print the results for each gene
for (gene in unique_genes) {
  cat("Tukey results for gene:", gene, "\n")
  print(tukey_results_list[[gene]])
  cat("\n")
}

## Tukey results for gene: ANXA1
##
##   Simultaneous Tests for General Linear Hypotheses
##
## Multiple Comparisons of Means: Tukey Contrasts
##
##
## Fit: glm(formula = Protein_Count ~ Tissue, family = gaussian, data = gene_
data)
##
## Linear Hypotheses:
##                Estimate Std. Error z value Pr(>|z|)
## EL - EF == 0  -130274    1621287   -0.08    0.936
## (Adjusted p values reported -- single-step method)
##
##
## Tukey results for gene: ANXA11
##
##   Simultaneous Tests for General Linear Hypotheses
##
## Multiple Comparisons of Means: Tukey Contrasts
##
##
## Fit: glm(formula = Protein_Count ~ Tissue, family = gaussian, data = gene_
data)
##
## Linear Hypotheses:
##                Estimate Std. Error z value Pr(>|z|)
## EL - EF == 0   100141     245672   0.408    0.684
## (Adjusted p values reported -- single-step method)
##
##
## Tukey results for gene: ANXA2
##
##   Simultaneous Tests for General Linear Hypotheses
##
## Multiple Comparisons of Means: Tukey Contrasts
##
##
## Fit: glm(formula = Protein_Count ~ Tissue, family = gaussian, data = gene_
```

```
data)
##
## Linear Hypotheses:
##              Estimate Std. Error z value Pr(>|z|)
## EL - EF == 0  2759436    8145705   0.339    0.735
## (Adjusted p values reported -- single-step method)
##
##
## Tukey results for gene: ANXA3
##
##   Simultaneous Tests for General Linear Hypotheses
##
## Multiple Comparisons of Means: Tukey Contrasts
##
##
## Fit: glm(formula = Protein_Count ~ Tissue, family = gaussian, data = gene_
data)
##
## Linear Hypotheses:
##              Estimate Std. Error z value Pr(>|z|)
## EL - EF == 0   749127     994976   0.753    0.452
## (Adjusted p values reported -- single-step method)
##
##
## Tukey results for gene: ANXA4
##
##   Simultaneous Tests for General Linear Hypotheses
##
## Multiple Comparisons of Means: Tukey Contrasts
##
##
## Fit: glm(formula = Protein_Count ~ Tissue, family = gaussian, data = gene_
data)
##
## Linear Hypotheses:
##              Estimate Std. Error z value Pr(>|z|)
## EL - EF == 0  -686217     592106  -1.159    0.246
## (Adjusted p values reported -- single-step method)
##
##
## Tukey results for gene: ANXA5
##
##   Simultaneous Tests for General Linear Hypotheses
##
## Multiple Comparisons of Means: Tukey Contrasts
##
##
## Fit: glm(formula = Protein_Count ~ Tissue, family = gaussian, data = gene_
data)
##
```

```
## Linear Hypotheses:
##              Estimate Std. Error z value Pr(>|z|)
## EL - EF == 0  -268492    2471715  -0.109    0.913
## (Adjusted p values reported -- single-step method)
##
##
## Tukey results for gene: ANXA6
##
##   Simultaneous Tests for General Linear Hypotheses
##
## Multiple Comparisons of Means: Tukey Contrasts
##
##
## Fit: glm(formula = Protein_Count ~ Tissue, family = gaussian, data = gene_
data)
##
## Linear Hypotheses:
##              Estimate Std. Error z value Pr(>|z|)
## EL - EF == 0   542299     533530   1.016    0.309
## (Adjusted p values reported -- single-step method)
##
##
## Tukey results for gene: ANXA7
##
##   Simultaneous Tests for General Linear Hypotheses
##
## Multiple Comparisons of Means: Tukey Contrasts
##
##
## Fit: glm(formula = Protein_Count ~ Tissue, family = gaussian, data = gene_
data)
##
## Linear Hypotheses:
##              Estimate Std. Error z value Pr(>|z|)
## EL - EF == 0   123528     467172   0.264    0.791
## (Adjusted p values reported -- single-step method)
##
##
## Tukey results for gene: C1QC
##
##   Simultaneous Tests for General Linear Hypotheses
##
## Multiple Comparisons of Means: Tukey Contrasts
##
##
## Fit: glm(formula = Protein_Count ~ Tissue, family = gaussian, data = gene_
data)
##
## Linear Hypotheses:
##              Estimate Std. Error z value Pr(>|z|)
```

```
## EL - EF == 0    37800      72216   0.523    0.601
## (Adjusted p values reported -- single-step method)
##
##
## Tukey results for gene: CLEC11A
##
##   Simultaneous Tests for General Linear Hypotheses
##
## Multiple Comparisons of Means: Tukey Contrasts
##
##
## Fit: glm(formula = Protein_Count ~ Tissue, family = gaussian, data = gene_
data)
##
## Linear Hypotheses:
##              Estimate Std. Error z value Pr(>|z|)
## EL - EF == 0   -49120     167576  -0.293    0.769
## (Adjusted p values reported -- single-step method)
##
##
## Tukey results for gene: CLEC3B
##
##   Simultaneous Tests for General Linear Hypotheses
##
## Multiple Comparisons of Means: Tukey Contrasts
##
##
## Fit: glm(formula = Protein_Count ~ Tissue, family = gaussian, data = gene_
data)
##
## Linear Hypotheses:
##              Estimate Std. Error z value Pr(>|z|)
## EL - EF == 0   -29664     784331  -0.038     0.97
## (Adjusted p values reported -- single-step method)
##
##
## Tukey results for gene: COLEC12
##
##   Simultaneous Tests for General Linear Hypotheses
##
## Multiple Comparisons of Means: Tukey Contrasts
##
##
## Fit: glm(formula = Protein_Count ~ Tissue, family = gaussian, data = gene_
data)
##
## Linear Hypotheses:
##              Estimate Std. Error z value Pr(>|z|)
## EL - EF == 0  -112857     114444  -0.986    0.324
## (Adjusted p values reported -- single-step method)
```

```
##
##
## Tukey results for gene: CSPG4
##
##   Simultaneous Tests for General Linear Hypotheses
##
## Multiple Comparisons of Means: Tukey Contrasts
##
##
## Fit: glm(formula = Protein_Count ~ Tissue, family = gaussian, data = gene_
data)
##
## Linear Hypotheses:
##              Estimate Std. Error z value Pr(>|z|)
## EL - EF == 0   212836     250858   0.848    0.396
## (Adjusted p values reported -- single-step method)
##
##
## Tukey results for gene: HPX
##
##   Simultaneous Tests for General Linear Hypotheses
##
## Multiple Comparisons of Means: Tukey Contrasts
##
##
## Fit: glm(formula = Protein_Count ~ Tissue, family = gaussian, data = gene_
data)
##
## Linear Hypotheses:
##              Estimate Std. Error z value Pr(>|z|)
## EL - EF == 0   586871     385163   1.524    0.128
## (Adjusted p values reported -- single-step method)
##
##
## Tukey results for gene: LGALS1
##
##   Simultaneous Tests for General Linear Hypotheses
##
## Multiple Comparisons of Means: Tukey Contrasts
##
##
## Fit: glm(formula = Protein_Count ~ Tissue, family = gaussian, data = gene_
data)
##
## Linear Hypotheses:
##              Estimate Std. Error z value Pr(>|z|)
## EL - EF == 0  -883222     597869  -1.477     0.14
## (Adjusted p values reported -- single-step method)
##
##
```

```
## Tukey results for gene: LGALS7
##
##   Simultaneous Tests for General Linear Hypotheses
##
## Multiple Comparisons of Means: Tukey Contrasts
##
##
## Fit: glm(formula = Protein_Count ~ Tissue, family = gaussian, data = gene_
data)
##
## Linear Hypotheses:
##               Estimate Std. Error z value Pr(>|z|)
## EL - EF == 0   -88172     112626  -0.783    0.434
## (Adjusted p values reported -- single-step method)
##
##
## Tukey results for gene: LGALS9
##
##   Simultaneous Tests for General Linear Hypotheses
##
## Multiple Comparisons of Means: Tukey Contrasts
##
##
## Fit: glm(formula = Protein_Count ~ Tissue, family = gaussian, data = gene_
data)
##
## Linear Hypotheses:
##               Estimate Std. Error z value Pr(>|z|)
## EL - EF == 0  -141621     349453  -0.405    0.685
## (Adjusted p values reported -- single-step method)
##
##
## Tukey results for gene: LMAN1
##
##   Simultaneous Tests for General Linear Hypotheses
##
## Multiple Comparisons of Means: Tukey Contrasts
##
##
## Fit: glm(formula = Protein_Count ~ Tissue, family = gaussian, data = gene_
data)
##
## Linear Hypotheses:
##               Estimate Std. Error z value Pr(>|z|)
## EL - EF == 0   -13481     102317  -0.132    0.895
## (Adjusted p values reported -- single-step method)
##
##
## Tukey results for gene: PLXNB1
##
```

```
##   Simultaneous Tests for General Linear Hypotheses
##
## Multiple Comparisons of Means: Tukey Contrasts
##
##
## Fit: glm(formula = Protein_Count ~ Tissue, family = gaussian, data = gene_
data)
##
## Linear Hypotheses:
##              Estimate Std. Error z value Pr(>|z|)
## EL - EF == 0    23315     112971   0.206    0.836
## (Adjusted p values reported -- single-step method)
##
##
## Tukey results for gene: PLXNB2
##
##   Simultaneous Tests for General Linear Hypotheses
##
## Multiple Comparisons of Means: Tukey Contrasts
##
##
## Fit: glm(formula = Protein_Count ~ Tissue, family = gaussian, data = gene_
data)
##
## Linear Hypotheses:
##              Estimate Std. Error z value Pr(>|z|)
## EL - EF == 0  -308181     299690  -1.028    0.304
## (Adjusted p values reported -- single-step method)
```

```r
#### Secreted Factors

# Read your cleaned joined data file into a data frame
cleaned_joined_data <- read.csv("cleaned_matrisome_endo.csv")

# Filter rows with Matrisome_Category "Secreted Factors"
Secreted_Factors_data <- cleaned_joined_data %>%
  filter(Matrisome_Category == "Secreted Factors")

# Reshape the data to have separate columns for each replicate and each tissu
e stage
organized_data <- Secreted_Factors_data %>%
  pivot_longer(cols = starts_with("EF") | starts_with("EL"),
               names_to = c("Tissue", "Stage"),
               names_sep = "\\.",
               values_to = "Protein_Count")
```

```
## Warning: Expected 2 pieces. Missing pieces filled with `NA` in 2 rows [1,
6].
```

```
# Save the reshaped data to a new file

write.csv(organized_data, "Secreted_factors_data_matrisome_endo_to_clean.csv"
, row.names = FALSE)

data2 <- read.csv("Secreted_factors_data_matrisome_endo_clean.csv")

png(filename="endo_Secreted_factors.png",
    width = 12, height = 6, units = "in",
    res = 600)

boxplot(Protein_Count ~ Tissue*Genes,
        data = data2,
        cex.axis = 0.8,
        col = viridis(2, alpha = 0.7),
        lwd = 0.7,
        ylab = "",
        ylim = c(0, 45000000),
        axes = FALSE,
        frame.plot = TRUE,
        xlab = "")

GROUP <- unique(data2$Genes)
groups <- 15
numbox <- 2
total <- groups * numbox
xpoints <- seq(median(1:numbox), total, numbox)

axis(2, seq(0, 45000000, by = 10000000), las = 1, cex.axis = 0.7)
axis(1, labels = GROUP, at = xpoints, las = 2, cex.axis = 0.7)

stripchart(data2$Protein_Count ~ data2$Tissue*data2$Genes,
           method = "jitter",
           pch = 19,
           col = "black",
           vertical = TRUE,
           add = TRUE,
           cex = 0.3)

title(ylab = expression("Protein abundance (RU)"), line = 3.0)
legend("top", fill = viridis(2), legend = c("Endometrium Follicular", "Endome
trium Luteal"), horiz = T, cex = 0.7)

dev.off()

## quartz_off_screen
##                 2
```

```
# Convert Tissue to factor
data2$Tissue <- as.factor(data2$Tissue)

# Create an empty list to store Tukey results
tukey_results_list <- list()

# Unique genes
unique_genes <- unique(data2$Genes)

# Loop over each gene
for (gene in unique_genes) {
  # Subset data for the current gene
  gene_data <- subset(data2, Genes == gene)

  # Fit the generalized linear model
  glm_model <- glm(Protein_Count ~ Tissue, data = gene_data, family = gaussia
n)

  # Perform Tukey test
  contrast_matrix <- glht(glm_model, linfct = mcp(Tissue = "Tukey"))
  tukey_results <- summary(contrast_matrix)

  # Store the results
  tukey_results_list[[gene]] <- tukey_results
}

# Print the results for each gene
for (gene in unique_genes) {
  cat("Tukey results for gene:", gene, "\n")
  print(tukey_results_list[[gene]])
  cat("\n")
}

## Tukey results for gene: ANGPT4
##
##   Simultaneous Tests for General Linear Hypotheses
##
## Multiple Comparisons of Means: Tukey Contrasts
##
##
## Fit: glm(formula = Protein_Count ~ Tissue, family = gaussian, data = gene_
data)
##
## Linear Hypotheses:
##              Estimate Std. Error z value Pr(>|z|)
## EL - EF == 0  -205103     163265  -1.256    0.209
## (Adjusted p values reported -- single-step method)
##
##
```

```
## Tukey results for gene: ANGPTL2
##
##   Simultaneous Tests for General Linear Hypotheses
##
## Multiple Comparisons of Means: Tukey Contrasts
##
##
## Fit: glm(formula = Protein_Count ~ Tissue, family = gaussian, data = gene_
data)
##
## Linear Hypotheses:
##              Estimate Std. Error z value Pr(>|z|)
## EL - EF == 0  -163425     270017  -0.605    0.545
## (Adjusted p values reported -- single-step method)
##
##
## Tukey results for gene: CHRD
##
##   Simultaneous Tests for General Linear Hypotheses
##
## Multiple Comparisons of Means: Tukey Contrasts
##
##
## Fit: glm(formula = Protein_Count ~ Tissue, family = gaussian, data = gene_
data)
##
## Linear Hypotheses:
##              Estimate Std. Error z value Pr(>|z|)
## EL - EF == 0   105815     120350   0.879    0.379
## (Adjusted p values reported -- single-step method)
##
##
## Tukey results for gene: GDF6
##
##   Simultaneous Tests for General Linear Hypotheses
##
## Multiple Comparisons of Means: Tukey Contrasts
##
##
## Fit: glm(formula = Protein_Count ~ Tissue, family = gaussian, data = gene_
data)
##
## Linear Hypotheses:
##              Estimate Std. Error z value Pr(>|z|)
## EL - EF == 0   -51026      34181  -1.493    0.135
## (Adjusted p values reported -- single-step method)
##
##
## Tukey results for gene: GDF7
##
```

```
##   Simultaneous Tests for General Linear Hypotheses
##
## Multiple Comparisons of Means: Tukey Contrasts
##
##
## Fit: glm(formula = Protein_Count ~ Tissue, family = gaussian, data = gene_
data)
##
## Linear Hypotheses:
##              Estimate Std. Error z value Pr(>|z|)
## EL - EF == 0  -138251     148263  -0.932    0.351
## (Adjusted p values reported -- single-step method)
##
##
## Tukey results for gene: HCFC1
##
##   Simultaneous Tests for General Linear Hypotheses
##
## Multiple Comparisons of Means: Tukey Contrasts
##
##
## Fit: glm(formula = Protein_Count ~ Tissue, family = gaussian, data = gene_
data)
##
## Linear Hypotheses:
##              Estimate Std. Error z value Pr(>|z|)
## EL - EF == 0   116783     124206    0.94    0.347
## (Adjusted p values reported -- single-step method)
##
##
## Tukey results for gene: IGF2
##
##   Simultaneous Tests for General Linear Hypotheses
##
## Multiple Comparisons of Means: Tukey Contrasts
##
##
## Fit: glm(formula = Protein_Count ~ Tissue, family = gaussian, data = gene_
data)
##
## Linear Hypotheses:
##              Estimate Std. Error z value Pr(>|z|)
## EL - EF == 0  -234947      95248  -2.467   0.0136 *
## ---
## Signif. codes:  0 '***' 0.001 '**' 0.01 '*' 0.05 '.' 0.1 ' ' 1
## (Adjusted p values reported -- single-step method)
##
##
## Tukey results for gene: IL17D
##
```

```
##   Simultaneous Tests for General Linear Hypotheses
##
## Multiple Comparisons of Means: Tukey Contrasts
##
##
## Fit: glm(formula = Protein_Count ~ Tissue, family = gaussian, data = gene_
data)
##
## Linear Hypotheses:
##              Estimate Std. Error z value Pr(>|z|)
## EL - EF == 0  -214431     197480  -1.086    0.278
## (Adjusted p values reported -- single-step method)
##
##
## Tukey results for gene: MDK
##
##   Simultaneous Tests for General Linear Hypotheses
##
## Multiple Comparisons of Means: Tukey Contrasts
##
##
## Fit: glm(formula = Protein_Count ~ Tissue, family = gaussian, data = gene_
data)
##
## Linear Hypotheses:
##              Estimate Std. Error z value Pr(>|z|)
## EL - EF == 0    52573     111864    0.47    0.638
## (Adjusted p values reported -- single-step method)
##
##
## Tukey results for gene: S100A10
##
##   Simultaneous Tests for General Linear Hypotheses
##
## Multiple Comparisons of Means: Tukey Contrasts
##
##
## Fit: glm(formula = Protein_Count ~ Tissue, family = gaussian, data = gene_
data)
##
## Linear Hypotheses:
##              Estimate Std. Error z value Pr(>|z|)
## EL - EF == 0  -112225     272094  -0.412     0.68
## (Adjusted p values reported -- single-step method)
##
##
## Tukey results for gene: S100A11
##
##   Simultaneous Tests for General Linear Hypotheses
##
```

```
## Multiple Comparisons of Means: Tukey Contrasts
##
##
## Fit: glm(formula = Protein_Count ~ Tissue, family = gaussian, data = gene_
data)
##
## Linear Hypotheses:
##               Estimate Std. Error z value Pr(>|z|)
## EL - EF == 0  -870868    2289167   -0.38    0.704
## (Adjusted p values reported -- single-step method)
##
##
## Tukey results for gene: S100A13
##
##   Simultaneous Tests for General Linear Hypotheses
##
## Multiple Comparisons of Means: Tukey Contrasts
##
##
## Fit: glm(formula = Protein_Count ~ Tissue, family = gaussian, data = gene_
data)
##
## Linear Hypotheses:
##               Estimate Std. Error z value Pr(>|z|)
## EL - EF == 0   120860     131106   0.922    0.357
## (Adjusted p values reported -- single-step method)
##
##
## Tukey results for gene: S100A14
##
##   Simultaneous Tests for General Linear Hypotheses
##
## Multiple Comparisons of Means: Tukey Contrasts
##
##
## Fit: glm(formula = Protein_Count ~ Tissue, family = gaussian, data = gene_
data)
##
## Linear Hypotheses:
##               Estimate Std. Error z value Pr(>|z|)
## EL - EF == 0    69386     128930   0.538     0.59
## (Adjusted p values reported -- single-step method)
##
##
## Tukey results for gene: S100A4
##
##   Simultaneous Tests for General Linear Hypotheses
##
## Multiple Comparisons of Means: Tukey Contrasts
##
```

```
##
## Fit: glm(formula = Protein_Count ~ Tissue, family = gaussian, data = gene_
data)
##
## Linear Hypotheses:
##                  Estimate Std. Error z value Pr(>|z|)
## EL - EF == 0 -10395800    6408853  -1.622    0.105
## (Adjusted p values reported -- single-step method)
##
##
## Tukey results for gene: TGFB1
##
##   Simultaneous Tests for General Linear Hypotheses
##
## Multiple Comparisons of Means: Tukey Contrasts
##
##
## Fit: glm(formula = Protein_Count ~ Tissue, family = gaussian, data = gene_
data)
##
## Linear Hypotheses:
##                Estimate Std. Error z value Pr(>|z|)
## EL - EF == 0   -20534      42871  -0.479    0.632
## (Adjusted p values reported -- single-step method)

#### Components proportions

data <- read.csv("endo_matrisome_final.csv")

# Subset the data for each tissue (EF and EL) separately
data_EF <- subset(data, Tissue == "EF")
data_EL <- subset(data, Tissue == "EL")

# Function to calculate proportions
calculate_proportions <- function(data_subset) {
  proportions <- data_subset %>%
    group_by(Matrisome_Category) %>%
    summarize(Proportion = sum(Protein_Count) / sum(data_subset$Protein_Count
) * 100)
  return(proportions)
}

# Calculate proportions for EF
proportions_EF <- calculate_proportions(data_EF)

# Calculate proportions for EL
proportions_EL <- calculate_proportions(data_EL)
```

```r
# Print proportions for EF
cat("Proportions for Matrisome categories in tissue EF:\n")

## Proportions for Matrisome categories in tissue EF:

print(proportions_EF)

## # A tibble: 6 × 2
##   Matrisome_Category Proportion
##   <chr>                   <dbl>
## 1 Collagens              81.8
## 2 ECM Glycoproteins      10.4
## 3 ECM Regulators          3.34
## 4 ECM-affiliated          0.673
## 5 Proteoglycans           3.10
## 6 Secreted Factors        0.604

# Print proportions for EL
cat("Proportions for Matrisome categories in tissue EL:\n")

## Proportions for Matrisome categories in tissue EL:

print(proportions_EL)

## # A tibble: 6 × 2
##   Matrisome_Category Proportion
##   <chr>                   <dbl>
## 1 Collagens              80.0
## 2 ECM Glycoproteins      10.3
## 3 ECM Regulators          4.08
## 4 ECM-affiliated          0.987
## 5 Proteoglycans           4.03
## 6 Secreted Factors        0.603

###Pie charts

# Define a function to create a pie chart with values and save it
create_and_save_pie_chart <- function(proportions, title, filename) {
  # Create a color palette using Viridis
  color_palette <- viridis(length(unique(proportions$Matrisome_Category)))

  # Create a pie chart
  pie_chart <- ggplot(proportions, aes(x = "", y = Proportion, fill = Matriso
me_Category)) +
    geom_bar(stat = "identity") +
    coord_polar("y", start = 1) +
    geom_text(aes(label = sprintf("%.1f%%", Proportion)), position = position
_stack(vjust = 0.5), color = "white", size = 6) +
    labs(title = title, x = NULL, y = NULL, fill = "Matrisome Category") +
    scale_fill_manual(values = color_palette) +  # Set the color palette
    theme_void() +
```

```
    theme(legend.position = "right")

  # Save the pie chart
  ggsave(filename, plot = pie_chart, width = 6, height = 6, units = "in", dpi
= 300)
}

# Create and save a pie chart for EF
create_and_save_pie_chart(proportions_EF, "Matrisome Categories Proportion in
Tissue EF", "pie_chart_EF.png")

# Create and save a pie chart for EL
create_and_save_pie_chart(proportions_EL, "Matrisome Categories Proportion in
Tissue EL", "pie_chart_EL.png")

#####Summary data

data <- read.csv("endo_matrisome_final.csv")

# Calculate mean and standard deviation for each Gene considering Tissue and
Matrisome_Category
gene_stats <- data %>%
  group_by(Genes, Tissue, Matrisome_Category) %>%
  summarize(
    Mean_Protein_Count = mean(Protein_Count),
    SD_Protein_Count = sd(Protein_Count)
  )

## `summarise()` has grouped output by 'Genes', 'Tissue'. You can override us
ing
## the `.groups` argument.

# Print the results
print(gene_stats)

## # A tibble: 384 × 5
## # Groups:   Genes, Tissue [384]
##    Genes    Tissue Matrisome_Category Mean_Protein_Count SD_Protein_Count
##    <chr>    <chr>  <chr>                           <dbl>            <dbl>
##  1 A2M      EF     ECM Regulators                4069618         2380194.
##  2 A2M      EL     ECM Regulators                3505148         1653646.
##  3 ACAN     EF     Proteoglycans                  215747.          196424.
##  4 ACAN     EL     Proteoglycans                  209617.          300420.
##  5 ADAM10   EF     ECM Regulators                 440078           603742.
##  6 ADAM10   EL     ECM Regulators                 340597.          335792.
##  7 ADAMTS5  EF     ECM Regulators                 537532           560520.
##  8 ADAMTS5  EL     ECM Regulators                 297736.          120777.
##  9 ADAMTSL5 EF     ECM Regulators                 627680.          669812.
## 10 ADAMTSL5 EL     ECM Regulators                 365962.          374114.
## # ℹ 374 more rows
```

```r
# Save the summarized data to a CSV file
write.csv(gene_stats, file = "CEndo_Summary.csv", row.names = FALSE)




####CORTEX

# Read your data into a data frame or data table
data <- read.csv("Cortex_all_count.csv")

# Remove rows with more than 5 NA values
clean_data <- data %>%
  filter(rowSums(is.na(.)) <= 5)

# Replace remaining NA values with 0 in all columns
clean_data <- clean_data %>%
  mutate_all(funs(replace(., is.na(.), 0)))
```

```
## Warning: `funs()` was deprecated in dplyr 0.8.0.
## ℹ Please use a list of either functions or lambdas:
##
## # Simple named list: list(mean = mean, median = median)
##
## # Auto named with `tibble::lst()`: tibble::lst(mean, median)
##
## # Using lambdas list(~ mean(., trim = .2), ~ median(., na.rm = TRUE))
## Call `lifecycle::last_lifecycle_warnings()` to see where this warning was
## generated.
```

```r
# Save the cleaned data to a new file
write.csv(clean_data, "cleaned_data_cortex.csv", row.names = FALSE)

### Matrisome categorization

# Read your first data file into a data frame or data table
data <- read.csv("cleaned_data_cortex.csv")

# Read your second dataset into another data frame or data table
matrisome_info <- read.csv("Bt_Matrisome.csv")

# Perform a left join based on the "Genes" column
joined_data <- left_join(data, matrisome_info, by = "Genes")

# Save the joined data to a new file
write.csv(joined_data, "cortex_matrisome.csv", row.names = FALSE)

# Remove rows where either Matrisome_Division or Matrisome_Category have NA v
```

```
alues
cleaned_joined_data <- joined_data %>%
  filter(!is.na(Matrisome_Division) & !is.na(Matrisome_Category))

# Save the cleaned data to a new file

write.csv(cleaned_joined_data, "cleaned_matrisome_cortex.csv", row.names = FA
LSE)


#######Ploting


#### Collagens
# Read your cleaned joined data file into a data frame
cleaned_joined_data <- read.csv("cleaned_matrisome_cortex.csv")

# Filter rows with Matrisome_Category "Collagens"
collagen_data <- cleaned_joined_data %>%
  filter(Matrisome_Category == "Collagens")

# Reshape the data to have separate columns for each replicate and each tissu
e stage
organized_data <- collagen_data %>%
  pivot_longer(cols = starts_with("CF") | starts_with("CL"),
               names_to = c("Tissue", "Stage"),
               names_sep = "\\.",
               values_to = "Protein_Count")

## Warning: Expected 2 pieces. Missing pieces filled with `NA` in 2 rows [1,
7].

# Save the reshaped data to a new file

write.csv(organized_data, "Collagens_data_matrisome_cortex_to_clean.csv", row
.names = FALSE)

data2 <- read.csv("Collagens_data_matrisome_cortex_clean.csv")

png(filename="cortex_collagens.png",
    width = 12, height = 6, units = "in",
    res = 600)

boxplot(Protein_Count ~ Tissue*Genes,
        data = data2,
        cex.axis = 0.8,
        col = viridis(2, alpha = 0.7),
        lwd = 0.7,
        ylab = "",
```

```
        ylim = c(0, 6000000000),
        axes = FALSE,
        frame.plot = TRUE,
        xlab = "")

GROUP <- unique(data2$Genes)
groups <- 24
numbox <- 2
total <- groups * numbox
xpoints <- seq(median(1:numbox), total, numbox)

axis(2, seq(0, 6000000000, by = 1000000000), las = 1, cex.axis = 0.7)
axis(1, labels = GROUP, at = xpoints, las = 2, cex.axis = 0.7)

stripchart(data2$Protein_Count ~ data2$Tissue*data2$Genes,
           method = "jitter",
           pch = 19,
           col = "black",
           vertical = TRUE,
           add = TRUE,
           cex = 0.3)

title(ylab = expression("Protein abundance (RU)"), line = 3.0)
legend("top", fill = viridis(2), legend = c("Cortex Follicular", "Cortex Lute
al"), horiz = T, cex = 0.7)

dev.off()

## quartz_off_screen
##                  2

# Convert Tissue to factor
data2$Tissue <- as.factor(data2$Tissue)

# Create an empty list to store Tukey results
tukey_results_list <- list()

# Unique genes
unique_genes <- unique(data2$Genes)

# Loop over each gene
for (gene in unique_genes) {
  # Subset data for the current gene
  gene_data <- subset(data2, Genes == gene)

  # Fit the generalized linear model
  glm_model <- glm(Protein_Count ~ Tissue, data = gene_data, family = gaussia
n)
```

```
  # Perform Tukey test
  contrast_matrix <- glht(glm_model, linfct = mcp(Tissue = "Tukey"))
  tukey_results <- summary(contrast_matrix)

  # Store the results
  tukey_results_list[[gene]] <- tukey_results
}

# Print the results for each gene
for (gene in unique_genes) {
  cat("Tukey results for gene:", gene, "\n")
  print(tukey_results_list[[gene]])
  cat("\n")
}

## Tukey results for gene: COL11A1
##
##   Simultaneous Tests for General Linear Hypotheses
##
## Multiple Comparisons of Means: Tukey Contrasts
##
##
## Fit: glm(formula = Protein_Count ~ Tissue, family = gaussian, data = gene_
data)
##
## Linear Hypotheses:
##              Estimate Std. Error z value Pr(>|z|)
## CL - CF == 0   477533     698268   0.684    0.494
## (Adjusted p values reported -- single-step method)
##
##
## Tukey results for gene: COL12A1
##
##   Simultaneous Tests for General Linear Hypotheses
##
## Multiple Comparisons of Means: Tukey Contrasts
##
##
## Fit: glm(formula = Protein_Count ~ Tissue, family = gaussian, data = gene_
data)
##
## Linear Hypotheses:
##              Estimate Std. Error z value Pr(>|z|)
## CL - CF == 0 11666363   14167042   0.823     0.41
## (Adjusted p values reported -- single-step method)
##
##
## Tukey results for gene: COL14A1
##
##   Simultaneous Tests for General Linear Hypotheses
```

```
##
## Multiple Comparisons of Means: Tukey Contrasts
##
##
## Fit: glm(formula = Protein_Count ~ Tissue, family = gaussian, data = gene_
data)
##
## Linear Hypotheses:
##              Estimate Std. Error z value Pr(>|z|)
## CL - CF == 0  3439593    7014117    0.49    0.624
## (Adjusted p values reported -- single-step method)
##
##
## Tukey results for gene: COL15A1
##
##   Simultaneous Tests for General Linear Hypotheses
##
## Multiple Comparisons of Means: Tukey Contrasts
##
##
## Fit: glm(formula = Protein_Count ~ Tissue, family = gaussian, data = gene_
data)
##
## Linear Hypotheses:
##              Estimate Std. Error z value Pr(>|z|)
## CL - CF == 0   157612    1166417   0.135    0.893
## (Adjusted p values reported -- single-step method)
##
##
## Tukey results for gene: COL16A1
##
##   Simultaneous Tests for General Linear Hypotheses
##
## Multiple Comparisons of Means: Tukey Contrasts
##
##
## Fit: glm(formula = Protein_Count ~ Tissue, family = gaussian, data = gene_
data)
##
## Linear Hypotheses:
##              Estimate Std. Error z value Pr(>|z|)
## CL - CF == 0  2716764    5154366   0.527    0.598
## (Adjusted p values reported -- single-step method)
##
##
## Tukey results for gene: COL18A1
##
##   Simultaneous Tests for General Linear Hypotheses
##
## Multiple Comparisons of Means: Tukey Contrasts
```

```
##
##
## Fit: glm(formula = Protein_Count ~ Tissue, family = gaussian, data = gene_
data)
##
## Linear Hypotheses:
##                Estimate Std. Error z value Pr(>|z|)
## CL - CF == 0  -678691    1108341  -0.612     0.54
## (Adjusted p values reported -- single-step method)
##
##
## Tukey results for gene: COL1A1
##
##   Simultaneous Tests for General Linear Hypotheses
##
## Multiple Comparisons of Means: Tukey Contrasts
##
##
## Fit: glm(formula = Protein_Count ~ Tissue, family = gaussian, data = gene_
data)
##
## Linear Hypotheses:
##                 Estimate Std. Error z value Pr(>|z|)
## CL - CF == 0 2.896e+08  1.011e+09   0.286    0.775
## (Adjusted p values reported -- single-step method)
##
##
## Tukey results for gene: COL1A2
##
##   Simultaneous Tests for General Linear Hypotheses
##
## Multiple Comparisons of Means: Tukey Contrasts
##
##
## Fit: glm(formula = Protein_Count ~ Tissue, family = gaussian, data = gene_
data)
##
## Linear Hypotheses:
##                 Estimate Std. Error z value Pr(>|z|)
## CL - CF == 0 100081567  494130609   0.203    0.839
## (Adjusted p values reported -- single-step method)
##
##
## Tukey results for gene: COL27A1
##
##   Simultaneous Tests for General Linear Hypotheses
##
## Multiple Comparisons of Means: Tukey Contrasts
##
##
```

```
## Fit: glm(formula = Protein_Count ~ Tissue, family = gaussian, data = gene_
data)
##
## Linear Hypotheses:
##               Estimate Std. Error z value Pr(>|z|)
## CL - CF == 0    85587      91121   0.939    0.348
## (Adjusted p values reported -- single-step method)
##
##
## Tukey results for gene: COL3A1
##
##   Simultaneous Tests for General Linear Hypotheses
##
## Multiple Comparisons of Means: Tukey Contrasts
##
##
## Fit: glm(formula = Protein_Count ~ Tissue, family = gaussian, data = gene_
data)
##
## Linear Hypotheses:
##               Estimate Std. Error z value Pr(>|z|)
## CL - CF == 0   332821    2069818   0.161    0.872
## (Adjusted p values reported -- single-step method)
##
##
## Tukey results for gene: COL4A1
##
##   Simultaneous Tests for General Linear Hypotheses
##
## Multiple Comparisons of Means: Tukey Contrasts
##
##
## Fit: glm(formula = Protein_Count ~ Tissue, family = gaussian, data = gene_
data)
##
## Linear Hypotheses:
##               Estimate Std. Error z value Pr(>|z|)
## CL - CF == 0  -521322    4756000   -0.11    0.913
## (Adjusted p values reported -- single-step method)
##
##
## Tukey results for gene: COL4A2
##
##   Simultaneous Tests for General Linear Hypotheses
##
## Multiple Comparisons of Means: Tukey Contrasts
##
##
## Fit: glm(formula = Protein_Count ~ Tissue, family = gaussian, data = gene_
data)
```

```
##
## Linear Hypotheses:
##               Estimate Std. Error z value Pr(>|z|)
## CL - CF == 0  -491610    2797415  -0.176    0.861
## (Adjusted p values reported -- single-step method)
##
##
## Tukey results for gene: COL4A3
##
##   Simultaneous Tests for General Linear Hypotheses
##
## Multiple Comparisons of Means: Tukey Contrasts
##
##
## Fit: glm(formula = Protein_Count ~ Tissue, family = gaussian, data = gene_
data)
##
## Linear Hypotheses:
##               Estimate Std. Error z value Pr(>|z|)
## CL - CF == 0  4242525   10505134   0.404    0.686
## (Adjusted p values reported -- single-step method)
##
##
## Tukey results for gene: COL4A4
##
##   Simultaneous Tests for General Linear Hypotheses
##
## Multiple Comparisons of Means: Tukey Contrasts
##
##
## Fit: glm(formula = Protein_Count ~ Tissue, family = gaussian, data = gene_
data)
##
## Linear Hypotheses:
##               Estimate Std. Error z value Pr(>|z|)
## CL - CF == 0   -37188     914713  -0.041    0.968
## (Adjusted p values reported -- single-step method)
##
##
## Tukey results for gene: COL4A5
##
##   Simultaneous Tests for General Linear Hypotheses
##
## Multiple Comparisons of Means: Tukey Contrasts
##
##
## Fit: glm(formula = Protein_Count ~ Tissue, family = gaussian, data = gene_
data)
##
## Linear Hypotheses:
```

```
##               Estimate Std. Error z value Pr(>|z|)
## CL - CF == 0 -1253628    7270124  -0.172    0.863
## (Adjusted p values reported -- single-step method)
##
##
## Tukey results for gene: COL4A6
##
##   Simultaneous Tests for General Linear Hypotheses
##
## Multiple Comparisons of Means: Tukey Contrasts
##
##
## Fit: glm(formula = Protein_Count ~ Tissue, family = gaussian, data = gene_
data)
##
## Linear Hypotheses:
##               Estimate Std. Error z value Pr(>|z|)
## CL - CF == 0 -1019728     892212  -1.143    0.253
## (Adjusted p values reported -- single-step method)
##
##
## Tukey results for gene: COL5A1
##
##   Simultaneous Tests for General Linear Hypotheses
##
## Multiple Comparisons of Means: Tukey Contrasts
##
##
## Fit: glm(formula = Protein_Count ~ Tissue, family = gaussian, data = gene_
data)
##
## Linear Hypotheses:
##               Estimate Std. Error z value Pr(>|z|)
## CL - CF == 0   897687    1879150   0.478    0.633
## (Adjusted p values reported -- single-step method)
##
##
## Tukey results for gene: COL5A2
##
##   Simultaneous Tests for General Linear Hypotheses
##
## Multiple Comparisons of Means: Tukey Contrasts
##
##
## Fit: glm(formula = Protein_Count ~ Tissue, family = gaussian, data = gene_
data)
##
## Linear Hypotheses:
##               Estimate Std. Error z value Pr(>|z|)
## CL - CF == 0  2412890    6919911   0.349    0.727
```

```
## (Adjusted p values reported -- single-step method)
##
##
## Tukey results for gene: COL5A3
##
##   Simultaneous Tests for General Linear Hypotheses
##
## Multiple Comparisons of Means: Tukey Contrasts
##
##
## Fit: glm(formula = Protein_Count ~ Tissue, family = gaussian, data = gene_
data)
##
## Linear Hypotheses:
##              Estimate Std. Error z value Pr(>|z|)
## CL - CF == 0   904569    3867638   0.234    0.815
## (Adjusted p values reported -- single-step method)
##
##
## Tukey results for gene: COL6A1
##
##   Simultaneous Tests for General Linear Hypotheses
##
## Multiple Comparisons of Means: Tukey Contrasts
##
##
## Fit: glm(formula = Protein_Count ~ Tissue, family = gaussian, data = gene_
data)
##
## Linear Hypotheses:
##              Estimate Std. Error z value Pr(>|z|)
## CL - CF == 0 11014067   20857617   0.528    0.597
## (Adjusted p values reported -- single-step method)
##
##
## Tukey results for gene: COL6A2
##
##   Simultaneous Tests for General Linear Hypotheses
##
## Multiple Comparisons of Means: Tukey Contrasts
##
##
## Fit: glm(formula = Protein_Count ~ Tissue, family = gaussian, data = gene_
data)
##
## Linear Hypotheses:
##              Estimate Std. Error z value Pr(>|z|)
## CL - CF == 0  8011653   11601863   0.691     0.49
## (Adjusted p values reported -- single-step method)
##
```

```
##
## Tukey results for gene: COL6A3
##
##   Simultaneous Tests for General Linear Hypotheses
##
## Multiple Comparisons of Means: Tukey Contrasts
##
##
## Fit: glm(formula = Protein_Count ~ Tissue, family = gaussian, data = gene_
data)
##
## Linear Hypotheses:
##              Estimate Std. Error z value Pr(>|z|)
## CL - CF == 0 29768967   41339474    0.72    0.471
## (Adjusted p values reported -- single-step method)
##
##
## Tukey results for gene: COL6A5
##
##   Simultaneous Tests for General Linear Hypotheses
##
## Multiple Comparisons of Means: Tukey Contrasts
##
##
## Fit: glm(formula = Protein_Count ~ Tissue, family = gaussian, data = gene_
data)
##
## Linear Hypotheses:
##              Estimate Std. Error z value Pr(>|z|)
## CL - CF == 0    91272     194030    0.47    0.638
## (Adjusted p values reported -- single-step method)
##
##
## Tukey results for gene: COL8A1
##
##   Simultaneous Tests for General Linear Hypotheses
##
## Multiple Comparisons of Means: Tukey Contrasts
##
##
## Fit: glm(formula = Protein_Count ~ Tissue, family = gaussian, data = gene_
data)
##
## Linear Hypotheses:
##              Estimate Std. Error z value Pr(>|z|)
## CL - CF == 0   438338     413477    1.06    0.289
## (Adjusted p values reported -- single-step method)

#### ECM Glycoproteins
# Read your cleaned joined data file into a data frame
```

```r
cleaned_joined_data <- read.csv("cleaned_matrisome_cortex.csv")

# Filter rows with Matrisome_Category "ECM Glycoproteins"
Glycoproteins_data <- cleaned_joined_data %>%
  filter(Matrisome_Category == "ECM Glycoproteins")

# Reshape the data to have separate columns for each replicate and each tissu
e stage
organized_data <- Glycoproteins_data %>%
  pivot_longer(cols = starts_with("CF") | starts_with("CL"),
               names_to = c("Tissue", "Stage"),
               names_sep = "\\.",
               values_to = "Protein_Count")

## Warning: Expected 2 pieces. Missing pieces filled with `NA` in 2 rows [1,
7].

# Save the reshaped data to a new file

write.csv(organized_data, "Glycoproteins_data_matrisome_cortex_to_clean.csv",
row.names = FALSE)

data2 <- read.csv("Glycoproteins_data_matrisome_cortex_clean.csv")

png(filename="cortex_ECM Glycoproteins.png",
    width = 12, height = 6, units = "in",
    res = 600)

boxplot(Protein_Count ~ Tissue*Genes,
        data = data2,
        cex.axis = 0.8,
        col = viridis(2, alpha = 0.7),
        lwd = 0.7,
        ylab = "",
        ylim = c(0, 700000000),
        axes = FALSE,
        frame.plot = TRUE,
        xlab = "")

GROUP <- unique(data2$Genes)
groups <- 83
numbox <- 2
total <- groups * numbox
xpoints <- seq(median(1:numbox), total, numbox)

axis(2, seq(0, 700000000, by = 100000000), las = 1, cex.axis = 0.7)
axis(1, labels = GROUP, at = xpoints, las = 2, cex.axis = 0.7)

stripchart(data2$Protein_Count ~ data2$Tissue*data2$Genes,
```

```
           method = "jitter",
           pch = 19,
           col = "black",
           vertical = TRUE,
           add = TRUE,
           cex = 0.3)

title(ylab = expression("Protein abundance (RU)"), line = 3.0)
legend("top", fill = viridis(2), legend = c("Cortex Follicular", "Cortex Lute
al"), horiz = T, cex = 0.7)

dev.off()

## quartz_off_screen
##                  2

# Convert Tissue to factor
data2$Tissue <- as.factor(data2$Tissue)

# Create an empty list to store Tukey results
tukey_results_list <- list()

# Unique genes
unique_genes <- unique(data2$Genes)

# Loop over each gene
for (gene in unique_genes) {
  # Subset data for the current gene
  gene_data <- subset(data2, Genes == gene)

  # Fit the generalized linear model
  glm_model <- glm(Protein_Count ~ Tissue, data = gene_data, family = gaussia
n)

  # Perform Tukey test
  contrast_matrix <- glht(glm_model, linfct = mcp(Tissue = "Tukey"))
  tukey_results <- summary(contrast_matrix)

  # Store the results
  tukey_results_list[[gene]] <- tukey_results
}

# Print the results for each gene
for (gene in unique_genes) {
  cat("Tukey results for gene:", gene, "\n")
  print(tukey_results_list[[gene]])
  cat("\n")
}
```

```
## Tukey results for gene: ABI3BP
##
##   Simultaneous Tests for General Linear Hypotheses
##
## Multiple Comparisons of Means: Tukey Contrasts
##
##
## Fit: glm(formula = Protein_Count ~ Tissue, family = gaussian, data = gene_
data)
##
## Linear Hypotheses:
##              Estimate Std. Error z value Pr(>|z|)
## CL - CF == 0  -314814     436225  -0.722     0.47
## (Adjusted p values reported -- single-step method)
##
##
## Tukey results for gene: AEBP1
##
##   Simultaneous Tests for General Linear Hypotheses
##
## Multiple Comparisons of Means: Tukey Contrasts
##
##
## Fit: glm(formula = Protein_Count ~ Tissue, family = gaussian, data = gene_
data)
##
## Linear Hypotheses:
##              Estimate Std. Error z value Pr(>|z|)
## CL - CF == 0   389787     956062   0.408    0.683
## (Adjusted p values reported -- single-step method)
##
##
## Tukey results for gene: AGRN
##
##   Simultaneous Tests for General Linear Hypotheses
##
## Multiple Comparisons of Means: Tukey Contrasts
##
##
## Fit: glm(formula = Protein_Count ~ Tissue, family = gaussian, data = gene_
data)
##
## Linear Hypotheses:
##              Estimate Std. Error z value Pr(>|z|)
## CL - CF == 0  -484765     411288  -1.179    0.239
## (Adjusted p values reported -- single-step method)
##
##
## Tukey results for gene: AHSG
##
```

```
##   Simultaneous Tests for General Linear Hypotheses
##
## Multiple Comparisons of Means: Tukey Contrasts
##
##
## Fit: glm(formula = Protein_Count ~ Tissue, family = gaussian, data = gene_
data)
##
## Linear Hypotheses:
##              Estimate Std. Error z value Pr(>|z|)
## CL - CF == 0  -221213     386374  -0.573    0.567
## (Adjusted p values reported -- single-step method)
##
##
## Tukey results for gene: CILP2
##
##   Simultaneous Tests for General Linear Hypotheses
##
## Multiple Comparisons of Means: Tukey Contrasts
##
##
## Fit: glm(formula = Protein_Count ~ Tissue, family = gaussian, data = gene_
data)
##
## Linear Hypotheses:
##              Estimate Std. Error z value Pr(>|z|)
## CL - CF == 0  5300853   27074639   0.196    0.845
## (Adjusted p values reported -- single-step method)
##
##
## Tukey results for gene: COCH
##
##   Simultaneous Tests for General Linear Hypotheses
##
## Multiple Comparisons of Means: Tukey Contrasts
##
##
## Fit: glm(formula = Protein_Count ~ Tissue, family = gaussian, data = gene_
data)
##
## Linear Hypotheses:
##              Estimate Std. Error z value Pr(>|z|)
## CL - CF == 0     1310     125244    0.01    0.992
## (Adjusted p values reported -- single-step method)
##
##
## Tukey results for gene: DPT
##
##   Simultaneous Tests for General Linear Hypotheses
##
```

```
## Multiple Comparisons of Means: Tukey Contrasts
##
##
## Fit: glm(formula = Protein_Count ~ Tissue, family = gaussian, data = gene_
data)
##
## Linear Hypotheses:
##               Estimate Std. Error z value Pr(>|z|)
## CL - CF == 0 -29002167  106790803  -0.272    0.786
## (Adjusted p values reported -- single-step method)
##
##
## Tukey results for gene: ECM1
##
##   Simultaneous Tests for General Linear Hypotheses
##
## Multiple Comparisons of Means: Tukey Contrasts
##
##
## Fit: glm(formula = Protein_Count ~ Tissue, family = gaussian, data = gene_
data)
##
## Linear Hypotheses:
##              Estimate Std. Error z value Pr(>|z|)
## CL - CF == 0  -177923    1753079  -0.101    0.919
## (Adjusted p values reported -- single-step method)
##
##
## Tukey results for gene: ECM2
##
##   Simultaneous Tests for General Linear Hypotheses
##
## Multiple Comparisons of Means: Tukey Contrasts
##
##
## Fit: glm(formula = Protein_Count ~ Tissue, family = gaussian, data = gene_
data)
##
## Linear Hypotheses:
##              Estimate Std. Error z value Pr(>|z|)
## CL - CF == 0   377102     442554   0.852    0.394
## (Adjusted p values reported -- single-step method)
##
##
## Tukey results for gene: EFEMP1
##
##   Simultaneous Tests for General Linear Hypotheses
##
## Multiple Comparisons of Means: Tukey Contrasts
##
```

```
##
## Fit: glm(formula = Protein_Count ~ Tissue, family = gaussian, data = gene_
data)
##
## Linear Hypotheses:
##               Estimate Std. Error z value Pr(>|z|)
## CL - CF == 0  -190958    1503249  -0.127    0.899
## (Adjusted p values reported -- single-step method)
##
##
## Tukey results for gene: EFEMP2
##
##   Simultaneous Tests for General Linear Hypotheses
##
## Multiple Comparisons of Means: Tukey Contrasts
##
##
## Fit: glm(formula = Protein_Count ~ Tissue, family = gaussian, data = gene_
data)
##
## Linear Hypotheses:
##               Estimate Std. Error z value Pr(>|z|)
## CL - CF == 0    94572     768296   0.123    0.902
## (Adjusted p values reported -- single-step method)
##
##
## Tukey results for gene: EGFLAM
##
##   Simultaneous Tests for General Linear Hypotheses
##
## Multiple Comparisons of Means: Tukey Contrasts
##
##
## Fit: glm(formula = Protein_Count ~ Tissue, family = gaussian, data = gene_
data)
##
## Linear Hypotheses:
##               Estimate Std. Error z value Pr(>|z|)
## CL - CF == 0  3788745    3316083   1.143    0.253
## (Adjusted p values reported -- single-step method)
##
##
## Tukey results for gene: EMID1
##
##   Simultaneous Tests for General Linear Hypotheses
##
## Multiple Comparisons of Means: Tukey Contrasts
##
##
## Fit: glm(formula = Protein_Count ~ Tissue, family = gaussian, data = gene_
```

```
data)
##
## Linear Hypotheses:
##               Estimate Std. Error z value Pr(>|z|)
## CL - CF == 0   667369     384675   1.735   0.0828 .
## ---
## Signif. codes:  0 '***' 0.001 '**' 0.01 '*' 0.05 '.' 0.1 ' ' 1
## (Adjusted p values reported -- single-step method)
##
##
## Tukey results for gene: EMILIN1
##
##   Simultaneous Tests for General Linear Hypotheses
##
## Multiple Comparisons of Means: Tukey Contrasts
##
##
## Fit: glm(formula = Protein_Count ~ Tissue, family = gaussian, data = gene_
data)
##
## Linear Hypotheses:
##               Estimate Std. Error z value Pr(>|z|)
## CL - CF == 0 -3874817    6081018  -0.637    0.524
## (Adjusted p values reported -- single-step method)
##
##
## Tukey results for gene: EMILIN2
##
##   Simultaneous Tests for General Linear Hypotheses
##
## Multiple Comparisons of Means: Tukey Contrasts
##
##
## Fit: glm(formula = Protein_Count ~ Tissue, family = gaussian, data = gene_
data)
##
## Linear Hypotheses:
##               Estimate Std. Error z value Pr(>|z|)
## CL - CF == 0   -44176     190712  -0.232    0.817
## (Adjusted p values reported -- single-step method)
##
##
## Tukey results for gene: EMILIN3
##
##   Simultaneous Tests for General Linear Hypotheses
##
## Multiple Comparisons of Means: Tukey Contrasts
##
##
## Fit: glm(formula = Protein_Count ~ Tissue, family = gaussian, data = gene_
```

```
data)
##
## Linear Hypotheses:
##                Estimate Std. Error z value Pr(>|z|)
## CL - CF == 0  -359866     414004  -0.869    0.385
## (Adjusted p values reported -- single-step method)
##
##
## Tukey results for gene: FBLN1
##
##   Simultaneous Tests for General Linear Hypotheses
##
## Multiple Comparisons of Means: Tukey Contrasts
##
##
## Fit: glm(formula = Protein_Count ~ Tissue, family = gaussian, data = gene_
data)
##
## Linear Hypotheses:
##                Estimate Std. Error z value Pr(>|z|)
## CL - CF == 0   132542    1096647   0.121    0.904
## (Adjusted p values reported -- single-step method)
##
##
## Tukey results for gene: FBLN2
##
##   Simultaneous Tests for General Linear Hypotheses
##
## Multiple Comparisons of Means: Tukey Contrasts
##
##
## Fit: glm(formula = Protein_Count ~ Tissue, family = gaussian, data = gene_
data)
##
## Linear Hypotheses:
##                Estimate Std. Error z value Pr(>|z|)
## CL - CF == 0   -99295     168927  -0.588    0.557
## (Adjusted p values reported -- single-step method)
##
##
## Tukey results for gene: FBLN5
##
##   Simultaneous Tests for General Linear Hypotheses
##
## Multiple Comparisons of Means: Tukey Contrasts
##
##
## Fit: glm(formula = Protein_Count ~ Tissue, family = gaussian, data = gene_
data)
##
```

```
## Linear Hypotheses:
##              Estimate Std. Error z value Pr(>|z|)
## CL - CF == 0   137126    1996090   0.069    0.945
## (Adjusted p values reported -- single-step method)
##
##
## Tukey results for gene: FBLN7
##
##   Simultaneous Tests for General Linear Hypotheses
##
## Multiple Comparisons of Means: Tukey Contrasts
##
##
## Fit: glm(formula = Protein_Count ~ Tissue, family = gaussian, data = gene_
data)
##
## Linear Hypotheses:
##              Estimate Std. Error z value Pr(>|z|)
## CL - CF == 0     9986      83782   0.119    0.905
## (Adjusted p values reported -- single-step method)
##
##
## Tukey results for gene: FBN1
##
##   Simultaneous Tests for General Linear Hypotheses
##
## Multiple Comparisons of Means: Tukey Contrasts
##
##
## Fit: glm(formula = Protein_Count ~ Tissue, family = gaussian, data = gene_
data)
##
## Linear Hypotheses:
##              Estimate Std. Error z value Pr(>|z|)
## CL - CF == 0   593237    1390166   0.427     0.67
## (Adjusted p values reported -- single-step method)
##
##
## Tukey results for gene: FBN2
##
##   Simultaneous Tests for General Linear Hypotheses
##
## Multiple Comparisons of Means: Tukey Contrasts
##
##
## Fit: glm(formula = Protein_Count ~ Tissue, family = gaussian, data = gene_
data)
##
## Linear Hypotheses:
##              Estimate Std. Error z value Pr(>|z|)
```

```
## CL - CF == 0   206216     320555   0.643     0.52
## (Adjusted p values reported -- single-step method)
##
##
## Tukey results for gene: FBN3
##
##   Simultaneous Tests for General Linear Hypotheses
##
## Multiple Comparisons of Means: Tukey Contrasts
##
##
## Fit: glm(formula = Protein_Count ~ Tissue, family = gaussian, data = gene_
data)
##
## Linear Hypotheses:
##              Estimate Std. Error z value Pr(>|z|)
## CL - CF == 0   646808     486859   1.329    0.184
## (Adjusted p values reported -- single-step method)
##
##
## Tukey results for gene: FGA
##
##   Simultaneous Tests for General Linear Hypotheses
##
## Multiple Comparisons of Means: Tukey Contrasts
##
##
## Fit: glm(formula = Protein_Count ~ Tissue, family = gaussian, data = gene_
data)
##
## Linear Hypotheses:
##              Estimate Std. Error z value Pr(>|z|)
## CL - CF == 0 -3008133    4011015   -0.75    0.453
## (Adjusted p values reported -- single-step method)
##
##
## Tukey results for gene: FGB
##
##   Simultaneous Tests for General Linear Hypotheses
##
## Multiple Comparisons of Means: Tukey Contrasts
##
##
## Fit: glm(formula = Protein_Count ~ Tissue, family = gaussian, data = gene_
data)
##
## Linear Hypotheses:
##              Estimate Std. Error z value Pr(>|z|)
## CL - CF == 0 -3452060    6236367  -0.554     0.58
## (Adjusted p values reported -- single-step method)
```

```
##
##
## Tukey results for gene: FGG
##
##   Simultaneous Tests for General Linear Hypotheses
##
## Multiple Comparisons of Means: Tukey Contrasts
##
##
## Fit: glm(formula = Protein_Count ~ Tissue, family = gaussian, data = gene_
data)
##
## Linear Hypotheses:
##              Estimate Std. Error z value Pr(>|z|)
## CL - CF == 0  -995798    1557127   -0.64    0.522
## (Adjusted p values reported -- single-step method)
##
##
## Tukey results for gene: FGL2
##
##   Simultaneous Tests for General Linear Hypotheses
##
## Multiple Comparisons of Means: Tukey Contrasts
##
##
## Fit: glm(formula = Protein_Count ~ Tissue, family = gaussian, data = gene_
data)
##
## Linear Hypotheses:
##              Estimate Std. Error z value Pr(>|z|)
## CL - CF == 0   -17596      99883  -0.176     0.86
## (Adjusted p values reported -- single-step method)
##
##
## Tukey results for gene: FN1
##
##   Simultaneous Tests for General Linear Hypotheses
##
## Multiple Comparisons of Means: Tukey Contrasts
##
##
## Fit: glm(formula = Protein_Count ~ Tissue, family = gaussian, data = gene_
data)
##
## Linear Hypotheses:
##              Estimate Std. Error z value Pr(>|z|)
## CL - CF == 0  1710171    3639145    0.47    0.638
## (Adjusted p values reported -- single-step method)
##
##
```

```
## Tukey results for gene: FRAS1
##
##   Simultaneous Tests for General Linear Hypotheses
##
## Multiple Comparisons of Means: Tukey Contrasts
##
##
## Fit: glm(formula = Protein_Count ~ Tissue, family = gaussian, data = gene_
data)
##
## Linear Hypotheses:
##              Estimate Std. Error z value Pr(>|z|)
## CL - CF == 0   419983     241247   1.741   0.0817 .
## ---
## Signif. codes:  0 '***' 0.001 '**' 0.01 '*' 0.05 '.' 0.1 ' ' 1
## (Adjusted p values reported -- single-step method)
##
##
## Tukey results for gene: HMCN1
##
##   Simultaneous Tests for General Linear Hypotheses
##
## Multiple Comparisons of Means: Tukey Contrasts
##
##
## Fit: glm(formula = Protein_Count ~ Tissue, family = gaussian, data = gene_
data)
##
## Linear Hypotheses:
##              Estimate Std. Error z value Pr(>|z|)
## CL - CF == 0   -51002      84206  -0.606    0.545
## (Adjusted p values reported -- single-step method)
##
##
## Tukey results for gene: IGFALS
##
##   Simultaneous Tests for General Linear Hypotheses
##
## Multiple Comparisons of Means: Tukey Contrasts
##
##
## Fit: glm(formula = Protein_Count ~ Tissue, family = gaussian, data = gene_
data)
##
## Linear Hypotheses:
##              Estimate Std. Error z value Pr(>|z|)
## CL - CF == 0    56996     154085    0.37    0.711
## (Adjusted p values reported -- single-step method)
##
##
```

```
## Tukey results for gene: IGFBP6
##
##   Simultaneous Tests for General Linear Hypotheses
##
## Multiple Comparisons of Means: Tukey Contrasts
##
##
## Fit: glm(formula = Protein_Count ~ Tissue, family = gaussian, data = gene_
data)
##
## Linear Hypotheses:
##              Estimate Std. Error z value Pr(>|z|)
## CL - CF == 0    25776      46477   0.555    0.579
## (Adjusted p values reported -- single-step method)
##
##
## Tukey results for gene: IGFBP7
##
##   Simultaneous Tests for General Linear Hypotheses
##
## Multiple Comparisons of Means: Tukey Contrasts
##
##
## Fit: glm(formula = Protein_Count ~ Tissue, family = gaussian, data = gene_
data)
##
## Linear Hypotheses:
##              Estimate Std. Error z value Pr(>|z|)
## CL - CF == 0  -198679     477706  -0.416    0.677
## (Adjusted p values reported -- single-step method)
##
##
## Tukey results for gene: IGSF10
##
##   Simultaneous Tests for General Linear Hypotheses
##
## Multiple Comparisons of Means: Tukey Contrasts
##
##
## Fit: glm(formula = Protein_Count ~ Tissue, family = gaussian, data = gene_
data)
##
## Linear Hypotheses:
##              Estimate Std. Error z value Pr(>|z|)
## CL - CF == 0     7949      47969   0.166    0.868
## (Adjusted p values reported -- single-step method)
##
##
## Tukey results for gene: LAMA1
##
```

```
##   Simultaneous Tests for General Linear Hypotheses
##
## Multiple Comparisons of Means: Tukey Contrasts
##
##
## Fit: glm(formula = Protein_Count ~ Tissue, family = gaussian, data = gene_
data)
##
## Linear Hypotheses:
##              Estimate Std. Error z value Pr(>|z|)
## CL - CF == 0  -187984     746301  -0.252    0.801
## (Adjusted p values reported -- single-step method)
##
##
## Tukey results for gene: LAMA2
##
##   Simultaneous Tests for General Linear Hypotheses
##
## Multiple Comparisons of Means: Tukey Contrasts
##
##
## Fit: glm(formula = Protein_Count ~ Tissue, family = gaussian, data = gene_
data)
##
## Linear Hypotheses:
##              Estimate Std. Error z value Pr(>|z|)
## CL - CF == 0   -66167     124682  -0.531    0.596
## (Adjusted p values reported -- single-step method)
##
##
## Tukey results for gene: LAMA3
##
##   Simultaneous Tests for General Linear Hypotheses
##
## Multiple Comparisons of Means: Tukey Contrasts
##
##
## Fit: glm(formula = Protein_Count ~ Tissue, family = gaussian, data = gene_
data)
##
## Linear Hypotheses:
##              Estimate Std. Error z value Pr(>|z|)
## CL - CF == 0    85991     224631   0.383    0.702
## (Adjusted p values reported -- single-step method)
##
##
## Tukey results for gene: LAMA4
##
##   Simultaneous Tests for General Linear Hypotheses
##
```

```
## Multiple Comparisons of Means: Tukey Contrasts
##
##
## Fit: glm(formula = Protein_Count ~ Tissue, family = gaussian, data = gene_
data)
##
## Linear Hypotheses:
##              Estimate Std. Error z value Pr(>|z|)
## CL - CF == 0  -245259     527524  -0.465    0.642
## (Adjusted p values reported -- single-step method)
##
##
## Tukey results for gene: LAMA5
##
##   Simultaneous Tests for General Linear Hypotheses
##
## Multiple Comparisons of Means: Tukey Contrasts
##
##
## Fit: glm(formula = Protein_Count ~ Tissue, family = gaussian, data = gene_
data)
##
## Linear Hypotheses:
##              Estimate Std. Error z value Pr(>|z|)
## CL - CF == 0  -376713     931299  -0.405    0.686
## (Adjusted p values reported -- single-step method)
##
##
## Tukey results for gene: LAMB1
##
##   Simultaneous Tests for General Linear Hypotheses
##
## Multiple Comparisons of Means: Tukey Contrasts
##
##
## Fit: glm(formula = Protein_Count ~ Tissue, family = gaussian, data = gene_
data)
##
## Linear Hypotheses:
##              Estimate Std. Error z value Pr(>|z|)
## CL - CF == 0  -446963     974800  -0.459    0.647
## (Adjusted p values reported -- single-step method)
##
##
## Tukey results for gene: LAMB2
##
##   Simultaneous Tests for General Linear Hypotheses
##
## Multiple Comparisons of Means: Tukey Contrasts
##
```

```
##
## Fit: glm(formula = Protein_Count ~ Tissue, family = gaussian, data = gene_
data)
##
## Linear Hypotheses:
##              Estimate Std. Error z value Pr(>|z|)
## CL - CF == 0  -606782    1989195  -0.305     0.76
## (Adjusted p values reported -- single-step method)
##
##
## Tukey results for gene: LAMB3
##
##   Simultaneous Tests for General Linear Hypotheses
##
## Multiple Comparisons of Means: Tukey Contrasts
##
##
## Fit: glm(formula = Protein_Count ~ Tissue, family = gaussian, data = gene_
data)
##
## Linear Hypotheses:
##              Estimate Std. Error z value Pr(>|z|)
## CL - CF == 0    61148      67720   0.903    0.367
## (Adjusted p values reported -- single-step method)
##
##
## Tukey results for gene: LAMC1
##
##   Simultaneous Tests for General Linear Hypotheses
##
## Multiple Comparisons of Means: Tukey Contrasts
##
##
## Fit: glm(formula = Protein_Count ~ Tissue, family = gaussian, data = gene_
data)
##
## Linear Hypotheses:
##              Estimate Std. Error z value Pr(>|z|)
## CL - CF == 0  -739463    1923752  -0.384    0.701
## (Adjusted p values reported -- single-step method)
##
##
## Tukey results for gene: LAMC3
##
##   Simultaneous Tests for General Linear Hypotheses
##
## Multiple Comparisons of Means: Tukey Contrasts
##
##
## Fit: glm(formula = Protein_Count ~ Tissue, family = gaussian, data = gene_
```

```
data)
##
## Linear Hypotheses:
##              Estimate Std. Error z value Pr(>|z|)
## CL - CF == 0   -31361     100143  -0.313    0.754
## (Adjusted p values reported -- single-step method)
##
##
## Tukey results for gene: LRG1
##
##   Simultaneous Tests for General Linear Hypotheses
##
## Multiple Comparisons of Means: Tukey Contrasts
##
##
## Fit: glm(formula = Protein_Count ~ Tissue, family = gaussian, data = gene_
data)
##
## Linear Hypotheses:
##              Estimate Std. Error z value Pr(>|z|)
## CL - CF == 0   -96459      78121  -1.235    0.217
## (Adjusted p values reported -- single-step method)
##
##
## Tukey results for gene: LTBP1
##
##   Simultaneous Tests for General Linear Hypotheses
##
## Multiple Comparisons of Means: Tukey Contrasts
##
##
## Fit: glm(formula = Protein_Count ~ Tissue, family = gaussian, data = gene_
data)
##
## Linear Hypotheses:
##              Estimate Std. Error z value Pr(>|z|)
## CL - CF == 0   -60651     337337   -0.18    0.857
## (Adjusted p values reported -- single-step method)
##
##
## Tukey results for gene: LTBP2
##
##   Simultaneous Tests for General Linear Hypotheses
##
## Multiple Comparisons of Means: Tukey Contrasts
##
##
## Fit: glm(formula = Protein_Count ~ Tissue, family = gaussian, data = gene_
data)
##
```

```
## Linear Hypotheses:
##              Estimate Std. Error z value Pr(>|z|)
## CL - CF == 0    35135     186813   0.188    0.851
## (Adjusted p values reported -- single-step method)
##
##
## Tukey results for gene: LTBP3
##
##   Simultaneous Tests for General Linear Hypotheses
##
## Multiple Comparisons of Means: Tukey Contrasts
##
##
## Fit: glm(formula = Protein_Count ~ Tissue, family = gaussian, data = gene_
data)
##
## Linear Hypotheses:
##              Estimate Std. Error z value Pr(>|z|)
## CL - CF == 0   226204     185835   1.217    0.224
## (Adjusted p values reported -- single-step method)
##
##
## Tukey results for gene: LTBP4
##
##   Simultaneous Tests for General Linear Hypotheses
##
## Multiple Comparisons of Means: Tukey Contrasts
##
##
## Fit: glm(formula = Protein_Count ~ Tissue, family = gaussian, data = gene_
data)
##
## Linear Hypotheses:
##              Estimate Std. Error z value Pr(>|z|)
## CL - CF == 0  -316480    2470935  -0.128    0.898
## (Adjusted p values reported -- single-step method)
##
##
## Tukey results for gene: MATN2
##
##   Simultaneous Tests for General Linear Hypotheses
##
## Multiple Comparisons of Means: Tukey Contrasts
##
##
## Fit: glm(formula = Protein_Count ~ Tissue, family = gaussian, data = gene_
data)
##
## Linear Hypotheses:
##              Estimate Std. Error z value Pr(>|z|)
```

```
## CL - CF == 0   161187     559448   0.288    0.773
## (Adjusted p values reported -- single-step method)
##
##
## Tukey results for gene: MFAP2
##
##   Simultaneous Tests for General Linear Hypotheses
##
## Multiple Comparisons of Means: Tukey Contrasts
##
##
## Fit: glm(formula = Protein_Count ~ Tissue, family = gaussian, data = gene_
data)
##
## Linear Hypotheses:
##              Estimate Std. Error z value Pr(>|z|)
## CL - CF == 0  3289057    8149281   0.404    0.687
## (Adjusted p values reported -- single-step method)
##
##
## Tukey results for gene: MFAP4
##
##   Simultaneous Tests for General Linear Hypotheses
##
## Multiple Comparisons of Means: Tukey Contrasts
##
##
## Fit: glm(formula = Protein_Count ~ Tissue, family = gaussian, data = gene_
data)
##
## Linear Hypotheses:
##              Estimate Std. Error z value Pr(>|z|)
## CL - CF == 0  3558280    4882529   0.729    0.466
## (Adjusted p values reported -- single-step method)
##
##
## Tukey results for gene: MFAP5
##
##   Simultaneous Tests for General Linear Hypotheses
##
## Multiple Comparisons of Means: Tukey Contrasts
##
##
## Fit: glm(formula = Protein_Count ~ Tissue, family = gaussian, data = gene_
data)
##
## Linear Hypotheses:
##              Estimate Std. Error z value Pr(>|z|)
## CL - CF == 0   -24746     266976  -0.093    0.926
## (Adjusted p values reported -- single-step method)
```

```
##
##
## Tukey results for gene: MFGE8
##
##   Simultaneous Tests for General Linear Hypotheses
##
## Multiple Comparisons of Means: Tukey Contrasts
##
##
## Fit: glm(formula = Protein_Count ~ Tissue, family = gaussian, data = gene_
data)
##
## Linear Hypotheses:
##              Estimate Std. Error z value Pr(>|z|)
## CL - CF == 0  -151529     166036  -0.913    0.361
## (Adjusted p values reported -- single-step method)
##
##
## Tukey results for gene: MGP
##
##   Simultaneous Tests for General Linear Hypotheses
##
## Multiple Comparisons of Means: Tukey Contrasts
##
##
## Fit: glm(formula = Protein_Count ~ Tissue, family = gaussian, data = gene_
data)
##
## Linear Hypotheses:
##              Estimate Std. Error z value Pr(>|z|)
## CL - CF == 0  2680834    5842367   0.459    0.646
## (Adjusted p values reported -- single-step method)
##
##
## Tukey results for gene: MMRN2
##
##   Simultaneous Tests for General Linear Hypotheses
##
## Multiple Comparisons of Means: Tukey Contrasts
##
##
## Fit: glm(formula = Protein_Count ~ Tissue, family = gaussian, data = gene_
data)
##
## Linear Hypotheses:
##              Estimate Std. Error z value Pr(>|z|)
## CL - CF == 0    -8060      65473  -0.123    0.902
## (Adjusted p values reported -- single-step method)
##
##
```

```
## Tukey results for gene: NID1
##
##   Simultaneous Tests for General Linear Hypotheses
##
## Multiple Comparisons of Means: Tukey Contrasts
##
##
## Fit: glm(formula = Protein_Count ~ Tissue, family = gaussian, data = gene_
data)
##
## Linear Hypotheses:
##              Estimate Std. Error z value Pr(>|z|)
## CL - CF == 0  4672057   21254992    0.22    0.826
## (Adjusted p values reported -- single-step method)
##
##
## Tukey results for gene: NID2
##
##   Simultaneous Tests for General Linear Hypotheses
##
## Multiple Comparisons of Means: Tukey Contrasts
##
##
## Fit: glm(formula = Protein_Count ~ Tissue, family = gaussian, data = gene_
data)
##
## Linear Hypotheses:
##              Estimate Std. Error z value Pr(>|z|)
## CL - CF == 0  -636830    4705067  -0.135    0.892
## (Adjusted p values reported -- single-step method)
##
##
## Tukey results for gene: PCOLCE
##
##   Simultaneous Tests for General Linear Hypotheses
##
## Multiple Comparisons of Means: Tukey Contrasts
##
##
## Fit: glm(formula = Protein_Count ~ Tissue, family = gaussian, data = gene_
data)
##
## Linear Hypotheses:
##              Estimate Std. Error z value Pr(>|z|)
## CL - CF == 0  -869483     745536  -1.166    0.244
## (Adjusted p values reported -- single-step method)
##
##
## Tukey results for gene: PCOLCE2
##
```

```
##   Simultaneous Tests for General Linear Hypotheses
##
## Multiple Comparisons of Means: Tukey Contrasts
##
##
## Fit: glm(formula = Protein_Count ~ Tissue, family = gaussian, data = gene_
data)
##
## Linear Hypotheses:
##              Estimate Std. Error z value Pr(>|z|)
## CL - CF == 0     2671      24460   0.109    0.913
## (Adjusted p values reported -- single-step method)
##
##
## Tukey results for gene: POSTN
##
##   Simultaneous Tests for General Linear Hypotheses
##
## Multiple Comparisons of Means: Tukey Contrasts
##
##
## Fit: glm(formula = Protein_Count ~ Tissue, family = gaussian, data = gene_
data)
##
## Linear Hypotheses:
##              Estimate Std. Error z value Pr(>|z|)
## CL - CF == 0   507475     794159   0.639    0.523
## (Adjusted p values reported -- single-step method)
##
##
## Tukey results for gene: PXDN
##
##   Simultaneous Tests for General Linear Hypotheses
##
## Multiple Comparisons of Means: Tukey Contrasts
##
##
## Fit: glm(formula = Protein_Count ~ Tissue, family = gaussian, data = gene_
data)
##
## Linear Hypotheses:
##              Estimate Std. Error z value Pr(>|z|)
## CL - CF == 0  -231255     137932  -1.677   0.0936 .
## ---
## Signif. codes:  0 '***' 0.001 '**' 0.01 '*' 0.05 '.' 0.1 ' ' 1
## (Adjusted p values reported -- single-step method)
##
##
## Tukey results for gene: RSPO3
##
```

```
##   Simultaneous Tests for General Linear Hypotheses
##
## Multiple Comparisons of Means: Tukey Contrasts
##
##
## Fit: glm(formula = Protein_Count ~ Tissue, family = gaussian, data = gene_
data)
##
## Linear Hypotheses:
##              Estimate Std. Error z value Pr(>|z|)
## CL - CF == 0  -190859     287532  -0.664    0.507
## (Adjusted p values reported -- single-step method)
##
##
## Tukey results for gene: SBSPON
##
##   Simultaneous Tests for General Linear Hypotheses
##
## Multiple Comparisons of Means: Tukey Contrasts
##
##
## Fit: glm(formula = Protein_Count ~ Tissue, family = gaussian, data = gene_
data)
##
## Linear Hypotheses:
##              Estimate Std. Error z value Pr(>|z|)
## CL - CF == 0  2268822    4243420   0.535    0.593
## (Adjusted p values reported -- single-step method)
##
##
## Tukey results for gene: SPARC
##
##   Simultaneous Tests for General Linear Hypotheses
##
## Multiple Comparisons of Means: Tukey Contrasts
##
##
## Fit: glm(formula = Protein_Count ~ Tissue, family = gaussian, data = gene_
data)
##
## Linear Hypotheses:
##              Estimate Std. Error z value Pr(>|z|)
## CL - CF == 0  -453827    1294172  -0.351    0.726
## (Adjusted p values reported -- single-step method)
##
##
## Tukey results for gene: SPARCL1
##
##   Simultaneous Tests for General Linear Hypotheses
##
```

```
## Multiple Comparisons of Means: Tukey Contrasts
##
##
## Fit: glm(formula = Protein_Count ~ Tissue, family = gaussian, data = gene_
data)
##
## Linear Hypotheses:
##              Estimate Std. Error z value Pr(>|z|)
## CL - CF == 0   543302     461534   1.177    0.239
## (Adjusted p values reported -- single-step method)
##
##
## Tukey results for gene: SPON1
##
##   Simultaneous Tests for General Linear Hypotheses
##
## Multiple Comparisons of Means: Tukey Contrasts
##
##
## Fit: glm(formula = Protein_Count ~ Tissue, family = gaussian, data = gene_
data)
##
## Linear Hypotheses:
##              Estimate Std. Error z value Pr(>|z|)
## CL - CF == 0  -195327    1060823  -0.184    0.854
## (Adjusted p values reported -- single-step method)
##
##
## Tukey results for gene: SRPX
##
##   Simultaneous Tests for General Linear Hypotheses
##
## Multiple Comparisons of Means: Tukey Contrasts
##
##
## Fit: glm(formula = Protein_Count ~ Tissue, family = gaussian, data = gene_
data)
##
## Linear Hypotheses:
##              Estimate Std. Error z value Pr(>|z|)
## CL - CF == 0    50277     135424   0.371     0.71
## (Adjusted p values reported -- single-step method)
##
##
## Tukey results for gene: SRPX2
##
##   Simultaneous Tests for General Linear Hypotheses
##
## Multiple Comparisons of Means: Tukey Contrasts
##
```

```
##
## Fit: glm(formula = Protein_Count ~ Tissue, family = gaussian, data = gene_
data)
##
## Linear Hypotheses:
##               Estimate Std. Error z value Pr(>|z|)
## CL - CF == 0  -285985     257103  -1.112    0.266
## (Adjusted p values reported -- single-step method)
##
##
## Tukey results for gene: SVEP1
##
##   Simultaneous Tests for General Linear Hypotheses
##
## Multiple Comparisons of Means: Tukey Contrasts
##
##
## Fit: glm(formula = Protein_Count ~ Tissue, family = gaussian, data = gene_
data)
##
## Linear Hypotheses:
##               Estimate Std. Error z value Pr(>|z|)
## CL - CF == 0   -14891     106945  -0.139    0.889
## (Adjusted p values reported -- single-step method)
##
##
## Tukey results for gene: TGFBI
##
##   Simultaneous Tests for General Linear Hypotheses
##
## Multiple Comparisons of Means: Tukey Contrasts
##
##
## Fit: glm(formula = Protein_Count ~ Tissue, family = gaussian, data = gene_
data)
##
## Linear Hypotheses:
##               Estimate Std. Error z value Pr(>|z|)
## CL - CF == 0 -1300552    3950557  -0.329    0.742
## (Adjusted p values reported -- single-step method)
##
##
## Tukey results for gene: THBS1
##
##   Simultaneous Tests for General Linear Hypotheses
##
## Multiple Comparisons of Means: Tukey Contrasts
##
##
## Fit: glm(formula = Protein_Count ~ Tissue, family = gaussian, data = gene_
```

```
data)
##
## Linear Hypotheses:
##               Estimate Std. Error z value Pr(>|z|)
## CL - CF == 0   -23887     542410  -0.044    0.965
## (Adjusted p values reported -- single-step method)
##
##
## Tukey results for gene: THBS2
##
##   Simultaneous Tests for General Linear Hypotheses
##
## Multiple Comparisons of Means: Tukey Contrasts
##
##
## Fit: glm(formula = Protein_Count ~ Tissue, family = gaussian, data = gene_
data)
##
## Linear Hypotheses:
##               Estimate Std. Error z value Pr(>|z|)
## CL - CF == 0 -2054484    2151392  -0.955     0.34
## (Adjusted p values reported -- single-step method)
##
##
## Tukey results for gene: THBS3
##
##   Simultaneous Tests for General Linear Hypotheses
##
## Multiple Comparisons of Means: Tukey Contrasts
##
##
## Fit: glm(formula = Protein_Count ~ Tissue, family = gaussian, data = gene_
data)
##
## Linear Hypotheses:
##               Estimate Std. Error z value Pr(>|z|)
## CL - CF == 0   -74273      53144  -1.398    0.162
## (Adjusted p values reported -- single-step method)
##
##
## Tukey results for gene: THSD4
##
##   Simultaneous Tests for General Linear Hypotheses
##
## Multiple Comparisons of Means: Tukey Contrasts
##
##
## Fit: glm(formula = Protein_Count ~ Tissue, family = gaussian, data = gene_
data)
##
```

```
## Linear Hypotheses:
##               Estimate Std. Error z value Pr(>|z|)
## CL - CF == 0   267126     886237   0.301    0.763
## (Adjusted p values reported -- single-step method)
##
##
## Tukey results for gene: TINAGL1
##
##   Simultaneous Tests for General Linear Hypotheses
##
## Multiple Comparisons of Means: Tukey Contrasts
##
##
## Fit: glm(formula = Protein_Count ~ Tissue, family = gaussian, data = gene_
data)
##
## Linear Hypotheses:
##                Estimate Std. Error z value Pr(>|z|)
## CL - CF == 0 -11871653   17031451  -0.697    0.486
## (Adjusted p values reported -- single-step method)
##
##
## Tukey results for gene: TNC
##
##   Simultaneous Tests for General Linear Hypotheses
##
## Multiple Comparisons of Means: Tukey Contrasts
##
##
## Fit: glm(formula = Protein_Count ~ Tissue, family = gaussian, data = gene_
data)
##
## Linear Hypotheses:
##               Estimate Std. Error z value Pr(>|z|)
## CL - CF == 0  -105047     343285  -0.306     0.76
## (Adjusted p values reported -- single-step method)
##
##
## Tukey results for gene: TNXB
##
##   Simultaneous Tests for General Linear Hypotheses
##
## Multiple Comparisons of Means: Tukey Contrasts
##
##
## Fit: glm(formula = Protein_Count ~ Tissue, family = gaussian, data = gene_
data)
##
## Linear Hypotheses:
##               Estimate Std. Error z value Pr(>|z|)
```

```
## CL - CF == 0   457585     343488   1.332    0.183
## (Adjusted p values reported -- single-step method)
##
##
## Tukey results for gene: VTN
##
##   Simultaneous Tests for General Linear Hypotheses
##
## Multiple Comparisons of Means: Tukey Contrasts
##
##
## Fit: glm(formula = Protein_Count ~ Tissue, family = gaussian, data = gene_
data)
##
## Linear Hypotheses:
##              Estimate Std. Error z value Pr(>|z|)
## CL - CF == 0 -1135137    1608232  -0.706     0.48
## (Adjusted p values reported -- single-step method)
##
##
## Tukey results for gene: VWA1
##
##   Simultaneous Tests for General Linear Hypotheses
##
## Multiple Comparisons of Means: Tukey Contrasts
##
##
## Fit: glm(formula = Protein_Count ~ Tissue, family = gaussian, data = gene_
data)
##
## Linear Hypotheses:
##              Estimate Std. Error z value Pr(>|z|)
## CL - CF == 0 -1035610    1773482  -0.584    0.559
## (Adjusted p values reported -- single-step method)
##
##
## Tukey results for gene: VWF
##
##   Simultaneous Tests for General Linear Hypotheses
##
## Multiple Comparisons of Means: Tukey Contrasts
##
##
## Fit: glm(formula = Protein_Count ~ Tissue, family = gaussian, data = gene_
data)
##
## Linear Hypotheses:
##              Estimate Std. Error z value Pr(>|z|)
## CL - CF == 0     6144      42810   0.144    0.886
## (Adjusted p values reported -- single-step method)
```

```
##
##
## Tukey results for gene: ZP3
##
##   Simultaneous Tests for General Linear Hypotheses
##
## Multiple Comparisons of Means: Tukey Contrasts
##
##
## Fit: glm(formula = Protein_Count ~ Tissue, family = gaussian, data = gene_
data)
##
## Linear Hypotheses:
##               Estimate Std. Error z value Pr(>|z|)
## CL - CF == 0     2862     843126   0.003    0.997
## (Adjusted p values reported -- single-step method)
##
##
## Tukey results for gene: ZP4
##
##   Simultaneous Tests for General Linear Hypotheses
##
## Multiple Comparisons of Means: Tukey Contrasts
##
##
## Fit: glm(formula = Protein_Count ~ Tissue, family = gaussian, data = gene_
data)
##
## Linear Hypotheses:
##               Estimate Std. Error z value Pr(>|z|)
## CL - CF == 0  -114853     645752  -0.178    0.859
## (Adjusted p values reported -- single-step method)
```

```
#### Proteoglycans

# Read your cleaned joined data file into a data frame
cleaned_joined_data <- read.csv("cleaned_matrisome_cortex.csv")

# Filter rows with Matrisome_Category "Proteoglycans"
Proteoglycans_data <- cleaned_joined_data %>%
  filter(Matrisome_Category == "Proteoglycans")

# Reshape the data to have separate columns for each replicate and each tissu
e stage
organized_data <- Proteoglycans_data %>%
  pivot_longer(cols = starts_with("CF") | starts_with("CL"),
               names_to = c("Tissue", "Stage"),
               names_sep = "\\.",
               values_to = "Protein_Count")
```

```
## Warning: Expected 2 pieces. Missing pieces filled with `NA` in 2 rows [1,
7].

# Save the reshaped data to a new file

write.csv(organized_data, "Proteoglycans_data_matrisome_cortex_to_clean.csv",
row.names = FALSE)

data2 <- read.csv("Proteoglycans_data_matrisome_cortex_clean.csv")

png(filename="cortex_Proteoglycans.png",
    width = 12, height = 6, units = "in",
    res = 600)

boxplot(Protein_Count ~ Tissue*Genes,
        data = data2,
        cex.axis = 0.8,
        col = viridis(2, alpha = 0.7),
        lwd = 0.7,
        ylab = "",
        ylim = c(0, 300000000),
        axes = FALSE,
        frame.plot = TRUE,
        xlab = "")

GROUP <- unique(data2$Genes)
groups <- 14
numbox <- 2
total <- groups * numbox
xpoints <- seq(median(1:numbox), total, numbox)

axis(2, seq(0, 300000000, by = 10000000), las = 1, cex.axis = 0.7)
axis(1, labels = GROUP, at = xpoints, las = 2, cex.axis = 0.7)

stripchart(data2$Protein_Count ~ data2$Tissue*data2$Genes,
           method = "jitter",
           pch = 19,
           col = "black",
           vertical = TRUE,
           add = TRUE,
           cex = 0.3)

title(ylab = expression("Protein abundance (RU)"), line = 3.0)
legend("top", fill = viridis(2), legend = c("Cortex Follicular", "Cortex Lute
al"), horiz = T, cex = 0.7)

dev.off()
```

```
## quartz_off_screen
##                 2

# Convert Tissue to factor
data2$Tissue <- as.factor(data2$Tissue)

# Create an empty list to store Tukey results
tukey_results_list <- list()

# Unique genes
unique_genes <- unique(data2$Genes)

# Loop over each gene
for (gene in unique_genes) {
  # Subset data for the current gene
  gene_data <- subset(data2, Genes == gene)

  # Fit the generalized linear model
  glm_model <- glm(Protein_Count ~ Tissue, data = gene_data, family = gaussia
n)

  # Perform Tukey test
  contrast_matrix <- glht(glm_model, linfct = mcp(Tissue = "Tukey"))
  tukey_results <- summary(contrast_matrix)

  # Store the results
  tukey_results_list[[gene]] <- tukey_results
}

# Print the results for each gene
for (gene in unique_genes) {
  cat("Tukey results for gene:", gene, "\n")
  print(tukey_results_list[[gene]])
  cat("\n")
}

## Tukey results for gene: ACAN
##
##   Simultaneous Tests for General Linear Hypotheses
##
## Multiple Comparisons of Means: Tukey Contrasts
##
##
## Fit: glm(formula = Protein_Count ~ Tissue, family = gaussian, data = gene_
data)
##
## Linear Hypotheses:
##               Estimate Std. Error z value Pr(>|z|)
## CL - CF == 0  -356790     373296  -0.956    0.339
## (Adjusted p values reported -- single-step method)
```

```
##
##
## Tukey results for gene: ASPN
##
##   Simultaneous Tests for General Linear Hypotheses
##
## Multiple Comparisons of Means: Tukey Contrasts
##
##
## Fit: glm(formula = Protein_Count ~ Tissue, family = gaussian, data = gene_
data)
##
## Linear Hypotheses:
##              Estimate Std. Error z value Pr(>|z|)
## CL - CF == 0  2062925    4033618   0.511    0.609
## (Adjusted p values reported -- single-step method)
##
##
## Tukey results for gene: BGN
##
##   Simultaneous Tests for General Linear Hypotheses
##
## Multiple Comparisons of Means: Tukey Contrasts
##
##
## Fit: glm(formula = Protein_Count ~ Tissue, family = gaussian, data = gene_
data)
##
## Linear Hypotheses:
##              Estimate Std. Error z value Pr(>|z|)
## CL - CF == 0    28555     297138   0.096    0.923
## (Adjusted p values reported -- single-step method)
##
##
## Tukey results for gene: DCN
##
##   Simultaneous Tests for General Linear Hypotheses
##
## Multiple Comparisons of Means: Tukey Contrasts
##
##
## Fit: glm(formula = Protein_Count ~ Tissue, family = gaussian, data = gene_
data)
##
## Linear Hypotheses:
##              Estimate Std. Error z value Pr(>|z|)
## CL - CF == 0   119012    1017912   0.117    0.907
## (Adjusted p values reported -- single-step method)
##
##
```

```
## Tukey results for gene: FMOD
##
##   Simultaneous Tests for General Linear Hypotheses
##
## Multiple Comparisons of Means: Tukey Contrasts
##
##
## Fit: glm(formula = Protein_Count ~ Tissue, family = gaussian, data = gene_
data)
##
## Linear Hypotheses:
##              Estimate Std. Error z value Pr(>|z|)
## CL - CF == 0 -6084703   13327346  -0.457    0.648
## (Adjusted p values reported -- single-step method)
##
##
## Tukey results for gene: HAPLN1
##
##   Simultaneous Tests for General Linear Hypotheses
##
## Multiple Comparisons of Means: Tukey Contrasts
##
##
## Fit: glm(formula = Protein_Count ~ Tissue, family = gaussian, data = gene_
data)
##
## Linear Hypotheses:
##              Estimate Std. Error z value Pr(>|z|)
## CL - CF == 0 -3282742    3518318  -0.933    0.351
## (Adjusted p values reported -- single-step method)
##
##
## Tukey results for gene: HAPLN3
##
##   Simultaneous Tests for General Linear Hypotheses
##
## Multiple Comparisons of Means: Tukey Contrasts
##
##
## Fit: glm(formula = Protein_Count ~ Tissue, family = gaussian, data = gene_
data)
##
## Linear Hypotheses:
##              Estimate Std. Error z value Pr(>|z|)
## CL - CF == 0 -1131853     836243  -1.353    0.176
## (Adjusted p values reported -- single-step method)
##
##
## Tukey results for gene: HSPG2
##
```

```
##   Simultaneous Tests for General Linear Hypotheses
##
## Multiple Comparisons of Means: Tukey Contrasts
##
##
## Fit: glm(formula = Protein_Count ~ Tissue, family = gaussian, data = gene_
data)
##
## Linear Hypotheses:
##               Estimate Std. Error z value Pr(>|z|)
## CL - CF == 0 -2436903    3494927  -0.697    0.486
## (Adjusted p values reported -- single-step method)
##
##
## Tukey results for gene: LUM
##
##   Simultaneous Tests for General Linear Hypotheses
##
## Multiple Comparisons of Means: Tukey Contrasts
##
##
## Fit: glm(formula = Protein_Count ~ Tissue, family = gaussian, data = gene_
data)
##
## Linear Hypotheses:
##               Estimate Std. Error z value Pr(>|z|)
## CL - CF == 0   730130    9721582   0.075     0.94
## (Adjusted p values reported -- single-step method)
##
##
## Tukey results for gene: OGN
##
##   Simultaneous Tests for General Linear Hypotheses
##
## Multiple Comparisons of Means: Tukey Contrasts
##
##
## Fit: glm(formula = Protein_Count ~ Tissue, family = gaussian, data = gene_
data)
##
## Linear Hypotheses:
##               Estimate Std. Error z value Pr(>|z|)
## CL - CF == 0 -6823051   10265358  -0.665    0.506
## (Adjusted p values reported -- single-step method)
##
##
## Tukey results for gene: PODN
##
##   Simultaneous Tests for General Linear Hypotheses
##
```

```
## Multiple Comparisons of Means: Tukey Contrasts
##
##
## Fit: glm(formula = Protein_Count ~ Tissue, family = gaussian, data = gene_
data)
##
## Linear Hypotheses:
##              Estimate Std. Error z value Pr(>|z|)
## CL - CF == 0    68681     173500   0.396    0.692
## (Adjusted p values reported -- single-step method)
##
##
## Tukey results for gene: PODNL1
##
##   Simultaneous Tests for General Linear Hypotheses
##
## Multiple Comparisons of Means: Tukey Contrasts
##
##
## Fit: glm(formula = Protein_Count ~ Tissue, family = gaussian, data = gene_
data)
##
## Linear Hypotheses:
##              Estimate Std. Error z value Pr(>|z|)
## CL - CF == 0    74658     113371   0.659     0.51
## (Adjusted p values reported -- single-step method)
##
##
## Tukey results for gene: PRELP
##
##   Simultaneous Tests for General Linear Hypotheses
##
## Multiple Comparisons of Means: Tukey Contrasts
##
##
## Fit: glm(formula = Protein_Count ~ Tissue, family = gaussian, data = gene_
data)
##
## Linear Hypotheses:
##              Estimate Std. Error z value Pr(>|z|)
## CL - CF == 0  2885686   16454285   0.175    0.861
## (Adjusted p values reported -- single-step method)
##
##
## Tukey results for gene: VCAN
##
##   Simultaneous Tests for General Linear Hypotheses
##
## Multiple Comparisons of Means: Tukey Contrasts
##
```

```
##
## Fit: glm(formula = Protein_Count ~ Tissue, family = gaussian, data = gene_
data)
##
## Linear Hypotheses:
##                Estimate Std. Error z value Pr(>|z|)
## CL - CF == 0 -7680733   52495280  -0.146    0.884
## (Adjusted p values reported -- single-step method)
```

```
#### ECM regulators

# Read your cleaned joined data file into a data frame
cleaned_joined_data <- read.csv("cleaned_matrisome_cortex.csv")

# Filter rows with Matrisome_Category "ECM regulators"
ECM_regulators_data <- cleaned_joined_data %>%
  filter(Matrisome_Category == "ECM Regulators")

# Reshape the data to have separate columns for each replicate and each tissu
e stage
organized_data <- ECM_regulators_data %>%
  pivot_longer(cols = starts_with("CF") | starts_with("CL"),
               names_to = c("Tissue", "Stage"),
               names_sep = "\\.",
               values_to = "Protein_Count")
```

```
## Warning: Expected 2 pieces. Missing pieces filled with `NA` in 2 rows [1,
7].
```

```
# Save the reshaped data to a new file

write.csv(organized_data, "ECM regulators_data_matrisome_cortex_to_clean.csv"
, row.names = FALSE)

data2 <- read.csv("ECM regulators_data_matrisome_cortex_clean.csv")

png(filename="cortex_ECM regulators.png",
    width = 12, height = 6, units = "in",
    res = 600)

boxplot(Protein_Count ~ Tissue*Genes,
        data = data2,
        cex.axis = 0.8,
        col = viridis(2, alpha = 0.7),
        lwd = 0.7,
        ylab = "",
        ylim = c(0, 300000000),
        axes = FALSE,
        frame.plot = TRUE,
        xlab = "")
```

```
GROUP <- unique(data2$Genes)
groups <- 52
numbox <- 2
total <- groups * numbox
xpoints <- seq(median(1:numbox), total, numbox)

axis(2, seq(0, 300000000, by = 10000000), las = 1, cex.axis = 0.7)
axis(1, labels = GROUP, at = xpoints, las = 2, cex.axis = 0.7)

stripchart(data2$Protein_Count ~ data2$Tissue*data2$Genes,
           method = "jitter",
           pch = 19,
           col = "black",
           vertical = TRUE,
           add = TRUE,
           cex = 0.3)

title(ylab = expression("Protein abundance (RU)"), line = 3.0)
legend("top", fill = viridis(2), legend = c("Cortex Follicular", "Cortex Lute
al"), horiz = T, cex = 0.7)

dev.off()

## quartz_off_screen
##                 2

# Convert Tissue to factor
data2$Tissue <- as.factor(data2$Tissue)

# Create an empty list to store Tukey results
tukey_results_list <- list()

# Unique genes
unique_genes <- unique(data2$Genes)

# Loop over each gene
for (gene in unique_genes) {
  # Subset data for the current gene
  gene_data <- subset(data2, Genes == gene)

  # Fit the generalized linear model
  glm_model <- glm(Protein_Count ~ Tissue, data = gene_data, family = gaussia
n)

  # Perform Tukey test
  contrast_matrix <- glht(glm_model, linfct = mcp(Tissue = "Tukey"))
  tukey_results <- summary(contrast_matrix)
```

```
  # Store the results
  tukey_results_list[[gene]] <- tukey_results
}

# Print the results for each gene
for (gene in unique_genes) {
  cat("Tukey results for gene:", gene, "\n")
  print(tukey_results_list[[gene]])
  cat("\n")
}

## Tukey results for gene: A2M
##
##   Simultaneous Tests for General Linear Hypotheses
##
## Multiple Comparisons of Means: Tukey Contrasts
##
##
## Fit: glm(formula = Protein_Count ~ Tissue, family = gaussian, data = gene_
data)
##
## Linear Hypotheses:
##               Estimate Std. Error z value Pr(>|z|)
## CL - CF == 0    -38625     526884  -0.073    0.942
## (Adjusted p values reported -- single-step method)
##
##
## Tukey results for gene: ADAMTSL4
##
##   Simultaneous Tests for General Linear Hypotheses
##
## Multiple Comparisons of Means: Tukey Contrasts
##
##
## Fit: glm(formula = Protein_Count ~ Tissue, family = gaussian, data = gene_
data)
##
## Linear Hypotheses:
##               Estimate Std. Error z value Pr(>|z|)
## CL - CF == 0    -89188     167807  -0.531    0.595
## (Adjusted p values reported -- single-step method)
##
##
## Tukey results for gene: ADAMTSL5
##
##   Simultaneous Tests for General Linear Hypotheses
##
## Multiple Comparisons of Means: Tukey Contrasts
##
##
```

```
## Fit: glm(formula = Protein_Count ~ Tissue, family = gaussian, data = gene_
data)
##
## Linear Hypotheses:
##              Estimate Std. Error z value Pr(>|z|)
## CL - CF == 0  -136751     124463  -1.099    0.272
## (Adjusted p values reported -- single-step method)
##
##
## Tukey results for gene: AGT
##
##   Simultaneous Tests for General Linear Hypotheses
##
## Multiple Comparisons of Means: Tukey Contrasts
##
##
## Fit: glm(formula = Protein_Count ~ Tissue, family = gaussian, data = gene_
data)
##
## Linear Hypotheses:
##              Estimate Std. Error z value Pr(>|z|)
## CL - CF == 0   -13047     138929  -0.094    0.925
## (Adjusted p values reported -- single-step method)
##
##
## Tukey results for gene: CELA3B
##
##   Simultaneous Tests for General Linear Hypotheses
##
## Multiple Comparisons of Means: Tukey Contrasts
##
##
## Fit: glm(formula = Protein_Count ~ Tissue, family = gaussian, data = gene_
data)
##
## Linear Hypotheses:
##              Estimate Std. Error z value Pr(>|z|)
## CL - CF == 0  1808461    3058909   0.591    0.554
## (Adjusted p values reported -- single-step method)
##
##
## Tukey results for gene: CSTB
##
##   Simultaneous Tests for General Linear Hypotheses
##
## Multiple Comparisons of Means: Tukey Contrasts
##
##
## Fit: glm(formula = Protein_Count ~ Tissue, family = gaussian, data = gene_
data)
```

```
##
## Linear Hypotheses:
##               Estimate Std. Error z value Pr(>|z|)
## CL - CF == 0  -130064     375688  -0.346    0.729
## (Adjusted p values reported -- single-step method)
##
##
## Tukey results for gene: CTSD
##
##   Simultaneous Tests for General Linear Hypotheses
##
## Multiple Comparisons of Means: Tukey Contrasts
##
##
## Fit: glm(formula = Protein_Count ~ Tissue, family = gaussian, data = gene_
data)
##
## Linear Hypotheses:
##               Estimate Std. Error z value Pr(>|z|)
## CL - CF == 0    13779     315451   0.044    0.965
## (Adjusted p values reported -- single-step method)
##
##
## Tukey results for gene: CTSF
##
##   Simultaneous Tests for General Linear Hypotheses
##
## Multiple Comparisons of Means: Tukey Contrasts
##
##
## Fit: glm(formula = Protein_Count ~ Tissue, family = gaussian, data = gene_
data)
##
## Linear Hypotheses:
##               Estimate Std. Error z value Pr(>|z|)
## CL - CF == 0    31379     310803   0.101     0.92
## (Adjusted p values reported -- single-step method)
##
##
## Tukey results for gene: CTSH
##
##   Simultaneous Tests for General Linear Hypotheses
##
## Multiple Comparisons of Means: Tukey Contrasts
##
##
## Fit: glm(formula = Protein_Count ~ Tissue, family = gaussian, data = gene_
data)
##
## Linear Hypotheses:
```

```
##               Estimate Std. Error z value Pr(>|z|)
## CL - CF == 0   -97954     173184  -0.566    0.572
## (Adjusted p values reported -- single-step method)
##
##
## Tukey results for gene: CTSZ
##
##   Simultaneous Tests for General Linear Hypotheses
##
## Multiple Comparisons of Means: Tukey Contrasts
##
##
## Fit: glm(formula = Protein_Count ~ Tissue, family = gaussian, data = gene_
data)
##
## Linear Hypotheses:
##               Estimate Std. Error z value Pr(>|z|)
## CL - CF == 0  -104979     131867  -0.796    0.426
## (Adjusted p values reported -- single-step method)
##
##
## Tukey results for gene: F10
##
##   Simultaneous Tests for General Linear Hypotheses
##
## Multiple Comparisons of Means: Tukey Contrasts
##
##
## Fit: glm(formula = Protein_Count ~ Tissue, family = gaussian, data = gene_
data)
##
## Linear Hypotheses:
##               Estimate Std. Error z value Pr(>|z|)
## CL - CF == 0   -57062     165890  -0.344    0.731
## (Adjusted p values reported -- single-step method)
##
##
## Tukey results for gene: F13B
##
##   Simultaneous Tests for General Linear Hypotheses
##
## Multiple Comparisons of Means: Tukey Contrasts
##
##
## Fit: glm(formula = Protein_Count ~ Tissue, family = gaussian, data = gene_
data)
##
## Linear Hypotheses:
##               Estimate Std. Error z value Pr(>|z|)
## CL - CF == 0  -202685      96664  -2.097    0.036 *
```

```
## ---
## Signif. codes:  0 '***' 0.001 '**' 0.01 '*' 0.05 '.' 0.1 ' ' 1
## (Adjusted p values reported -- single-step method)
##
##
## Tukey results for gene: F2
##
##   Simultaneous Tests for General Linear Hypotheses
##
## Multiple Comparisons of Means: Tukey Contrasts
##
##
## Fit: glm(formula = Protein_Count ~ Tissue, family = gaussian, data = gene_
data)
##
## Linear Hypotheses:
##              Estimate Std. Error z value Pr(>|z|)
## CL - CF == 0  -222621     189805  -1.173    0.241
## (Adjusted p values reported -- single-step method)
##
##
## Tukey results for gene: F9
##
##   Simultaneous Tests for General Linear Hypotheses
##
## Multiple Comparisons of Means: Tukey Contrasts
##
##
## Fit: glm(formula = Protein_Count ~ Tissue, family = gaussian, data = gene_
data)
##
## Linear Hypotheses:
##              Estimate Std. Error z value Pr(>|z|)
## CL - CF == 0  -121692     168252  -0.723     0.47
## (Adjusted p values reported -- single-step method)
##
##
## Tukey results for gene: HABP2
##
##   Simultaneous Tests for General Linear Hypotheses
##
## Multiple Comparisons of Means: Tukey Contrasts
##
##
## Fit: glm(formula = Protein_Count ~ Tissue, family = gaussian, data = gene_
data)
##
## Linear Hypotheses:
##              Estimate Std. Error z value Pr(>|z|)
## CL - CF == 0   -24359      53083  -0.459    0.646
```

```
## (Adjusted p values reported -- single-step method)
##
##
## Tukey results for gene: HRG
##
##   Simultaneous Tests for General Linear Hypotheses
##
## Multiple Comparisons of Means: Tukey Contrasts
##
##
## Fit: glm(formula = Protein_Count ~ Tissue, family = gaussian, data = gene_
data)
##
## Linear Hypotheses:
##              Estimate Std. Error z value Pr(>|z|)
## CL - CF == 0   715139     789986   0.905    0.365
## (Adjusted p values reported -- single-step method)
##
##
## Tukey results for gene: HTRA1
##
##   Simultaneous Tests for General Linear Hypotheses
##
## Multiple Comparisons of Means: Tukey Contrasts
##
##
## Fit: glm(formula = Protein_Count ~ Tissue, family = gaussian, data = gene_
data)
##
## Linear Hypotheses:
##              Estimate Std. Error z value Pr(>|z|)
## CL - CF == 0  -109641     150187   -0.73    0.465
## (Adjusted p values reported -- single-step method)
##
##
## Tukey results for gene: ITIH1
##
##   Simultaneous Tests for General Linear Hypotheses
##
## Multiple Comparisons of Means: Tukey Contrasts
##
##
## Fit: glm(formula = Protein_Count ~ Tissue, family = gaussian, data = gene_
data)
##
## Linear Hypotheses:
##              Estimate Std. Error z value Pr(>|z|)
## CL - CF == 0 -1442186     989304  -1.458    0.145
## (Adjusted p values reported -- single-step method)
##
```

```
##
## Tukey results for gene: ITIH2
##
##   Simultaneous Tests for General Linear Hypotheses
##
## Multiple Comparisons of Means: Tukey Contrasts
##
##
## Fit: glm(formula = Protein_Count ~ Tissue, family = gaussian, data = gene_
data)
##
## Linear Hypotheses:
##              Estimate Std. Error z value Pr(>|z|)
## CL - CF == 0  -193933     362185  -0.535    0.592
## (Adjusted p values reported -- single-step method)
##
##
## Tukey results for gene: ITIH3
##
##   Simultaneous Tests for General Linear Hypotheses
##
## Multiple Comparisons of Means: Tukey Contrasts
##
##
## Fit: glm(formula = Protein_Count ~ Tissue, family = gaussian, data = gene_
data)
##
## Linear Hypotheses:
##              Estimate Std. Error z value Pr(>|z|)
## CL - CF == 0   -85136      89981  -0.946    0.344
## (Adjusted p values reported -- single-step method)
##
##
## Tukey results for gene: ITIH4
##
##   Simultaneous Tests for General Linear Hypotheses
##
## Multiple Comparisons of Means: Tukey Contrasts
##
##
## Fit: glm(formula = Protein_Count ~ Tissue, family = gaussian, data = gene_
data)
##
## Linear Hypotheses:
##              Estimate Std. Error z value Pr(>|z|)
## CL - CF == 0 -1986720    1700484  -1.168    0.243
## (Adjusted p values reported -- single-step method)
##
##
## Tukey results for gene: ITIH5
```

```
##
##   Simultaneous Tests for General Linear Hypotheses
##
## Multiple Comparisons of Means: Tukey Contrasts
##
##
## Fit: glm(formula = Protein_Count ~ Tissue, family = gaussian, data = gene_
data)
##
## Linear Hypotheses:
##              Estimate Std. Error z value Pr(>|z|)
## CL - CF == 0    63871     128409   0.497    0.619
## (Adjusted p values reported -- single-step method)
##
##
## Tukey results for gene: KNG1
##
##   Simultaneous Tests for General Linear Hypotheses
##
## Multiple Comparisons of Means: Tukey Contrasts
##
##
## Fit: glm(formula = Protein_Count ~ Tissue, family = gaussian, data = gene_
data)
##
## Linear Hypotheses:
##              Estimate Std. Error z value Pr(>|z|)
## CL - CF == 0    31612      72249   0.438    0.662
## (Adjusted p values reported -- single-step method)
##
##
## Tukey results for gene: LOX
##
##   Simultaneous Tests for General Linear Hypotheses
##
## Multiple Comparisons of Means: Tukey Contrasts
##
##
## Fit: glm(formula = Protein_Count ~ Tissue, family = gaussian, data = gene_
data)
##
## Linear Hypotheses:
##              Estimate Std. Error z value Pr(>|z|)
## CL - CF == 0   879608    3719972   0.236    0.813
## (Adjusted p values reported -- single-step method)
##
##
## Tukey results for gene: LOXL1
##
##   Simultaneous Tests for General Linear Hypotheses
```

```
##
## Multiple Comparisons of Means: Tukey Contrasts
##
##
## Fit: glm(formula = Protein_Count ~ Tissue, family = gaussian, data = gene_
data)
##
## Linear Hypotheses:
##              Estimate Std. Error z value Pr(>|z|)
## CL - CF == 0  1091276    1965674   0.555    0.579
## (Adjusted p values reported -- single-step method)
##
##
## Tukey results for gene: LOXL3
##
##   Simultaneous Tests for General Linear Hypotheses
##
## Multiple Comparisons of Means: Tukey Contrasts
##
##
## Fit: glm(formula = Protein_Count ~ Tissue, family = gaussian, data = gene_
data)
##
## Linear Hypotheses:
##              Estimate Std. Error z value Pr(>|z|)
## CL - CF == 0    -9549      58212  -0.164     0.87
## (Adjusted p values reported -- single-step method)
##
##
## Tukey results for gene: MASP2
##
##   Simultaneous Tests for General Linear Hypotheses
##
## Multiple Comparisons of Means: Tukey Contrasts
##
##
## Fit: glm(formula = Protein_Count ~ Tissue, family = gaussian, data = gene_
data)
##
## Linear Hypotheses:
##              Estimate Std. Error z value Pr(>|z|)
## CL - CF == 0  1475626    1076683   1.371    0.171
## (Adjusted p values reported -- single-step method)
##
##
## Tukey results for gene: MMP14
##
##   Simultaneous Tests for General Linear Hypotheses
##
## Multiple Comparisons of Means: Tukey Contrasts
```

```
##
##
## Fit: glm(formula = Protein_Count ~ Tissue, family = gaussian, data = gene_
data)
##
## Linear Hypotheses:
##              Estimate Std. Error z value Pr(>|z|)
## CL - CF == 0    46437     226348   0.205    0.837
## (Adjusted p values reported -- single-step method)
##
##
## Tukey results for gene: MMP2
##
##   Simultaneous Tests for General Linear Hypotheses
##
## Multiple Comparisons of Means: Tukey Contrasts
##
##
## Fit: glm(formula = Protein_Count ~ Tissue, family = gaussian, data = gene_
data)
##
## Linear Hypotheses:
##              Estimate Std. Error z value Pr(>|z|)
## CL - CF == 0  -111194     224935  -0.494    0.621
## (Adjusted p values reported -- single-step method)
##
##
## Tukey results for gene: MMP23
##
##   Simultaneous Tests for General Linear Hypotheses
##
## Multiple Comparisons of Means: Tukey Contrasts
##
##
## Fit: glm(formula = Protein_Count ~ Tissue, family = gaussian, data = gene_
data)
##
## Linear Hypotheses:
##              Estimate Std. Error z value Pr(>|z|)
## CL - CF == 0    16510      92332   0.179    0.858
## (Adjusted p values reported -- single-step method)
##
##
## Tukey results for gene: MMP3
##
##   Simultaneous Tests for General Linear Hypotheses
##
## Multiple Comparisons of Means: Tukey Contrasts
##
##
```

```
## Fit: glm(formula = Protein_Count ~ Tissue, family = gaussian, data = gene_
data)
##
## Linear Hypotheses:
##              Estimate Std. Error z value Pr(>|z|)
## CL - CF == 0   -70012     144661  -0.484    0.628
## (Adjusted p values reported -- single-step method)
##
##
## Tukey results for gene: P4HA2
##
##   Simultaneous Tests for General Linear Hypotheses
##
## Multiple Comparisons of Means: Tukey Contrasts
##
##
## Fit: glm(formula = Protein_Count ~ Tissue, family = gaussian, data = gene_
data)
##
## Linear Hypotheses:
##              Estimate Std. Error z value Pr(>|z|)
## CL - CF == 0    45951     123080   0.373    0.709
## (Adjusted p values reported -- single-step method)
##
##
## Tukey results for gene: PLG
##
##   Simultaneous Tests for General Linear Hypotheses
##
## Multiple Comparisons of Means: Tukey Contrasts
##
##
## Fit: glm(formula = Protein_Count ~ Tissue, family = gaussian, data = gene_
data)
##
## Linear Hypotheses:
##              Estimate Std. Error z value Pr(>|z|)
## CL - CF == 0  -200232     826490  -0.242    0.809
## (Adjusted p values reported -- single-step method)
##
##
## Tukey results for gene: PLOD1
##
##   Simultaneous Tests for General Linear Hypotheses
##
## Multiple Comparisons of Means: Tukey Contrasts
##
##
## Fit: glm(formula = Protein_Count ~ Tissue, family = gaussian, data = gene_
data)
```

```
##
## Linear Hypotheses:
##               Estimate Std. Error z value Pr(>|z|)
## CL - CF == 0  -243974     293334  -0.832    0.406
## (Adjusted p values reported -- single-step method)
##
##
## Tukey results for gene: PLOD3
##
##   Simultaneous Tests for General Linear Hypotheses
##
## Multiple Comparisons of Means: Tukey Contrasts
##
##
## Fit: glm(formula = Protein_Count ~ Tissue, family = gaussian, data = gene_
data)
##
## Linear Hypotheses:
##               Estimate Std. Error z value Pr(>|z|)
## CL - CF == 0    15724     114931   0.137    0.891
## (Adjusted p values reported -- single-step method)
##
##
## Tukey results for gene: PRSS1
##
##   Simultaneous Tests for General Linear Hypotheses
##
## Multiple Comparisons of Means: Tukey Contrasts
##
##
## Fit: glm(formula = Protein_Count ~ Tissue, family = gaussian, data = gene_
data)
##
## Linear Hypotheses:
##               Estimate Std. Error z value Pr(>|z|)
## CL - CF == 0   750049    1921093    0.39    0.696
## (Adjusted p values reported -- single-step method)
##
##
## Tukey results for gene: SERPINA1
##
##   Simultaneous Tests for General Linear Hypotheses
##
## Multiple Comparisons of Means: Tukey Contrasts
##
##
## Fit: glm(formula = Protein_Count ~ Tissue, family = gaussian, data = gene_
data)
##
## Linear Hypotheses:
```

```
##                Estimate Std. Error z value Pr(>|z|)
## CL - CF == 0    11296     161084    0.07    0.944
## (Adjusted p values reported -- single-step method)
##
##
## Tukey results for gene: SERPINA10
##
##   Simultaneous Tests for General Linear Hypotheses
##
## Multiple Comparisons of Means: Tukey Contrasts
##
##
## Fit: glm(formula = Protein_Count ~ Tissue, family = gaussian, data = gene_
data)
##
## Linear Hypotheses:
##                Estimate Std. Error z value Pr(>|z|)
## CL - CF == 0   -71163      77100  -0.923    0.356
## (Adjusted p values reported -- single-step method)
##
##
## Tukey results for gene: SERPINA3-8
##
##   Simultaneous Tests for General Linear Hypotheses
##
## Multiple Comparisons of Means: Tukey Contrasts
##
##
## Fit: glm(formula = Protein_Count ~ Tissue, family = gaussian, data = gene_
data)
##
## Linear Hypotheses:
##                Estimate Std. Error z value Pr(>|z|)
## CL - CF == 0   -43530      59634   -0.73    0.465
## (Adjusted p values reported -- single-step method)
##
##
## Tukey results for gene: SERPINA5
##
##   Simultaneous Tests for General Linear Hypotheses
##
## Multiple Comparisons of Means: Tukey Contrasts
##
##
## Fit: glm(formula = Protein_Count ~ Tissue, family = gaussian, data = gene_
data)
##
## Linear Hypotheses:
##                Estimate Std. Error z value Pr(>|z|)
## CL - CF == 0   111190     217524   0.511    0.609
```

```
## (Adjusted p values reported -- single-step method)
##
##
## Tukey results for gene: SERPINB6
##
##   Simultaneous Tests for General Linear Hypotheses
##
## Multiple Comparisons of Means: Tukey Contrasts
##
##
## Fit: glm(formula = Protein_Count ~ Tissue, family = gaussian, data = gene_
data)
##
## Linear Hypotheses:
##              Estimate Std. Error z value Pr(>|z|)
## CL - CF == 0    17585     195494    0.09    0.928
## (Adjusted p values reported -- single-step method)
##
##
## Tukey results for gene: SERPINC1
##
##   Simultaneous Tests for General Linear Hypotheses
##
## Multiple Comparisons of Means: Tukey Contrasts
##
##
## Fit: glm(formula = Protein_Count ~ Tissue, family = gaussian, data = gene_
data)
##
## Linear Hypotheses:
##              Estimate Std. Error z value Pr(>|z|)
## CL - CF == 0   147010     672582   0.219    0.827
## (Adjusted p values reported -- single-step method)
##
##
## Tukey results for gene: SERPIND1
##
##   Simultaneous Tests for General Linear Hypotheses
##
## Multiple Comparisons of Means: Tukey Contrasts
##
##
## Fit: glm(formula = Protein_Count ~ Tissue, family = gaussian, data = gene_
data)
##
## Linear Hypotheses:
##              Estimate Std. Error z value Pr(>|z|)
## CL - CF == 0   -31269     194626  -0.161    0.872
## (Adjusted p values reported -- single-step method)
##
```

```
##
## Tukey results for gene: SERPINE2
##
##   Simultaneous Tests for General Linear Hypotheses
##
## Multiple Comparisons of Means: Tukey Contrasts
##
##
## Fit: glm(formula = Protein_Count ~ Tissue, family = gaussian, data = gene_
data)
##
## Linear Hypotheses:
##              Estimate Std. Error z value Pr(>|z|)
## CL - CF == 0    12849     102632   0.125      0.9
## (Adjusted p values reported -- single-step method)
##
##
## Tukey results for gene: SERPINF1
##
##   Simultaneous Tests for General Linear Hypotheses
##
## Multiple Comparisons of Means: Tukey Contrasts
##
##
## Fit: glm(formula = Protein_Count ~ Tissue, family = gaussian, data = gene_
data)
##
## Linear Hypotheses:
##              Estimate Std. Error z value Pr(>|z|)
## CL - CF == 0    34218     585426   0.058    0.953
## (Adjusted p values reported -- single-step method)
##
##
## Tukey results for gene: SERPINF2
##
##   Simultaneous Tests for General Linear Hypotheses
##
## Multiple Comparisons of Means: Tukey Contrasts
##
##
## Fit: glm(formula = Protein_Count ~ Tissue, family = gaussian, data = gene_
data)
##
## Linear Hypotheses:
##              Estimate Std. Error z value Pr(>|z|)
## CL - CF == 0   -16528     166753  -0.099    0.921
## (Adjusted p values reported -- single-step method)
##
##
## Tukey results for gene: SERPING1
```

```
##
##   Simultaneous Tests for General Linear Hypotheses
##
## Multiple Comparisons of Means: Tukey Contrasts
##
##
## Fit: glm(formula = Protein_Count ~ Tissue, family = gaussian, data = gene_
data)
##
## Linear Hypotheses:
##               Estimate Std. Error z value Pr(>|z|)
## CL - CF == 0   -38714     206582  -0.187    0.851
## (Adjusted p values reported -- single-step method)
##
##
## Tukey results for gene: SERPINH1
##
##   Simultaneous Tests for General Linear Hypotheses
##
## Multiple Comparisons of Means: Tukey Contrasts
##
##
## Fit: glm(formula = Protein_Count ~ Tissue, family = gaussian, data = gene_
data)
##
## Linear Hypotheses:
##               Estimate Std. Error z value Pr(>|z|)
## CL - CF == 0   727990    4787748   0.152    0.879
## (Adjusted p values reported -- single-step method)
##
##
## Tukey results for gene: SERPINI2
##
##   Simultaneous Tests for General Linear Hypotheses
##
## Multiple Comparisons of Means: Tukey Contrasts
##
##
## Fit: glm(formula = Protein_Count ~ Tissue, family = gaussian, data = gene_
data)
##
## Linear Hypotheses:
##               Estimate Std. Error z value Pr(>|z|)
## CL - CF == 0    12881     184299    0.07    0.944
## (Adjusted p values reported -- single-step method)
##
##
## Tukey results for gene: TGM1
##
##   Simultaneous Tests for General Linear Hypotheses
```

```
##
## Multiple Comparisons of Means: Tukey Contrasts
##
##
## Fit: glm(formula = Protein_Count ~ Tissue, family = gaussian, data = gene_
data)
##
## Linear Hypotheses:
##                Estimate Std. Error z value Pr(>|z|)
## CL - CF == 0   -83555     427056  -0.196    0.845
## (Adjusted p values reported -- single-step method)
##
##
## Tukey results for gene: TGM2
##
##   Simultaneous Tests for General Linear Hypotheses
##
## Multiple Comparisons of Means: Tukey Contrasts
##
##
## Fit: glm(formula = Protein_Count ~ Tissue, family = gaussian, data = gene_
data)
##
## Linear Hypotheses:
##                 Estimate Std. Error z value Pr(>|z|)
## CL - CF == 0 -15958007   39115540  -0.408    0.683
## (Adjusted p values reported -- single-step method)
##
##
## Tukey results for gene: TIMP2
##
##   Simultaneous Tests for General Linear Hypotheses
##
## Multiple Comparisons of Means: Tukey Contrasts
##
##
## Fit: glm(formula = Protein_Count ~ Tissue, family = gaussian, data = gene_
data)
##
## Linear Hypotheses:
##                Estimate Std. Error z value Pr(>|z|)
## CL - CF == 0     4828      94952   0.051    0.959
## (Adjusted p values reported -- single-step method)

#### ECM-affiliated

# Read your cleaned joined data file into a data frame
cleaned_joined_data <- read.csv("cleaned_matrisome_cortex.csv")

# Filter rows with Matrisome_Category "ECM-affiliated"
```

```
ECMaffiliated_data <- cleaned_joined_data %>%
  filter(Matrisome_Category == "ECM-affiliated")

# Reshape the data to have separate columns for each replicate and each tissu
e stage
organized_data <- ECMaffiliated_data %>%
  pivot_longer(cols = starts_with("CF") | starts_with("CL"),
               names_to = c("Tissue", "Stage"),
               names_sep = "\\.",
               values_to = "Protein_Count")

## Warning: Expected 2 pieces. Missing pieces filled with `NA` in 2 rows [1,
7].

# Save the reshaped data to a new file

write.csv(organized_data, "ECM-affiliated_data_matrisome_cortex_to_clean.csv"
, row.names = FALSE)

data2 <- read.csv("ECM-affiliated_data_matrisome_cortex_clean.csv")

png(filename="cortex_ECM-affiliated.png",
    width = 12, height = 6, units = "in",
    res = 600)

boxplot(Protein_Count ~ Tissue*Genes,
        data = data2,
        cex.axis = 0.8,
        col = viridis(2, alpha = 0.7),
        lwd = 0.7,
        ylab = "",
        ylim = c(0, 60000000),
        axes = FALSE,
        frame.plot = TRUE,
        xlab = "")

GROUP <- unique(data2$Genes)
groups <- 22
numbox <- 2
total <- groups * numbox
xpoints <- seq(median(1:numbox), total, numbox)

axis(2, seq(0, 60000000, by = 1000000), las = 1, cex.axis = 0.7)
axis(1, labels = GROUP, at = xpoints, las = 2, cex.axis = 0.7)

stripchart(data2$Protein_Count ~ data2$Tissue*data2$Genes,
           method = "jitter",
           pch = 19,
           col = "black",
           vertical = TRUE,
```

```
           add = TRUE,
           cex = 0.3)

title(ylab = expression("Protein abundance (RU)"), line = 3.0)
legend("top", fill = viridis(2), legend = c("Cortex Follicular", "Cortex Lute
al"), horiz = T, cex = 0.7)

dev.off()

## quartz_off_screen
##                  2

# Convert Tissue to factor
data2$Tissue <- as.factor(data2$Tissue)

# Create an empty list to store Tukey results
tukey_results_list <- list()

# Unique genes
unique_genes <- unique(data2$Genes)

# Loop over each gene
for (gene in unique_genes) {
  # Subset data for the current gene
  gene_data <- subset(data2, Genes == gene)

  # Fit the generalized linear model
  glm_model <- glm(Protein_Count ~ Tissue, data = gene_data, family = gaussia
n)

  # Perform Tukey test
  contrast_matrix <- glht(glm_model, linfct = mcp(Tissue = "Tukey"))
  tukey_results <- summary(contrast_matrix)

  # Store the results
  tukey_results_list[[gene]] <- tukey_results
}

# Print the results for each gene
for (gene in unique_genes) {
  cat("Tukey results for gene:", gene, "\n")
  print(tukey_results_list[[gene]])
  cat("\n")
}

## Tukey results for gene: ANXA1
##
##   Simultaneous Tests for General Linear Hypotheses
##
## Multiple Comparisons of Means: Tukey Contrasts
```

```
##
##
## Fit: glm(formula = Protein_Count ~ Tissue, family = gaussian, data = gene_
data)
##
## Linear Hypotheses:
##              Estimate Std. Error z value Pr(>|z|)
## CL - CF == 0   -33938     910363  -0.037     0.97
## (Adjusted p values reported -- single-step method)
##
##
## Tukey results for gene: ANXA11
##
##   Simultaneous Tests for General Linear Hypotheses
##
## Multiple Comparisons of Means: Tukey Contrasts
##
##
## Fit: glm(formula = Protein_Count ~ Tissue, family = gaussian, data = gene_
data)
##
## Linear Hypotheses:
##              Estimate Std. Error z value Pr(>|z|)
## CL - CF == 0  -133414     389275  -0.343    0.732
## (Adjusted p values reported -- single-step method)
##
##
## Tukey results for gene: ANXA2
##
##   Simultaneous Tests for General Linear Hypotheses
##
## Multiple Comparisons of Means: Tukey Contrasts
##
##
## Fit: glm(formula = Protein_Count ~ Tissue, family = gaussian, data = gene_
data)
##
## Linear Hypotheses:
##              Estimate Std. Error z value Pr(>|z|)
## CL - CF == 0  -569451    3770264  -0.151     0.88
## (Adjusted p values reported -- single-step method)
##
##
## Tukey results for gene: ANXA3
##
##   Simultaneous Tests for General Linear Hypotheses
##
## Multiple Comparisons of Means: Tukey Contrasts
##
##
```

```
## Fit: glm(formula = Protein_Count ~ Tissue, family = gaussian, data = gene_
data)
##
## Linear Hypotheses:
##              Estimate Std. Error z value Pr(>|z|)
## CL - CF == 0   -88796     218376  -0.407    0.684
## (Adjusted p values reported -- single-step method)
##
##
## Tukey results for gene: ANXA4
##
##   Simultaneous Tests for General Linear Hypotheses
##
## Multiple Comparisons of Means: Tukey Contrasts
##
##
## Fit: glm(formula = Protein_Count ~ Tissue, family = gaussian, data = gene_
data)
##
## Linear Hypotheses:
##              Estimate Std. Error z value Pr(>|z|)
## CL - CF == 0   -29513     524486  -0.056    0.955
## (Adjusted p values reported -- single-step method)
##
##
## Tukey results for gene: ANXA5
##
##   Simultaneous Tests for General Linear Hypotheses
##
## Multiple Comparisons of Means: Tukey Contrasts
##
##
## Fit: glm(formula = Protein_Count ~ Tissue, family = gaussian, data = gene_
data)
##
## Linear Hypotheses:
##              Estimate Std. Error z value Pr(>|z|)
## CL - CF == 0 -4869888   10506262  -0.464    0.643
## (Adjusted p values reported -- single-step method)
##
##
## Tukey results for gene: ANXA6
##
##   Simultaneous Tests for General Linear Hypotheses
##
## Multiple Comparisons of Means: Tukey Contrasts
##
##
## Fit: glm(formula = Protein_Count ~ Tissue, family = gaussian, data = gene_
data)
```

```
##
## Linear Hypotheses:
##               Estimate Std. Error z value Pr(>|z|)
## CL - CF == 0  -295012     682902  -0.432    0.666
## (Adjusted p values reported -- single-step method)
##
##
## Tukey results for gene: ANXA7
##
##   Simultaneous Tests for General Linear Hypotheses
##
## Multiple Comparisons of Means: Tukey Contrasts
##
##
## Fit: glm(formula = Protein_Count ~ Tissue, family = gaussian, data = gene_
data)
##
## Linear Hypotheses:
##               Estimate Std. Error z value Pr(>|z|)
## CL - CF == 0   -22209     247644   -0.09    0.929
## (Adjusted p values reported -- single-step method)
##
##
## Tukey results for gene: C1QTNF3
##
##   Simultaneous Tests for General Linear Hypotheses
##
## Multiple Comparisons of Means: Tukey Contrasts
##
##
## Fit: glm(formula = Protein_Count ~ Tissue, family = gaussian, data = gene_
data)
##
## Linear Hypotheses:
##               Estimate Std. Error z value Pr(>|z|)
## CL - CF == 0     8078      37840   0.213    0.831
## (Adjusted p values reported -- single-step method)
##
##
## Tukey results for gene: C1QTNF5
##
##   Simultaneous Tests for General Linear Hypotheses
##
## Multiple Comparisons of Means: Tukey Contrasts
##
##
## Fit: glm(formula = Protein_Count ~ Tissue, family = gaussian, data = gene_
data)
##
## Linear Hypotheses:
```

```
##               Estimate Std. Error z value Pr(>|z|)
## CL - CF == 0   -72477     264382  -0.274    0.784
## (Adjusted p values reported -- single-step method)
##
##
## Tukey results for gene: C1QTNF7
##
##   Simultaneous Tests for General Linear Hypotheses
##
## Multiple Comparisons of Means: Tukey Contrasts
##
##
## Fit: glm(formula = Protein_Count ~ Tissue, family = gaussian, data = gene_
data)
##
## Linear Hypotheses:
##               Estimate Std. Error z value Pr(>|z|)
## CL - CF == 0    -5823     114125  -0.051    0.959
## (Adjusted p values reported -- single-step method)
##
##
## Tukey results for gene: CLEC11A
##
##   Simultaneous Tests for General Linear Hypotheses
##
## Multiple Comparisons of Means: Tukey Contrasts
##
##
## Fit: glm(formula = Protein_Count ~ Tissue, family = gaussian, data = gene_
data)
##
## Linear Hypotheses:
##               Estimate Std. Error z value Pr(>|z|)
## CL - CF == 0    65450     300260   0.218    0.827
## (Adjusted p values reported -- single-step method)
##
##
## Tukey results for gene: CLEC3B
##
##   Simultaneous Tests for General Linear Hypotheses
##
## Multiple Comparisons of Means: Tukey Contrasts
##
##
## Fit: glm(formula = Protein_Count ~ Tissue, family = gaussian, data = gene_
data)
##
## Linear Hypotheses:
##               Estimate Std. Error z value Pr(>|z|)
## CL - CF == 0  -914130    1558036  -0.587    0.557
```

```
## (Adjusted p values reported -- single-step method)
##
##
## Tukey results for gene: COLEC12
##
##   Simultaneous Tests for General Linear Hypotheses
##
## Multiple Comparisons of Means: Tukey Contrasts
##
##
## Fit: glm(formula = Protein_Count ~ Tissue, family = gaussian, data = gene_
data)
##
## Linear Hypotheses:
##              Estimate Std. Error z value Pr(>|z|)
## CL - CF == 0   -67298     239847  -0.281    0.779
## (Adjusted p values reported -- single-step method)
##
##
## Tukey results for gene: CSPG4
##
##   Simultaneous Tests for General Linear Hypotheses
##
## Multiple Comparisons of Means: Tukey Contrasts
##
##
## Fit: glm(formula = Protein_Count ~ Tissue, family = gaussian, data = gene_
data)
##
## Linear Hypotheses:
##              Estimate Std. Error z value Pr(>|z|)
## CL - CF == 0   113746     158046    0.72    0.472
## (Adjusted p values reported -- single-step method)
##
##
## Tukey results for gene: HPX
##
##   Simultaneous Tests for General Linear Hypotheses
##
## Multiple Comparisons of Means: Tukey Contrasts
##
##
## Fit: glm(formula = Protein_Count ~ Tissue, family = gaussian, data = gene_
data)
##
## Linear Hypotheses:
##              Estimate Std. Error z value Pr(>|z|)
## CL - CF == 0 -1038150     724755  -1.432    0.152
## (Adjusted p values reported -- single-step method)
##
```

```
##
## Tukey results for gene: LGALS1
##
##   Simultaneous Tests for General Linear Hypotheses
##
## Multiple Comparisons of Means: Tukey Contrasts
##
##
## Fit: glm(formula = Protein_Count ~ Tissue, family = gaussian, data = gene_
data)
##
## Linear Hypotheses:
##              Estimate Std. Error z value Pr(>|z|)
## CL - CF == 0  -966111     944295  -1.023    0.306
## (Adjusted p values reported -- single-step method)
##
##
## Tukey results for gene: LGALS9
##
##   Simultaneous Tests for General Linear Hypotheses
##
## Multiple Comparisons of Means: Tukey Contrasts
##
##
## Fit: glm(formula = Protein_Count ~ Tissue, family = gaussian, data = gene_
data)
##
## Linear Hypotheses:
##              Estimate Std. Error z value Pr(>|z|)
## CL - CF == 0    -5072     270826  -0.019    0.985
## (Adjusted p values reported -- single-step method)
##
##
## Tukey results for gene: LMAN1
##
##   Simultaneous Tests for General Linear Hypotheses
##
## Multiple Comparisons of Means: Tukey Contrasts
##
##
## Fit: glm(formula = Protein_Count ~ Tissue, family = gaussian, data = gene_
data)
##
## Linear Hypotheses:
##              Estimate Std. Error z value Pr(>|z|)
## CL - CF == 0    32897      58306   0.564    0.573
## (Adjusted p values reported -- single-step method)
##
##
## Tukey results for gene: OVGP1
```

```
##
##   Simultaneous Tests for General Linear Hypotheses
##
## Multiple Comparisons of Means: Tukey Contrasts
##
##
## Fit: glm(formula = Protein_Count ~ Tissue, family = gaussian, data = gene_
data)
##
## Linear Hypotheses:
##               Estimate Std. Error z value Pr(>|z|)
## CL - CF == 0  1779224    2184882   0.814    0.415
## (Adjusted p values reported -- single-step method)
##
##
## Tukey results for gene: PLXNB1
##
##   Simultaneous Tests for General Linear Hypotheses
##
## Multiple Comparisons of Means: Tukey Contrasts
##
##
## Fit: glm(formula = Protein_Count ~ Tissue, family = gaussian, data = gene_
data)
##
## Linear Hypotheses:
##               Estimate Std. Error z value Pr(>|z|)
## CL - CF == 0  -140010     103496  -1.353    0.176
## (Adjusted p values reported -- single-step method)
##
##
## Tukey results for gene: PLXNB2
##
##   Simultaneous Tests for General Linear Hypotheses
##
## Multiple Comparisons of Means: Tukey Contrasts
##
##
## Fit: glm(formula = Protein_Count ~ Tissue, family = gaussian, data = gene_
data)
##
## Linear Hypotheses:
##               Estimate Std. Error z value Pr(>|z|)
## CL - CF == 0  -251402     221060  -1.137    0.255
## (Adjusted p values reported -- single-step method)
```

```
#### Secreted Factors

# Read your cleaned joined data file into a data frame
cleaned_joined_data <- read.csv("cleaned_matrisome_cortex.csv")
```

```
# Filter rows with Matrisome_Category "Secreted Factors"
Secreted_Factors_data <- cleaned_joined_data %>%
  filter(Matrisome_Category == "Secreted Factors")

# Reshape the data to have separate columns for each replicate and each tissu
e stage
organized_data <- Secreted_Factors_data %>%
  pivot_longer(cols = starts_with("CF") | starts_with("CL"),
               names_to = c("Tissue", "Stage"),
               names_sep = "\\.",
               values_to = "Protein_Count")

## Warning: Expected 2 pieces. Missing pieces filled with `NA` in 2 rows [1,
7].

# Save the reshaped data to a new file

write.csv(organized_data, "Secreted_factors_data_matrisome_cortex_to_clean.cs
v", row.names = FALSE)

data2 <- read.csv("Secreted_factors_data_matrisome_cortex_clean.csv")

png(filename="cortex_Secreted_factors.png",
    width = 12, height = 6, units = "in",
    res = 600)

boxplot(Protein_Count ~ Tissue*Genes,
        data = data2,
        cex.axis = 0.8,
        col = viridis(2, alpha = 0.7),
        lwd = 0.7,
        ylab = "",
        ylim = c(0, 8000000),
        axes = FALSE,
        frame.plot = TRUE,
        xlab = "")

GROUP <- unique(data2$Genes)
groups <- 17
numbox <- 2
total <- groups * numbox
xpoints <- seq(median(1:numbox), total, numbox)

axis(2, seq(0, 8000000, by = 1000000), las = 1, cex.axis = 0.7)
axis(1, labels = GROUP, at = xpoints, las = 2, cex.axis = 0.7)

stripchart(data2$Protein_Count ~ data2$Tissue*data2$Genes,
           method = "jitter",
```

```
          pch = 19,
          col = "black",
          vertical = TRUE,
          add = TRUE,
          cex = 0.3)

title(ylab = expression("Protein abundance (RU)"), line = 3.0)
legend("top", fill = viridis(2), legend = c("Cortex Follicular", "Cortex Lute
al"), horiz = T, cex = 0.7)

dev.off()

## quartz_off_screen
##                 2

# Convert Tissue to factor
data2$Tissue <- as.factor(data2$Tissue)

# Create an empty list to store Tukey results
tukey_results_list <- list()

# Unique genes
unique_genes <- unique(data2$Genes)

# Loop over each gene
for (gene in unique_genes) {
  # Subset data for the current gene
  gene_data <- subset(data2, Genes == gene)

  # Fit the generalized linear model
  glm_model <- glm(Protein_Count ~ Tissue, data = gene_data, family = gaussia
n)

  # Perform Tukey test
  contrast_matrix <- glht(glm_model, linfct = mcp(Tissue = "Tukey"))
  tukey_results <- summary(contrast_matrix)

  # Store the results
  tukey_results_list[[gene]] <- tukey_results
}

# Print the results for each gene
for (gene in unique_genes) {
  cat("Tukey results for gene:", gene, "\n")
  print(tukey_results_list[[gene]])
  cat("\n")
}

## Tukey results for gene: ANGPT4
##
```

```
##   Simultaneous Tests for General Linear Hypotheses
##
## Multiple Comparisons of Means: Tukey Contrasts
##
##
## Fit: glm(formula = Protein_Count ~ Tissue, family = gaussian, data = gene_
data)
##
## Linear Hypotheses:
##              Estimate Std. Error z value Pr(>|z|)
## CL - CF == 0  -145301     222299  -0.654    0.513
## (Adjusted p values reported -- single-step method)
##
##
## Tukey results for gene: ANGPTL2
##
##   Simultaneous Tests for General Linear Hypotheses
##
## Multiple Comparisons of Means: Tukey Contrasts
##
##
## Fit: glm(formula = Protein_Count ~ Tissue, family = gaussian, data = gene_
data)
##
## Linear Hypotheses:
##              Estimate Std. Error z value Pr(>|z|)
## CL - CF == 0  -405618     224010  -1.811   0.0702 .
## ---
## Signif. codes:  0 '***' 0.001 '**' 0.01 '*' 0.05 '.' 0.1 ' ' 1
## (Adjusted p values reported -- single-step method)
##
##
## Tukey results for gene: ANGPTL6
##
##   Simultaneous Tests for General Linear Hypotheses
##
## Multiple Comparisons of Means: Tukey Contrasts
##
##
## Fit: glm(formula = Protein_Count ~ Tissue, family = gaussian, data = gene_
data)
##
## Linear Hypotheses:
##              Estimate Std. Error z value Pr(>|z|)
## CL - CF == 0  -119985     227831  -0.527    0.598
## (Adjusted p values reported -- single-step method)
##
##
## Tukey results for gene: CHRD
##
```

```
##   Simultaneous Tests for General Linear Hypotheses
##
## Multiple Comparisons of Means: Tukey Contrasts
##
##
## Fit: glm(formula = Protein_Count ~ Tissue, family = gaussian, data = gene_
data)
##
## Linear Hypotheses:
##              Estimate Std. Error z value Pr(>|z|)
## CL - CF == 0  -159640     110446  -1.445    0.148
## (Adjusted p values reported -- single-step method)
##
##
## Tukey results for gene: HCFC1
##
##   Simultaneous Tests for General Linear Hypotheses
##
## Multiple Comparisons of Means: Tukey Contrasts
##
##
## Fit: glm(formula = Protein_Count ~ Tissue, family = gaussian, data = gene_
data)
##
## Linear Hypotheses:
##              Estimate Std. Error z value Pr(>|z|)
## CL - CF == 0   -36142     189156  -0.191    0.848
## (Adjusted p values reported -- single-step method)
##
##
## Tukey results for gene: IGF2
##
##   Simultaneous Tests for General Linear Hypotheses
##
## Multiple Comparisons of Means: Tukey Contrasts
##
##
## Fit: glm(formula = Protein_Count ~ Tissue, family = gaussian, data = gene_
data)
##
## Linear Hypotheses:
##              Estimate Std. Error z value Pr(>|z|)
## CL - CF == 0  -174894     119034  -1.469    0.142
## (Adjusted p values reported -- single-step method)
##
##
## Tukey results for gene: IL17D
##
##   Simultaneous Tests for General Linear Hypotheses
##
```

```
## Multiple Comparisons of Means: Tukey Contrasts
##
##
## Fit: glm(formula = Protein_Count ~ Tissue, family = gaussian, data = gene_
data)
##
## Linear Hypotheses:
##              Estimate Std. Error z value Pr(>|z|)
## CL - CF == 0  -157421     193529  -0.813    0.416
## (Adjusted p values reported -- single-step method)
##
##
## Tukey results for gene: INSL3
##
##   Simultaneous Tests for General Linear Hypotheses
##
## Multiple Comparisons of Means: Tukey Contrasts
##
##
## Fit: glm(formula = Protein_Count ~ Tissue, family = gaussian, data = gene_
data)
##
## Linear Hypotheses:
##              Estimate Std. Error z value Pr(>|z|)
## CL - CF == 0   -18476      97520  -0.189     0.85
## (Adjusted p values reported -- single-step method)
##
##
## Tukey results for gene: PDGFC
##
##   Simultaneous Tests for General Linear Hypotheses
##
## Multiple Comparisons of Means: Tukey Contrasts
##
##
## Fit: glm(formula = Protein_Count ~ Tissue, family = gaussian, data = gene_
data)
##
## Linear Hypotheses:
##              Estimate Std. Error z value Pr(>|z|)
## CL - CF == 0   -83523      99253  -0.842      0.4
## (Adjusted p values reported -- single-step method)
##
##
## Tukey results for gene: S100A10
##
##   Simultaneous Tests for General Linear Hypotheses
##
## Multiple Comparisons of Means: Tukey Contrasts
##
```

```
##
## Fit: glm(formula = Protein_Count ~ Tissue, family = gaussian, data = gene_
data)
##
## Linear Hypotheses:
##              Estimate Std. Error z value Pr(>|z|)
## CL - CF == 0   -50570     226960  -0.223    0.824
## (Adjusted p values reported -- single-step method)
##
##
## Tukey results for gene: S100A11
##
##   Simultaneous Tests for General Linear Hypotheses
##
## Multiple Comparisons of Means: Tukey Contrasts
##
##
## Fit: glm(formula = Protein_Count ~ Tissue, family = gaussian, data = gene_
data)
##
## Linear Hypotheses:
##              Estimate Std. Error z value Pr(>|z|)
## CL - CF == 0   659463     914082   0.721    0.471
## (Adjusted p values reported -- single-step method)
##
##
## Tukey results for gene: S100A4
##
##   Simultaneous Tests for General Linear Hypotheses
##
## Multiple Comparisons of Means: Tukey Contrasts
##
##
## Fit: glm(formula = Protein_Count ~ Tissue, family = gaussian, data = gene_
data)
##
## Linear Hypotheses:
##              Estimate Std. Error z value Pr(>|z|)
## CL - CF == 0   266979     674195   0.396    0.692
## (Adjusted p values reported -- single-step method)
##
##
## Tukey results for gene: TGFB1
##
##   Simultaneous Tests for General Linear Hypotheses
##
## Multiple Comparisons of Means: Tukey Contrasts
##
##
## Fit: glm(formula = Protein_Count ~ Tissue, family = gaussian, data = gene_
```

```
data)
##
## Linear Hypotheses:
##               Estimate Std. Error z value Pr(>|z|)
## CL - CF == 0   -46135     164118  -0.281    0.779
## (Adjusted p values reported -- single-step method)
##
##
## Tukey results for gene: TGFB2
##
##   Simultaneous Tests for General Linear Hypotheses
##
## Multiple Comparisons of Means: Tukey Contrasts
##
##
## Fit: glm(formula = Protein_Count ~ Tissue, family = gaussian, data = gene_
data)
##
## Linear Hypotheses:
##               Estimate Std. Error z value Pr(>|z|)
## CL - CF == 0   224577    1242728   0.181    0.857
## (Adjusted p values reported -- single-step method)
##
##
## Tukey results for gene: TGFB3
##
##   Simultaneous Tests for General Linear Hypotheses
##
## Multiple Comparisons of Means: Tukey Contrasts
##
##
## Fit: glm(formula = Protein_Count ~ Tissue, family = gaussian, data = gene_
data)
##
## Linear Hypotheses:
##               Estimate Std. Error z value Pr(>|z|)
## CL - CF == 0   118207     932750   0.127    0.899
## (Adjusted p values reported -- single-step method)
##
##
## Tukey results for gene: WNT4
##
##   Simultaneous Tests for General Linear Hypotheses
##
## Multiple Comparisons of Means: Tukey Contrasts
##
##
## Fit: glm(formula = Protein_Count ~ Tissue, family = gaussian, data = gene_
data)
##
```

```
## Linear Hypotheses:
##              Estimate Std. Error z value Pr(>|z|)
## CL - CF == 0   -56647      88241  -0.642    0.521
## (Adjusted p values reported -- single-step method)
##
##
## Tukey results for gene: WNT6
##
##   Simultaneous Tests for General Linear Hypotheses
##
## Multiple Comparisons of Means: Tukey Contrasts
##
##
## Fit: glm(formula = Protein_Count ~ Tissue, family = gaussian, data = gene_
data)
##
## Linear Hypotheses:
##              Estimate Std. Error z value Pr(>|z|)
## CL - CF == 0   -76491     166685  -0.459    0.646
## (Adjusted p values reported -- single-step method)
```

```
#### Components proportions

data <- read.csv("cortex_matrisome_final.csv")

# Subset the data for each tissue (CF and CL) separately
data_CF <- subset(data, Tissue == "CF")
data_CL <- subset(data, Tissue == "CL")

# Function to calculate proportions
calculate_proportions <- function(data_subset) {
  proportions <- data_subset %>%
    group_by(Matrisome_Category) %>%
    summarize(Proportion = sum(Protein_Count) / sum(data_subset$Protein_Count
) * 100)
  return(proportions)
}

# Calculate proportions for CF
proportions_CF <- calculate_proportions(data_CF)

# Calculate proportions for CL
proportions_CL <- calculate_proportions(data_CL)

# Print proportions for CF
cat("Proportions for Matrisome categories in tissue CF:\n")
```

```
## Proportions for Matrisome categories in tissue CF:
```

```
print(proportions_CF)
```

```
## # A tibble: 6 × 2
##   Matrisome_Category Proportion
##   <chr>                   <dbl>
## 1 Collagens              76.8
## 2 ECM Glycoproteins      13.1
## 3 ECM Regulators          3.36
## 4 ECM-affiliated          0.744
## 5 Proteoglycans           5.77
## 6 Secreted Factors        0.248

# Print proportions for CL
cat("Proportions for Matrisome categories in tissue CL:\n")

## Proportions for Matrisome categories in tissue CL:

print(proportions_CL)

## # A tibble: 6 × 2
##   Matrisome_Category Proportion
##   <chr>                   <dbl>
## 1 Collagens              78.9
## 2 ECM Glycoproteins      12.0
## 3 ECM Regulators          3.02
## 4 ECM-affiliated          0.613
## 5 Proteoglycans           5.21
## 6 Secreted Factors        0.232

###Pie charts

# Define a function to create a pie chart with values and save it
create_and_save_pie_chart <- function(proportions, title, filename) {
  # Create a color palette using Viridis
  color_palette <- viridis(length(unique(proportions$Matrisome_Category)))

  # Create a pie chart
  pie_chart <- ggplot(proportions, aes(x = "", y = Proportion, fill = Matriso
me_Category)) +
    geom_bar(stat = "identity") +
    coord_polar("y", start = 1) +
    geom_text(aes(label = sprintf("%.1f%%", Proportion)), position = position
_stack(vjust = 0.5), color = "white", size = 6) +
    labs(title = title, x = NULL, y = NULL, fill = "Matrisome Category") +
    scale_fill_manual(values = color_palette) +  # Set the color palette
    theme_void() +
    theme(legend.position = "right")

  # Save the pie chart
  ggsave(filename, plot = pie_chart, width = 6, height = 6, units = "in", dpi
= 300)
}
```

```r
# Create and save a pie chart for CF
create_and_save_pie_chart(proportions_CF, "Matrisome Categories Proportion in
Tissue CF", "pie_chart_CF.png")

# Create and save a pie chart for CL
create_and_save_pie_chart(proportions_CL, "Matrisome Categories Proportion in
Tissue CL", "pie_chart_CL.png")


#####Summary data

data <- read.csv("cortex_matrisome_final.csv")

# Calculate mean and standard deviation for each Gene considering Tissue and
Matrisome_Category
gene_stats <- data %>%
  group_by(Genes, Tissue, Matrisome_Category) %>%
  summarize(
    Mean_Protein_Count = mean(Protein_Count),
    SD_Protein_Count = sd(Protein_Count)
  )

## `summarise()` has grouped output by 'Genes', 'Tissue'. You can override us
ing
## the `.groups` argument.

# Print the results
print(gene_stats)

## # A tibble: 424 × 5
## # Groups:   Genes, Tissue [424]
##    Genes    Tissue Matrisome_Category Mean_Protein_Count SD_Protein_Count
##    <chr>    <chr>  <chr>                           <dbl>            <dbl>
##  1 A2M      CF     ECM Regulators               2333823.         1062902.
##  2 A2M      CL     ECM Regulators               2295198           539720.
##  3 ABI3BP   CF     ECM Glycoproteins             899573.          877969.
##  4 ABI3BP   CL     ECM Glycoproteins             584759.          451847.
##  5 ACAN     CF     Proteoglycans                 911989.          771834.
##  6 ACAN     CL     Proteoglycans                 555198.          332330.
##  7 ADAMTSL4 CF     ECM Regulators               1063472.          245931.
##  8 ADAMTSL4 CL     ECM Regulators                974284.          311757.
##  9 ADAMTSL5 CF     ECM Regulators                236358.          246096.
## 10 ADAMTSL5 CL     ECM Regulators                 99607.          139118.
## # ℹ 414 more rows

# Save the summarized data to a CSV file
write.csv(gene_stats, file = "Cortex_Summary.csv", row.names = FALSE)
```

```
###MEDULLA

# Read your data into a data frame or data table
data <- read.csv("Medulla_all_count.csv")

# Remove rows with more than 6 NA values
clean_data <- data %>%
  filter(rowSums(is.na(.)) <= 6)

# Replace remaining NA values with 0 in all columns
clean_data <- clean_data %>%
  mutate_all(funs(replace(., is.na(.), 0)))

## Warning: `funs()` was deprecated in dplyr 0.8.0.
## ℹ Please use a list of either functions or lambdas:
##
## # Simple named list: list(mean = mean, median = median)
##
## # Auto named with `tibble::lst()`: tibble::lst(mean, median)
##
## # Using lambdas list(~ mean(., trim = .2), ~ median(., na.rm = TRUE))
## Call `lifecycle::last_lifecycle_warnings()` to see where this warning was
## generated.

# Save the cleaned data to a new file
write.csv(clean_data, "cleaned_data_medulla.csv", row.names = FALSE)


### Matrisome categorization

# Read your first data file into a data frame or data table
data <- read.csv("cleaned_data_medulla.csv")

# Read your second dataset into another data frame or data table
matrisome_info <- read.csv("Bt_Matrisome.csv")

# Perform a left join based on the "Genes" column
joined_data <- left_join(data, matrisome_info, by = "Genes")

# Save the joined data to a new file
write.csv(joined_data, "medulla_matrisome.csv", row.names = FALSE)

# Remove rows where either Matrisome_Division or Matrisome_Category have NA v
alues
cleaned_joined_data <- joined_data %>%
  filter(!is.na(Matrisome_Division) & !is.na(Matrisome_Category))

# Save the cleaned data to a new file
```

```
write.csv(cleaned_joined_data, "cleaned_matrisome_medu.csv", row.names = FALS
E)

#######Ploting


#### Collagens
# Read your cleaned joined data file into a data frame
cleaned_joined_data <- read.csv("cleaned_matrisome_medu.csv")

# Filter rows with Matrisome_Category "Collagens"
collagen_data <- cleaned_joined_data %>%
  filter(Matrisome_Category == "Collagens")

# Reshape the data to have separate columns for each replicate and each tissu
e stage
organized_data <- collagen_data %>%
  pivot_longer(cols = starts_with("MF") | starts_with("ML"),
               names_to = c("Tissue", "Stage"),
               names_sep = "\\.",
               values_to = "Protein_Count")

## Warning: Expected 2 pieces. Missing pieces filled with `NA` in 2 rows [1,
6].

# Save the reshaped data to a new file

write.csv(organized_data, "Collagens_matrisome_medu_to_clean.csv", row.names
= FALSE)

data2 <- read.csv("Collagens_matrisome_medu_clean.csv")

png(filename="medu_collagens.png",
    width = 12, height = 6, units = "in",
    res = 600)

boxplot(Protein_Count ~ Tissue*Genes,
        data = data2,
        cex.axis = 0.8,
        col = viridis(2, alpha = 0.7),
        lwd = 0.7,
        ylab = "",
        ylim = c(0, 4500000000),
        axes = FALSE,
        frame.plot = TRUE,
        xlab = "")

GROUP <- unique(data2$Genes)
groups <- 25
```

```
numbox <- 2
total <- groups * numbox
xpoints <- seq(median(1:numbox), total, numbox)

axis(2, seq(0, 4500000000, by = 1000000000), las = 1, cex.axis = 0.7)
axis(1, labels = GROUP, at = xpoints, las = 2, cex.axis = 0.7)

stripchart(data2$Protein_Count ~ data2$Tissue*data2$Genes,
           method = "jitter",
           pch = 19,
           col = "black",
           vertical = TRUE,
           add = TRUE,
           cex = 0.3)

title(ylab = expression("Protein abundance (RU)"), line = 3.0)
legend("top", fill = viridis(2), legend = c("Medulla Follicular", "Medulla Lu
teal"), horiz = T, cex = 0.7)

dev.off()

## quartz_off_screen
##                 2

# Convert Tissue to factor
data2$Tissue <- as.factor(data2$Tissue)

# Create an empty list to store Tukey results
tukey_results_list <- list()

# Unique genes
unique_genes <- unique(data2$Genes)

# Loop over each gene
for (gene in unique_genes) {
  # Subset data for the current gene
  gene_data <- subset(data2, Genes == gene)

  # Fit the generalized linear model
  glm_model <- glm(Protein_Count ~ Tissue, data = gene_data, family = gaussia
n)

  # Perform Tukey test
  contrast_matrix <- glht(glm_model, linfct = mcp(Tissue = "Tukey"))
  tukey_results <- summary(contrast_matrix)

  # Store the results
  tukey_results_list[[gene]] <- tukey_results
}
```

```r
# Print the results for each gene
for (gene in unique_genes) {
  cat("Tukey results for gene:", gene, "\n")
  print(tukey_results_list[[gene]])
  cat("\n")
}
```

```
## Tukey results for gene: COL11A1
##
##   Simultaneous Tests for General Linear Hypotheses
##
## Multiple Comparisons of Means: Tukey Contrasts
##
##
## Fit: glm(formula = Protein_Count ~ Tissue, family = gaussian, data = gene_
data)
##
## Linear Hypotheses:
##                Estimate Std. Error z value Pr(>|z|)
## ML - MF == 0  -217200     503303  -0.432    0.666
## (Adjusted p values reported -- single-step method)
##
##
## Tukey results for gene: COL12A1
##
##   Simultaneous Tests for General Linear Hypotheses
##
## Multiple Comparisons of Means: Tukey Contrasts
##
##
## Fit: glm(formula = Protein_Count ~ Tissue, family = gaussian, data = gene_
data)
##
## Linear Hypotheses:
##                Estimate Std. Error z value Pr(>|z|)
## ML - MF == 0  7952440    7622549   1.043    0.297
## (Adjusted p values reported -- single-step method)
##
##
## Tukey results for gene: COL14A1
##
##   Simultaneous Tests for General Linear Hypotheses
##
## Multiple Comparisons of Means: Tukey Contrasts
##
##
## Fit: glm(formula = Protein_Count ~ Tissue, family = gaussian, data = gene_
data)
##
```

```
## Linear Hypotheses:
##              Estimate Std. Error z value Pr(>|z|)
## ML - MF == 0  7082980   10311803   0.687    0.492
## (Adjusted p values reported -- single-step method)
##
##
## Tukey results for gene: COL15A1
##
##   Simultaneous Tests for General Linear Hypotheses
##
## Multiple Comparisons of Means: Tukey Contrasts
##
##
## Fit: glm(formula = Protein_Count ~ Tissue, family = gaussian, data = gene_
data)
##
## Linear Hypotheses:
##              Estimate Std. Error z value Pr(>|z|)
## ML - MF == 0   963182    1267887    0.76    0.447
## (Adjusted p values reported -- single-step method)
##
##
## Tukey results for gene: COL16A1
##
##   Simultaneous Tests for General Linear Hypotheses
##
## Multiple Comparisons of Means: Tukey Contrasts
##
##
## Fit: glm(formula = Protein_Count ~ Tissue, family = gaussian, data = gene_
data)
##
## Linear Hypotheses:
##              Estimate Std. Error z value Pr(>|z|)
## ML - MF == 0 -2733852    3495806  -0.782    0.434
## (Adjusted p values reported -- single-step method)
##
##
## Tukey results for gene: COL18A1
##
##   Simultaneous Tests for General Linear Hypotheses
##
## Multiple Comparisons of Means: Tukey Contrasts
##
##
## Fit: glm(formula = Protein_Count ~ Tissue, family = gaussian, data = gene_
data)
##
## Linear Hypotheses:
##              Estimate Std. Error z value Pr(>|z|)
```

```
## ML - MF == 0  3562794    2422047   1.471    0.141
## (Adjusted p values reported -- single-step method)
##
##
## Tukey results for gene: COL1A1
##
##   Simultaneous Tests for General Linear Hypotheses
##
## Multiple Comparisons of Means: Tukey Contrasts
##
##
## Fit: glm(formula = Protein_Count ~ Tissue, family = gaussian, data = gene_
data)
##
## Linear Hypotheses:
##                Estimate Std. Error z value Pr(>|z|)
## ML - MF == 0 -372162400  977583328  -0.381    0.703
## (Adjusted p values reported -- single-step method)
##
##
## Tukey results for gene: COL1A2
##
##   Simultaneous Tests for General Linear Hypotheses
##
## Multiple Comparisons of Means: Tukey Contrasts
##
##
## Fit: glm(formula = Protein_Count ~ Tissue, family = gaussian, data = gene_
data)
##
## Linear Hypotheses:
##                Estimate Std. Error z value Pr(>|z|)
## ML - MF == 0 -268341200  443450186  -0.605    0.545
## (Adjusted p values reported -- single-step method)
##
##
## Tukey results for gene: COL27A1
##
##   Simultaneous Tests for General Linear Hypotheses
##
## Multiple Comparisons of Means: Tukey Contrasts
##
##
## Fit: glm(formula = Protein_Count ~ Tissue, family = gaussian, data = gene_
data)
##
## Linear Hypotheses:
##              Estimate Std. Error z value Pr(>|z|)
## ML - MF == 0   -69591      87508  -0.795    0.426
## (Adjusted p values reported -- single-step method)
```

```
##
##
## Tukey results for gene: COL2A1
##
##   Simultaneous Tests for General Linear Hypotheses
##
## Multiple Comparisons of Means: Tukey Contrasts
##
##
## Fit: glm(formula = Protein_Count ~ Tissue, family = gaussian, data = gene_
data)
##
## Linear Hypotheses:
##              Estimate Std. Error z value Pr(>|z|)
## ML - MF == 0    52120     366156   0.142    0.887
## (Adjusted p values reported -- single-step method)
##
##
## Tukey results for gene: COL3A1
##
##   Simultaneous Tests for General Linear Hypotheses
##
## Multiple Comparisons of Means: Tukey Contrasts
##
##
## Fit: glm(formula = Protein_Count ~ Tissue, family = gaussian, data = gene_
data)
##
## Linear Hypotheses:
##              Estimate Std. Error z value Pr(>|z|)
## ML - MF == 0  -771322    1897018  -0.407    0.684
## (Adjusted p values reported -- single-step method)
##
##
## Tukey results for gene: COL4A1
##
##   Simultaneous Tests for General Linear Hypotheses
##
## Multiple Comparisons of Means: Tukey Contrasts
##
##
## Fit: glm(formula = Protein_Count ~ Tissue, family = gaussian, data = gene_
data)
##
## Linear Hypotheses:
##              Estimate Std. Error z value Pr(>|z|)
## ML - MF == 0   633452    3802083   0.167    0.868
## (Adjusted p values reported -- single-step method)
##
##
```

```
## Tukey results for gene: COL4A2
##
##   Simultaneous Tests for General Linear Hypotheses
##
## Multiple Comparisons of Means: Tukey Contrasts
##
##
## Fit: glm(formula = Protein_Count ~ Tissue, family = gaussian, data = gene_
data)
##
## Linear Hypotheses:
##              Estimate Std. Error z value Pr(>|z|)
## ML - MF == 0     5576    1665375   0.003    0.997
## (Adjusted p values reported -- single-step method)
##
##
## Tukey results for gene: COL4A3
##
##   Simultaneous Tests for General Linear Hypotheses
##
## Multiple Comparisons of Means: Tukey Contrasts
##
##
## Fit: glm(formula = Protein_Count ~ Tissue, family = gaussian, data = gene_
data)
##
## Linear Hypotheses:
##              Estimate Std. Error z value Pr(>|z|)
## ML - MF == 0  -994750    3004797  -0.331    0.741
## (Adjusted p values reported -- single-step method)
##
##
## Tukey results for gene: COL4A4
##
##   Simultaneous Tests for General Linear Hypotheses
##
## Multiple Comparisons of Means: Tukey Contrasts
##
##
## Fit: glm(formula = Protein_Count ~ Tissue, family = gaussian, data = gene_
data)
##
## Linear Hypotheses:
##              Estimate Std. Error z value Pr(>|z|)
## ML - MF == 0    -4269     518439  -0.008    0.993
## (Adjusted p values reported -- single-step method)
##
##
## Tukey results for gene: COL4A5
##
```

```
##   Simultaneous Tests for General Linear Hypotheses
##
## Multiple Comparisons of Means: Tukey Contrasts
##
##
## Fit: glm(formula = Protein_Count ~ Tissue, family = gaussian, data = gene_
data)
##
## Linear Hypotheses:
##              Estimate Std. Error z value Pr(>|z|)
## ML - MF == 0 -1599210    4337963  -0.369    0.712
## (Adjusted p values reported -- single-step method)
##
##
## Tukey results for gene: COL4A6
##
##   Simultaneous Tests for General Linear Hypotheses
##
## Multiple Comparisons of Means: Tukey Contrasts
##
##
## Fit: glm(formula = Protein_Count ~ Tissue, family = gaussian, data = gene_
data)
##
## Linear Hypotheses:
##              Estimate Std. Error z value Pr(>|z|)
## ML - MF == 0 -1336624    1398150  -0.956    0.339
## (Adjusted p values reported -- single-step method)
##
##
## Tukey results for gene: COL5A1
##
##   Simultaneous Tests for General Linear Hypotheses
##
## Multiple Comparisons of Means: Tukey Contrasts
##
##
## Fit: glm(formula = Protein_Count ~ Tissue, family = gaussian, data = gene_
data)
##
## Linear Hypotheses:
##              Estimate Std. Error z value Pr(>|z|)
## ML - MF == 0  -617162    1292975  -0.477    0.633
## (Adjusted p values reported -- single-step method)
##
##
## Tukey results for gene: COL5A2
##
##   Simultaneous Tests for General Linear Hypotheses
##
```

```
## Multiple Comparisons of Means: Tukey Contrasts
##
##
## Fit: glm(formula = Protein_Count ~ Tissue, family = gaussian, data = gene_
data)
##
## Linear Hypotheses:
##               Estimate Std. Error z value Pr(>|z|)
## ML - MF == 0 -3775022    7561710  -0.499    0.618
## (Adjusted p values reported -- single-step method)
##
##
## Tukey results for gene: COL5A3
##
##   Simultaneous Tests for General Linear Hypotheses
##
## Multiple Comparisons of Means: Tukey Contrasts
##
##
## Fit: glm(formula = Protein_Count ~ Tissue, family = gaussian, data = gene_
data)
##
## Linear Hypotheses:
##               Estimate Std. Error z value Pr(>|z|)
## ML - MF == 0   247336    3373047   0.073    0.942
## (Adjusted p values reported -- single-step method)
##
##
## Tukey results for gene: COL6A1
##
##   Simultaneous Tests for General Linear Hypotheses
##
## Multiple Comparisons of Means: Tukey Contrasts
##
##
## Fit: glm(formula = Protein_Count ~ Tissue, family = gaussian, data = gene_
data)
##
## Linear Hypotheses:
##                Estimate Std. Error z value Pr(>|z|)
## ML - MF == 0 -16282480   51086523  -0.319     0.75
## (Adjusted p values reported -- single-step method)
##
##
## Tukey results for gene: COL6A2
##
##   Simultaneous Tests for General Linear Hypotheses
##
## Multiple Comparisons of Means: Tukey Contrasts
##
```

```
##
## Fit: glm(formula = Protein_Count ~ Tissue, family = gaussian, data = gene_
data)
##
## Linear Hypotheses:
##                 Estimate Std. Error z value Pr(>|z|)
## ML - MF == 0 -9209760   25805207  -0.357    0.721
## (Adjusted p values reported -- single-step method)
##
##
## Tukey results for gene: COL6A3
##
##   Simultaneous Tests for General Linear Hypotheses
##
## Multiple Comparisons of Means: Tukey Contrasts
##
##
## Fit: glm(formula = Protein_Count ~ Tissue, family = gaussian, data = gene_
data)
##
## Linear Hypotheses:
##                 Estimate Std. Error z value Pr(>|z|)
## ML - MF == 0 20782600   28327408   0.734    0.463
## (Adjusted p values reported -- single-step method)
##
##
## Tukey results for gene: COL6A5
##
##   Simultaneous Tests for General Linear Hypotheses
##
## Multiple Comparisons of Means: Tukey Contrasts
##
##
## Fit: glm(formula = Protein_Count ~ Tissue, family = gaussian, data = gene_
data)
##
## Linear Hypotheses:
##                 Estimate Std. Error z value Pr(>|z|)
## ML - MF == 0   183568     436588    0.42    0.674
## (Adjusted p values reported -- single-step method)
##
##
## Tukey results for gene: COL8A1
##
##   Simultaneous Tests for General Linear Hypotheses
##
## Multiple Comparisons of Means: Tukey Contrasts
##
##
## Fit: glm(formula = Protein_Count ~ Tissue, family = gaussian, data = gene_
```

```
data)
##
## Linear Hypotheses:
##               Estimate Std. Error z value Pr(>|z|)
## ML - MF == 0  -125293     256510  -0.488    0.625
## (Adjusted p values reported -- single-step method)
```

```
#### ECM Glycoproteins
# Read your cleaned joined data file into a data frame
cleaned_joined_data <- read.csv("cleaned_matrisome_medu.csv")

# Filter rows with Matrisome_Category "ECM Glycoproteins"
Glycoproteins_data <- cleaned_joined_data %>%
  filter(Matrisome_Category == "ECM Glycoproteins")

# Reshape the data to have separate columns for each replicate and each tissu
e stage
organized_data <- Glycoproteins_data %>%
  pivot_longer(cols = starts_with("MF") | starts_with("ML"),
               names_to = c("Tissue", "Stage"),
               names_sep = "\\.",
               values_to = "Protein_Count")
```

```
## Warning: Expected 2 pieces. Missing pieces filled with `NA` in 2 rows [1,
6].
```

```
# Save the reshaped data to a new file

write.csv(organized_data, "Glycoproteins_data_matrisome_medu_to_clean.csv", r
ow.names = FALSE)

data2 <- read.csv("Glycoproteins_data_matrisome_medu_clean.csv")

png(filename="medu_ECM Glycoproteins.png",
    width = 12, height = 6, units = "in",
    res = 600)

boxplot(Protein_Count ~ Tissue*Genes,
        data = data2,
        cex.axis = 0.8,
        col = viridis(2, alpha = 0.7),
        lwd = 0.7,
        ylab = "",
        ylim = c(0, 800000000),
        axes = FALSE,
        frame.plot = TRUE,
        xlab = "")

GROUP <- unique(data2$Genes)
groups <- 90
```

```
numbox <- 2
total <- groups * numbox
xpoints <- seq(median(1:numbox), total, numbox)

axis(2, seq(0, 800000000, by = 100000000), las = 1, cex.axis = 0.7)
axis(1, labels = GROUP, at = xpoints, las = 2, cex.axis = 0.7)

stripchart(data2$Protein_Count ~ data2$Tissue*data2$Genes,
           method = "jitter",
           pch = 19,
           col = "black",
           vertical = TRUE,
           add = TRUE,
           cex = 0.3)

title(ylab = expression("Protein abundance (RU)"), line = 3.0)
legend("top", fill = viridis(2), legend = c("Medulla Follicular", "Medulla Lu
teal"), horiz = T, cex = 0.7)

dev.off()

## quartz_off_screen
##                 2

# Convert Tissue to factor
data2$Tissue <- as.factor(data2$Tissue)

# Create an empty list to store Tukey results
tukey_results_list <- list()

# Unique genes
unique_genes <- unique(data2$Genes)

# Loop over each gene
for (gene in unique_genes) {
  # Subset data for the current gene
  gene_data <- subset(data2, Genes == gene)

  # Fit the generalized linear model
  glm_model <- glm(Protein_Count ~ Tissue, data = gene_data, family = gaussia
n)

  # Perform Tukey test
  contrast_matrix <- glht(glm_model, linfct = mcp(Tissue = "Tukey"))
  tukey_results <- summary(contrast_matrix)

  # Store the results
  tukey_results_list[[gene]] <- tukey_results
}
```

```r
# Print the results for each gene
for (gene in unique_genes) {
  cat("Tukey results for gene:", gene, "\n")
  print(tukey_results_list[[gene]])
  cat("\n")
}
```

```
## Tukey results for gene: ABI3BP
##
##   Simultaneous Tests for General Linear Hypotheses
##
## Multiple Comparisons of Means: Tukey Contrasts
##
##
## Fit: glm(formula = Protein_Count ~ Tissue, family = gaussian, data = gene_
data)
##
## Linear Hypotheses:
##              Estimate Std. Error z value Pr(>|z|)
## ML - MF == 0   -11493     159458  -0.072    0.943
## (Adjusted p values reported -- single-step method)
##
##
## Tukey results for gene: AEBP1
##
##   Simultaneous Tests for General Linear Hypotheses
##
## Multiple Comparisons of Means: Tukey Contrasts
##
##
## Fit: glm(formula = Protein_Count ~ Tissue, family = gaussian, data = gene_
data)
##
## Linear Hypotheses:
##              Estimate Std. Error z value Pr(>|z|)
## ML - MF == 0  -194346     846930  -0.229    0.819
## (Adjusted p values reported -- single-step method)
##
##
## Tukey results for gene: AGRN
##
##   Simultaneous Tests for General Linear Hypotheses
##
## Multiple Comparisons of Means: Tukey Contrasts
##
##
## Fit: glm(formula = Protein_Count ~ Tissue, family = gaussian, data = gene_
data)
##
```

```
## Linear Hypotheses:
##              Estimate Std. Error z value Pr(>|z|)
## ML - MF == 0  -457298     290936  -1.572    0.116
## (Adjusted p values reported -- single-step method)
##
##
## Tukey results for gene: AHSG
##
##   Simultaneous Tests for General Linear Hypotheses
##
## Multiple Comparisons of Means: Tukey Contrasts
##
##
## Fit: glm(formula = Protein_Count ~ Tissue, family = gaussian, data = gene_
data)
##
## Linear Hypotheses:
##              Estimate Std. Error z value Pr(>|z|)
## ML - MF == 0  -450119     292453  -1.539    0.124
## (Adjusted p values reported -- single-step method)
##
##
## Tukey results for gene: CILP2
##
##   Simultaneous Tests for General Linear Hypotheses
##
## Multiple Comparisons of Means: Tukey Contrasts
##
##
## Fit: glm(formula = Protein_Count ~ Tissue, family = gaussian, data = gene_
data)
##
## Linear Hypotheses:
##              Estimate Std. Error z value Pr(>|z|)
## ML - MF == 0  8146220   21199608   0.384    0.701
## (Adjusted p values reported -- single-step method)
##
##
## Tukey results for gene: COCH
##
##   Simultaneous Tests for General Linear Hypotheses
##
## Multiple Comparisons of Means: Tukey Contrasts
##
##
## Fit: glm(formula = Protein_Count ~ Tissue, family = gaussian, data = gene_
data)
##
## Linear Hypotheses:
##              Estimate Std. Error z value Pr(>|z|)
```

```
## ML - MF == 0   110967     272043   0.408    0.683
## (Adjusted p values reported -- single-step method)
##
##
## Tukey results for gene: CRELD2
##
##   Simultaneous Tests for General Linear Hypotheses
##
## Multiple Comparisons of Means: Tukey Contrasts
##
##
## Fit: glm(formula = Protein_Count ~ Tissue, family = gaussian, data = gene_
data)
##
## Linear Hypotheses:
##              Estimate Std. Error z value Pr(>|z|)
## ML - MF == 0   -11429      27290  -0.419    0.675
## (Adjusted p values reported -- single-step method)
##
##
## Tukey results for gene: CTHRC1
##
##   Simultaneous Tests for General Linear Hypotheses
##
## Multiple Comparisons of Means: Tukey Contrasts
##
##
## Fit: glm(formula = Protein_Count ~ Tissue, family = gaussian, data = gene_
data)
##
## Linear Hypotheses:
##              Estimate Std. Error z value Pr(>|z|)
## ML - MF == 0    34387      38207     0.9    0.368
## (Adjusted p values reported -- single-step method)
##
##
## Tukey results for gene: DPT
##
##   Simultaneous Tests for General Linear Hypotheses
##
## Multiple Comparisons of Means: Tukey Contrasts
##
##
## Fit: glm(formula = Protein_Count ~ Tissue, family = gaussian, data = gene_
data)
##
## Linear Hypotheses:
##               Estimate Std. Error z value Pr(>|z|)
## ML - MF == 0 -67559800  123582120  -0.547    0.585
## (Adjusted p values reported -- single-step method)
```

```
##
##
## Tukey results for gene: ECM1
##
##   Simultaneous Tests for General Linear Hypotheses
##
## Multiple Comparisons of Means: Tukey Contrasts
##
##
## Fit: glm(formula = Protein_Count ~ Tissue, family = gaussian, data = gene_
data)
##
## Linear Hypotheses:
##              Estimate Std. Error z value Pr(>|z|)
## ML - MF == 0  -624430    1156381   -0.54    0.589
## (Adjusted p values reported -- single-step method)
##
##
## Tukey results for gene: ECM2
##
##   Simultaneous Tests for General Linear Hypotheses
##
## Multiple Comparisons of Means: Tukey Contrasts
##
##
## Fit: glm(formula = Protein_Count ~ Tissue, family = gaussian, data = gene_
data)
##
## Linear Hypotheses:
##              Estimate Std. Error z value Pr(>|z|)
## ML - MF == 0    69716     309707   0.225    0.822
## (Adjusted p values reported -- single-step method)
##
##
## Tukey results for gene: EFEMP1
##
##   Simultaneous Tests for General Linear Hypotheses
##
## Multiple Comparisons of Means: Tukey Contrasts
##
##
## Fit: glm(formula = Protein_Count ~ Tissue, family = gaussian, data = gene_
data)
##
## Linear Hypotheses:
##              Estimate Std. Error z value Pr(>|z|)
## ML - MF == 0  -236822    1385795  -0.171    0.864
## (Adjusted p values reported -- single-step method)
##
##
```

```
## Tukey results for gene: EFEMP2
##
##   Simultaneous Tests for General Linear Hypotheses
##
## Multiple Comparisons of Means: Tukey Contrasts
##
##
## Fit: glm(formula = Protein_Count ~ Tissue, family = gaussian, data = gene_
data)
##
## Linear Hypotheses:
##              Estimate Std. Error z value Pr(>|z|)
## ML - MF == 0    43836    1031016   0.043    0.966
## (Adjusted p values reported -- single-step method)
##
##
## Tukey results for gene: EGFLAM
##
##   Simultaneous Tests for General Linear Hypotheses
##
## Multiple Comparisons of Means: Tukey Contrasts
##
##
## Fit: glm(formula = Protein_Count ~ Tissue, family = gaussian, data = gene_
data)
##
## Linear Hypotheses:
##              Estimate Std. Error z value Pr(>|z|)
## ML - MF == 0    58785     108302   0.543    0.587
## (Adjusted p values reported -- single-step method)
##
##
## Tukey results for gene: EMID1
##
##   Simultaneous Tests for General Linear Hypotheses
##
## Multiple Comparisons of Means: Tukey Contrasts
##
##
## Fit: glm(formula = Protein_Count ~ Tissue, family = gaussian, data = gene_
data)
##
## Linear Hypotheses:
##              Estimate Std. Error z value Pr(>|z|)
## ML - MF == 0    58328     367931   0.159    0.874
## (Adjusted p values reported -- single-step method)
##
##
## Tukey results for gene: EMILIN1
##
```

```
##   Simultaneous Tests for General Linear Hypotheses
##
## Multiple Comparisons of Means: Tukey Contrasts
##
##
## Fit: glm(formula = Protein_Count ~ Tissue, family = gaussian, data = gene_
data)
##
## Linear Hypotheses:
##               Estimate Std. Error z value Pr(>|z|)
## ML - MF == 0 -4858684    6040504  -0.804    0.421
## (Adjusted p values reported -- single-step method)
##
##
## Tukey results for gene: EMILIN2
##
##   Simultaneous Tests for General Linear Hypotheses
##
## Multiple Comparisons of Means: Tukey Contrasts
##
##
## Fit: glm(formula = Protein_Count ~ Tissue, family = gaussian, data = gene_
data)
##
## Linear Hypotheses:
##               Estimate Std. Error z value Pr(>|z|)
## ML - MF == 0   -32662     148038  -0.221    0.825
## (Adjusted p values reported -- single-step method)
##
##
## Tukey results for gene: EMILIN3
##
##   Simultaneous Tests for General Linear Hypotheses
##
## Multiple Comparisons of Means: Tukey Contrasts
##
##
## Fit: glm(formula = Protein_Count ~ Tissue, family = gaussian, data = gene_
data)
##
## Linear Hypotheses:
##               Estimate Std. Error z value Pr(>|z|)
## ML - MF == 0   -39672     180742  -0.219    0.826
## (Adjusted p values reported -- single-step method)
##
##
## Tukey results for gene: FBLN1
##
##   Simultaneous Tests for General Linear Hypotheses
##
```

```
## Multiple Comparisons of Means: Tukey Contrasts
##
##
## Fit: glm(formula = Protein_Count ~ Tissue, family = gaussian, data = gene_
data)
##
## Linear Hypotheses:
##                Estimate Std. Error z value Pr(>|z|)
## ML - MF == 0   111640    1097659   0.102    0.919
## (Adjusted p values reported -- single-step method)
##
##
## Tukey results for gene: FBLN2
##
##   Simultaneous Tests for General Linear Hypotheses
##
## Multiple Comparisons of Means: Tukey Contrasts
##
##
## Fit: glm(formula = Protein_Count ~ Tissue, family = gaussian, data = gene_
data)
##
## Linear Hypotheses:
##                Estimate Std. Error z value Pr(>|z|)
## ML - MF == 0    40948      87024   0.471    0.638
## (Adjusted p values reported -- single-step method)
##
##
## Tukey results for gene: FBLN5
##
##   Simultaneous Tests for General Linear Hypotheses
##
## Multiple Comparisons of Means: Tukey Contrasts
##
##
## Fit: glm(formula = Protein_Count ~ Tissue, family = gaussian, data = gene_
data)
##
## Linear Hypotheses:
##                Estimate Std. Error z value Pr(>|z|)
## ML - MF == 0  3549180    8243993   0.431    0.667
## (Adjusted p values reported -- single-step method)
##
##
## Tukey results for gene: FBLN7
##
##   Simultaneous Tests for General Linear Hypotheses
##
## Multiple Comparisons of Means: Tukey Contrasts
##
```

```
##
## Fit: glm(formula = Protein_Count ~ Tissue, family = gaussian, data = gene_
data)
##
## Linear Hypotheses:
##               Estimate Std. Error z value Pr(>|z|)
## ML - MF == 0    89167     154373   0.578    0.564
## (Adjusted p values reported -- single-step method)
##
##
## Tukey results for gene: FBN1
##
##   Simultaneous Tests for General Linear Hypotheses
##
## Multiple Comparisons of Means: Tukey Contrasts
##
##
## Fit: glm(formula = Protein_Count ~ Tissue, family = gaussian, data = gene_
data)
##
## Linear Hypotheses:
##               Estimate Std. Error z value Pr(>|z|)
## ML - MF == 0   235226    3321996   0.071    0.944
## (Adjusted p values reported -- single-step method)
##
##
## Tukey results for gene: FBN2
##
##   Simultaneous Tests for General Linear Hypotheses
##
## Multiple Comparisons of Means: Tukey Contrasts
##
##
## Fit: glm(formula = Protein_Count ~ Tissue, family = gaussian, data = gene_
data)
##
## Linear Hypotheses:
##               Estimate Std. Error z value Pr(>|z|)
## ML - MF == 0   -19190     508469  -0.038     0.97
## (Adjusted p values reported -- single-step method)
##
##
## Tukey results for gene: FBN3
##
##   Simultaneous Tests for General Linear Hypotheses
##
## Multiple Comparisons of Means: Tukey Contrasts
##
##
## Fit: glm(formula = Protein_Count ~ Tissue, family = gaussian, data = gene_
```

```
data)
##
## Linear Hypotheses:
##                Estimate Std. Error z value Pr(>|z|)
## ML - MF == 0 -1045584     745665  -1.402    0.161
## (Adjusted p values reported -- single-step method)
##
##
## Tukey results for gene: FGA
##
##   Simultaneous Tests for General Linear Hypotheses
##
## Multiple Comparisons of Means: Tukey Contrasts
##
##
## Fit: glm(formula = Protein_Count ~ Tissue, family = gaussian, data = gene_
data)
##
## Linear Hypotheses:
##                Estimate Std. Error z value Pr(>|z|)
## ML - MF == 0 -1605860    3254688  -0.493    0.622
## (Adjusted p values reported -- single-step method)
##
##
## Tukey results for gene: FGB
##
##   Simultaneous Tests for General Linear Hypotheses
##
## Multiple Comparisons of Means: Tukey Contrasts
##
##
## Fit: glm(formula = Protein_Count ~ Tissue, family = gaussian, data = gene_
data)
##
## Linear Hypotheses:
##                Estimate Std. Error z value Pr(>|z|)
## ML - MF == 0 -3199360    5177869  -0.618    0.537
## (Adjusted p values reported -- single-step method)
##
##
## Tukey results for gene: FGG
##
##   Simultaneous Tests for General Linear Hypotheses
##
## Multiple Comparisons of Means: Tukey Contrasts
##
##
## Fit: glm(formula = Protein_Count ~ Tissue, family = gaussian, data = gene_
data)
##
```

```
## Linear Hypotheses:
##                Estimate Std. Error z value Pr(>|z|)
## ML - MF == 0 -1341776    1359936  -0.987    0.324
## (Adjusted p values reported -- single-step method)
##
##
## Tukey results for gene: FGL2
##
##   Simultaneous Tests for General Linear Hypotheses
##
## Multiple Comparisons of Means: Tukey Contrasts
##
##
## Fit: glm(formula = Protein_Count ~ Tissue, family = gaussian, data = gene_
data)
##
## Linear Hypotheses:
##                Estimate Std. Error z value Pr(>|z|)
## ML - MF == 0   109572     215444   0.509    0.611
## (Adjusted p values reported -- single-step method)
##
##
## Tukey results for gene: FN1
##
##   Simultaneous Tests for General Linear Hypotheses
##
## Multiple Comparisons of Means: Tukey Contrasts
##
##
## Fit: glm(formula = Protein_Count ~ Tissue, family = gaussian, data = gene_
data)
##
## Linear Hypotheses:
##                Estimate Std. Error z value Pr(>|z|)
## ML - MF == 0    19882    2076147    0.01    0.992
## (Adjusted p values reported -- single-step method)
##
##
## Tukey results for gene: FNDC1
##
##   Simultaneous Tests for General Linear Hypotheses
##
## Multiple Comparisons of Means: Tukey Contrasts
##
##
## Fit: glm(formula = Protein_Count ~ Tissue, family = gaussian, data = gene_
data)
##
## Linear Hypotheses:
##                Estimate Std. Error z value Pr(>|z|)
```

```
## ML - MF == 0    38340      25211   1.521    0.128
## (Adjusted p values reported -- single-step method)
##
##
## Tukey results for gene: IGFALS
##
##   Simultaneous Tests for General Linear Hypotheses
##
## Multiple Comparisons of Means: Tukey Contrasts
##
##
## Fit: glm(formula = Protein_Count ~ Tissue, family = gaussian, data = gene_
data)
##
## Linear Hypotheses:
##              Estimate Std. Error z value Pr(>|z|)
## ML - MF == 0   160746      70388   2.284   0.0224 *
## ---
## Signif. codes:  0 '***' 0.001 '**' 0.01 '*' 0.05 '.' 0.1 ' ' 1
## (Adjusted p values reported -- single-step method)
##
##
## Tukey results for gene: IGFBP5
##
##   Simultaneous Tests for General Linear Hypotheses
##
## Multiple Comparisons of Means: Tukey Contrasts
##
##
## Fit: glm(formula = Protein_Count ~ Tissue, family = gaussian, data = gene_
data)
##
## Linear Hypotheses:
##              Estimate Std. Error z value Pr(>|z|)
## ML - MF == 0   -43125     103012  -0.419    0.675
## (Adjusted p values reported -- single-step method)
##
##
## Tukey results for gene: IGFBP6
##
##   Simultaneous Tests for General Linear Hypotheses
##
## Multiple Comparisons of Means: Tukey Contrasts
##
##
## Fit: glm(formula = Protein_Count ~ Tissue, family = gaussian, data = gene_
data)
##
## Linear Hypotheses:
##              Estimate Std. Error z value Pr(>|z|)
```

```
## ML - MF == 0    94787      87077   1.089    0.276
## (Adjusted p values reported -- single-step method)
##
##
## Tukey results for gene: IGFBP7
##
##   Simultaneous Tests for General Linear Hypotheses
##
## Multiple Comparisons of Means: Tukey Contrasts
##
##
## Fit: glm(formula = Protein_Count ~ Tissue, family = gaussian, data = gene_
data)
##
## Linear Hypotheses:
##              Estimate Std. Error z value Pr(>|z|)
## ML - MF == 0   444451     564507   0.787    0.431
## (Adjusted p values reported -- single-step method)
##
##
## Tukey results for gene: IGSF10
##
##   Simultaneous Tests for General Linear Hypotheses
##
## Multiple Comparisons of Means: Tukey Contrasts
##
##
## Fit: glm(formula = Protein_Count ~ Tissue, family = gaussian, data = gene_
data)
##
## Linear Hypotheses:
##              Estimate Std. Error z value Pr(>|z|)
## ML - MF == 0    41055      33988   1.208    0.227
## (Adjusted p values reported -- single-step method)
##
##
## Tukey results for gene: LAMA1
##
##   Simultaneous Tests for General Linear Hypotheses
##
## Multiple Comparisons of Means: Tukey Contrasts
##
##
## Fit: glm(formula = Protein_Count ~ Tissue, family = gaussian, data = gene_
data)
##
## Linear Hypotheses:
##              Estimate Std. Error z value Pr(>|z|)
## ML - MF == 0  -181264     255225   -0.71    0.478
## (Adjusted p values reported -- single-step method)
```

```
##
##
## Tukey results for gene: LAMA2
##
##   Simultaneous Tests for General Linear Hypotheses
##
## Multiple Comparisons of Means: Tukey Contrasts
##
##
## Fit: glm(formula = Protein_Count ~ Tissue, family = gaussian, data = gene_
data)
##
## Linear Hypotheses:
##              Estimate Std. Error z value Pr(>|z|)
## ML - MF == 0   -80553      98662  -0.816    0.414
## (Adjusted p values reported -- single-step method)
##
##
## Tukey results for gene: LAMA3
##
##   Simultaneous Tests for General Linear Hypotheses
##
## Multiple Comparisons of Means: Tukey Contrasts
##
##
## Fit: glm(formula = Protein_Count ~ Tissue, family = gaussian, data = gene_
data)
##
## Linear Hypotheses:
##              Estimate Std. Error z value Pr(>|z|)
## ML - MF == 0   -13120     171667  -0.076    0.939
## (Adjusted p values reported -- single-step method)
##
##
## Tukey results for gene: LAMA4
##
##   Simultaneous Tests for General Linear Hypotheses
##
## Multiple Comparisons of Means: Tukey Contrasts
##
##
## Fit: glm(formula = Protein_Count ~ Tissue, family = gaussian, data = gene_
data)
##
## Linear Hypotheses:
##              Estimate Std. Error z value Pr(>|z|)
## ML - MF == 0  -470160    1341206  -0.351    0.726
## (Adjusted p values reported -- single-step method)
##
##
```

```
## Tukey results for gene: LAMA5
##
##   Simultaneous Tests for General Linear Hypotheses
##
## Multiple Comparisons of Means: Tukey Contrasts
##
##
## Fit: glm(formula = Protein_Count ~ Tissue, family = gaussian, data = gene_
data)
##
## Linear Hypotheses:
##               Estimate Std. Error z value Pr(>|z|)
## ML - MF == 0  -533840    1056154  -0.505    0.613
## (Adjusted p values reported -- single-step method)
##
##
## Tukey results for gene: LAMB1
##
##   Simultaneous Tests for General Linear Hypotheses
##
## Multiple Comparisons of Means: Tukey Contrasts
##
##
## Fit: glm(formula = Protein_Count ~ Tissue, family = gaussian, data = gene_
data)
##
## Linear Hypotheses:
##               Estimate Std. Error z value Pr(>|z|)
## ML - MF == 0  -576616    1024164  -0.563    0.573
## (Adjusted p values reported -- single-step method)
##
##
## Tukey results for gene: LAMB2
##
##   Simultaneous Tests for General Linear Hypotheses
##
## Multiple Comparisons of Means: Tukey Contrasts
##
##
## Fit: glm(formula = Protein_Count ~ Tissue, family = gaussian, data = gene_
data)
##
## Linear Hypotheses:
##               Estimate Std. Error z value Pr(>|z|)
## ML - MF == 0 -1224814    2896457  -0.423    0.672
## (Adjusted p values reported -- single-step method)
##
##
## Tukey results for gene: LAMC1
##
```

```
##   Simultaneous Tests for General Linear Hypotheses
##
## Multiple Comparisons of Means: Tukey Contrasts
##
##
## Fit: glm(formula = Protein_Count ~ Tissue, family = gaussian, data = gene_
data)
##
## Linear Hypotheses:
##              Estimate Std. Error z value Pr(>|z|)
## ML - MF == 0 -2192992    2924482   -0.75    0.453
## (Adjusted p values reported -- single-step method)
##
##
## Tukey results for gene: LAMC3
##
##   Simultaneous Tests for General Linear Hypotheses
##
## Multiple Comparisons of Means: Tukey Contrasts
##
##
## Fit: glm(formula = Protein_Count ~ Tissue, family = gaussian, data = gene_
data)
##
## Linear Hypotheses:
##              Estimate Std. Error z value Pr(>|z|)
## ML - MF == 0  -148836      37221  -3.999 6.37e-05 ***
## ---
## Signif. codes:  0 '***' 0.001 '**' 0.01 '*' 0.05 '.' 0.1 ' ' 1
## (Adjusted p values reported -- single-step method)
##
##
## Tukey results for gene: LRG1
##
##   Simultaneous Tests for General Linear Hypotheses
##
## Multiple Comparisons of Means: Tukey Contrasts
##
##
## Fit: glm(formula = Protein_Count ~ Tissue, family = gaussian, data = gene_
data)
##
## Linear Hypotheses:
##              Estimate Std. Error z value Pr(>|z|)
## ML - MF == 0   -71636      59176  -1.211    0.226
## (Adjusted p values reported -- single-step method)
##
##
## Tukey results for gene: LTBP1
##
```

```
##   Simultaneous Tests for General Linear Hypotheses
##
## Multiple Comparisons of Means: Tukey Contrasts
##
##
## Fit: glm(formula = Protein_Count ~ Tissue, family = gaussian, data = gene_
data)
##
## Linear Hypotheses:
##               Estimate Std. Error z value Pr(>|z|)
## ML - MF == 0   102080     488313   0.209    0.834
## (Adjusted p values reported -- single-step method)
##
##
## Tukey results for gene: LTBP2
##
##   Simultaneous Tests for General Linear Hypotheses
##
## Multiple Comparisons of Means: Tukey Contrasts
##
##
## Fit: glm(formula = Protein_Count ~ Tissue, family = gaussian, data = gene_
data)
##
## Linear Hypotheses:
##               Estimate Std. Error z value Pr(>|z|)
## ML - MF == 0   731848     710454    1.03    0.303
## (Adjusted p values reported -- single-step method)
##
##
## Tukey results for gene: LTBP3
##
##   Simultaneous Tests for General Linear Hypotheses
##
## Multiple Comparisons of Means: Tukey Contrasts
##
##
## Fit: glm(formula = Protein_Count ~ Tissue, family = gaussian, data = gene_
data)
##
## Linear Hypotheses:
##               Estimate Std. Error z value Pr(>|z|)
## ML - MF == 0    47942     123051    0.39    0.697
## (Adjusted p values reported -- single-step method)
##
##
## Tukey results for gene: LTBP4
##
##   Simultaneous Tests for General Linear Hypotheses
##
```

```
## Multiple Comparisons of Means: Tukey Contrasts
##
##
## Fit: glm(formula = Protein_Count ~ Tissue, family = gaussian, data = gene_
data)
##
## Linear Hypotheses:
##              Estimate Std. Error z value Pr(>|z|)
## ML - MF == 0  -247144    1896011   -0.13    0.896
## (Adjusted p values reported -- single-step method)
##
##
## Tukey results for gene: MATN2
##
##   Simultaneous Tests for General Linear Hypotheses
##
## Multiple Comparisons of Means: Tukey Contrasts
##
##
## Fit: glm(formula = Protein_Count ~ Tissue, family = gaussian, data = gene_
data)
##
## Linear Hypotheses:
##              Estimate Std. Error z value Pr(>|z|)
## ML - MF == 0   360202     467635    0.77    0.441
## (Adjusted p values reported -- single-step method)
##
##
## Tukey results for gene: MFAP2
##
##   Simultaneous Tests for General Linear Hypotheses
##
## Multiple Comparisons of Means: Tukey Contrasts
##
##
## Fit: glm(formula = Protein_Count ~ Tissue, family = gaussian, data = gene_
data)
##
## Linear Hypotheses:
##              Estimate Std. Error z value Pr(>|z|)
## ML - MF == 0   516692    3401194   0.152    0.879
## (Adjusted p values reported -- single-step method)
##
##
## Tukey results for gene: MFAP4
##
##   Simultaneous Tests for General Linear Hypotheses
##
## Multiple Comparisons of Means: Tukey Contrasts
##
```

```
##
## Fit: glm(formula = Protein_Count ~ Tissue, family = gaussian, data = gene_
data)
##
## Linear Hypotheses:
##                Estimate Std. Error z value Pr(>|z|)
## ML - MF == 0 -5732140    7603126  -0.754    0.451
## (Adjusted p values reported -- single-step method)
##
##
## Tukey results for gene: MFAP5
##
##   Simultaneous Tests for General Linear Hypotheses
##
## Multiple Comparisons of Means: Tukey Contrasts
##
##
## Fit: glm(formula = Protein_Count ~ Tissue, family = gaussian, data = gene_
data)
##
## Linear Hypotheses:
##                Estimate Std. Error z value Pr(>|z|)
## ML - MF == 0     9183     438755   0.021    0.983
## (Adjusted p values reported -- single-step method)
##
##
## Tukey results for gene: MFGE8
##
##   Simultaneous Tests for General Linear Hypotheses
##
## Multiple Comparisons of Means: Tukey Contrasts
##
##
## Fit: glm(formula = Protein_Count ~ Tissue, family = gaussian, data = gene_
data)
##
## Linear Hypotheses:
##                Estimate Std. Error z value Pr(>|z|)
## ML - MF == 0   -76374     203790  -0.375    0.708
## (Adjusted p values reported -- single-step method)
##
##
## Tukey results for gene: MGP
##
##   Simultaneous Tests for General Linear Hypotheses
##
## Multiple Comparisons of Means: Tukey Contrasts
##
##
## Fit: glm(formula = Protein_Count ~ Tissue, family = gaussian, data = gene_
```

```
data)
##
## Linear Hypotheses:
##                Estimate Std. Error z value Pr(>|z|)
## ML - MF == 0  -473838    3941359   -0.12    0.904
## (Adjusted p values reported -- single-step method)
##
##
## Tukey results for gene: MMRN2
##
##   Simultaneous Tests for General Linear Hypotheses
##
## Multiple Comparisons of Means: Tukey Contrasts
##
##
## Fit: glm(formula = Protein_Count ~ Tissue, family = gaussian, data = gene_
data)
##
## Linear Hypotheses:
##                Estimate Std. Error z value Pr(>|z|)
## ML - MF == 0   -59926      70934  -0.845    0.398
## (Adjusted p values reported -- single-step method)
##
##
## Tukey results for gene: MXRA5
##
##   Simultaneous Tests for General Linear Hypotheses
##
## Multiple Comparisons of Means: Tukey Contrasts
##
##
## Fit: glm(formula = Protein_Count ~ Tissue, family = gaussian, data = gene_
data)
##
## Linear Hypotheses:
##                Estimate Std. Error z value Pr(>|z|)
## ML - MF == 0   132406      35504   3.729 0.000192 ***
## ---
## Signif. codes:  0 '***' 0.001 '**' 0.01 '*' 0.05 '.' 0.1 ' ' 1
## (Adjusted p values reported -- single-step method)
##
##
## Tukey results for gene: NID1
##
##   Simultaneous Tests for General Linear Hypotheses
##
## Multiple Comparisons of Means: Tukey Contrasts
##
##
## Fit: glm(formula = Protein_Count ~ Tissue, family = gaussian, data = gene_
```

```
data)
##
## Linear Hypotheses:
##                Estimate Std. Error z value Pr(>|z|)
## ML - MF == 0 -9303980   13358446  -0.696    0.486
## (Adjusted p values reported -- single-step method)
##
##
## Tukey results for gene: NID2
##
##   Simultaneous Tests for General Linear Hypotheses
##
## Multiple Comparisons of Means: Tukey Contrasts
##
##
## Fit: glm(formula = Protein_Count ~ Tissue, family = gaussian, data = gene_
data)
##
## Linear Hypotheses:
##                Estimate Std. Error z value Pr(>|z|)
## ML - MF == 0 -3208000    3941210  -0.814    0.416
## (Adjusted p values reported -- single-step method)
##
##
## Tukey results for gene: NPNT
##
##   Simultaneous Tests for General Linear Hypotheses
##
## Multiple Comparisons of Means: Tukey Contrasts
##
##
## Fit: glm(formula = Protein_Count ~ Tissue, family = gaussian, data = gene_
data)
##
## Linear Hypotheses:
##                Estimate Std. Error z value Pr(>|z|)
## ML - MF == 0   128461      97505   1.317    0.188
## (Adjusted p values reported -- single-step method)
##
##
## Tukey results for gene: NTN4
##
##   Simultaneous Tests for General Linear Hypotheses
##
## Multiple Comparisons of Means: Tukey Contrasts
##
##
## Fit: glm(formula = Protein_Count ~ Tissue, family = gaussian, data = gene_
data)
##
```

```
## Linear Hypotheses:
##                Estimate Std. Error z value Pr(>|z|)
## ML - MF == 0   -40092      54540  -0.735    0.462
## (Adjusted p values reported -- single-step method)
##
##
## Tukey results for gene: PAPLN
##
##   Simultaneous Tests for General Linear Hypotheses
##
## Multiple Comparisons of Means: Tukey Contrasts
##
##
## Fit: glm(formula = Protein_Count ~ Tissue, family = gaussian, data = gene_
data)
##
## Linear Hypotheses:
##                Estimate Std. Error z value Pr(>|z|)
## ML - MF == 0   112134      85262   1.315    0.188
## (Adjusted p values reported -- single-step method)
##
##
## Tukey results for gene: PCOLCE
##
##   Simultaneous Tests for General Linear Hypotheses
##
## Multiple Comparisons of Means: Tukey Contrasts
##
##
## Fit: glm(formula = Protein_Count ~ Tissue, family = gaussian, data = gene_
data)
##
## Linear Hypotheses:
##                Estimate Std. Error z value Pr(>|z|)
## ML - MF == 0   342759     137780   2.488   0.0129 *
## ---
## Signif. codes:  0 '***' 0.001 '**' 0.01 '*' 0.05 '.' 0.1 ' ' 1
## (Adjusted p values reported -- single-step method)
##
##
## Tukey results for gene: POSTN
##
##   Simultaneous Tests for General Linear Hypotheses
##
## Multiple Comparisons of Means: Tukey Contrasts
##
##
## Fit: glm(formula = Protein_Count ~ Tissue, family = gaussian, data = gene_
data)
##
```

```
## Linear Hypotheses:
##              Estimate Std. Error z value Pr(>|z|)
## ML - MF == 0   104068     401096   0.259    0.795
## (Adjusted p values reported -- single-step method)
##
##
## Tukey results for gene: PXDN
##
##   Simultaneous Tests for General Linear Hypotheses
##
## Multiple Comparisons of Means: Tukey Contrasts
##
##
## Fit: glm(formula = Protein_Count ~ Tissue, family = gaussian, data = gene_
data)
##
## Linear Hypotheses:
##              Estimate Std. Error z value Pr(>|z|)
## ML - MF == 0   257635     147069   1.752   0.0798 .
## ---
## Signif. codes:  0 '***' 0.001 '**' 0.01 '*' 0.05 '.' 0.1 ' ' 1
## (Adjusted p values reported -- single-step method)
##
##
## Tukey results for gene: RSPO3
##
##   Simultaneous Tests for General Linear Hypotheses
##
## Multiple Comparisons of Means: Tukey Contrasts
##
##
## Fit: glm(formula = Protein_Count ~ Tissue, family = gaussian, data = gene_
data)
##
## Linear Hypotheses:
##              Estimate Std. Error z value Pr(>|z|)
## ML - MF == 0  -294797     482901   -0.61    0.542
## (Adjusted p values reported -- single-step method)
##
##
## Tukey results for gene: SBSPON
##
##   Simultaneous Tests for General Linear Hypotheses
##
## Multiple Comparisons of Means: Tukey Contrasts
##
##
## Fit: glm(formula = Protein_Count ~ Tissue, family = gaussian, data = gene_
data)
##
```

```
## Linear Hypotheses:
##              Estimate Std. Error z value Pr(>|z|)
## ML - MF == 0 -2628124    3628262  -0.724    0.469
## (Adjusted p values reported -- single-step method)
##
##
## Tukey results for gene: SMOC1
##
##   Simultaneous Tests for General Linear Hypotheses
##
## Multiple Comparisons of Means: Tukey Contrasts
##
##
## Fit: glm(formula = Protein_Count ~ Tissue, family = gaussian, data = gene_
data)
##
## Linear Hypotheses:
##              Estimate Std. Error z value Pr(>|z|)
## ML - MF == 0    71359     119148   0.599    0.549
## (Adjusted p values reported -- single-step method)
##
##
## Tukey results for gene: SMOC2
##
##   Simultaneous Tests for General Linear Hypotheses
##
## Multiple Comparisons of Means: Tukey Contrasts
##
##
## Fit: glm(formula = Protein_Count ~ Tissue, family = gaussian, data = gene_
data)
##
## Linear Hypotheses:
##              Estimate Std. Error z value Pr(>|z|)
## ML - MF == 0   -12670      50125  -0.253      0.8
## (Adjusted p values reported -- single-step method)
##
##
## Tukey results for gene: SPARC
##
##   Simultaneous Tests for General Linear Hypotheses
##
## Multiple Comparisons of Means: Tukey Contrasts
##
##
## Fit: glm(formula = Protein_Count ~ Tissue, family = gaussian, data = gene_
data)
##
## Linear Hypotheses:
##              Estimate Std. Error z value Pr(>|z|)
```

```
## ML - MF == 0  1252248    1238867   1.011    0.312
## (Adjusted p values reported -- single-step method)
##
##
## Tukey results for gene: SPARCL1
##
##   Simultaneous Tests for General Linear Hypotheses
##
## Multiple Comparisons of Means: Tukey Contrasts
##
##
## Fit: glm(formula = Protein_Count ~ Tissue, family = gaussian, data = gene_
data)
##
## Linear Hypotheses:
##              Estimate Std. Error z value Pr(>|z|)
## ML - MF == 0   138474     175024   0.791    0.429
## (Adjusted p values reported -- single-step method)
##
##
## Tukey results for gene: SPON1
##
##   Simultaneous Tests for General Linear Hypotheses
##
## Multiple Comparisons of Means: Tukey Contrasts
##
##
## Fit: glm(formula = Protein_Count ~ Tissue, family = gaussian, data = gene_
data)
##
## Linear Hypotheses:
##              Estimate Std. Error z value Pr(>|z|)
## ML - MF == 0   823502    2734496   0.301    0.763
## (Adjusted p values reported -- single-step method)
##
##
## Tukey results for gene: SPON2
##
##   Simultaneous Tests for General Linear Hypotheses
##
## Multiple Comparisons of Means: Tukey Contrasts
##
##
## Fit: glm(formula = Protein_Count ~ Tissue, family = gaussian, data = gene_
data)
##
## Linear Hypotheses:
##              Estimate Std. Error z value Pr(>|z|)
## ML - MF == 0   -24779      87776  -0.282    0.778
## (Adjusted p values reported -- single-step method)
```

```
##
##
## Tukey results for gene: SRPX
##
##   Simultaneous Tests for General Linear Hypotheses
##
## Multiple Comparisons of Means: Tukey Contrasts
##
##
## Fit: glm(formula = Protein_Count ~ Tissue, family = gaussian, data = gene_
data)
##
## Linear Hypotheses:
##              Estimate Std. Error z value Pr(>|z|)
## ML - MF == 0   181167     135570   1.336    0.181
## (Adjusted p values reported -- single-step method)
##
##
## Tukey results for gene: SRPX2
##
##   Simultaneous Tests for General Linear Hypotheses
##
## Multiple Comparisons of Means: Tukey Contrasts
##
##
## Fit: glm(formula = Protein_Count ~ Tissue, family = gaussian, data = gene_
data)
##
## Linear Hypotheses:
##              Estimate Std. Error z value Pr(>|z|)
## ML - MF == 0   245430     295998   0.829    0.407
## (Adjusted p values reported -- single-step method)
##
##
## Tukey results for gene: SVEP1
##
##   Simultaneous Tests for General Linear Hypotheses
##
## Multiple Comparisons of Means: Tukey Contrasts
##
##
## Fit: glm(formula = Protein_Count ~ Tissue, family = gaussian, data = gene_
data)
##
## Linear Hypotheses:
##              Estimate Std. Error z value Pr(>|z|)
## ML - MF == 0   142838     169746   0.841      0.4
## (Adjusted p values reported -- single-step method)
##
##
```

```
## Tukey results for gene: TGFBI
##
##   Simultaneous Tests for General Linear Hypotheses
##
## Multiple Comparisons of Means: Tukey Contrasts
##
##
## Fit: glm(formula = Protein_Count ~ Tissue, family = gaussian, data = gene_
data)
##
## Linear Hypotheses:
##              Estimate Std. Error z value Pr(>|z|)
## ML - MF == 0  4248980    5143211   0.826    0.409
## (Adjusted p values reported -- single-step method)
##
##
## Tukey results for gene: THBS1
##
##   Simultaneous Tests for General Linear Hypotheses
##
## Multiple Comparisons of Means: Tukey Contrasts
##
##
## Fit: glm(formula = Protein_Count ~ Tissue, family = gaussian, data = gene_
data)
##
## Linear Hypotheses:
##              Estimate Std. Error z value Pr(>|z|)
## ML - MF == 0   340194     405548   0.839    0.402
## (Adjusted p values reported -- single-step method)
##
##
## Tukey results for gene: THBS2
##
##   Simultaneous Tests for General Linear Hypotheses
##
## Multiple Comparisons of Means: Tukey Contrasts
##
##
## Fit: glm(formula = Protein_Count ~ Tissue, family = gaussian, data = gene_
data)
##
## Linear Hypotheses:
##              Estimate Std. Error z value Pr(>|z|)
## ML - MF == 0  -277255     468669  -0.592    0.554
## (Adjusted p values reported -- single-step method)
##
##
## Tukey results for gene: THBS3
##
```

```
##   Simultaneous Tests for General Linear Hypotheses
##
## Multiple Comparisons of Means: Tukey Contrasts
##
##
## Fit: glm(formula = Protein_Count ~ Tissue, family = gaussian, data = gene_
data)
##
## Linear Hypotheses:
##               Estimate Std. Error z value Pr(>|z|)
## ML - MF == 0    60077      38133   1.575    0.115
## (Adjusted p values reported -- single-step method)
##
##
## Tukey results for gene: THSD4
##
##   Simultaneous Tests for General Linear Hypotheses
##
## Multiple Comparisons of Means: Tukey Contrasts
##
##
## Fit: glm(formula = Protein_Count ~ Tissue, family = gaussian, data = gene_
data)
##
## Linear Hypotheses:
##               Estimate Std. Error z value Pr(>|z|)
## ML - MF == 0    63900     908781    0.07    0.944
## (Adjusted p values reported -- single-step method)
##
##
## Tukey results for gene: TINAGL1
##
##   Simultaneous Tests for General Linear Hypotheses
##
## Multiple Comparisons of Means: Tukey Contrasts
##
##
## Fit: glm(formula = Protein_Count ~ Tissue, family = gaussian, data = gene_
data)
##
## Linear Hypotheses:
##                Estimate Std. Error z value Pr(>|z|)
## ML - MF == 0 -17449600   22867362  -0.763    0.445
## (Adjusted p values reported -- single-step method)
##
##
## Tukey results for gene: TNC
##
##   Simultaneous Tests for General Linear Hypotheses
##
```

```
## Multiple Comparisons of Means: Tukey Contrasts
##
##
## Fit: glm(formula = Protein_Count ~ Tissue, family = gaussian, data = gene_
data)
##
## Linear Hypotheses:
##              Estimate Std. Error z value Pr(>|z|)
## ML - MF == 0   -22824     111217  -0.205    0.837
## (Adjusted p values reported -- single-step method)
##
##
## Tukey results for gene: TNXB
##
##   Simultaneous Tests for General Linear Hypotheses
##
## Multiple Comparisons of Means: Tukey Contrasts
##
##
## Fit: glm(formula = Protein_Count ~ Tissue, family = gaussian, data = gene_
data)
##
## Linear Hypotheses:
##              Estimate Std. Error z value Pr(>|z|)
## ML - MF == 0   604102     538525   1.122    0.262
## (Adjusted p values reported -- single-step method)
##
##
## Tukey results for gene: TSKU
##
##   Simultaneous Tests for General Linear Hypotheses
##
## Multiple Comparisons of Means: Tukey Contrasts
##
##
## Fit: glm(formula = Protein_Count ~ Tissue, family = gaussian, data = gene_
data)
##
## Linear Hypotheses:
##              Estimate Std. Error z value Pr(>|z|)
## ML - MF == 0   -39483      57481  -0.687    0.492
## (Adjusted p values reported -- single-step method)
##
##
## Tukey results for gene: VTN
##
##   Simultaneous Tests for General Linear Hypotheses
##
## Multiple Comparisons of Means: Tukey Contrasts
##
```

```
##
## Fit: glm(formula = Protein_Count ~ Tissue, family = gaussian, data = gene_
data)
##
## Linear Hypotheses:
##               Estimate Std. Error z value Pr(>|z|)
## ML - MF == 0 -2142800    3355086  -0.639    0.523
## (Adjusted p values reported -- single-step method)
##
##
## Tukey results for gene: VWA1
##
##   Simultaneous Tests for General Linear Hypotheses
##
## Multiple Comparisons of Means: Tukey Contrasts
##
##
## Fit: glm(formula = Protein_Count ~ Tissue, family = gaussian, data = gene_
data)
##
## Linear Hypotheses:
##               Estimate Std. Error z value Pr(>|z|)
## ML - MF == 0  1096978    1453246   0.755     0.45
## (Adjusted p values reported -- single-step method)
##
##
## Tukey results for gene: VWA5A
##
##   Simultaneous Tests for General Linear Hypotheses
##
## Multiple Comparisons of Means: Tukey Contrasts
##
##
## Fit: glm(formula = Protein_Count ~ Tissue, family = gaussian, data = gene_
data)
##
## Linear Hypotheses:
##               Estimate Std. Error z value Pr(>|z|)
## ML - MF == 0    24784      82060   0.302    0.763
## (Adjusted p values reported -- single-step method)
##
##
## Tukey results for gene: VWF
##
##   Simultaneous Tests for General Linear Hypotheses
##
## Multiple Comparisons of Means: Tukey Contrasts
##
##
## Fit: glm(formula = Protein_Count ~ Tissue, family = gaussian, data = gene_
```

```
data)
##
## Linear Hypotheses:
##              Estimate Std. Error z value Pr(>|z|)
## ML - MF == 0   -51864     156404  -0.332     0.74
## (Adjusted p values reported -- single-step method)

#### Proteoglycans

# Read your cleaned joined data file into a data frame
cleaned_joined_data <- read.csv("cleaned_matrisome_medu.csv")

# Filter rows with Matrisome_Category "Proteoglycans"
Proteoglycans_data <- cleaned_joined_data %>%
  filter(Matrisome_Category == "Proteoglycans")

# Reshape the data to have separate columns for each replicate and each tissu
e stage
organized_data <- Proteoglycans_data %>%
  pivot_longer(cols = starts_with("MF") | starts_with("ML"),
               names_to = c("Tissue", "Stage"),
               names_sep = "\\.",
               values_to = "Protein_Count")

## Warning: Expected 2 pieces. Missing pieces filled with `NA` in 2 rows [1,
6].

# Save the reshaped data to a new file

write.csv(organized_data, "Proteoglycans_data_matrisome_medu_to_clean.csv", r
ow.names = FALSE)

data2 <- read.csv("Proteoglycans_data_matrisome_medu_clean.csv")

png(filename="medu_Proteoglycans.png",
    width = 12, height = 6, units = "in",
    res = 600)

boxplot(Protein_Count ~ Tissue*Genes,
        data = data2,
        cex.axis = 0.8,
        col = viridis(2, alpha = 0.7),
        lwd = 0.7,
        ylab = "",
        ylim = c(0, 400000000),
        axes = FALSE,
        frame.plot = TRUE,
        xlab = "")

GROUP <- unique(data2$Genes)
```

```r
groups <- 15
numbox <- 2
total <- groups * numbox
xpoints <- seq(median(1:numbox), total, numbox)

axis(2, seq(0, 400000000, by = 20000000), las = 1, cex.axis = 0.7)
axis(1, labels = GROUP, at = xpoints, las = 2, cex.axis = 0.7)

stripchart(data2$Protein_Count ~ data2$Tissue*data2$Genes,
           method = "jitter",
           pch = 19,
           col = "black",
           vertical = TRUE,
           add = TRUE,
           cex = 0.3)

title(ylab = expression("Protein abundance (RU)"), line = 3.0)
legend("top", fill = viridis(2), legend = c("Medulla Follicular", "Medulla Lu
teal"), horiz = T, cex = 0.7)

dev.off()

## quartz_off_screen
##                  2

# Convert Tissue to factor
data2$Tissue <- as.factor(data2$Tissue)

# Create an empty list to store Tukey results
tukey_results_list <- list()

# Unique genes
unique_genes <- unique(data2$Genes)

# Loop over each gene
for (gene in unique_genes) {
  # Subset data for the current gene
  gene_data <- subset(data2, Genes == gene)

  # Fit the generalized linear model
  glm_model <- glm(Protein_Count ~ Tissue, data = gene_data, family = gaussia
n)

  # Perform Tukey test
  contrast_matrix <- glht(glm_model, linfct = mcp(Tissue = "Tukey"))
  tukey_results <- summary(contrast_matrix)

  # Store the results
  tukey_results_list[[gene]] <- tukey_results
```

```
}

# Print the results for each gene
for (gene in unique_genes) {
  cat("Tukey results for gene:", gene, "\n")
  print(tukey_results_list[[gene]])
  cat("\n")
}

## Tukey results for gene: ACAN
##
##   Simultaneous Tests for General Linear Hypotheses
##
## Multiple Comparisons of Means: Tukey Contrasts
##
##
## Fit: glm(formula = Protein_Count ~ Tissue, family = gaussian, data = gene_
data)
##
## Linear Hypotheses:
##                Estimate Std. Error z value Pr(>|z|)
## ML - MF == 0  -126902     705102   -0.18    0.857
## (Adjusted p values reported -- single-step method)
##
##
## Tukey results for gene: ASPN
##
##   Simultaneous Tests for General Linear Hypotheses
##
## Multiple Comparisons of Means: Tukey Contrasts
##
##
## Fit: glm(formula = Protein_Count ~ Tissue, family = gaussian, data = gene_
data)
##
## Linear Hypotheses:
##                Estimate Std. Error z value Pr(>|z|)
## ML - MF == 0  5879840    9714097   0.605    0.545
## (Adjusted p values reported -- single-step method)
##
##
## Tukey results for gene: BGN
##
##   Simultaneous Tests for General Linear Hypotheses
##
## Multiple Comparisons of Means: Tukey Contrasts
##
##
## Fit: glm(formula = Protein_Count ~ Tissue, family = gaussian, data = gene_
data)
```

```
##
## Linear Hypotheses:
##               Estimate Std. Error z value Pr(>|z|)
## ML - MF == 0  1241014    1910118    0.65    0.516
## (Adjusted p values reported -- single-step method)
##
##
## Tukey results for gene: DCN
##
##   Simultaneous Tests for General Linear Hypotheses
##
## Multiple Comparisons of Means: Tukey Contrasts
##
##
## Fit: glm(formula = Protein_Count ~ Tissue, family = gaussian, data = gene_
data)
##
## Linear Hypotheses:
##               Estimate Std. Error z value Pr(>|z|)
## ML - MF == 0  2898486    2379302   1.218    0.223
## (Adjusted p values reported -- single-step method)
##
##
## Tukey results for gene: FMOD
##
##   Simultaneous Tests for General Linear Hypotheses
##
## Multiple Comparisons of Means: Tukey Contrasts
##
##
## Fit: glm(formula = Protein_Count ~ Tissue, family = gaussian, data = gene_
data)
##
## Linear Hypotheses:
##               Estimate Std. Error z value Pr(>|z|)
## ML - MF == 0 19978720   43221422   0.462    0.644
## (Adjusted p values reported -- single-step method)
##
##
## Tukey results for gene: HAPLN1
##
##   Simultaneous Tests for General Linear Hypotheses
##
## Multiple Comparisons of Means: Tukey Contrasts
##
##
## Fit: glm(formula = Protein_Count ~ Tissue, family = gaussian, data = gene_
data)
##
## Linear Hypotheses:
```

```
##               Estimate Std. Error z value Pr(>|z|)
## ML - MF == 0   110021     276806   0.397    0.691
## (Adjusted p values reported -- single-step method)
##
##
## Tukey results for gene: HAPLN3
##
##   Simultaneous Tests for General Linear Hypotheses
##
## Multiple Comparisons of Means: Tukey Contrasts
##
##
## Fit: glm(formula = Protein_Count ~ Tissue, family = gaussian, data = gene_
data)
##
## Linear Hypotheses:
##               Estimate Std. Error z value Pr(>|z|)
## ML - MF == 0  -250402     275014  -0.911    0.363
## (Adjusted p values reported -- single-step method)
##
##
## Tukey results for gene: HSPG2
##
##   Simultaneous Tests for General Linear Hypotheses
##
## Multiple Comparisons of Means: Tukey Contrasts
##
##
## Fit: glm(formula = Protein_Count ~ Tissue, family = gaussian, data = gene_
data)
##
## Linear Hypotheses:
##               Estimate Std. Error z value Pr(>|z|)
## ML - MF == 0   -82740    4520167  -0.018    0.985
## (Adjusted p values reported -- single-step method)
##
##
## Tukey results for gene: KERA
##
##   Simultaneous Tests for General Linear Hypotheses
##
## Multiple Comparisons of Means: Tukey Contrasts
##
##
## Fit: glm(formula = Protein_Count ~ Tissue, family = gaussian, data = gene_
data)
##
## Linear Hypotheses:
##               Estimate Std. Error z value Pr(>|z|)
## ML - MF == 0    98312      59296   1.658   0.0973 .
```

```
## ---
## Signif. codes:  0 '***' 0.001 '**' 0.01 '*' 0.05 '.' 0.1 ' ' 1
## (Adjusted p values reported -- single-step method)
##
##
## Tukey results for gene: LUM
##
##   Simultaneous Tests for General Linear Hypotheses
##
## Multiple Comparisons of Means: Tukey Contrasts
##
##
## Fit: glm(formula = Protein_Count ~ Tissue, family = gaussian, data = gene_
data)
##
## Linear Hypotheses:
##              Estimate Std. Error z value Pr(>|z|)
## ML - MF == 0  1029600   54682215   0.019    0.985
## (Adjusted p values reported -- single-step method)
##
##
## Tukey results for gene: OGN
##
##   Simultaneous Tests for General Linear Hypotheses
##
## Multiple Comparisons of Means: Tukey Contrasts
##
##
## Fit: glm(formula = Protein_Count ~ Tissue, family = gaussian, data = gene_
data)
##
## Linear Hypotheses:
##              Estimate Std. Error z value Pr(>|z|)
## ML - MF == 0 10579200    9210030   1.149    0.251
## (Adjusted p values reported -- single-step method)
##
##
## Tukey results for gene: PODN
##
##   Simultaneous Tests for General Linear Hypotheses
##
## Multiple Comparisons of Means: Tukey Contrasts
##
##
## Fit: glm(formula = Protein_Count ~ Tissue, family = gaussian, data = gene_
data)
##
## Linear Hypotheses:
##              Estimate Std. Error z value Pr(>|z|)
## ML - MF == 0   172250     216865   0.794    0.427
```

```
## (Adjusted p values reported -- single-step method)
##
##
## Tukey results for gene: PODNL1
##
##   Simultaneous Tests for General Linear Hypotheses
##
## Multiple Comparisons of Means: Tukey Contrasts
##
##
## Fit: glm(formula = Protein_Count ~ Tissue, family = gaussian, data = gene_
data)
##
## Linear Hypotheses:
##              Estimate Std. Error z value Pr(>|z|)
## ML - MF == 0    -5470      58072  -0.094    0.925
## (Adjusted p values reported -- single-step method)
##
##
## Tukey results for gene: PRELP
##
##   Simultaneous Tests for General Linear Hypotheses
##
## Multiple Comparisons of Means: Tukey Contrasts
##
##
## Fit: glm(formula = Protein_Count ~ Tissue, family = gaussian, data = gene_
data)
##
## Linear Hypotheses:
##              Estimate Std. Error z value Pr(>|z|)
## ML - MF == 0  3098420   18213927    0.17    0.865
## (Adjusted p values reported -- single-step method)
##
##
## Tukey results for gene: VCAN
##
##   Simultaneous Tests for General Linear Hypotheses
##
## Multiple Comparisons of Means: Tukey Contrasts
##
##
## Fit: glm(formula = Protein_Count ~ Tissue, family = gaussian, data = gene_
data)
##
## Linear Hypotheses:
##              Estimate Std. Error z value Pr(>|z|)
## ML - MF == 0 -6403820    6591325  -0.972    0.331
## (Adjusted p values reported -- single-step method)
```

```r
#### ECM regulators

# Read your cleaned joined data file into a data frame
cleaned_joined_data <- read.csv("cleaned_matrisome_medu.csv")

# Filter rows with Matrisome_Category "ECM regulators"
ECM_regulators_data <- cleaned_joined_data %>%
  filter(Matrisome_Category == "ECM Regulators")

# Reshape the data to have separate columns for each replicate and each tissu
e stage
organized_data <- ECM_regulators_data %>%
  pivot_longer(cols = starts_with("MF") | starts_with("ML"),
               names_to = c("Tissue", "Stage"),
               names_sep = "\\.",
               values_to = "Protein_Count")

## Warning: Expected 2 pieces. Missing pieces filled with `NA` in 2 rows [1,
6].

# Save the reshaped data to a new file

write.csv(organized_data, "ECM regulators_data_matrisome_medu_to_clean.csv",
row.names = FALSE)

data2 <- read.csv("ECM regulators_data_matrisome_medu_clean.csv")

png(filename="medu_ECM regulators.png",
    width = 12, height = 6, units = "in",
    res = 600)

boxplot(Protein_Count ~ Tissue*Genes,
        data = data2,
        cex.axis = 0.8,
        col = viridis(2, alpha = 0.7),
        lwd = 0.7,
        ylab = "",
        ylim = c(0, 350000000),
        axes = FALSE,
        frame.plot = TRUE,
        xlab = "")

GROUP <- unique(data2$Genes)
groups <- 70
numbox <- 2
total <- groups * numbox
xpoints <- seq(median(1:numbox), total, numbox)

axis(2, seq(0, 350000000, by = 10000000), las = 1, cex.axis = 0.7)
```

```
axis(1, labels = GROUP, at = xpoints, las = 2, cex.axis = 0.7)

stripchart(data2$Protein_Count ~ data2$Tissue*data2$Genes,
           method = "jitter",
           pch = 19,
           col = "black",
           vertical = TRUE,
           add = TRUE,
           cex = 0.3)

title(ylab = expression("Protein abundance (RU)"), line = 3.0)
legend("top", fill = viridis(2), legend = c("Medulla Follicular", "Medulla Lu
teal"), horiz = T, cex = 0.7)

dev.off()

## quartz_off_screen
##                 2

# Convert Tissue to factor
data2$Tissue <- as.factor(data2$Tissue)

# Create an empty list to store Tukey results
tukey_results_list <- list()

# Unique genes
unique_genes <- unique(data2$Genes)

# Loop over each gene
for (gene in unique_genes) {
  # Subset data for the current gene
  gene_data <- subset(data2, Genes == gene)

  # Fit the generalized linear model
  glm_model <- glm(Protein_Count ~ Tissue, data = gene_data, family = gaussia
n)

  # Perform Tukey test
  contrast_matrix <- glht(glm_model, linfct = mcp(Tissue = "Tukey"))
  tukey_results <- summary(contrast_matrix)

  # Store the results
  tukey_results_list[[gene]] <- tukey_results
}

# Print the results for each gene
for (gene in unique_genes) {
  cat("Tukey results for gene:", gene, "\n")
  print(tukey_results_list[[gene]])
```

```
  cat("\n")
}

## Tukey results for gene: A2M
##
##   Simultaneous Tests for General Linear Hypotheses
##
## Multiple Comparisons of Means: Tukey Contrasts
##
##
## Fit: glm(formula = Protein_Count ~ Tissue, family = gaussian, data = gene_
data)
##
## Linear Hypotheses:
##                Estimate Std. Error z value Pr(>|z|)
## ML - MF == 0  -852086     797576  -1.068    0.285
## (Adjusted p values reported -- single-step method)
##
##
## Tukey results for gene: ADAM10
##
##   Simultaneous Tests for General Linear Hypotheses
##
## Multiple Comparisons of Means: Tukey Contrasts
##
##
## Fit: glm(formula = Protein_Count ~ Tissue, family = gaussian, data = gene_
data)
##
## Linear Hypotheses:
##                Estimate Std. Error z value Pr(>|z|)
## ML - MF == 0    39588     121177   0.327    0.744
## (Adjusted p values reported -- single-step method)
##
##
## Tukey results for gene: ADAMTSL1
##
##   Simultaneous Tests for General Linear Hypotheses
##
## Multiple Comparisons of Means: Tukey Contrasts
##
##
## Fit: glm(formula = Protein_Count ~ Tissue, family = gaussian, data = gene_
data)
##
## Linear Hypotheses:
##                Estimate Std. Error z value Pr(>|z|)
## ML - MF == 0    22458      46017   0.488    0.626
## (Adjusted p values reported -- single-step method)
##
```

```
##
## Tukey results for gene: ADAMTSL4
##
##   Simultaneous Tests for General Linear Hypotheses
##
## Multiple Comparisons of Means: Tukey Contrasts
##
##
## Fit: glm(formula = Protein_Count ~ Tissue, family = gaussian, data = gene_
data)
##
## Linear Hypotheses:
##              Estimate Std. Error z value Pr(>|z|)
## ML - MF == 0   123802      91630   1.351    0.177
## (Adjusted p values reported -- single-step method)
##
##
## Tukey results for gene: ADAMTSL5
##
##   Simultaneous Tests for General Linear Hypotheses
##
## Multiple Comparisons of Means: Tukey Contrasts
##
##
## Fit: glm(formula = Protein_Count ~ Tissue, family = gaussian, data = gene_
data)
##
## Linear Hypotheses:
##              Estimate Std. Error z value Pr(>|z|)
## ML - MF == 0   -31838     168131  -0.189     0.85
## (Adjusted p values reported -- single-step method)
##
##
## Tukey results for gene: AGT
##
##   Simultaneous Tests for General Linear Hypotheses
##
## Multiple Comparisons of Means: Tukey Contrasts
##
##
## Fit: glm(formula = Protein_Count ~ Tissue, family = gaussian, data = gene_
data)
##
## Linear Hypotheses:
##              Estimate Std. Error z value Pr(>|z|)
## ML - MF == 0     1970      95875   0.021    0.984
## (Adjusted p values reported -- single-step method)
##
##
## Tukey results for gene: CD109
```

```
##
##   Simultaneous Tests for General Linear Hypotheses
##
## Multiple Comparisons of Means: Tukey Contrasts
##
##
## Fit: glm(formula = Protein_Count ~ Tissue, family = gaussian, data = gene_
data)
##
## Linear Hypotheses:
##              Estimate Std. Error z value Pr(>|z|)
## ML - MF == 0    77158     106988   0.721    0.471
## (Adjusted p values reported -- single-step method)
##
##
## Tukey results for gene: CELA1
##
##   Simultaneous Tests for General Linear Hypotheses
##
## Multiple Comparisons of Means: Tukey Contrasts
##
##
## Fit: glm(formula = Protein_Count ~ Tissue, family = gaussian, data = gene_
data)
##
## Linear Hypotheses:
##              Estimate Std. Error z value Pr(>|z|)
## ML - MF == 0    52986      89728   0.591    0.555
## (Adjusted p values reported -- single-step method)
##
##
## Tukey results for gene: CELA3B
##
##   Simultaneous Tests for General Linear Hypotheses
##
## Multiple Comparisons of Means: Tukey Contrasts
##
##
## Fit: glm(formula = Protein_Count ~ Tissue, family = gaussian, data = gene_
data)
##
## Linear Hypotheses:
##              Estimate Std. Error z value Pr(>|z|)
## ML - MF == 0 -1041341    3286664  -0.317    0.751
## (Adjusted p values reported -- single-step method)
##
##
## Tukey results for gene: CST6
##
##   Simultaneous Tests for General Linear Hypotheses
```

```
##
## Multiple Comparisons of Means: Tukey Contrasts
##
##
## Fit: glm(formula = Protein_Count ~ Tissue, family = gaussian, data = gene_
data)
##
## Linear Hypotheses:
##              Estimate Std. Error z value Pr(>|z|)
## ML - MF == 0  -306130     343637  -0.891    0.373
## (Adjusted p values reported -- single-step method)
##
##
## Tukey results for gene: CSTB
##
##   Simultaneous Tests for General Linear Hypotheses
##
## Multiple Comparisons of Means: Tukey Contrasts
##
##
## Fit: glm(formula = Protein_Count ~ Tissue, family = gaussian, data = gene_
data)
##
## Linear Hypotheses:
##              Estimate Std. Error z value Pr(>|z|)
## ML - MF == 0   147566     161878   0.912    0.362
## (Adjusted p values reported -- single-step method)
##
##
## Tukey results for gene: CTSC
##
##   Simultaneous Tests for General Linear Hypotheses
##
## Multiple Comparisons of Means: Tukey Contrasts
##
##
## Fit: glm(formula = Protein_Count ~ Tissue, family = gaussian, data = gene_
data)
##
## Linear Hypotheses:
##              Estimate Std. Error z value Pr(>|z|)
## ML - MF == 0    38592      40609    0.95    0.342
## (Adjusted p values reported -- single-step method)
##
##
## Tukey results for gene: CTSD
##
##   Simultaneous Tests for General Linear Hypotheses
##
## Multiple Comparisons of Means: Tukey Contrasts
```

```
##
##
## Fit: glm(formula = Protein_Count ~ Tissue, family = gaussian, data = gene_
data)
##
## Linear Hypotheses:
##               Estimate Std. Error z value Pr(>|z|)
## ML - MF == 0  -191235     172153  -1.111    0.267
## (Adjusted p values reported -- single-step method)
##
##
## Tukey results for gene: CTSF
##
##   Simultaneous Tests for General Linear Hypotheses
##
## Multiple Comparisons of Means: Tukey Contrasts
##
##
## Fit: glm(formula = Protein_Count ~ Tissue, family = gaussian, data = gene_
data)
##
## Linear Hypotheses:
##               Estimate Std. Error z value Pr(>|z|)
## ML - MF == 0  -182419     282581  -0.646    0.519
## (Adjusted p values reported -- single-step method)
##
##
## Tukey results for gene: CTSH
##
##   Simultaneous Tests for General Linear Hypotheses
##
## Multiple Comparisons of Means: Tukey Contrasts
##
##
## Fit: glm(formula = Protein_Count ~ Tissue, family = gaussian, data = gene_
data)
##
## Linear Hypotheses:
##               Estimate Std. Error z value Pr(>|z|)
## ML - MF == 0   376994     315318   1.196    0.232
## (Adjusted p values reported -- single-step method)
##
##
## Tukey results for gene: CTSS
##
##   Simultaneous Tests for General Linear Hypotheses
##
## Multiple Comparisons of Means: Tukey Contrasts
##
##
```

```
## Fit: glm(formula = Protein_Count ~ Tissue, family = gaussian, data = gene_
data)
##
## Linear Hypotheses:
##                Estimate Std. Error z value Pr(>|z|)
## ML - MF == 0    15243      82331   0.185    0.853
## (Adjusted p values reported -- single-step method)
##
##
## Tukey results for gene: CTSW
##
##   Simultaneous Tests for General Linear Hypotheses
##
## Multiple Comparisons of Means: Tukey Contrasts
##
##
## Fit: glm(formula = Protein_Count ~ Tissue, family = gaussian, data = gene_
data)
##
## Linear Hypotheses:
##                Estimate Std. Error z value Pr(>|z|)
## ML - MF == 0   -28826     137080   -0.21    0.833
## (Adjusted p values reported -- single-step method)
##
##
## Tukey results for gene: CTSZ
##
##   Simultaneous Tests for General Linear Hypotheses
##
## Multiple Comparisons of Means: Tukey Contrasts
##
##
## Fit: glm(formula = Protein_Count ~ Tissue, family = gaussian, data = gene_
data)
##
## Linear Hypotheses:
##                Estimate Std. Error z value Pr(>|z|)
## ML - MF == 0   -51824     104630  -0.495     0.62
## (Adjusted p values reported -- single-step method)
##
##
## Tukey results for gene: F10
##
##   Simultaneous Tests for General Linear Hypotheses
##
## Multiple Comparisons of Means: Tukey Contrasts
##
##
## Fit: glm(formula = Protein_Count ~ Tissue, family = gaussian, data = gene_
data)
```

```
##
## Linear Hypotheses:
##                Estimate Std. Error z value Pr(>|z|)
## ML - MF == 0   -73889     171872   -0.43    0.667
## (Adjusted p values reported -- single-step method)
##
##
## Tukey results for gene: F12
##
##   Simultaneous Tests for General Linear Hypotheses
##
## Multiple Comparisons of Means: Tukey Contrasts
##
##
## Fit: glm(formula = Protein_Count ~ Tissue, family = gaussian, data = gene_
data)
##
## Linear Hypotheses:
##                Estimate Std. Error z value Pr(>|z|)
## ML - MF == 0    46396      55041   0.843    0.399
## (Adjusted p values reported -- single-step method)
##
##
## Tukey results for gene: F13B
##
##   Simultaneous Tests for General Linear Hypotheses
##
## Multiple Comparisons of Means: Tukey Contrasts
##
##
## Fit: glm(formula = Protein_Count ~ Tissue, family = gaussian, data = gene_
data)
##
## Linear Hypotheses:
##                Estimate Std. Error z value Pr(>|z|)
## ML - MF == 0   -53436      69585  -0.768    0.443
## (Adjusted p values reported -- single-step method)
##
##
## Tukey results for gene: F2
##
##   Simultaneous Tests for General Linear Hypotheses
##
## Multiple Comparisons of Means: Tukey Contrasts
##
##
## Fit: glm(formula = Protein_Count ~ Tissue, family = gaussian, data = gene_
data)
##
## Linear Hypotheses:
```

```
##               Estimate Std. Error z value Pr(>|z|)
## ML - MF == 0  -317234     445475  -0.712    0.476
## (Adjusted p values reported -- single-step method)
##
##
## Tukey results for gene: F9
##
##   Simultaneous Tests for General Linear Hypotheses
##
## Multiple Comparisons of Means: Tukey Contrasts
##
##
## Fit: glm(formula = Protein_Count ~ Tissue, family = gaussian, data = gene_
data)
##
## Linear Hypotheses:
##               Estimate Std. Error z value Pr(>|z|)
## ML - MF == 0    -1304     152473  -0.009    0.993
## (Adjusted p values reported -- single-step method)
##
##
## Tukey results for gene: FETUB
##
##   Simultaneous Tests for General Linear Hypotheses
##
## Multiple Comparisons of Means: Tukey Contrasts
##
##
## Fit: glm(formula = Protein_Count ~ Tissue, family = gaussian, data = gene_
data)
##
## Linear Hypotheses:
##               Estimate Std. Error z value Pr(>|z|)
## ML - MF == 0  -178233      92010  -1.937   0.0527 .
## ---
## Signif. codes:  0 '***' 0.001 '**' 0.01 '*' 0.05 '.' 0.1 ' ' 1
## (Adjusted p values reported -- single-step method)
##
##
## Tukey results for gene: HABP2
##
##   Simultaneous Tests for General Linear Hypotheses
##
## Multiple Comparisons of Means: Tukey Contrasts
##
##
## Fit: glm(formula = Protein_Count ~ Tissue, family = gaussian, data = gene_
data)
##
## Linear Hypotheses:
```

```
##                Estimate Std. Error z value Pr(>|z|)
## ML - MF == 0     7440      64984   0.114    0.909
## (Adjusted p values reported -- single-step method)
##
##
## Tukey results for gene: HRG
##
##   Simultaneous Tests for General Linear Hypotheses
##
## Multiple Comparisons of Means: Tukey Contrasts
##
##
## Fit: glm(formula = Protein_Count ~ Tissue, family = gaussian, data = gene_
data)
##
## Linear Hypotheses:
##                Estimate Std. Error z value Pr(>|z|)
## ML - MF == 0  1473546    1456583   1.012    0.312
## (Adjusted p values reported -- single-step method)
##
##
## Tukey results for gene: HTRA1
##
##   Simultaneous Tests for General Linear Hypotheses
##
## Multiple Comparisons of Means: Tukey Contrasts
##
##
## Fit: glm(formula = Protein_Count ~ Tissue, family = gaussian, data = gene_
data)
##
## Linear Hypotheses:
##                Estimate Std. Error z value Pr(>|z|)
## ML - MF == 0   -11888      74696  -0.159    0.874
## (Adjusted p values reported -- single-step method)
##
##
## Tukey results for gene: HTRA3
##
##   Simultaneous Tests for General Linear Hypotheses
##
## Multiple Comparisons of Means: Tukey Contrasts
##
##
## Fit: glm(formula = Protein_Count ~ Tissue, family = gaussian, data = gene_
data)
##
## Linear Hypotheses:
##                Estimate Std. Error z value Pr(>|z|)
## ML - MF == 0     4818      91336   0.053    0.958
```

```
## (Adjusted p values reported -- single-step method)
##
##
## Tukey results for gene: HTRA4
##
##   Simultaneous Tests for General Linear Hypotheses
##
## Multiple Comparisons of Means: Tukey Contrasts
##
##
## Fit: glm(formula = Protein_Count ~ Tissue, family = gaussian, data = gene_
data)
##
## Linear Hypotheses:
##              Estimate Std. Error z value Pr(>|z|)
## ML - MF == 0     5849      29106   0.201    0.841
## (Adjusted p values reported -- single-step method)
##
##
## Tukey results for gene: ITIH1
##
##   Simultaneous Tests for General Linear Hypotheses
##
## Multiple Comparisons of Means: Tukey Contrasts
##
##
## Fit: glm(formula = Protein_Count ~ Tissue, family = gaussian, data = gene_
data)
##
## Linear Hypotheses:
##              Estimate Std. Error z value Pr(>|z|)
## ML - MF == 0 -1093792     893743  -1.224    0.221
## (Adjusted p values reported -- single-step method)
##
##
## Tukey results for gene: ITIH2
##
##   Simultaneous Tests for General Linear Hypotheses
##
## Multiple Comparisons of Means: Tukey Contrasts
##
##
## Fit: glm(formula = Protein_Count ~ Tissue, family = gaussian, data = gene_
data)
##
## Linear Hypotheses:
##              Estimate Std. Error z value Pr(>|z|)
## ML - MF == 0  -318428    1121154  -0.284    0.776
## (Adjusted p values reported -- single-step method)
##
```

```
##
## Tukey results for gene: ITIH3
##
##   Simultaneous Tests for General Linear Hypotheses
##
## Multiple Comparisons of Means: Tukey Contrasts
##
##
## Fit: glm(formula = Protein_Count ~ Tissue, family = gaussian, data = gene_
data)
##
## Linear Hypotheses:
##                Estimate Std. Error z value Pr(>|z|)
## ML - MF == 0    52180      52482   0.994     0.32
## (Adjusted p values reported -- single-step method)
##
##
## Tukey results for gene: ITIH4
##
##   Simultaneous Tests for General Linear Hypotheses
##
## Multiple Comparisons of Means: Tukey Contrasts
##
##
## Fit: glm(formula = Protein_Count ~ Tissue, family = gaussian, data = gene_
data)
##
## Linear Hypotheses:
##                Estimate Std. Error z value Pr(>|z|)
## ML - MF == 0 -1869198    1042577  -1.793    0.073 .
## ---
## Signif. codes:  0 '***' 0.001 '**' 0.01 '*' 0.05 '.' 0.1 ' ' 1
## (Adjusted p values reported -- single-step method)
##
##
## Tukey results for gene: ITIH5
##
##   Simultaneous Tests for General Linear Hypotheses
##
## Multiple Comparisons of Means: Tukey Contrasts
##
##
## Fit: glm(formula = Protein_Count ~ Tissue, family = gaussian, data = gene_
data)
##
## Linear Hypotheses:
##                Estimate Std. Error z value Pr(>|z|)
## ML - MF == 0   -10293      36126  -0.285    0.776
## (Adjusted p values reported -- single-step method)
##
```

```
##
## Tukey results for gene: KNG1
##
##   Simultaneous Tests for General Linear Hypotheses
##
## Multiple Comparisons of Means: Tukey Contrasts
##
##
## Fit: glm(formula = Protein_Count ~ Tissue, family = gaussian, data = gene_
data)
##
## Linear Hypotheses:
##              Estimate Std. Error z value Pr(>|z|)
## ML - MF == 0  -111218     113925  -0.976    0.329
## (Adjusted p values reported -- single-step method)
##
##
## Tukey results for gene: KNG2
##
##   Simultaneous Tests for General Linear Hypotheses
##
## Multiple Comparisons of Means: Tukey Contrasts
##
##
## Fit: glm(formula = Protein_Count ~ Tissue, family = gaussian, data = gene_
data)
##
## Linear Hypotheses:
##              Estimate Std. Error z value Pr(>|z|)
## ML - MF == 0  -116083     142083  -0.817    0.414
## (Adjusted p values reported -- single-step method)
##
##
## Tukey results for gene: LOX
##
##   Simultaneous Tests for General Linear Hypotheses
##
## Multiple Comparisons of Means: Tukey Contrasts
##
##
## Fit: glm(formula = Protein_Count ~ Tissue, family = gaussian, data = gene_
data)
##
## Linear Hypotheses:
##              Estimate Std. Error z value Pr(>|z|)
## ML - MF == 0   351520    6170448   0.057    0.955
## (Adjusted p values reported -- single-step method)
##
##
## Tukey results for gene: LOXL1
```

```
##
##   Simultaneous Tests for General Linear Hypotheses
##
## Multiple Comparisons of Means: Tukey Contrasts
##
##
## Fit: glm(formula = Protein_Count ~ Tissue, family = gaussian, data = gene_
data)
##
## Linear Hypotheses:
##               Estimate Std. Error z value Pr(>|z|)
## ML - MF == 0  -111756    1871531   -0.06    0.952
## (Adjusted p values reported -- single-step method)
##
##
## Tukey results for gene: LOXL3
##
##   Simultaneous Tests for General Linear Hypotheses
##
## Multiple Comparisons of Means: Tukey Contrasts
##
##
## Fit: glm(formula = Protein_Count ~ Tissue, family = gaussian, data = gene_
data)
##
## Linear Hypotheses:
##               Estimate Std. Error z value Pr(>|z|)
## ML - MF == 0    10144      86033   0.118    0.906
## (Adjusted p values reported -- single-step method)
##
##
## Tukey results for gene: MASP2
##
##   Simultaneous Tests for General Linear Hypotheses
##
## Multiple Comparisons of Means: Tukey Contrasts
##
##
## Fit: glm(formula = Protein_Count ~ Tissue, family = gaussian, data = gene_
data)
##
## Linear Hypotheses:
##               Estimate Std. Error z value Pr(>|z|)
## ML - MF == 0 -1198952    1212266  -0.989    0.323
## (Adjusted p values reported -- single-step method)
##
##
## Tukey results for gene: MMP14
##
##   Simultaneous Tests for General Linear Hypotheses
```

```
##
## Multiple Comparisons of Means: Tukey Contrasts
##
##
## Fit: glm(formula = Protein_Count ~ Tissue, family = gaussian, data = gene_
data)
##
## Linear Hypotheses:
##              Estimate Std. Error z value Pr(>|z|)
## ML - MF == 0   208350     121006   1.722   0.0851 .
## ---
## Signif. codes:  0 '***' 0.001 '**' 0.01 '*' 0.05 '.' 0.1 ' ' 1
## (Adjusted p values reported -- single-step method)
##
##
## Tukey results for gene: MMP2
##
##   Simultaneous Tests for General Linear Hypotheses
##
## Multiple Comparisons of Means: Tukey Contrasts
##
##
## Fit: glm(formula = Protein_Count ~ Tissue, family = gaussian, data = gene_
data)
##
## Linear Hypotheses:
##              Estimate Std. Error z value Pr(>|z|)
## ML - MF == 0    20148     332955   0.061    0.952
## (Adjusted p values reported -- single-step method)
##
##
## Tukey results for gene: MMP23
##
##   Simultaneous Tests for General Linear Hypotheses
##
## Multiple Comparisons of Means: Tukey Contrasts
##
##
## Fit: glm(formula = Protein_Count ~ Tissue, family = gaussian, data = gene_
data)
##
## Linear Hypotheses:
##              Estimate Std. Error z value Pr(>|z|)
## ML - MF == 0    70127     142508   0.492    0.623
## (Adjusted p values reported -- single-step method)
##
##
## Tukey results for gene: MMP3
##
##   Simultaneous Tests for General Linear Hypotheses
```

```
##
## Multiple Comparisons of Means: Tukey Contrasts
##
##
## Fit: glm(formula = Protein_Count ~ Tissue, family = gaussian, data = gene_
data)
##
## Linear Hypotheses:
##              Estimate Std. Error z value Pr(>|z|)
## ML - MF == 0  -112664      71559  -1.574    0.115
## (Adjusted p values reported -- single-step method)
##
##
## Tukey results for gene: P4HA1
##
##   Simultaneous Tests for General Linear Hypotheses
##
## Multiple Comparisons of Means: Tukey Contrasts
##
##
## Fit: glm(formula = Protein_Count ~ Tissue, family = gaussian, data = gene_
data)
##
## Linear Hypotheses:
##              Estimate Std. Error z value Pr(>|z|)
## ML - MF == 0    15475     147959   0.105    0.917
## (Adjusted p values reported -- single-step method)
##
##
## Tukey results for gene: P4HA2
##
##   Simultaneous Tests for General Linear Hypotheses
##
## Multiple Comparisons of Means: Tukey Contrasts
##
##
## Fit: glm(formula = Protein_Count ~ Tissue, family = gaussian, data = gene_
data)
##
## Linear Hypotheses:
##              Estimate Std. Error z value Pr(>|z|)
## ML - MF == 0   115044      89600   1.284    0.199
## (Adjusted p values reported -- single-step method)
##
##
## Tukey results for gene: PLG
##
##   Simultaneous Tests for General Linear Hypotheses
##
## Multiple Comparisons of Means: Tukey Contrasts
```

```
##
##
## Fit: glm(formula = Protein_Count ~ Tissue, family = gaussian, data = gene_
data)
##
## Linear Hypotheses:
##              Estimate Std. Error z value Pr(>|z|)
## ML - MF == 0   841928    1014071    0.83    0.406
## (Adjusted p values reported -- single-step method)
##
##
## Tukey results for gene: PLOD1
##
##   Simultaneous Tests for General Linear Hypotheses
##
## Multiple Comparisons of Means: Tukey Contrasts
##
##
## Fit: glm(formula = Protein_Count ~ Tissue, family = gaussian, data = gene_
data)
##
## Linear Hypotheses:
##              Estimate Std. Error z value Pr(>|z|)
## ML - MF == 0  -257072     273971  -0.938    0.348
## (Adjusted p values reported -- single-step method)
##
##
## Tukey results for gene: PLOD3
##
##   Simultaneous Tests for General Linear Hypotheses
##
## Multiple Comparisons of Means: Tukey Contrasts
##
##
## Fit: glm(formula = Protein_Count ~ Tissue, family = gaussian, data = gene_
data)
##
## Linear Hypotheses:
##              Estimate Std. Error z value Pr(>|z|)
## ML - MF == 0   -41110     107129  -0.384    0.701
## (Adjusted p values reported -- single-step method)
##
##
## Tukey results for gene: PRSS1
##
##   Simultaneous Tests for General Linear Hypotheses
##
## Multiple Comparisons of Means: Tukey Contrasts
##
##
```

```
## Fit: glm(formula = Protein_Count ~ Tissue, family = gaussian, data = gene_
data)
##
## Linear Hypotheses:
##              Estimate Std. Error z value Pr(>|z|)
## ML - MF == 0  -312192    1760473  -0.177    0.859
## (Adjusted p values reported -- single-step method)
##
##
## Tukey results for gene: SERPINA1
##
##   Simultaneous Tests for General Linear Hypotheses
##
## Multiple Comparisons of Means: Tukey Contrasts
##
##
## Fit: glm(formula = Protein_Count ~ Tissue, family = gaussian, data = gene_
data)
##
## Linear Hypotheses:
##              Estimate Std. Error z value Pr(>|z|)
## ML - MF == 0  -109574      82344  -1.331    0.183
## (Adjusted p values reported -- single-step method)
##
##
## Tukey results for gene: SERPINA10
##
##   Simultaneous Tests for General Linear Hypotheses
##
## Multiple Comparisons of Means: Tukey Contrasts
##
##
## Fit: glm(formula = Protein_Count ~ Tissue, family = gaussian, data = gene_
data)
##
## Linear Hypotheses:
##              Estimate Std. Error z value Pr(>|z|)
## ML - MF == 0    -3735      57922  -0.064    0.949
## (Adjusted p values reported -- single-step method)
##
##
## Tukey results for gene: SERPINA3-8
##
##   Simultaneous Tests for General Linear Hypotheses
##
## Multiple Comparisons of Means: Tukey Contrasts
##
##
## Fit: glm(formula = Protein_Count ~ Tissue, family = gaussian, data = gene_
data)
```

```
##
## Linear Hypotheses:
##               Estimate Std. Error z value Pr(>|z|)
## ML - MF == 0   -33886      51544  -0.657    0.511
## (Adjusted p values reported -- single-step method)
##
##
## Tukey results for gene: SERPINA5
##
##   Simultaneous Tests for General Linear Hypotheses
##
## Multiple Comparisons of Means: Tukey Contrasts
##
##
## Fit: glm(formula = Protein_Count ~ Tissue, family = gaussian, data = gene_
data)
##
## Linear Hypotheses:
##               Estimate Std. Error z value Pr(>|z|)
## ML - MF == 0    71047     117524   0.605    0.545
## (Adjusted p values reported -- single-step method)
##
##
## Tukey results for gene: SERPINA7
##
##   Simultaneous Tests for General Linear Hypotheses
##
## Multiple Comparisons of Means: Tukey Contrasts
##
##
## Fit: glm(formula = Protein_Count ~ Tissue, family = gaussian, data = gene_
data)
##
## Linear Hypotheses:
##               Estimate Std. Error z value Pr(>|z|)
## ML - MF == 0    16222      28515   0.569    0.569
## (Adjusted p values reported -- single-step method)
##
##
## Tukey results for gene: SERPINB1
##
##   Simultaneous Tests for General Linear Hypotheses
##
## Multiple Comparisons of Means: Tukey Contrasts
##
##
## Fit: glm(formula = Protein_Count ~ Tissue, family = gaussian, data = gene_
data)
##
## Linear Hypotheses:
```

```
##               Estimate Std. Error z value Pr(>|z|)
## ML - MF == 0    27152      19328   1.405     0.16
## (Adjusted p values reported -- single-step method)
##
##
## Tukey results for gene: SERPINB6
##
##   Simultaneous Tests for General Linear Hypotheses
##
## Multiple Comparisons of Means: Tukey Contrasts
##
##
## Fit: glm(formula = Protein_Count ~ Tissue, family = gaussian, data = gene_
data)
##
## Linear Hypotheses:
##               Estimate Std. Error z value Pr(>|z|)
## ML - MF == 0    29078     188316   0.154    0.877
## (Adjusted p values reported -- single-step method)
##
##
## Tukey results for gene: SERPINB9
##
##   Simultaneous Tests for General Linear Hypotheses
##
## Multiple Comparisons of Means: Tukey Contrasts
##
##
## Fit: glm(formula = Protein_Count ~ Tissue, family = gaussian, data = gene_
data)
##
## Linear Hypotheses:
##               Estimate Std. Error z value Pr(>|z|)
## ML - MF == 0    62921      78162   0.805    0.421
## (Adjusted p values reported -- single-step method)
##
##
## Tukey results for gene: SERPINC1
##
##   Simultaneous Tests for General Linear Hypotheses
##
## Multiple Comparisons of Means: Tukey Contrasts
##
##
## Fit: glm(formula = Protein_Count ~ Tissue, family = gaussian, data = gene_
data)
##
## Linear Hypotheses:
##               Estimate Std. Error z value Pr(>|z|)
## ML - MF == 0 -1184172     711228  -1.665   0.0959 .
```

```
## ---
## Signif. codes:  0 '***' 0.001 '**' 0.01 '*' 0.05 '.' 0.1 ' ' 1
## (Adjusted p values reported -- single-step method)
##
##
## Tukey results for gene: SERPIND1
##
##   Simultaneous Tests for General Linear Hypotheses
##
## Multiple Comparisons of Means: Tukey Contrasts
##
##
## Fit: glm(formula = Protein_Count ~ Tissue, family = gaussian, data = gene_
data)
##
## Linear Hypotheses:
##              Estimate Std. Error z value Pr(>|z|)
## ML - MF == 0   227318     316799   0.718    0.473
## (Adjusted p values reported -- single-step method)
##
##
## Tukey results for gene: SERPINE2
##
##   Simultaneous Tests for General Linear Hypotheses
##
## Multiple Comparisons of Means: Tukey Contrasts
##
##
## Fit: glm(formula = Protein_Count ~ Tissue, family = gaussian, data = gene_
data)
##
## Linear Hypotheses:
##              Estimate Std. Error z value Pr(>|z|)
## ML - MF == 0   -71677     134902  -0.531    0.595
## (Adjusted p values reported -- single-step method)
##
##
## Tukey results for gene: SERPINF1
##
##   Simultaneous Tests for General Linear Hypotheses
##
## Multiple Comparisons of Means: Tukey Contrasts
##
##
## Fit: glm(formula = Protein_Count ~ Tissue, family = gaussian, data = gene_
data)
##
## Linear Hypotheses:
##              Estimate Std. Error z value Pr(>|z|)
## ML - MF == 0   749728    1352328   0.554    0.579
```

```
## (Adjusted p values reported -- single-step method)
##
##
## Tukey results for gene: SERPINF2
##
##   Simultaneous Tests for General Linear Hypotheses
##
## Multiple Comparisons of Means: Tukey Contrasts
##
##
## Fit: glm(formula = Protein_Count ~ Tissue, family = gaussian, data = gene_
data)
##
## Linear Hypotheses:
##              Estimate Std. Error z value Pr(>|z|)
## ML - MF == 0  -302241     137507  -2.198   0.0279 *
## ---
## Signif. codes:  0 '***' 0.001 '**' 0.01 '*' 0.05 '.' 0.1 ' ' 1
## (Adjusted p values reported -- single-step method)
##
##
## Tukey results for gene: SERPING1
##
##   Simultaneous Tests for General Linear Hypotheses
##
## Multiple Comparisons of Means: Tukey Contrasts
##
##
## Fit: glm(formula = Protein_Count ~ Tissue, family = gaussian, data = gene_
data)
##
## Linear Hypotheses:
##              Estimate Std. Error z value Pr(>|z|)
## ML - MF == 0    43628     413011   0.106    0.916
## (Adjusted p values reported -- single-step method)
##
##
## Tukey results for gene: SERPINH1
##
##   Simultaneous Tests for General Linear Hypotheses
##
## Multiple Comparisons of Means: Tukey Contrasts
##
##
## Fit: glm(formula = Protein_Count ~ Tissue, family = gaussian, data = gene_
data)
##
## Linear Hypotheses:
##              Estimate Std. Error z value Pr(>|z|)
## ML - MF == 0   990220    2459745   0.403    0.687
```

```
## (Adjusted p values reported -- single-step method)
##
##
## Tukey results for gene: SERPINI2
##
##   Simultaneous Tests for General Linear Hypotheses
##
## Multiple Comparisons of Means: Tukey Contrasts
##
##
## Fit: glm(formula = Protein_Count ~ Tissue, family = gaussian, data = gene_
data)
##
## Linear Hypotheses:
##              Estimate Std. Error z value Pr(>|z|)
## ML - MF == 0   154328     193412   0.798    0.425
## (Adjusted p values reported -- single-step method)
##
##
## Tukey results for gene: SULF2
##
##   Simultaneous Tests for General Linear Hypotheses
##
## Multiple Comparisons of Means: Tukey Contrasts
##
##
## Fit: glm(formula = Protein_Count ~ Tissue, family = gaussian, data = gene_
data)
##
## Linear Hypotheses:
##              Estimate Std. Error z value Pr(>|z|)
## ML - MF == 0     1780      82099   0.022    0.983
## (Adjusted p values reported -- single-step method)
##
##
## Tukey results for gene: TGM1
##
##   Simultaneous Tests for General Linear Hypotheses
##
## Multiple Comparisons of Means: Tukey Contrasts
##
##
## Fit: glm(formula = Protein_Count ~ Tissue, family = gaussian, data = gene_
data)
##
## Linear Hypotheses:
##              Estimate Std. Error z value Pr(>|z|)
## ML - MF == 0    76295     168451   0.453    0.651
## (Adjusted p values reported -- single-step method)
##
```

```
##
## Tukey results for gene: TGM2
##
##   Simultaneous Tests for General Linear Hypotheses
##
## Multiple Comparisons of Means: Tukey Contrasts
##
##
## Fit: glm(formula = Protein_Count ~ Tissue, family = gaussian, data = gene_
data)
##
## Linear Hypotheses:
##                 Estimate Std. Error z value Pr(>|z|)
## ML - MF == 0 -49352820   47388159  -1.041    0.298
## (Adjusted p values reported -- single-step method)
##
##
## Tukey results for gene: TIMP2
##
##   Simultaneous Tests for General Linear Hypotheses
##
## Multiple Comparisons of Means: Tukey Contrasts
##
##
## Fit: glm(formula = Protein_Count ~ Tissue, family = gaussian, data = gene_
data)
##
## Linear Hypotheses:
##                Estimate Std. Error z value Pr(>|z|)
## ML - MF == 0    54235     113856   0.476    0.634
## (Adjusted p values reported -- single-step method)

#### ECM-affiliated

# Read your cleaned joined data file into a data frame
cleaned_joined_data <- read.csv("cleaned_matrisome_medu.csv")

# Filter rows with Matrisome_Category "ECM-affiliated"
ECMaffiliated_data <- cleaned_joined_data %>%
  filter(Matrisome_Category == "ECM-affiliated")

# Reshape the data to have separate columns for each replicate and each tissu
e stage
organized_data <- ECMaffiliated_data %>%
  pivot_longer(cols = starts_with("MF") | starts_with("ML"),
               names_to = c("Tissue", "Stage"),
               names_sep = "\\.",
               values_to = "Protein_Count")
```

```
## Warning: Expected 2 pieces. Missing pieces filled with `NA` in 2 rows [1,
6].

# Save the reshaped data to a new file

write.csv(organized_data, "ECM-affiliated_data_matrisome_medu_to_clean.csv",
row.names = FALSE)

data2 <- read.csv("ECM-affiliated_data_matrisome_medu_clean.csv")

png(filename="medu_ECM-affiliated.png",
    width = 12, height = 6, units = "in",
    res = 600)

boxplot(Protein_Count ~ Tissue*Genes,
        data = data2,
        cex.axis = 0.8,
        col = viridis(2, alpha = 0.7),
        lwd = 0.7,
        ylab = "",
        ylim = c(0, 20000000),
        axes = FALSE,
        frame.plot = TRUE,
        xlab = "")

GROUP <- unique(data2$Genes)
groups <- 23
numbox <- 2
total <- groups * numbox
xpoints <- seq(median(1:numbox), total, numbox)

axis(2, seq(0, 20000000, by = 1000000), las = 1, cex.axis = 0.7)
axis(1, labels = GROUP, at = xpoints, las = 2, cex.axis = 0.7)

stripchart(data2$Protein_Count ~ data2$Tissue*data2$Genes,
           method = "jitter",
           pch = 19,
           col = "black",
           vertical = TRUE,
           add = TRUE,
           cex = 0.3)

title(ylab = expression("Protein abundance (RU)"), line = 3.0)
legend("top", fill = viridis(2), legend = c("Medulla Follicular", "Medulla Lu
teal"), horiz = T, cex = 0.7)

dev.off()
```

```
## quartz_off_screen
##                  2

# Convert Tissue to factor
data2$Tissue <- as.factor(data2$Tissue)

# Create an empty list to store Tukey results
tukey_results_list <- list()

# Unique genes
unique_genes <- unique(data2$Genes)

# Loop over each gene
for (gene in unique_genes) {
  # Subset data for the current gene
  gene_data <- subset(data2, Genes == gene)

  # Fit the generalized linear model
  glm_model <- glm(Protein_Count ~ Tissue, data = gene_data, family = gaussia
n)

  # Perform Tukey test
  contrast_matrix <- glht(glm_model, linfct = mcp(Tissue = "Tukey"))
  tukey_results <- summary(contrast_matrix)

  # Store the results
  tukey_results_list[[gene]] <- tukey_results
}

# Print the results for each gene
for (gene in unique_genes) {
  cat("Tukey results for gene:", gene, "\n")
  print(tukey_results_list[[gene]])
  cat("\n")
}

## Tukey results for gene: ANXA1
##
##   Simultaneous Tests for General Linear Hypotheses
##
## Multiple Comparisons of Means: Tukey Contrasts
##
##
## Fit: glm(formula = Protein_Count ~ Tissue, family = gaussian, data = gene_
data)
##
## Linear Hypotheses:
##              Estimate Std. Error z value Pr(>|z|)
## ML - MF == 0   694386     541875   1.281      0.2
## (Adjusted p values reported -- single-step method)
```

```
##
##
## Tukey results for gene: ANXA11
##
##   Simultaneous Tests for General Linear Hypotheses
##
## Multiple Comparisons of Means: Tukey Contrasts
##
##
## Fit: glm(formula = Protein_Count ~ Tissue, family = gaussian, data = gene_
data)
##
## Linear Hypotheses:
##              Estimate Std. Error z value Pr(>|z|)
## ML - MF == 0   108313     249758   0.434    0.665
## (Adjusted p values reported -- single-step method)
##
##
## Tukey results for gene: ANXA2
##
##   Simultaneous Tests for General Linear Hypotheses
##
## Multiple Comparisons of Means: Tukey Contrasts
##
##
## Fit: glm(formula = Protein_Count ~ Tissue, family = gaussian, data = gene_
data)
##
## Linear Hypotheses:
##              Estimate Std. Error z value Pr(>|z|)
## ML - MF == 0  3478362    2572603   1.352    0.176
## (Adjusted p values reported -- single-step method)
##
##
## Tukey results for gene: ANXA3
##
##   Simultaneous Tests for General Linear Hypotheses
##
## Multiple Comparisons of Means: Tukey Contrasts
##
##
## Fit: glm(formula = Protein_Count ~ Tissue, family = gaussian, data = gene_
data)
##
## Linear Hypotheses:
##              Estimate Std. Error z value Pr(>|z|)
## ML - MF == 0   125063     232412   0.538    0.591
## (Adjusted p values reported -- single-step method)
##
##
```

```
## Tukey results for gene: ANXA4
##
##   Simultaneous Tests for General Linear Hypotheses
##
## Multiple Comparisons of Means: Tukey Contrasts
##
##
## Fit: glm(formula = Protein_Count ~ Tissue, family = gaussian, data = gene_
data)
##
## Linear Hypotheses:
##              Estimate Std. Error z value Pr(>|z|)
## ML - MF == 0  -130079     122290  -1.064    0.287
## (Adjusted p values reported -- single-step method)
##
##
## Tukey results for gene: ANXA5
##
##   Simultaneous Tests for General Linear Hypotheses
##
## Multiple Comparisons of Means: Tukey Contrasts
##
##
## Fit: glm(formula = Protein_Count ~ Tissue, family = gaussian, data = gene_
data)
##
## Linear Hypotheses:
##              Estimate Std. Error z value Pr(>|z|)
## ML - MF == 0  -508527    3261214  -0.156    0.876
## (Adjusted p values reported -- single-step method)
##
##
## Tukey results for gene: ANXA6
##
##   Simultaneous Tests for General Linear Hypotheses
##
## Multiple Comparisons of Means: Tukey Contrasts
##
##
## Fit: glm(formula = Protein_Count ~ Tissue, family = gaussian, data = gene_
data)
##
## Linear Hypotheses:
##              Estimate Std. Error z value Pr(>|z|)
## ML - MF == 0    58064     472734   0.123    0.902
## (Adjusted p values reported -- single-step method)
##
##
## Tukey results for gene: ANXA7
##
```

```
##   Simultaneous Tests for General Linear Hypotheses
##
## Multiple Comparisons of Means: Tukey Contrasts
##
##
## Fit: glm(formula = Protein_Count ~ Tissue, family = gaussian, data = gene_
data)
##
## Linear Hypotheses:
##              Estimate Std. Error z value Pr(>|z|)
## ML - MF == 0   174194     239205   0.728    0.466
## (Adjusted p values reported -- single-step method)
##
##
## Tukey results for gene: C1QA
##
##   Simultaneous Tests for General Linear Hypotheses
##
## Multiple Comparisons of Means: Tukey Contrasts
##
##
## Fit: glm(formula = Protein_Count ~ Tissue, family = gaussian, data = gene_
data)
##
## Linear Hypotheses:
##              Estimate Std. Error z value Pr(>|z|)
## ML - MF == 0     7350      10062    0.73    0.465
## (Adjusted p values reported -- single-step method)
##
##
## Tukey results for gene: C1QC
##
##   Simultaneous Tests for General Linear Hypotheses
##
## Multiple Comparisons of Means: Tukey Contrasts
##
##
## Fit: glm(formula = Protein_Count ~ Tissue, family = gaussian, data = gene_
data)
##
## Linear Hypotheses:
##              Estimate Std. Error z value Pr(>|z|)
## ML - MF == 0    47714     137392   0.347    0.728
## (Adjusted p values reported -- single-step method)
##
##
## Tukey results for gene: C1QTNF3
##
##   Simultaneous Tests for General Linear Hypotheses
##
```

```
## Multiple Comparisons of Means: Tukey Contrasts
##
##
## Fit: glm(formula = Protein_Count ~ Tissue, family = gaussian, data = gene_
data)
##
## Linear Hypotheses:
##              Estimate Std. Error z value Pr(>|z|)
## ML - MF == 0     8962      35330   0.254      0.8
## (Adjusted p values reported -- single-step method)
##
##
## Tukey results for gene: C1QTNF5
##
##   Simultaneous Tests for General Linear Hypotheses
##
## Multiple Comparisons of Means: Tukey Contrasts
##
##
## Fit: glm(formula = Protein_Count ~ Tissue, family = gaussian, data = gene_
data)
##
## Linear Hypotheses:
##              Estimate Std. Error z value Pr(>|z|)
## ML - MF == 0   -30911     254369  -0.122    0.903
## (Adjusted p values reported -- single-step method)
##
##
## Tukey results for gene: C1QTNF7
##
##   Simultaneous Tests for General Linear Hypotheses
##
## Multiple Comparisons of Means: Tukey Contrasts
##
##
## Fit: glm(formula = Protein_Count ~ Tissue, family = gaussian, data = gene_
data)
##
## Linear Hypotheses:
##              Estimate Std. Error z value Pr(>|z|)
## ML - MF == 0    57995     151059   0.384    0.701
## (Adjusted p values reported -- single-step method)
##
##
## Tukey results for gene: CLEC11A
##
##   Simultaneous Tests for General Linear Hypotheses
##
## Multiple Comparisons of Means: Tukey Contrasts
##
```

```
##
## Fit: glm(formula = Protein_Count ~ Tissue, family = gaussian, data = gene_
data)
##
## Linear Hypotheses:
##               Estimate Std. Error z value Pr(>|z|)
## ML - MF == 0  -234555     231632  -1.013    0.311
## (Adjusted p values reported -- single-step method)
##
##
## Tukey results for gene: CLEC3B
##
##   Simultaneous Tests for General Linear Hypotheses
##
## Multiple Comparisons of Means: Tukey Contrasts
##
##
## Fit: glm(formula = Protein_Count ~ Tissue, family = gaussian, data = gene_
data)
##
## Linear Hypotheses:
##               Estimate Std. Error z value Pr(>|z|)
## ML - MF == 0  1048742    1358188   0.772     0.44
## (Adjusted p values reported -- single-step method)
##
##
## Tukey results for gene: COLEC12
##
##   Simultaneous Tests for General Linear Hypotheses
##
## Multiple Comparisons of Means: Tukey Contrasts
##
##
## Fit: glm(formula = Protein_Count ~ Tissue, family = gaussian, data = gene_
data)
##
## Linear Hypotheses:
##               Estimate Std. Error z value Pr(>|z|)
## ML - MF == 0   -71344     120060  -0.594    0.552
## (Adjusted p values reported -- single-step method)
##
##
## Tukey results for gene: CSPG4
##
##   Simultaneous Tests for General Linear Hypotheses
##
## Multiple Comparisons of Means: Tukey Contrasts
##
##
## Fit: glm(formula = Protein_Count ~ Tissue, family = gaussian, data = gene_
```

```
data)
##
## Linear Hypotheses:
##              Estimate Std. Error z value Pr(>|z|)
## ML - MF == 0    60185     110908   0.543    0.587
## (Adjusted p values reported -- single-step method)
##
##
## Tukey results for gene: HPX
##
##   Simultaneous Tests for General Linear Hypotheses
##
## Multiple Comparisons of Means: Tukey Contrasts
##
##
## Fit: glm(formula = Protein_Count ~ Tissue, family = gaussian, data = gene_
data)
##
## Linear Hypotheses:
##              Estimate Std. Error z value Pr(>|z|)
## ML - MF == 0 -1275680     818488  -1.559    0.119
## (Adjusted p values reported -- single-step method)
##
##
## Tukey results for gene: LGALS1
##
##   Simultaneous Tests for General Linear Hypotheses
##
## Multiple Comparisons of Means: Tukey Contrasts
##
##
## Fit: glm(formula = Protein_Count ~ Tissue, family = gaussian, data = gene_
data)
##
## Linear Hypotheses:
##              Estimate Std. Error z value Pr(>|z|)
## ML - MF == 0  -549206     936670  -0.586    0.558
## (Adjusted p values reported -- single-step method)
##
##
## Tukey results for gene: LGALS9
##
##   Simultaneous Tests for General Linear Hypotheses
##
## Multiple Comparisons of Means: Tukey Contrasts
##
##
## Fit: glm(formula = Protein_Count ~ Tissue, family = gaussian, data = gene_
data)
##
```

```
## Linear Hypotheses:
##              Estimate Std. Error z value Pr(>|z|)
## ML - MF == 0    26532     240860    0.11    0.912
## (Adjusted p values reported -- single-step method)
##
##
## Tukey results for gene: LMAN1
##
##   Simultaneous Tests for General Linear Hypotheses
##
## Multiple Comparisons of Means: Tukey Contrasts
##
##
## Fit: glm(formula = Protein_Count ~ Tissue, family = gaussian, data = gene_
data)
##
## Linear Hypotheses:
##              Estimate Std. Error z value Pr(>|z|)
## ML - MF == 0    35714      57982   0.616    0.538
## (Adjusted p values reported -- single-step method)
##
##
## Tukey results for gene: PLXNB1
##
##   Simultaneous Tests for General Linear Hypotheses
##
## Multiple Comparisons of Means: Tukey Contrasts
##
##
## Fit: glm(formula = Protein_Count ~ Tissue, family = gaussian, data = gene_
data)
##
## Linear Hypotheses:
##              Estimate Std. Error z value Pr(>|z|)
## ML - MF == 0  -116028      49353  -2.351   0.0187 *
## ---
## Signif. codes:  0 '***' 0.001 '**' 0.01 '*' 0.05 '.' 0.1 ' ' 1
## (Adjusted p values reported -- single-step method)
##
##
## Tukey results for gene: PLXNB2
##
##   Simultaneous Tests for General Linear Hypotheses
##
## Multiple Comparisons of Means: Tukey Contrasts
##
##
## Fit: glm(formula = Protein_Count ~ Tissue, family = gaussian, data = gene_
data)
##
```

```
## Linear Hypotheses:
##                Estimate Std. Error z value Pr(>|z|)
## ML - MF == 0  -270166     175813  -1.537    0.124
## (Adjusted p values reported -- single-step method)
```

```r
#### Secreted Factors

# Read your cleaned joined data file into a data frame
cleaned_joined_data <- read.csv("cleaned_matrisome_medu.csv")

# Filter rows with Matrisome_Category "Secreted Factors"
Secreted_Factors_data <- cleaned_joined_data %>%
  filter(Matrisome_Category == "Secreted Factors")

# Reshape the data to have separate columns for each replicate and each tissu
e stage
organized_data <- Secreted_Factors_data %>%
  pivot_longer(cols = starts_with("MF") | starts_with("ML"),
               names_to = c("Tissue", "Stage"),
               names_sep = "\\.",
               values_to = "Protein_Count")
```

```
## Warning: Expected 2 pieces. Missing pieces filled with `NA` in 2 rows [1,
6].
```

```r
# Save the reshaped data to a new file

write.csv(organized_data, "Secreted_factors_data_matrisome_medu_to_clean.csv"
, row.names = FALSE)

data2 <- read.csv("Secreted_factors_data_matrisome_medu_clean.csv")

png(filename="medu_Secreted_factors.png",
    width = 12, height = 6, units = "in",
    res = 600)

boxplot(Protein_Count ~ Tissue*Genes,
        data = data2,
        cex.axis = 0.8,
        col = viridis(2, alpha = 0.7),
        lwd = 0.7,
        ylab = "",
        ylim = c(0, 25000000),
        axes = FALSE,
        frame.plot = TRUE,
        xlab = "")

GROUP <- unique(data2$Genes)
groups <- 27
numbox <- 2
```

```r
total <- groups * numbox
xpoints <- seq(median(1:numbox), total, numbox)

axis(2, seq(0, 25000000, by = 5000000), las = 1, cex.axis = 0.7)
axis(1, labels = GROUP, at = xpoints, las = 2, cex.axis = 0.7)

stripchart(data2$Protein_Count ~ data2$Tissue*data2$Genes,
           method = "jitter",
           pch = 19,
           col = "black",
           vertical = TRUE,
           add = TRUE,
           cex = 0.3)

title(ylab = expression("Protein abundance (RU)"), line = 3.0)
legend("top", fill = viridis(2), legend = c("Medulla Follicular", "Medulla Lu
teal"), horiz = T, cex = 0.7)

dev.off()

## quartz_off_screen
##                 2

# Convert Tissue to factor
data2$Tissue <- as.factor(data2$Tissue)

# Create an empty list to store Tukey results
tukey_results_list <- list()

# Unique genes
unique_genes <- unique(data2$Genes)

# Loop over each gene
for (gene in unique_genes) {
  # Subset data for the current gene
  gene_data <- subset(data2, Genes == gene)

  # Fit the generalized linear model
  glm_model <- glm(Protein_Count ~ Tissue, data = gene_data, family = gaussia
n)

  # Perform Tukey test
  contrast_matrix <- glht(glm_model, linfct = mcp(Tissue = "Tukey"))
  tukey_results <- summary(contrast_matrix)

  # Store the results
  tukey_results_list[[gene]] <- tukey_results
}
```

```
# Print the results for each gene
for (gene in unique_genes) {
  cat("Tukey results for gene:", gene, "\n")
  print(tukey_results_list[[gene]])
  cat("\n")
}
```

```
## Tukey results for gene: ANGPT4
##
##   Simultaneous Tests for General Linear Hypotheses
##
## Multiple Comparisons of Means: Tukey Contrasts
##
##
## Fit: glm(formula = Protein_Count ~ Tissue, family = gaussian, data = gene_
data)
##
## Linear Hypotheses:
##                Estimate Std. Error z value Pr(>|z|)
## ML - MF == 0  -224547      200140  -1.122    0.262
## (Adjusted p values reported -- single-step method)
##
##
## Tukey results for gene: ANGPTL2
##
##   Simultaneous Tests for General Linear Hypotheses
##
## Multiple Comparisons of Means: Tukey Contrasts
##
##
## Fit: glm(formula = Protein_Count ~ Tissue, family = gaussian, data = gene_
data)
##
## Linear Hypotheses:
##                Estimate Std. Error z value Pr(>|z|)
## ML - MF == 0    36004      107681   0.334    0.738
## (Adjusted p values reported -- single-step method)
##
##
## Tukey results for gene: ANGPTL6
##
##   Simultaneous Tests for General Linear Hypotheses
##
## Multiple Comparisons of Means: Tukey Contrasts
##
##
## Fit: glm(formula = Protein_Count ~ Tissue, family = gaussian, data = gene_
data)
##
## Linear Hypotheses:
```

```
##               Estimate Std. Error z value Pr(>|z|)
## ML - MF == 0    35379      67882   0.521    0.602
## (Adjusted p values reported -- single-step method)
##
##
## Tukey results for gene: CHRD
##
##   Simultaneous Tests for General Linear Hypotheses
##
## Multiple Comparisons of Means: Tukey Contrasts
##
##
## Fit: glm(formula = Protein_Count ~ Tissue, family = gaussian, data = gene_
data)
##
## Linear Hypotheses:
##               Estimate Std. Error z value Pr(>|z|)
## ML - MF == 0   -22412      93444   -0.24     0.81
## (Adjusted p values reported -- single-step method)
##
##
## Tukey results for gene: CXCL12
##
##   Simultaneous Tests for General Linear Hypotheses
##
## Multiple Comparisons of Means: Tukey Contrasts
##
##
## Fit: glm(formula = Protein_Count ~ Tissue, family = gaussian, data = gene_
data)
##
## Linear Hypotheses:
##               Estimate Std. Error z value Pr(>|z|)
## ML - MF == 0    16974      82293   0.206    0.837
## (Adjusted p values reported -- single-step method)
##
##
## Tukey results for gene: FRZB
##
##   Simultaneous Tests for General Linear Hypotheses
##
## Multiple Comparisons of Means: Tukey Contrasts
##
##
## Fit: glm(formula = Protein_Count ~ Tissue, family = gaussian, data = gene_
data)
##
## Linear Hypotheses:
##               Estimate Std. Error z value Pr(>|z|)
## ML - MF == 0   117160     122779   0.954     0.34
```

```
## (Adjusted p values reported -- single-step method)
##
##
## Tukey results for gene: FST
##
##   Simultaneous Tests for General Linear Hypotheses
##
## Multiple Comparisons of Means: Tukey Contrasts
##
##
## Fit: glm(formula = Protein_Count ~ Tissue, family = gaussian, data = gene_
data)
##
## Linear Hypotheses:
##              Estimate Std. Error z value Pr(>|z|)
## ML - MF == 0    12686     146004   0.087    0.931
## (Adjusted p values reported -- single-step method)
##
##
## Tukey results for gene: HCFC1
##
##   Simultaneous Tests for General Linear Hypotheses
##
## Multiple Comparisons of Means: Tukey Contrasts
##
##
## Fit: glm(formula = Protein_Count ~ Tissue, family = gaussian, data = gene_
data)
##
## Linear Hypotheses:
##              Estimate Std. Error z value Pr(>|z|)
## ML - MF == 0    50012     152020   0.329    0.742
## (Adjusted p values reported -- single-step method)
##
##
## Tukey results for gene: HGFAC
##
##   Simultaneous Tests for General Linear Hypotheses
##
## Multiple Comparisons of Means: Tukey Contrasts
##
##
## Fit: glm(formula = Protein_Count ~ Tissue, family = gaussian, data = gene_
data)
##
## Linear Hypotheses:
##              Estimate Std. Error z value Pr(>|z|)
## ML - MF == 0    22401     118499   0.189     0.85
## (Adjusted p values reported -- single-step method)
##
```

```
##
## Tukey results for gene: IGF2
##
##   Simultaneous Tests for General Linear Hypotheses
##
## Multiple Comparisons of Means: Tukey Contrasts
##
##
## Fit: glm(formula = Protein_Count ~ Tissue, family = gaussian, data = gene_
data)
##
## Linear Hypotheses:
##              Estimate Std. Error z value Pr(>|z|)
## ML - MF == 0   -84935     107172  -0.793    0.428
## (Adjusted p values reported -- single-step method)
##
##
## Tukey results for gene: IL17D
##
##   Simultaneous Tests for General Linear Hypotheses
##
## Multiple Comparisons of Means: Tukey Contrasts
##
##
## Fit: glm(formula = Protein_Count ~ Tissue, family = gaussian, data = gene_
data)
##
## Linear Hypotheses:
##              Estimate Std. Error z value Pr(>|z|)
## ML - MF == 0   -83455     121135  -0.689    0.491
## (Adjusted p values reported -- single-step method)
##
##
## Tukey results for gene: INHBB
##
##   Simultaneous Tests for General Linear Hypotheses
##
## Multiple Comparisons of Means: Tukey Contrasts
##
##
## Fit: glm(formula = Protein_Count ~ Tissue, family = gaussian, data = gene_
data)
##
## Linear Hypotheses:
##              Estimate Std. Error z value Pr(>|z|)
## ML - MF == 0    45775      42426   1.079    0.281
## (Adjusted p values reported -- single-step method)
##
##
## Tukey results for gene: INSL3
```

```
##
##   Simultaneous Tests for General Linear Hypotheses
##
## Multiple Comparisons of Means: Tukey Contrasts
##
##
## Fit: glm(formula = Protein_Count ~ Tissue, family = gaussian, data = gene_
data)
##
## Linear Hypotheses:
##              Estimate Std. Error z value Pr(>|z|)
## ML - MF == 0  -111698      95155  -1.174     0.24
## (Adjusted p values reported -- single-step method)
##
##
## Tukey results for gene: MDK
##
##   Simultaneous Tests for General Linear Hypotheses
##
## Multiple Comparisons of Means: Tukey Contrasts
##
##
## Fit: glm(formula = Protein_Count ~ Tissue, family = gaussian, data = gene_
data)
##
## Linear Hypotheses:
##              Estimate Std. Error z value Pr(>|z|)
## ML - MF == 0   -86418     259888  -0.333    0.739
## (Adjusted p values reported -- single-step method)
##
##
## Tukey results for gene: MST1
##
##   Simultaneous Tests for General Linear Hypotheses
##
## Multiple Comparisons of Means: Tukey Contrasts
##
##
## Fit: glm(formula = Protein_Count ~ Tissue, family = gaussian, data = gene_
data)
##
## Linear Hypotheses:
##              Estimate Std. Error z value Pr(>|z|)
## ML - MF == 0    54935     131535   0.418    0.676
## (Adjusted p values reported -- single-step method)
##
##
## Tukey results for gene: PTN
##
##   Simultaneous Tests for General Linear Hypotheses
```

```
##
## Multiple Comparisons of Means: Tukey Contrasts
##
##
## Fit: glm(formula = Protein_Count ~ Tissue, family = gaussian, data = gene_
data)
##
## Linear Hypotheses:
##              Estimate Std. Error z value Pr(>|z|)
## ML - MF == 0  -365067     264218  -1.382    0.167
## (Adjusted p values reported -- single-step method)
##
##
## Tukey results for gene: S100A10
##
##   Simultaneous Tests for General Linear Hypotheses
##
## Multiple Comparisons of Means: Tukey Contrasts
##
##
## Fit: glm(formula = Protein_Count ~ Tissue, family = gaussian, data = gene_
data)
##
## Linear Hypotheses:
##              Estimate Std. Error z value Pr(>|z|)
## ML - MF == 0    72159     152343   0.474    0.636
## (Adjusted p values reported -- single-step method)
##
##
## Tukey results for gene: S100A11
##
##   Simultaneous Tests for General Linear Hypotheses
##
## Multiple Comparisons of Means: Tukey Contrasts
##
##
## Fit: glm(formula = Protein_Count ~ Tissue, family = gaussian, data = gene_
data)
##
## Linear Hypotheses:
##              Estimate Std. Error z value Pr(>|z|)
## ML - MF == 0   504310    1068411   0.472    0.637
## (Adjusted p values reported -- single-step method)
##
##
## Tukey results for gene: S100A12
##
##   Simultaneous Tests for General Linear Hypotheses
##
## Multiple Comparisons of Means: Tukey Contrasts
```

```
##
##
## Fit: glm(formula = Protein_Count ~ Tissue, family = gaussian, data = gene_
data)
##
## Linear Hypotheses:
##              Estimate Std. Error z value Pr(>|z|)
## ML - MF == 0    56256      44980   1.251    0.211
## (Adjusted p values reported -- single-step method)
##
##
## Tukey results for gene: S100A13
##
##   Simultaneous Tests for General Linear Hypotheses
##
## Multiple Comparisons of Means: Tukey Contrasts
##
##
## Fit: glm(formula = Protein_Count ~ Tissue, family = gaussian, data = gene_
data)
##
## Linear Hypotheses:
##              Estimate Std. Error z value Pr(>|z|)
## ML - MF == 0    36215     225925    0.16    0.873
## (Adjusted p values reported -- single-step method)
##
##
## Tukey results for gene: S100A14
##
##   Simultaneous Tests for General Linear Hypotheses
##
## Multiple Comparisons of Means: Tukey Contrasts
##
##
## Fit: glm(formula = Protein_Count ~ Tissue, family = gaussian, data = gene_
data)
##
## Linear Hypotheses:
##              Estimate Std. Error z value Pr(>|z|)
## ML - MF == 0     2114      79774   0.027    0.979
## (Adjusted p values reported -- single-step method)
##
##
## Tukey results for gene: S100A16
##
##   Simultaneous Tests for General Linear Hypotheses
##
## Multiple Comparisons of Means: Tukey Contrasts
##
##
```

```
## Fit: glm(formula = Protein_Count ~ Tissue, family = gaussian, data = gene_
data)
##
## Linear Hypotheses:
##              Estimate Std. Error z value Pr(>|z|)
## ML - MF == 0    55123      68853   0.801    0.423
## (Adjusted p values reported -- single-step method)
##
##
## Tukey results for gene: S100A4
##
##   Simultaneous Tests for General Linear Hypotheses
##
## Multiple Comparisons of Means: Tukey Contrasts
##
##
## Fit: glm(formula = Protein_Count ~ Tissue, family = gaussian, data = gene_
data)
##
## Linear Hypotheses:
##              Estimate Std. Error z value Pr(>|z|)
## ML - MF == 0  3423660    3123589   1.096    0.273
## (Adjusted p values reported -- single-step method)
##
##
## Tukey results for gene: TGFB1
##
##   Simultaneous Tests for General Linear Hypotheses
##
## Multiple Comparisons of Means: Tukey Contrasts
##
##
## Fit: glm(formula = Protein_Count ~ Tissue, family = gaussian, data = gene_
data)
##
## Linear Hypotheses:
##              Estimate Std. Error z value Pr(>|z|)
## ML - MF == 0   151988      93612   1.624    0.104
## (Adjusted p values reported -- single-step method)
##
##
## Tukey results for gene: TGFB2
##
##   Simultaneous Tests for General Linear Hypotheses
##
## Multiple Comparisons of Means: Tukey Contrasts
##
##
## Fit: glm(formula = Protein_Count ~ Tissue, family = gaussian, data = gene_
data)
```

```
##
## Linear Hypotheses:
##                Estimate Std. Error z value Pr(>|z|)
## ML - MF == 0    -80506     524242  -0.154    0.878
## (Adjusted p values reported -- single-step method)
##
##
## Tukey results for gene: TGFB3
##
##   Simultaneous Tests for General Linear Hypotheses
##
## Multiple Comparisons of Means: Tukey Contrasts
##
##
## Fit: glm(formula = Protein_Count ~ Tissue, family = gaussian, data = gene_
data)
##
## Linear Hypotheses:
##                Estimate Std. Error z value Pr(>|z|)
## ML - MF == 0     29307     482940   0.061    0.952
## (Adjusted p values reported -- single-step method)
##
##
## Tukey results for gene: WNT6
##
##   Simultaneous Tests for General Linear Hypotheses
##
## Multiple Comparisons of Means: Tukey Contrasts
##
##
## Fit: glm(formula = Protein_Count ~ Tissue, family = gaussian, data = gene_
data)
##
## Linear Hypotheses:
##                Estimate Std. Error z value Pr(>|z|)
## ML - MF == 0     32969     104531   0.315    0.752
## (Adjusted p values reported -- single-step method)

#### Components proportions

data <- read.csv("medulla_matrisome_final.csv")

# Subset the data for each tissue (MF and ML) separately
data_MF <- subset(data, Tissue == "MF")
data_ML <- subset(data, Tissue == "ML")

# Function to calculate proportions
calculate_proportions <- function(data_subset) {
  proportions <- data_subset %>%
    group_by(Matrisome_Category) %>%
```

```
    summarize(Proportion = sum(Protein_Count) / sum(data_subset$Protein_Count
) * 100)
  return(proportions)
}

# Calculate proportions for MF
proportions_MF <- calculate_proportions(data_MF)

# Calculate proportions for ML
proportions_ML <- calculate_proportions(data_ML)

# Print proportions for MF
cat("Proportions for Matrisome categories in tissue MF:\n")

## Proportions for Matrisome categories in tissue MF:

print(proportions_MF)

## # A tibble: 6 × 2
##   Matrisome_Category Proportion
##   <chr>                   <dbl>
## 1 Collagens              70.7
## 2 ECM Glycoproteins      15.3
## 3 ECM Regulators          5.35
## 4 ECM-affiliated          0.750
## 5 Proteoglycans           7.52
## 6 Secreted Factors        0.370

# Print proportions for ML
cat("Proportions for Matrisome categories in tissue ML:\n")

## Proportions for Matrisome categories in tissue ML:

print(proportions_ML)

## # A tibble: 6 × 2
##   Matrisome_Category Proportion
##   <chr>                   <dbl>
## 1 Collagens              68.9
## 2 ECM Glycoproteins      15.5
## 3 ECM Regulators          5.13
## 4 ECM-affiliated          0.865
## 5 Proteoglycans           9.10
## 6 Secreted Factors        0.478

###Pie charts

# Define a function to create a pie chart with values and save it
create_and_save_pie_chart <- function(proportions, title, filename) {
  # Create a color palette using Viridis
  color_palette <- viridis(length(unique(proportions$Matrisome_Category)))
```

```
  # Create a pie chart
  pie_chart <- ggplot(proportions, aes(x = "", y = Proportion, fill = Matriso
me_Category)) +
    geom_bar(stat = "identity") +
    coord_polar("y", start = 1) +
    geom_text(aes(label = sprintf("%.1f%%", Proportion)), position = position
_stack(vjust = 0.5), color = "white", size = 6) +
    labs(title = title, x = NULL, y = NULL, fill = "Matrisome Category") +
    scale_fill_manual(values = color_palette) +  # Set the color palette
    theme_void() +
    theme(legend.position = "right")

  # Save the pie chart
  ggsave(filename, plot = pie_chart, width = 6, height = 6, units = "in", dpi
= 300)
}

# Create and save a pie chart for MF
create_and_save_pie_chart(proportions_MF, "Matrisome Categories Proportion in
Tissue MF", "pie_chart_MF.png")

# Create and save a pie chart for ML
create_and_save_pie_chart(proportions_ML, "Matrisome Categories Proportion in
Tissue ML", "pie_chart_ML.png")

#####Summary data

data <- read.csv("medulla_matrisome_final.csv")

# Calculate mean and standard deviation for each Gene considering Tissue and
Matrisome_Category
gene_stats <- data %>%
  group_by(Genes, Tissue, Matrisome_Category) %>%
  summarize(
    Mean_Protein_Count = mean(Protein_Count),
    SD_Protein_Count = sd(Protein_Count)
  )

## `summarise()` has grouped output by 'Genes', 'Tissue'. You can override us
ing
## the `.groups` argument.

# Print the results
print(gene_stats)

## # A tibble: 500 × 5
## # Groups:   Genes, Tissue [500]
##    Genes    Tissue Matrisome_Category Mean_Protein_Count SD_Protein_Count
##    <chr>    <chr>  <chr>                           <dbl>            <dbl>
```

```
##   1 A2M      MF     ECM Regulators                4217704          1439391.
##   2 A2M      ML     ECM Regulators                3365618          1052989.
##   3 ABI3BP   MF     ECM Glycoproteins              331748.          295472.
##   4 ABI3BP   ML     ECM Glycoproteins              320254.          199576.
##   5 ACAN     MF     Proteoglycans                 1854652.          783951.
##   6 ACAN     ML     Proteoglycans                 1727749.         1367941.
##   7 ADAM10   MF     ECM Regulators                 164932           229723.
##   8 ADAM10   ML     ECM Regulators                 204520.          143690.
##   9 ADAMTSL1 MF     ECM Regulators                  92176.           54265.
##  10 ADAMTSL1 ML     ECM Regulators                 114634.           87425.
## # ℹ 490 more rows

# Save the summarized data to a CSV file
write.csv(gene_stats, file = "CMedulla_Summary.csv", row.names = FALSE)
```
