## Supplementary material for "Developing and characterising decellularized extracellular matrix hydrogels to bio-fabricate female reproductive tissues": Suppl. FIle 2

### Supplementary file 1

**Suppl. Table 1.** List of matrisome proteins differentially abundant between follicular

| Tissue | Gene name | Matrisome division | Matrisome category | Follicular *vs* Luteal phase | Adj. P-value |
|---|---|---|---|---|---|
| **Endometrium** | COL2A1 | Core matrisome | Collagens | down regulated | 0.0489 |
| | COL4A1 | Core matrisome | Collagens | up regulated | 0.0497 |
| | COL4A2 | Core matrisome | Collagens | up regulated | 0.0243 |
| | FBN1 | Core matrisome | ECM Glycoproteins | up regulated | 0.049 |
| | KNG2 | Matrisome associated | ECM Regulators | up regulated | 0.0552 |
| | MMP3 | Matrisome associated | ECM Regulators | up regulated | 0.0532 |
| | IGF2 | Matrisome associated | ECM Regulators | up regulated | 0.0136 |
| **Oviduct** | COL3A1 | Core matrisome | Collagens | up regulated | 0.0081 |
| | HAPLN1 | Core matrisome | Proteoglycans | up regulated | 0.0144 |
| | THBS1 | Core matrisome | ECM Glycoproteins | up regulated | 0.0036 |
| | CTSS | Matrisome associated | ECM Regulators | up regulated | 0.0374 |
| **Cortex** | F13B | Matrisome associated | ECM Regulators | up regulated | 0.036 |
| | ANGPTL2 | Matrisome associated | Secreted factors | up regulated | 0.0702 |
| **Medulla** | IGFALS | Core matrisome | ECM Glycoproteins | down regulated | 0.0224 |
| | LAMC3 | Core matrisome | ECM Glycoproteins | up regulated | 0.0006 |
| | MXRA5 | Core matrisome | ECM Glycoproteins | down regulated | 0.0001 |
| | PCOLCE | Core matrisome | ECM Glycoproteins | down regulated | 0.0129 |
| | FETUB | Matrisome associated | ECM Regulators | up regulated | 0.0527 |
| | SERPINF2 | Matrisome associated | ECM Regulators | up regulated | 0.0279 |
| | PLXNB1 | Matrisome associated | ECM-affiliated | up regulated | 0.0187 |

and luteal estrus phases for each tissue investigated

**Suppl. Table 2.** Comparison of gelation kinetic parameters for the different tissues hydrogels supplemented with 0, 0.5 or 1% of alginate

| | $T_{lag}$ (min)* | $T_{1/2}$ (min)** | $T_1$ (min)*** |
|---|---|---|---|
| Cortex 0% | $2.7^{a,c}$ | $13.1^{a,b,c}$ | 40.7 |
| Cortex 0.5% | $2.0^{a,c}$ | $23.6^{b}$ | 52.7 |
| Cortex 1% | $2.7^{a,c}$ | $6^{c}$ | 32 |
| Endometrium 0% | $12.0^{a,b}$ | $21.7^{a,b,c}$ | 46 |
| Endometrium 0.5% | $13.0^{b,d}$ | $22.7^{a,b}$ | 45 |
| Endometrium 1% | $11.0^{b}$ | $21.8^{a,b}$ | 53 |
| Medulla 0% | $2.0^{a,c}$ | $10.1^{a,c}$ | 32 |
| Medulla 0.5% | $2.0^{a,c}$ | $14.9^{a,b,c}$ | 55 |
| Medulla 1% | $2.0^{a,c}$ | $12.5^{a,b,c}$ | 55 |
| Oviduct 0% | $2.0^{a,c}$ | $14.1^{a,c}$ | 44 |
| Oviduct 0.5% | $6.0^{a,b,c}$ | $17.4^{a,b,c}$ | 43 |
| Oviduct 1% | $6.0^{a,b,c}$ | $16.5^{a,b,c}$ | 40 |

Data represents mean, letters indicates statistical differences for each column separately ($p<0.05$)

*point at which the linear section of the gelation curve intersects with 0% absorbance

**moment when absorbance reaches 50%

***time to complete gelation (when absorbance hits 100%)

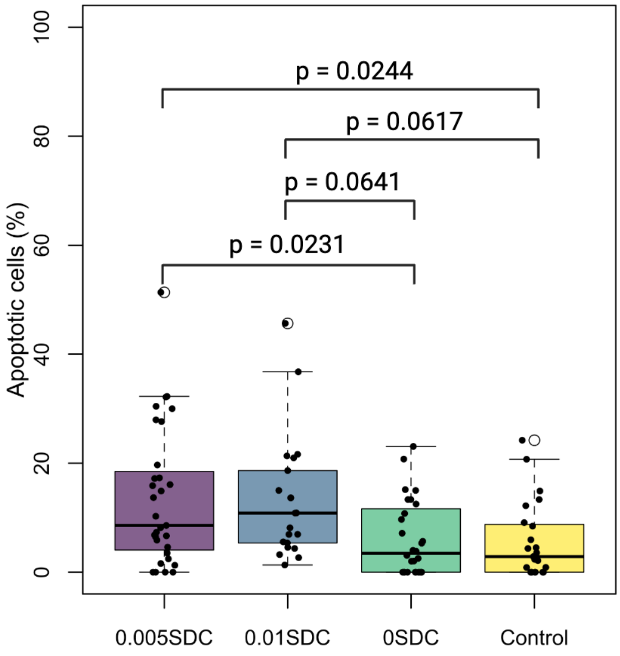


**Suppl. Figure 1.** SDC toxicity in embryos. Percentage of apoptotic cells in blastocysts cultured with or without dECM with varying SDC concentrations (0, 0.005, and 0.01%).

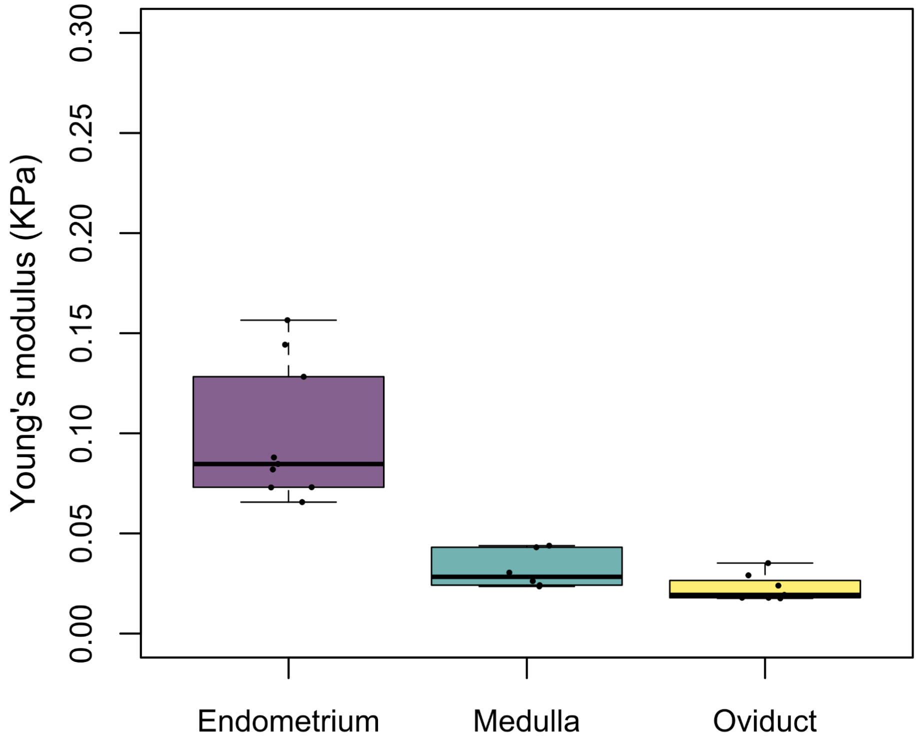


**Suppl. Figure 2.** Young´s modulus of EndoECM, MedECM and OvaECM (10 mg/ml) not-enriched with alginate.

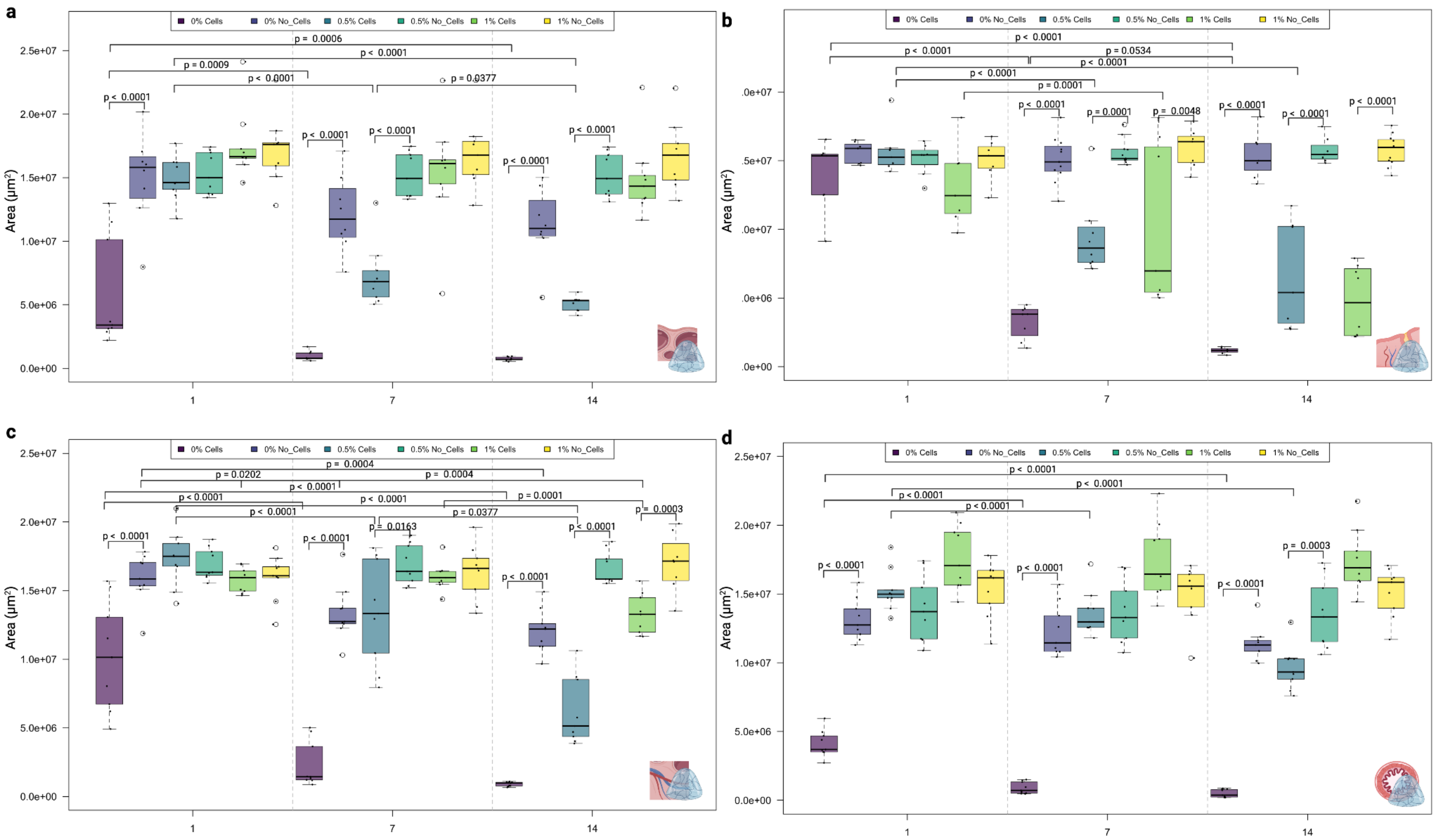


**Suppl. Figure 3.** Hydrogel stability over time with and without cells. Area measurement of CorECM (a), EndoECM (b), MedECM (c), and OviECM (d) enriched with 1 % alginate, 0.5 % alginate, and non-alginate, embedded with or without ovarian cortex, endometrial, ovarian medulla, and oviductal stroma cells, respectively, over a 14 days period.
